# Lipidomic analysis reveals differences in the extent of remyelination in the brain and spinal cord

**DOI:** 10.1101/2023.07.24.550351

**Authors:** Nishama De Silva Mohotti, Hiroko Kobayashi, Jenna M. Williams, Rashmi Binjawadagi, Michel P. Evertsen, Ethan G. Christ, Meredith D. Hartley

**Affiliations:** Department of Chemistry, University of Kansas, 2030 Becker Drive, Lawrence, Kansas 66047, United States

**Keywords:** Lipidomics, myelin, demyelination, remyelination

## Abstract

During demyelination, lipid-rich myelin debris is released in the central nervous system (CNS) and must be phagocytosed and processed before new myelin can form. Although myelin comprises over 70% lipids, relatively little is known about how the CNS lipidome changes during demyelination and remyelination. In this study, we obtained a longitudinal lipidomic profile of the brain, spinal cord, and serum using a genetic mouse model of demyelination, known as *Plp1*-iCKO-*Myrf* mice. This model has distinct phases of demyelination and remyelination over the course of 24 weeks, in which loss of motor function peaks during demyelination. Using principal component analysis (PCA) and volcano plots, we have demonstrated that the brain and spinal cord have different remyelination capabilities and that this is reflected in different lipidomic profiles over time. We observed that plasmalogens (ether-linked phosphatidylserine and ether-linked phosphatidylcholine) were elevated specifically during the early stages of active demyelination. In addition, we identified lipids in the brain that were altered when mice were treated with a remyelinating drug, which may be CNS biomarkers of remyelination. The results of this study provide new insights into how the lipidome changes in response to demyelination, which will enable future studies to elucidate mechanisms of lipid regulation during demyelination and remyelination.

## INTRODUCTION

Most lipids in the central nervous system (CNS) reside in myelin membranes in the brain and spinal cord.^1^ Myelin sheaths surround and insulate axons to facilitate efficient, saltatory signal transduction. Myelin is formed by the extension of an oligodendrocyte lipid bilayer, which pinches off and wraps around the axon to form a multilamellar myelin sheath. Demyelination occurs when an oligodendrocyte or myelin is damaged and the myelin sheath degrades, which leads to a release of myelin lipid debris into the CNS parenchyma. In order for remyelination to occur, the damaged myelin must be removed from the lesion site through microglia phagocytosis.^2^

Lipidomics is a powerful tool that enables the profiling of lipid dynamics in demyelination, which has the potential to provide insights into how lipids are regulated during myelin damage and repair. Previous studies performed in gliotoxin models of demyelination, such as the cuprizone model, have revealed changes in the brain lipidome during demyelination.^3–5^ However, the interpretation of these previous studies is limited by fast remyelination, which can occur simultaneously with demyelination in the cuprizone model.^6^ Rapid remyelination also makes it challenging to obtain longitudinal lipidomics data from distinct phases of demyelination and remyelination.

The goal of this study is to obtain longitudinal data on how the lipidomes of brain, spinal cord, and serum change during demyelination and remyelination. To achieve, this goal we have used a genetic mouse model of demyelination (*Plp1*-iCKO-*Myrf*), in which widespread demyelination occurs in both the brain and the spinal cord and is followed by a discrete phase of remyelination.^7, 8^ The *Plp1*-iCKO-*Myrf* model enables us to directly compare changes in the multiple CNS tissues at the same time. Demyelination in the *Plp1*-iCKO-*Myrf* mouse strain is based on the deletion of myelin regulation factor (*Myrf*), which is a transcription factor required for the expression of myelin genes and the maintenance of a healthy myelin sheath.^9^ Loss of *Myrf* results in the gradual degradation of the myelin sheath over time. *Myrf* is not deleted in oligodendrocyte precursor cells (OPCs), and the OPCs are available to proliferate, differentiate into oligodendrocytes, and remyelinate the CNS.

In this study, we used the *Plp1*-iCKO-*Myrf* mouse strain to perform unbiased lipidomics to profile brain, spinal cord, and serum during demyelination and remyelination. We also performed lipidomics on *Plp1*-iCKO-*Myrf* mice treated with Sob-AM2, which is a thyroid hormone agonist^10–12^ that has improved remyelination and motor recovery in *Plp1*-iCKO-*Myrf* mice.^7^ The lipidomics data have provided several new insights into CNS lipid dynamics during myelin damage and repair. The brain lipidome showed substantial changes that were the greatest near peak demyelination and showed partial normalization with remyelination. In contrast, the spinal cord lipidome was greatly altered at all timepoints including those associated with motor recovery and remyelination in the brain. The differences between the brain and spinal cord lipidomes correlated with histological analysis revealing that the spinal cord has poor remyelination in this model relative to the brain. In addition, we observed that demyelination phases were associated with both decreased and increased lipid species, but that later timepoints associated with remyelination did not have any elevated lipid levels. This study represents the first longitudinal profile detailing how CNS lipids change during the time course of myelin damage and repair in multiple tissues simultaneously.

## METHODS

### Animal husbandry

Male and female C57BL6/J *Plp1*-iCKO-*Myrf* mice (*Myrf fl/fl*; *Plp1-CreERT*)^8^ were bred at the University of Kansas by crossing *Myrf(fl/fl)*^9^ mice with *Plp1-CreERT*^13^ mice. All mice were housed in a climate-controlled room (24 ± 1 °C) with a constant 12 h light / dark cycle (12h on, 12h off) with food and water *ad libitum*. At 8 weeks of age, mice were injected intraperitoneally with 2 mg of tamoxifen (100 μl of 20 mg/ml tamoxifen) in corn oil daily for 5 consecutive days. Both *Cre* negative mice, which do not lose *Myrf* and undergo demyelination, and *Cre* positive mice, which do undergo demyelination, were used in all experiments. Mice were randomly assigned into the control or Sob-AM2 treatment groups. The control group received control chow (Envigo Teklad 2016 diet) and the Sob-AM2 treatment group received chow containing 420 µg/kg chow of Sob-AM2 (nominal daily dose of 84 µg/kg body weight) starting 2 weeks after tamoxifen injections. Mice on control chow were euthanized at 4 timepoints: 6, 12, 18, and 24 weeks post-tamoxifen injections. Mice treated with Sob-AM2 were euthanized at 12 and 18 weeks post-tamoxifen. Euthanasia was performed by carbon dioxide inhalation and cervical dislocation, and serum, brain, and spinal cord tissues were collected immediately. Mice for histological analysis were euthanized by intracardiac perfusion with HBSS (Hanks’ Balanced Salt Solution) buffer followed by 4% paraformaldehyde in HBSS. All experiments were approved by the Institutional Animal Care and Use Committee at the University of Kansas.

### Rotarod analysis

All mice were trained on the rotarod 2–4 days before the first experiment. Training consisted of three 2-minute trials at 8 rpm with 15-minute rest intervals between trials. Motor function testing was performed at 6, 12, 18, and 24 weeks post-tamoxifen using a Rotarod (IITC Life Science). Testing was performed with three 5-minute trials with the following program: 0:00-4:00, ramp from 8-40 rpm; 4:00-5:00 held at 40 rpm. The mice were allowed to return to their cage for at least 15 minutes between trials. If mice stayed on the rod for the full 5 minutes, then 300 seconds was recorded as the latency. If mice held onto the rod and rotated around once, then that was counted as a fall. Time to fall (latency) was recorded for the three trials and averaged for each mouse at each timepoint.

### Black gold II staining

Free floating sections (40 µm) of brain and lumbar spinal cord were obtained using a Vibratome (Leica VT 1200S). Sections were mounted on positively charged slides and dried overnight at room temperature. Black Gold staining was performed according to the manufacturer’s protocol with the following modifications. Mounted sections were rehydrated in de-ionized water for 2 minutes. Then the slides were transferred into a 0.3% Black Gold ® II (Histo-Chem) solution for 3-5 hours until desired myelin impregnation was observed. Then the slides were rinsed with de-ionized water and transferred into a 1% sodium thiosulfate solution for 3 minutes. Slides were again rinsed with de-ionized water three times. Sections were dehydrated for 30 seconds in each ethanol solution (50%, 75%, 85%, 95%, 100%). Finally, the slides were immersed in xylene for 1-2 minutes and secured with coverslips using Cytoseal XYL mounting media (Thermo Fisher). Mounted slides were heated on a slide warmer for 2-3 hours at 60 °C. Sections were imaged using a slide reader (BioTek Cytation 5) at x4 or x20 magnification. Multiple images were taken and stitched together to obtain the whole image.

### Lipid extraction

A modified Bligh-Dyer protocol was used to extract lipids from the brain and spinal cord tissue.^14^ Brain homogenates (either 300 mg/ml or 50 mg/ml) were prepared with ice cold water using a Bead Mill homogenizer (Bead Ruptor Elite, Omni international, USA). Spinal cord homogenates were prepared at 65 mg/ml in cold water. Immediately after homogenization, the brain homogenates were diluted with cold water (150 µL of 300 mg/ml brain homogenate was mixed with 800 µl of cold water. For the 50 mg/ml brain homogenates, 1 mL of homogenate was used directly. Spinal cord homogenates were diluted 10-fold (100 μl of the spinal cord homogenates with 900 μl of cold water). For all samples, 10 μl of the diluted homogenate was removed and stored at −80°C for protein quantification using a BCA assay. The diluted tissue homogenates were combined with a mixture of chloroform (containing 0.01% butylated hydroxytoluene, BHT): methanol: water (3:2:1) in glass tubes. After shaking and vortexing thoroughly, the mixture was centrifuged (Sorvall ST 40R Centrifuge, Thermo Fisher Scientific) at 1300 rpm for 10 minutes. The lower layer was carefully removed and saved in a glass tube. The remaining top layer was further extracted twice with 1.25 ml chloroform with 0.01% BHT; the lower layers were carefully removed and combined. The combined lower layer was then washed with 300 μl of 1 M KCl followed by 300 μl of water and vacuum dried completely (Savant SpeedVac SPD130DLX vacuum concentrator, Thermo Fisher Scientific, USA) to obtain the dried lipid extract.

Serum (3 μl) was added directly to a vial containing 1.2 mL of chloroform: methanol: 300 mM ammonium acetate in water (300:665:35). The contents were mixed thoroughly, and the vials were centrifuged for 5 min at low speed in a clinical centrifuge to pellet proteins before submitting samples to the mass spectrometer.

### Mass spectrometry analysis

Quantification of phospholipids was performed by direct infusion triple quadrupole mass spectrometry on a Sciex 4000 QTrap at the Kansas Lipidomic Research Center at Kansas State University. Briefly, dried lipid samples (brain and spinal cord) were dissolved in 1 mL of chloroform and serum samples were used directly as prepared above. Aliquots were mixed with internal standards and solvents, and analysis was carried out as previously described.^15^ Internal standards are indicated in Table S1. Lipid measurements were performed using the acquisition and data processing parameters indicated in Table S1. Internal standards were from the same class as the analytes, and corrections for differences in response between the internal standards employed and SPLASH Lipidomix (Avanti Polar Lipids) were applied. For sphingomyelin (SM) and phosphatidylcholine (PC), standards were used and a correction for the response of a standard SM vs standard PCs was applied. No additional corrections for variation in response of the instrument to individual analytes vs their standards were applied. Thus, data are reported as normalized mass spectral intensity where a value of 1 indicates the intensity of 1 nmol of internal standard (Tables S2-S4).

The mass spectrometry analysis of the samples from the 6, 12, and 18 week groups (including control and Sob-AM2 treatment groups) were analyzed separately from the 24 week group. Brain, spinal cord, and serum were analyzed on separate days. To account for differences in the data due to the day of analysis, samples from the 12 and 18 week groups were analyzed again during the 24 week analysis and the levels were used to normalize the 24 week data. Serum samples from 24 weeks were not analyzed.

### Data analysis

Data were analyzed both as individual lipid species and as total lipids, in which the different fatty acyl derivatives were summed (Tables S2-S4). Quality controls were prepared by pooling a portion of all samples for a given analytical run. Then 5 quality control samples were analyzed at intervals during the sample run. The coefficient of variation (CV) of the quality control for each lipid species was determined (Tables S2-S4). Any lipid species with a CV higher than 0.3 was not included in the subsequent data analysis. Similarly, all lipid totals with a CV higher than 0.3 were removed from further analysis except for total lysophosphatidylethanolamine (LPE), which had sufficient CV in the brain, but not the spinal cord. Since the CV of total LPE was near 0.3 in the spinal cord (0.41 during weeks 6, 12, and 18, 0.38 for week 24), we included the data for comparison to the brain.

The data from 5 mice (for all tissues) were excluded from data analysis due to errors in genotyping. One spinal cord sample was excluded from data analysis due to a mistake during the lipid extraction procedure.

The total lipid levels for *Cre* negative and *Cre* positive mice were first compared at all 4 timepoints using the Holm-Šídák correction method for multiple t-tests. In addition, *Cre* negative mice on control chow were compared to *Cre* negative mice treated with SobAM2 chow and *Cre* positive mice on control chow were compared to *Cre* positive mice treated with SobAM2 chow using the Holm-Šídák correction method for multiple t-tests. The results of the statistical analyses including P values are contained in Tables S5-S7.

### Principal Component Analysis (PCA)

Using the total lipid data, principal component analysis (PCA) was performed for each week comparing *Cre* negative and *Cre* positive samples to determine how the samples cluster. PCA was also performed on all week 6-18 brain samples grouped by phenotype: healthy (all *Cre* negative), demyelination (week 6 and 12 *Cre* positive), and remyelination (week 18 *Cre* positive). To analyze the effects of Sob-AM2 treatment, PCA was performed with datasets that included (1) *Cre* negative mice administered control chow and *Cre* negative mice administered SobAM2 chow at week 12 or 18, and (2) *Cre* positive mice administered control chow and *Cre* positive mice administered SobAM2 chow at week 12 or 18. In addition, PCA was also performed on all week 12 samples (both genotypes and both treatment conditions) and all week 18 samples (both genotypes and both treatment conditions). The analyses were performed using R Statistical Software (v4.2.3) and the factoextra package (v1.0.7).^16, 17^ The code is available in the Supplementary Information.

### Volcano plot analysis

The individual lipid species were analyzed in the following groups (*Cre* negative versus *Cre* positive for all timepoints, *Cre* negative mice on control chow versus *Cre* negative mice on SobAM2 chow at week 12 or 18, and *Cre* positive mice on control chow and *Cre* positive mice on SobAM2 chow at week 12 or 18. The data was log_2_ transformed prior to analysis and lipid species that included values of “0” were dropped from the data at each timepoint. The fold change (FC) and P values from individual t tests were determined for all comparisons and volcano plots were prepared plotting -Log_10_(P-value) on the y-axis versus Log_2_(FC) on the x-axis. The Log_2_(FC) was set at 0.5 and the p-value rejection threshold was determined for each timepoint using a permutation-based false discovery proportion estimate method.^18^ The rejection thresholds were determined through the R package permFDP (v0.1.0)^19^ and are listed in Tables S8-S10. The volcano plots were generated using the R package enhanced Volcano (v1.13.2).^20^ The individual FC and P-values for all comparisons are included in Tables S8-S10 and the lipids that meet the FC and P-value thresholds are listed in Tables S11-S13.

## RESULTS AND DISCUSSION

### Remyelination in the spinal cord is impaired relative to remyelination in the brain

*Plp1*-iCKO-*Myrf* mice were allowed to undergo normal developmental myelination prior to the induction of demyelination at 8 weeks of age. Demyelination was initiated by 5 daily doses of tamoxifen, which deleted the *Myrf* gene by conditionally activating *Cre* recombinase in mature oligodendrocytes (*Plp1* promoter),^8^ but not in OPCs. In the model, demyelination in the brain begins around week 4 post-tamoxifen and typically peaks at week 10 post-tamoxifen. Demyelination correlates with increasing loss of motor function, and from 10-14 weeks post-tamoxifen, the mice show maximum motor loss with significant hindlimb weakness. Remyelination in the brain starts around week 12 post-tamoxifen with steady improvements in motor ability from weeks 14-20 post-tamoxifen.^7^

Motor ability was assessed using an accelerating rotarod motor test (Fig. 1A) at weeks 6, 12, 18, and 24 post-tamoxifen. Demyelination decreased motor function as mice had reduced latency, or time spent on the rotarod, at all timepoints. The mice experienced the greatest loss of motor function at week 12, which corresponds to the peak demyelination observed in the brain. Only partial recovery of motor function was observed by week 24. These results are consistent with previous observations in this model.^7^

**Figure 1.**
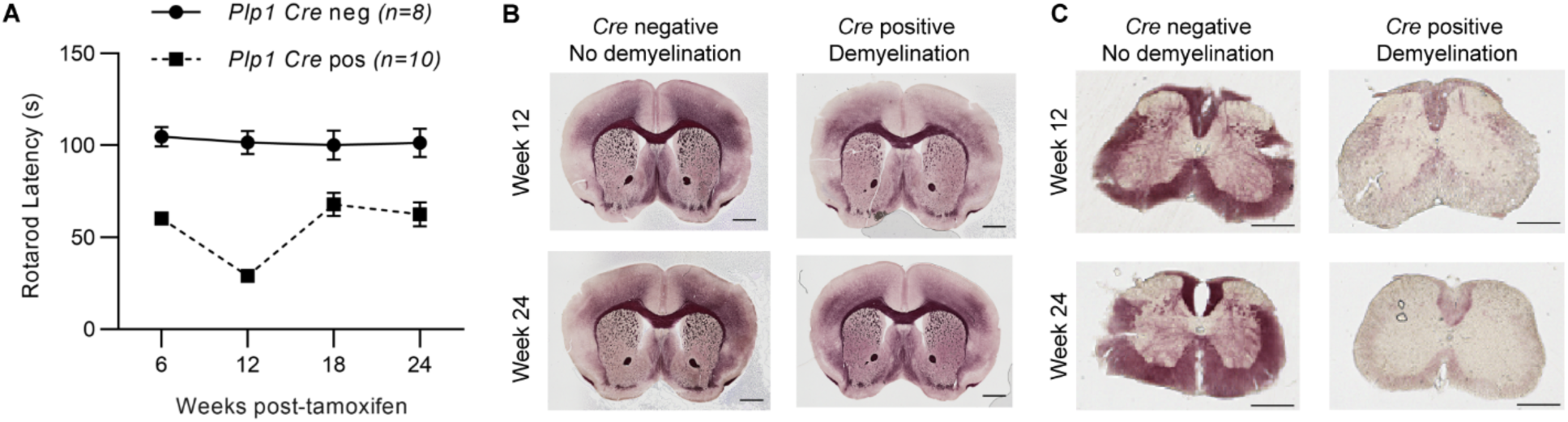
Demyelination in *Plp1*-iCKO-*Myrf* mice is followed by remyelination in the brain, and impaired remyelination in the spinal cord. (A) Motor dysfunction was assessed by rotarod testing. Rotarod latency (time to fall) was recorded at weeks 6, 12, 18, and 24 post-tamoxifen. Data is plotted as mean ± SEM. (B and C) Brain and spinal cord tissue sections were stained with BlackGold, a myelin lipid stain. Sections were imaged at weeks 12 and 24 post-tamoxifen in *Cre* negative (healthy) mice and *Cre* positive (demyelination) mice. Scale bars in B represent 1 mm and in C represent 0.5 mm.

Previous studies focused primarily on the time course of demyelination and remyelination in the brain. To determine the effects of demyelination in the spinal cord, myelin was stained in both brain and spinal cord tissue sections using BlackGold from mice euthanized at peak motor disability (week 12) and motor recovery (week 24). The brain showed demyelination at week 12 followed by remyelination at week 24 (Fig. 1B).^7^ In striking contrast to the brain, spinal cord showed similar levels of demyelination at week 12 and week 24 (Fig. 1C) suggesting that remyelination is impaired in the spinal cord relative to the brain. The contrast between the brain and spinal cord provides a unique opportunity to compare the lipidomes of robust remyelination in the brain with impaired remyelination in the spinal cord.

### Brain and spinal cord have unique lipidomic signatures during demyelination and remyelination

Brains and spinal cords were isolated from both *Plp1*-iCKO-*Myrf* mice at 6, 12, 18, and 24 weeks post-tamoxifen in two genotypes: *Cre* negative (healthy, no demyelination) and *Cre* positive (demyelination). These timepoints represented active demyelination (6 weeks), peak demyelination (12 weeks), and remyelination (18 and 24 weeks). The tissues were homogenized in water and extracted using a modified Bligh-Dyer protocol.^14, 15^ The tissues were analyzed by a lipidomics platform that included 12 major classes of lipids (Tables S2-S4). For each class, individual lipids were measured (ranging from 14:0 – 22:0 for monoacylated lipids and 28:1 – 44:2 for diacylated lipids). Quality controls prepared from pooled samples were measured five times, and any individual lipids that had high coefficient of variations in the quality controls (> 0.3) were excluded from subsequent analyses.

The individual lipids for each class were summed and the totals were plotted for the four timepoints to visualize how the overall lipidome of the brain (Fig. 2) and spinal cord (Fig. 3) change with demyelination and remyelination. The lipid levels for each sample are in Tables S2 and S3, and statistical analyses are available in Tables S5 and S6.

**Figure 2.**
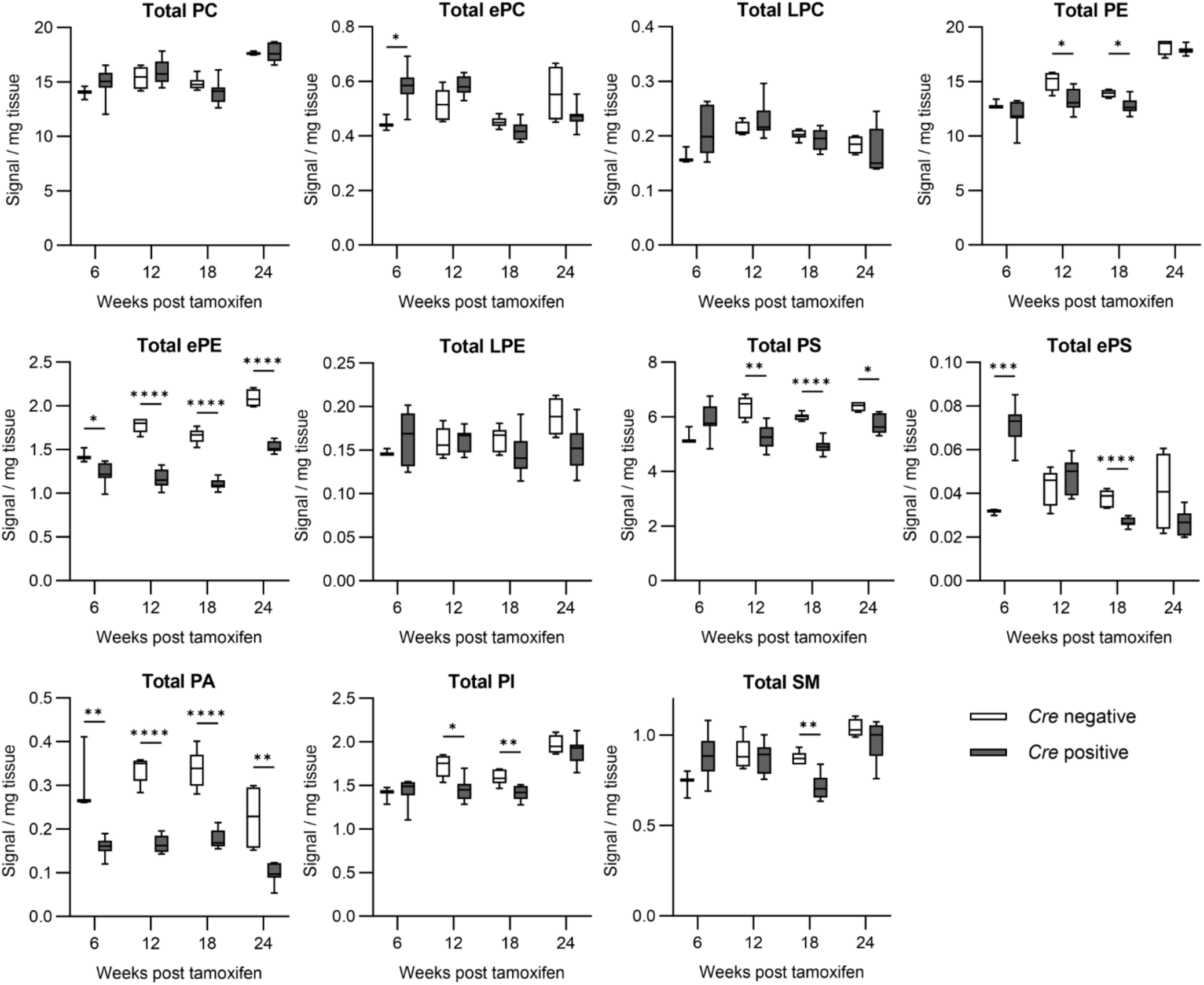
Total lipid levels in brain during demyelination and remyelination. The total lipid levels for 11 major classes are shown. All individual lipids were measured for each class and the values were summed for this figure. The data are plotted as box and whisker plots with the bars representing the minimum and maximum values. Statistical analysis was performed with multiple t-tests comparing *Cre* negative and *Cre* positive at each timepoint using a Holm-Šídák correction for multiple comparisons. The sample numbers are the following (Cre negative: 6 weeks n = 3, 12 weeks n = 5, 18 weeks n = 6, 24 weeks n = 4; Cre positive: 6 weeks n = 8, 12 weeks n = 8, 18 weeks n = 8, 24 weeks n = 7). Abbreviations: PC, phosphatidylcholine; ePC, ether-linked PC; LPC, lyso-PC; PE, phosphatidylethanolamine; ePE, ether-linked PE; LPE, lyso-PE; PS, phosphatidylserine; ePS, ether-linked PS; PA, phosphatidic acid; PI, phosphatidylinositol; SM, sphingomyelin.

**Figure 3.**
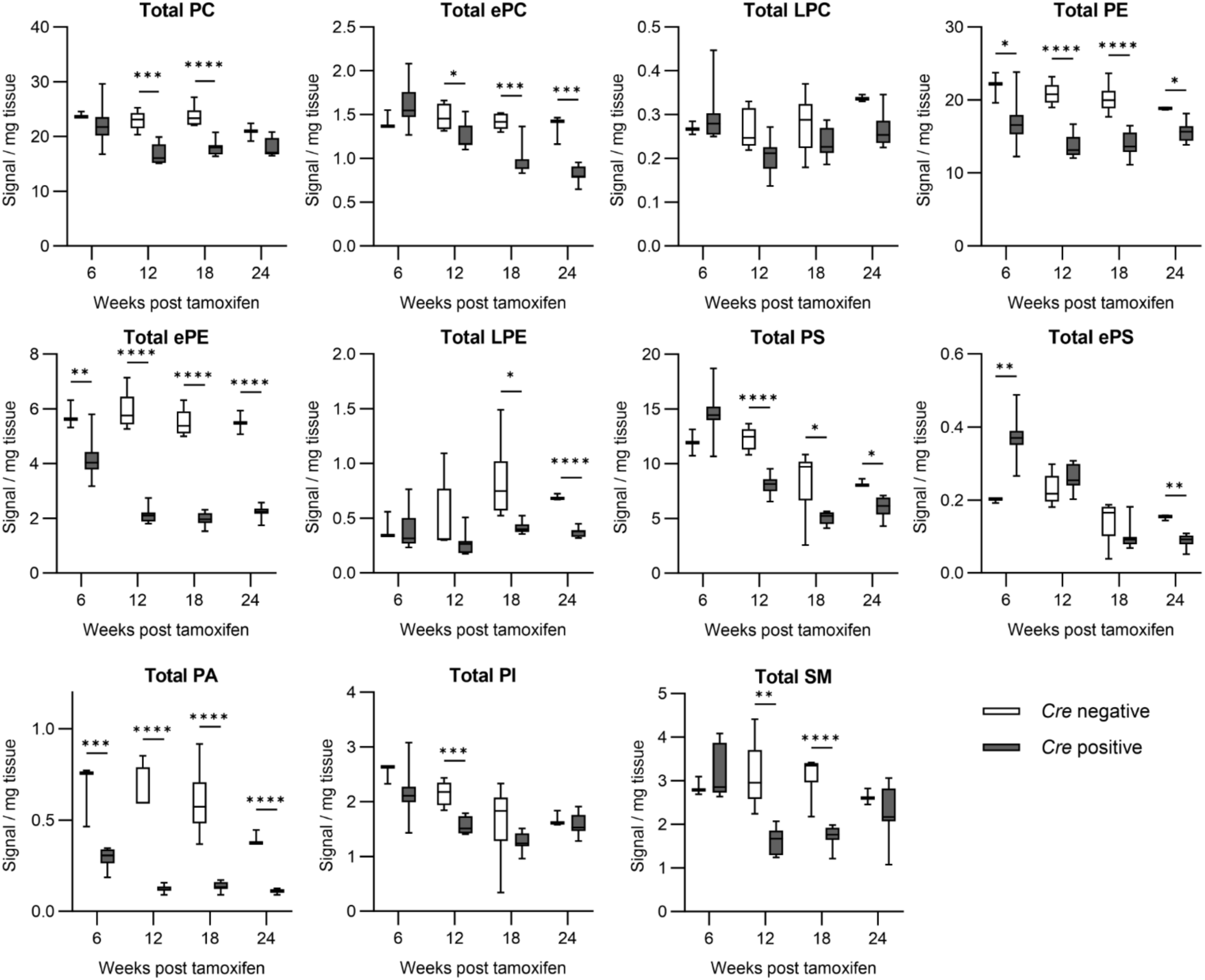
Total lipid levels in spinal cord tissue after demyelination. The total lipid levels for 11 major classes are shown. All individual lipids were measured for each class and the values were summed for this figure. The data are plotted as box and whisker plots with the bars representing the minimum and maximum values. Statistical analysis was performed with multiple t-tests comparing *Cre* negative and *Cre* positive at each timepoint using a Holm-Šídák correction for multiple comparisons. The sample numbers are the following (Cre negative: 6 weeks n = 3, 12 weeks n = 5, 18 weeks n = 6, 24 weeks n = 3; Cre positive: 6 weeks n = 8, 12 weeks n = 8, 18 weeks n = 9, 24 weeks n = 7). Abbreviations: PC, phosphatidylcholine; ePC, ether-linked PC; LPC, lyso-PC; PE, phosphatidylethanolamine; ePE, ether-linked PE; LPE, lyso-PE; PS, phosphatidylserine; ePS, ether-linked PS; PA, phosphatidic acid; PI, phosphatidylinositol; SM, sphingomyelin.

Analysis of Figs. 2 and 3 revealed four major trends that are similar for both brain and spinal cord. First, ether-linked phosphatidylethanolamine (ePE), phosphatidylserine (PS), and phosphatidic acid (PA) were reduced at most or all timepoints. This suggests that demyelination causes chronic reductions in these lipid classes that are not restored even with robust remyelination in the brain. Second, phosphatidylethanolamine (PE), phosphatidylinositol (PI), and sphingomyelin (SM) decreased with demyelination (weeks 12 and 18) and increased to normal levels at later timepoints (week 24), which suggests a role for these lipids in remyelination. Third, phosphatidylcholine (PC), lysophosphatidylcholine (LPC), and lyso-PE (LPE) were mostly unaffected by demyelination. This may imply that these lipids are less abundant in myelin relative to other lipid classes. Finally, ether-linked PC (ePC) in the brain and ether-linked PS (ePS) in both the brain and spinal cord were elevated during active demyelination (week 6), but had normalized or decreased levels at later timepoints. The role of ether-linked lipids, or plasmalogens, in neurological disease is still unclear.^21^ Plasmalogens have been implicated both as mediators of inflammation and as potential antioxidants.^22^ Further studies will be needed to understand how ePC and ePS are modulating demyelination and remyelination in the *Plp1*-iCKO-*Myrf* model.

In addition to identifying trends in lipid classes, the total lipid analysis enabled us to directly define how the lipidomes of brain and spinal cord respond to demyelination (Figs. 2 and 3). Myelin lipids comprise a larger portion of the spinal cord, which is consistent with our observations that the spinal cord showed more dramatic fold changes in almost every lipid class as compared to the brain. Histological analysis revealed that the brain showed robust remyelination (Fig. 1A) as compared to poor remyelination in the spinal cord (Fig. 1B). The impaired remyelination in the spinal cord is also clear from the lipidomic results. In the brain, 8 of 11 lipid classes are at normal levels by week 24, whereas in the spinal cord, only 4 of the 11 lipid classes are at normal levels at week 24.

### Principal component analysis (PCA) discriminates between the lipidomes of healthy, demyelinated, and remyelinated CNS tissues

The total lipid data in Figs. 2 and 3 was further analyzed by PCA to determine if tissue that has undergone demyelination can be distinguished from healthy tissue based on changes to the lipidome. Unsupervised PCA was performed to classify the data at each timepoint in the brain (Fig. 4A-D) and spinal cord (Fig. 4E-H). The PCA score plots were annotated with 95% confidence clusters to group the samples. The healthy (*Cre* negative) and demyelination (*Cre* positive) groups clustered distinctly at all timepoints in both brain and spinal cord. The PCA score plot of brain at week 24 showed the least discrimination between the two groups, which is consistent with the robust remyelination and recovery of most lipid classes by this timepoint in the brain. PCA biplots shown in Figs. S1 and S2 further illustrate lipid class trends, variability, and contribution to each principal component.

**Figure 4.**
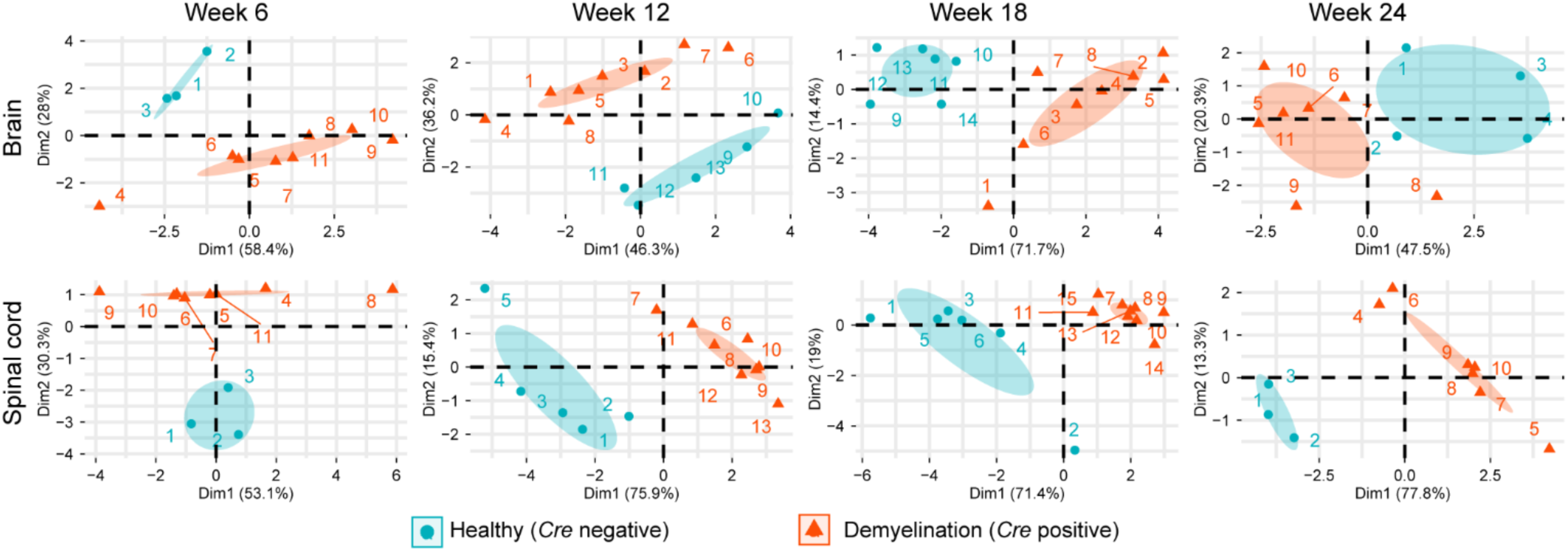
Principal component analysis (PCA) discriminated between healthy and demyelinated CNS lipidomes. PCA was performed on each timepoint (6, 12, 18, and 24 weeks post-tamoxifen) in both tissues (brain and spinal cord). The total levels for each lipid class were used in this analysis (Tables S2 and S3). The genotypes cluster distinctly at all timepoints with the brain at week 24 showing the least discrimination.

Additional PCA was performed to determine if demyelination could be discriminated from remyelination in the brain. This analysis was performed only on weeks 6, 12, and 18 brain samples as they were all analyzed by mass spectrometry on the same day, and the brain had more robust remyelination than the spinal cord. The samples were divided into the following three groups based on phenotype: 1. Healthy (all *Cre* negative, weeks 6, 12, and 18), 2. Demyelination (*Cre* positive, weeks 6 and 12), and 3. Remyelination (*Cre* positive, week 18). PCA analysis revealed that the three groups clustered separately with clear contrast between healthy, demyelinating, and remyelinating lipidomes (Fig. 5). Interestingly, the remyelinating lipidome grouped halfway between the normal and demyelinated lipidome groups.

**Figure 5.**
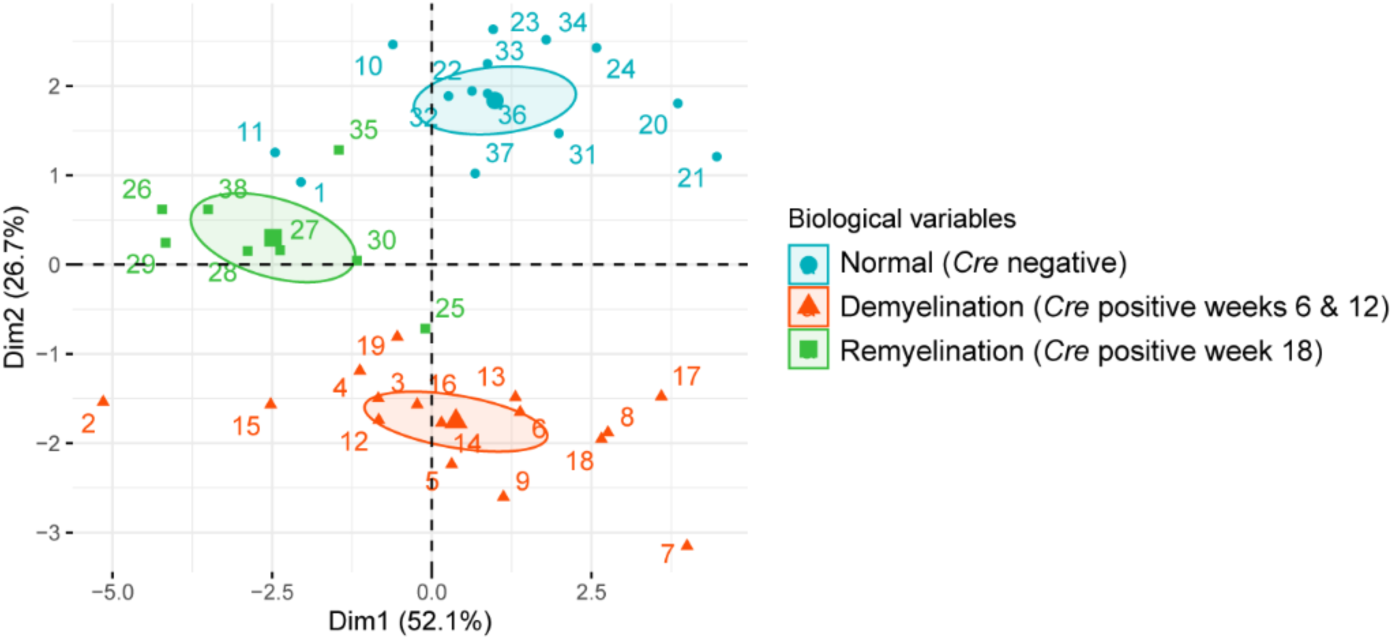
PCA clustered healthy, demyelinating, and remyelinating lipidomes separately. Brain samples were grouped by phenotype: 1. Normal (all *Cre* negative, weeks 6, 12, and 18), 2. Demyelination (*Cre* positive, weeks 6 and 12), and 3. Remyelination (*Cre* positive, week 18). The total levels for each lipid class were used in this analysis (Tables S2).

To verify that the PCA clustering was not an artifact of mass spectrometry sample order, the same set of samples was grouped into 3 sets based on sample order during the analysis. As shown in Fig. S3, PCA did not discriminate between the three groups and there was greater overlap. These results support the assertion that differences observed between phenotypes in Figs. 4-5 are based on biological differences and not based on mass spectrometry sample order.

### Volcano plot analysis suggests that demyelination is associated with elevated levels of unsaturated lipids

To identify changes to individual lipid species at each timepoint, we prepared volcano plots showing all lipids that met the coefficient of variation cut-off. The Log_2_ fold change was set at 0.5 and the -Log_10_ P-value cut-off thresholds (Tables S8-S9) were determined using a permutation-based false discovery proportion estimation method.^18^ The individual lipids identified as significantly changing are listed in Tables S11 and S12.

Examining the volcano plots revealed that increased levels of individual lipids were primarily observed at earlier timepoints during demyelination whereas decreased levels were observed at all timepoints (Fig. 6). These trends also supported the observation that the lipidome of the brain showed changes consistent with remyelination, whereas the lipidome of the spinal cord had more chronic changes associated with demyelination. In the brain, lipids with increased levels were only observed in weeks 6 and 12, whereas in the spinal cord, elevated lipids were observed at all timepoints (Table 1). The higher number of increased lipids is consistent with the more persistent demyelination observed in the spinal cord and suggests that the elevated lipid species may be biomarkers associated with active demyelination states. Further support for remyelination in the brain comes from the number of reduced lipids. In the brain, 41 lipids were reduced at peak demyelination (week 12), whereas 79 lipids were reduced in the spinal cord at the same timepoint. By week 24, the brain had only 21 differentially identified lipids, whereas spinal cord still had more than 60.

**Figure 6.**
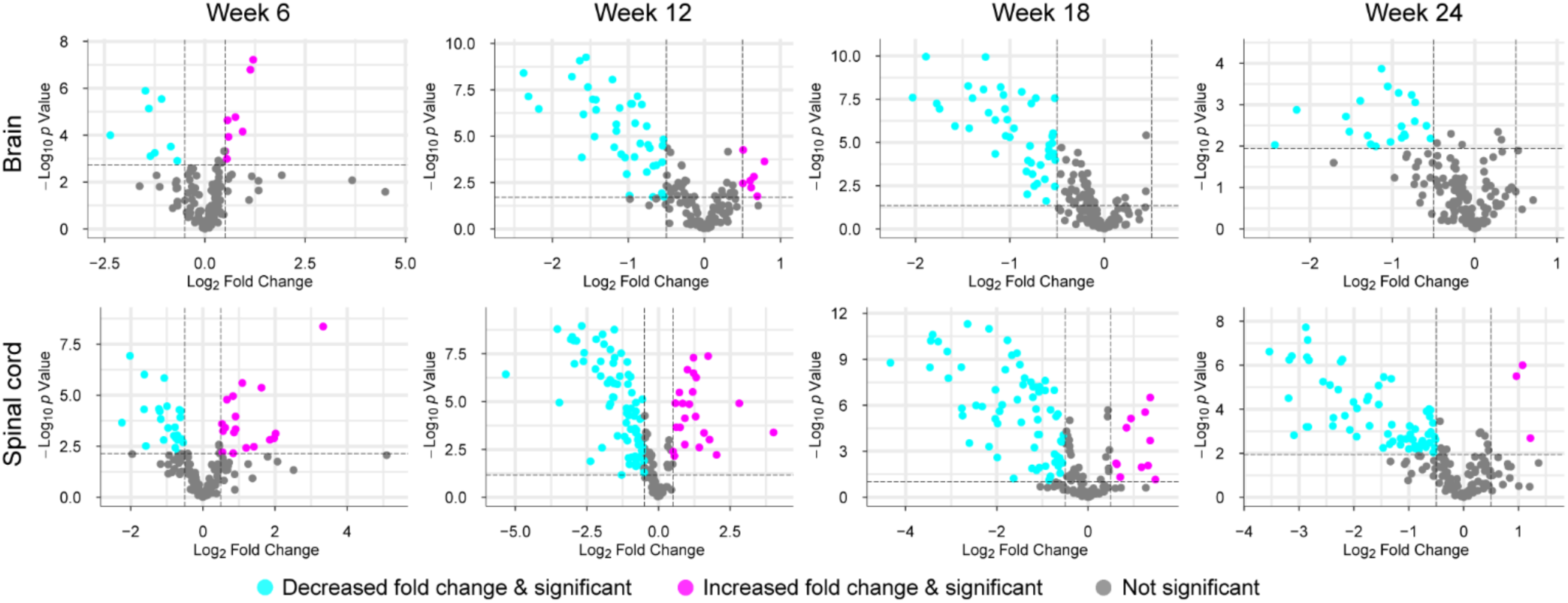
Volcano plot analyses of individual lipid changes during demyelination. All lipids for each comparison that met the coefficient of variation threshold were included in this analysis. The x-axis shows Log_2_(Fold Change), and the threshold was set at 0.5. -Log_10_(P-value) is plotted on the y-axis and the threshold was determined using a permutation-based false discovery estimation method. The individual thresholds are listed in Tables S8-S9.

**Table 1:** Number of individual lipids increasing or decreasing during the demyelination disease course.

| Timepoint | Brain |  | Spinal cord |  |
| --- | --- | --- | --- | --- |
|  | Increased | Decreased | Increased | Decreased |
| 6 weeks | 8 | 8 | 18 | 24 |
| 12 weeks | 7 | 41 | 23 | 79 |
| 18 weeks | 0 | 45 | 11 | 66 |
| 24 weeks | 0 | 21 | 3 | 63 |

To better catalogue which lipids were changing and how they were changing, we prepared Table 2, which contains the fold change for all individual lipids that are identified as significant in the volcano plot analyses and change in both the brain and the spinal cord. The brain has 62 individual lipids that change in at least one of the four timepoints, and the spinal cord has 121 individual lipids that change in at least one of the four timepoints. Of the 62 lipids that change in the brain, 58 also change in the spinal cord dataset, and these are included in Table 2. The high overlap between the identities of the altered lipids between brain and spinal cord suggests that the observed changes are biologically relevant and related to the demyelination phenotype. Fig. S4 contains the fold change data for the lipids that are changed only in the brain or the spinal cord.

**Table 2:** Fold change values for all individual lipids that change significantly both in brain and spinal cord as determined by volcano plot analyses.

| Lipid species | m/z | Brains |  |  |  | Spinal cords |  |  |  |
| --- | --- | --- | --- | --- | --- | --- | --- | --- | --- |
|  |  | Week 6 | Week 12 | Week 18 | Week 24 | Week 6 | Week 12 | Week 18 | Week 24 |
| ePC(34:2) | 744.6 | 1.694 | - | 0.665 | - | 1.442 | 0.639 | 0.342 | 0.259 |
| ePC(36:2) | 772.6 | - | - | 0.665 | - | - | 0.477 | 0.314 | 0.305 |
| ePC(36:3) | 770.6 | - | 0.440 | 0.571 | - | 1.585 | 0.635 | 0.521 | - |
| ePC(36:4) | 768.6 | 1.919 | 1.729 | - | - | 2.663 | 3.317 | 2.575 | - |
| ePC(36:5) | 766.6 | 1.423 | - | - | - | 1.816 | 2.284 | - | - |
| ePC(38:2) | 800.6 | - | 0.598 | 0.449 | 0.492 | 0.622 | 0.246 | 0.147 | 0.192 |
| ePC(38:4) | 796.6 | - | 1.531 | - | - | 1.870 | 2.328 | 0.149 | - |
| ePC(38:5) | 794.6 | - | 1.619 | - | - | 1.788 | 3.006 | 1.532 | - |
| ePC(40:2) | 828.7 | - | 0.493 | 0.653 | 0.540 | - | 0.304 | 0.245 | 0.260 |
| ePE(34:1) | 704.6 | - | 0.608 | 0.682 | - | 0.573 | 0.340 | 0.355 | 0.393 |
| ePE(34:2) | 702.5 | - | 0.512 | 0.515 | 0.556 | - | 0.335 | 0.340 | 0.341 |
| ePE(36:1) | 732.6 | - | 0.432 | 0.449 | 0.604 | 0.585 | 0.258 | 0.222 | 0.249 |
| ePE(36:2) | 730.6 | - | 0.346 | 0.418 | 0.552 | - | 0.267 | 0.294 | 0.364 |
| ePE(36:3) | 728.6 | - | 0.321 | 0.380 | 0.527 | - | 0.349 | 0.433 | 0.594 |
| ePE(38:1) | 760.6 | - | 0.372 | 0.334 | 0.481 | 0.475 | 0.124 | 0.090 | 0.110 |
| ePE(38:2) | 758.6 | - | 0.201 | 0.270 | 0.382 | - | 0.129 | 0.092 | 0.109 |
| ePE(38:3) | 756.6 | 0.421 | 0.192 | 0.245 | 0.338 | 0.513 | 0.163 | 0.118 | 0.142 |
| ePE(40:2) | 786.6 | - | 0.367 | 0.368 | 0.588 | 0.436 | 0.120 | 0.094 | 0.114 |
| ePS(38:1) | 804.6 | - | 0.529 | 0.499 | - | - | 0.460 | 0.427 | 0.384 |
| ePS(38:2) | 802.6 | - | - | 0.568 | - | 1.457 | 0.582 | - | 0.169 |
| LPE(18:1) | 480.3 | - | - | 0.682 | - | - | 0.605 | 0.525 | - |
| PA(34:1) | 692.5 | - | 0.462 | 0.485 | 0.417 | - | 0.165 | 0.182 | 0.205 |
| PA(36:2) | 718.5 | - | 0.332 | 0.428 | 0.348 | 0.334 | 0.137 | 0.253 | 0.408 |
| PA(38:2) | 746.5 | 0.195 | 0.221 | 0.292 | 0.223 | - | 0.025 | 0.050 | 0.086 |
| PA(38:4) | 742.5 | - | 0.689 | 0.679 | - | - | 0.312 | 0.467 | 0.500 |
| PC(36:5) | 780.5 | 1.504 | 1.519 | - | - | 2.295 | 2.452 | 2.382 | - |
| PC(38:1) | 816.6 | - | 0.628 | 0.626 | - | - | 0.310 | 0.271 | 0.300 |
| PC(40:2) | 842.7 | - | 0.485 | 0.491 | - | 0.614 | 0.247 | 0.284 | 0.342 |
| PC(42:2) | 870.7 | - | 0.468 | 0.598 | - | - | 0.319 | 0.373 | 0.540 |
| PC(42:9) | 856.6 | - | - | 0.585 | - | 0.660 | 0.567 | 0.363 | - |
| PE(36:1) | 746.6 | 0.473 | 0.543 | 0.695 | - | 0.429 | 0.343 | 0.458 | 0.658 |
| PE(36:2) | 744.5 | - | 0.682 | - | - | 0.591 | 0.404 | 0.474 | 0.661 |
| PE(36:5) | 738.5 | - | 1.421 | - | - | - | 2.103 | 2.573 | 2.105 |
| PE(38:1) | 774.6 | 0.621 | 0.565 | 0.695 | 0.666 | 0.324 | 0.156 | 0.160 | 0.223 |
| PE(38:2) | 772.6 | 0.39 | 0.34 | 0.605 | 0.541 | 0.246 | 0.086 | 0.102 | 0.136 |
| PE(40:2) | 800.6 | 0.38 | 0.327 | 0.412 | 0.456 | 0.323 | 0.091 | 0.146 | 0.185 |
| PE(40:5) | 794.6 | - | - | - | 0.186 | 0.211 | - | 0.200 | 0.237 |
| PE(42:7) | 818.6 | - | 0.691 | - | - | - | 0.486 | 0.435 | 0.505 |
| PI(34:1) | 854.5 | - | 0.591 | 0.697 | - | - | 0.669 | - | - |
| PI(36:1) | 882.6 | 0.358 | 0.373 | 0.585 | - | - | 0.440 | - | - |
| PI(36:2) | 880.6 | 0.555 | 0.448 | 0.59 | - | - | 0.510 | - | - |
| PI(40:4) | 932.6 | - | 0.532 | 0.688 | - | 0.470 | 0.459 | - | 0.694 |
| PI(40:5) | 930.6 | - | - | 0.611 | - | - | 0.217 | 0.323 | 0.361 |
| PS(36:1) | 790.6 | - | 0.518 | 0.545 | 0.689 | - | 0.524 | - | - |
| PS(36:4) | 784.5 | - | 1.57 | - | - | 10.076 | 7.046 | 2.255 | - |
| PS(38:1) | 818.6 | - | 0.299 | 0.298 | 0.435 | 0.683 | 0.221 | 0.165 | 0.139 |
| PS(38:2) | 816.6 | - | 0.448 | 0.467 | - | - | 0.312 | 0.222 | 0.216 |
| PS(38:3) | 814.6 | 1.48 | - | - | - | 1.782 | - | - | - |
| PS(38:4) | 812.5 | 2.304 | - | - | - | 3.079 | 2.012 | 1.633 | - |
| PS(38:5) | 810.5 | 2.196 | - | - | - | 3.619 | 1.790 | - | - |
| PS(40:1) | 846.6 | - | 0.679 | 0.687 | - | - | 0.524 | - | - |
| PS(40:2) | 844.6 | - | 0.362 | 0.477 | - | - | 0.369 | 0.457 | 0.566 |
| PS(40:8) | 832.5 | - | 0.592 | 0.668 | - | - | 0.579 | - | 0.640 |
| PS(42:5) | 866.6 | - | - | 0.646 | - | - | 0.704 | - | - |
| PS(42:7) | 862.6 | - | 0.647 | 0.695 | - | - | 0.513 | 0.252 | 0.450 |
| PS(42:9) | 858.5 | - | 0.681 | 0.564 | 0.606 | - | 0.587 | - | - |
| SM(16:0) | 703.6 | - | 1.424 | - | - | 1.531 | 1.684 | 1.552 | 1.947 |
| SM(24:1) | 813.7 | - | 0.557 | 0.482 | - | - | 0.463 | 0.454 | - |

The volcano plots (Fig. 6) and Table 2 provide a different perspective that complements the analysis performed on the total lipid levels (Figs. 2-4). For example, the total level of ePS was elevated in both the brain and spinal cord at 6 weeks (Figs. 2-3), however, no individual ePS lipids were identified in the brain at week 6 and only two were identified in the spinal cord (ePS 36:1 and 38:2). Another example is that several PS species were elevated at week 6 in the brain (1.4-2.3 fold) and in the spinal cord (1.7-10.1 fold), but total PS was not significantly elevated. Table 2 also provides further insights into the observation that certain lipid species are increased during active demyelination. At week 6, almost all lipids that were increasing are unsaturated, and many are highly unsaturated with 3, 4, or 5 double bonds (observed in PC, ePC, PS, and ePS). It has been recently recognized that polyunsaturated fatty acids may serve as precursors for a class of lipids known as specialized proresolving mediators that reduce oxidative stress and inflammation, and lead to improved tissue regeneration.^23, 24^ Further studies will be needed to determine if unsaturated lipids are mediating remyelination in the *Plp1*-iCKO-*Myrf* mice.

Several observations from our dataset are consistent with results from other studies in different demyelination models. A limited lipidomics panel focusing on phosphatidylcholine (PC) in cuprizone treated mice, revealed a decrease in PC 36:1 in the corpus callosum and cortex, and this decrease was also observed in post-mortem multiple sclerosis brain tissue.^5^ We also observed that PC 36:1 was decreased at demyelinating timepoints in the brain (weeks 6 and 12) and spinal cord (weeks 6, 12, and 18). Another study used stereotaxic injection of lysolecithin into the corpus callosum to induce demyelination and analyzed the lipids in the lesion site with desorption electrospray ionization-MS imaging. They primarily observed changes to PCs and PEs. For example, PC 36:1 is also decreased in this study, but they observe changes in several other PC and PE species that are distinct from our results.^4^ One confounding factor is that the lysolecithin model relies on an injection of lysolecithin (or LPC), which may lead to lipid alterations that are unique to the lysolecithin detergent used in this model.

Other studies in the cuprizone model have examined more lipid classes during remyelination revealing persistent changes in the prefrontal cortex after demyelination and remyelination. The largest changes included increased PS and decreased LPE.^3, 25^ We also observed increases in PS during demyelination, but these increases diminished at later timepoints. We did not observe major changes in LPE, and overall, we observed few changes in lyso-lipids, which contain a single acyl chain. Most changes were in the diacylated phospholipids (PS, PE, etc.) and ether-linked phospholipids (ePS, ePE, etc.).

### Treatment with a remyelinating drug reveals lipid biomarkers of remyelination

Sob-AM2 is a CNS-penetrating thyroid hormone agonist that has been shown to promote remyelination in the brains of *Plp1*-iCKO-*Myrf* mice.^7^ To define how the lipidome is altered by treatment with this remyelinating agent, the *Plp1*-iCKO-*Myrf* mice were administered chow containing Sob-AM2 (84 ug/kg nominal daily dose) starting at week 2 post-tamoxifen, a Sob-AM2 treatment method that was used and validated previously. Mice were euthanized at week 12 and week 18 and brain and spinal cord tissues were analyzed by the lipidomics panel at the same time as the untreated *Cre* negative and *Cre* positive samples.

To identify drug-related changes to the lipidome, the total lipid levels were analyzed with pairwise comparisons (*Cre* negative on control chow versus *Cre* negative on SobAM2 chow and *Cre* positive on control chow versus *Cre* positive on SobAM2 chow) at weeks 12 and 18. Sob-AM2 treatment elicited significant changes in several brain lipid classes (Fig. 7) but had a minimal impact on lipids in the spinal cord (Fig. S5).

**Figure 7.**
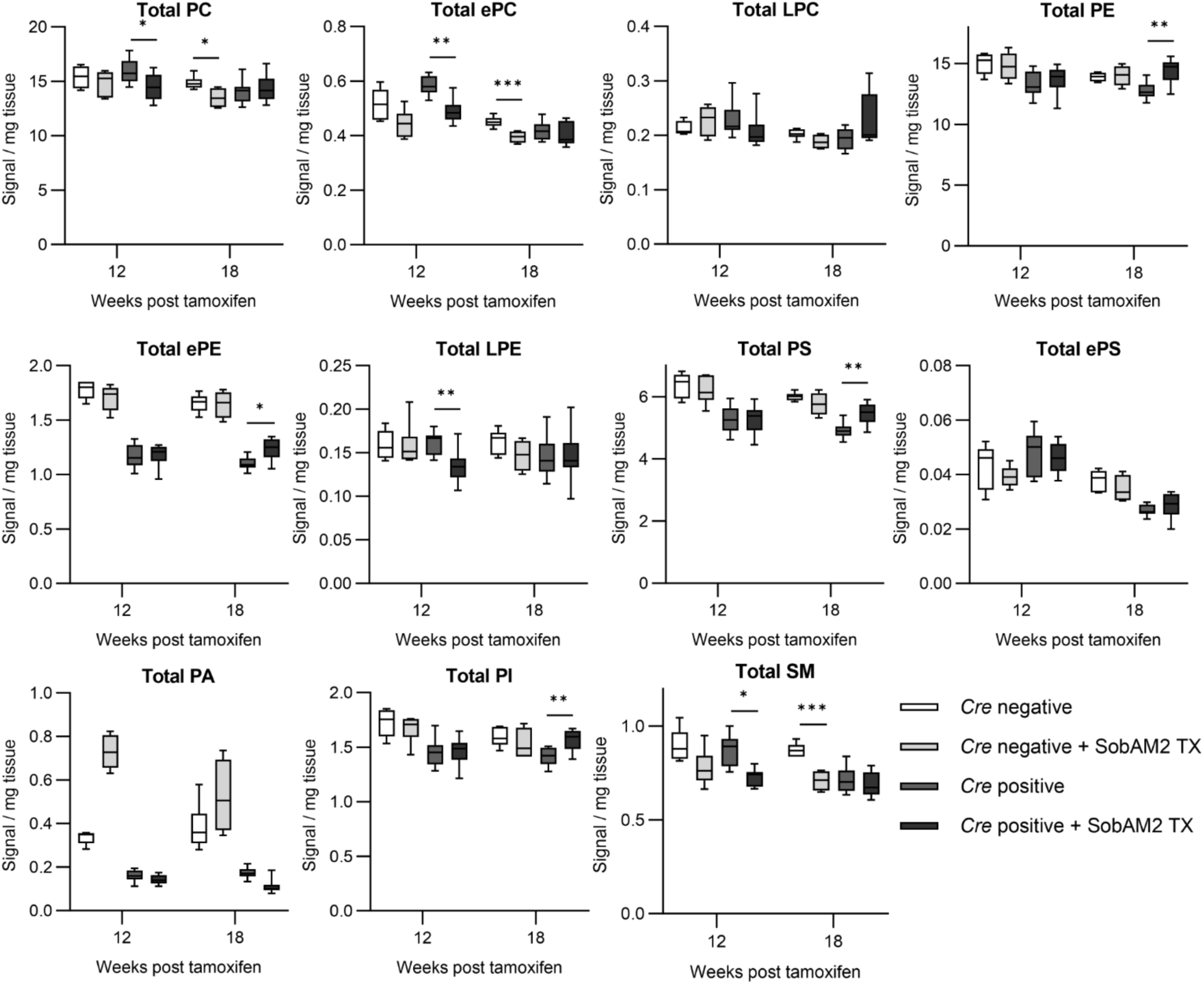
Total lipid levels in brain tissue at 12- and 18-weeks post-tamoxifen with Sob-AM2 treatment (TX). The total lipid levels for 11 major classes of lipids are shown. All individual lipids were measured for each class and the values were summed for this figure. The data are plotted as box and whisker plots with the bars representing the minimum and maximum values. Statistical analysis was performed with multiple t-tests comparing the different genotypes with and without Sob-AM2 treatment at each timepoint using a Holm-Šídák correction for multiple comparisons. The sample numbers are the following (*Cre* negative control: 12 weeks n = 5, 18 weeks n = 6; *Cre* negative SobAM2: 12 weeks n = 6, 18 weeks n = 5; *Cre* positive control: 12 weeks n = 8, 18 weeks n = 9; *Cre* positive SobAM2: 12 weeks n = 8, 18 weeks n = 9). Abbreviations: PC, phosphatidylcholine; ePC, ether-linked PC; LPC, lyso-PC; PE, phosphatidylethanolamine; ePE, ether-linked PE; LPE, lyso-PE; PS, phosphatidylserine; ePS, ether-linked PS; PA, phosphatidic acid; PI, phosphatidylinositol; SM, sphingomyelin.

Our goal was to identify Sob-AM2 induced lipid changes that may be relevant to the remyelinating effects of the drug. Four classes (PE, ePE, PS, and PI) all showed a similar and interesting trend. Above, we noted that all four classes were decreased in the brains experiencing demyelination during week 12 and week 18 (Fig. 2). With the addition of Sob-AM2 treatment, we observed increases in the total levels of PE, ePE, PS, and PI at week 18 (Fig. 7). Critically, this only occurred in *Cre* positive mice that had undergone demyelination; drug treatment had no effect in the absence of demyelination. Sob-AM2 treatment effectively normalizes the amounts of PE, PS, and PI, which correlates with Sob-AM2 inducing remyelination in the brain by 18 weeks post-tamoxifen. This also suggests that total PE, PS, and PI levels may correlate with both demyelination and remyelination. This is further corroborated by the results in the spinal cord (Fig. 3), which showed substantial drops in PE and PS during all measured timepoints.

To further confirm the drug-induced changes on the lipidome and their relevance to drug-induced remyelination, we performed PCA on the four groups (*Cre* positive and *Cre* negative with and without Sob-AM2) in brain and spinal cord at week 12 and week 18 (Fig. 8 and Fig. S6). In the PCA score plots, the lipidome of brains from *Cre* positive mice treated with Sob-AM2 cluster separately from *Cre* positive mice that received control (Fig. 8A). This supports the assertion that remyelination induced by Sob-AM2 correlates with an altered lipidomic profile.

**Figure 8.**
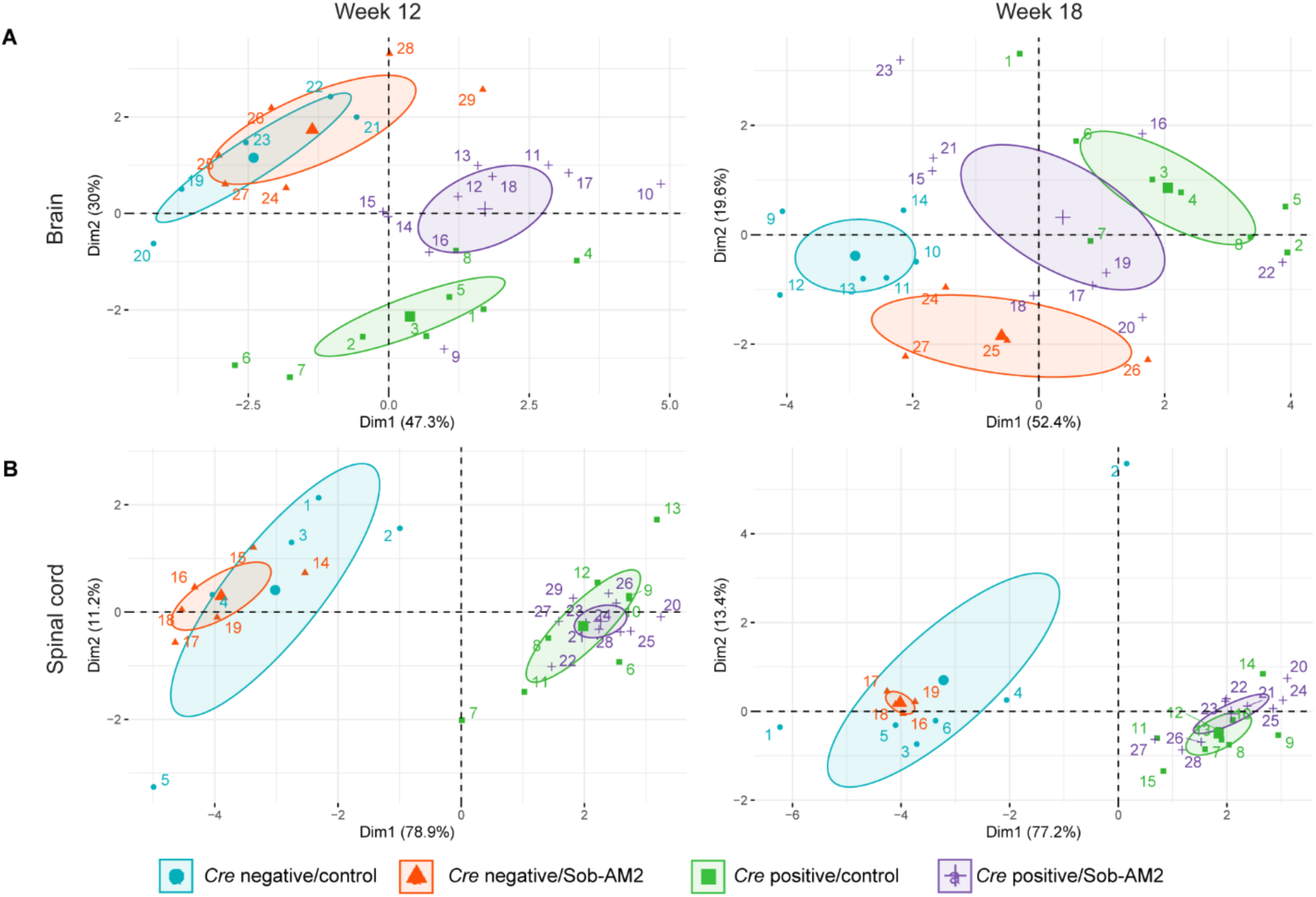
Principal component analysis (PCA) discriminates between the brain lipidomes of mice treated with control or Sob-AM2. PCA was performed on each timepoint (12 and 18 weeks post-tamoxifen) in (A) brain and (B) spinal cord. The samples were clustered into four groups based on genotype (*Cre* negative and *Cre* positive) and treatment (control and Sob-AM2). The total levels for each lipid class were used in this analysis (Tables S2-S3).

In contrast to the brain, Sob-AM2 had a limited effect on the lipidome of the spinal cord. This was supported both by the total lipid analysis (Fig. S5) and the grouped PCA score plot (Fig. 8B). The PCA showed that control and Sob-AM2 treated groups clustered identically for *Cre* negative and *Cre* positive mice. Previously, it was shown that Sob-AM2 showed good efficacy in brain remyelination in the *Plp1*-iCKO-*Myrf* mice,^7^ but the effect of Sob-AM2 on the spinal cord was not thoroughly characterized. The lack of a Sob-AM2 treatment effect in the spinal cord is explained in part by the extensive demyelination observed in the spinal cords of *Plp1*-iCKO-*Myrf* mice (Fig. 1), which could limit the efficacy of any remyelination therapy. Further research is necessary to understand why the remyelination in the spinal cord is impaired relative to the brain in the *Plp1*-iCKO-*Myrf* model.

Although changes were observed in the brain with Sob-AM2 treatment at the total lipid level (Fig. 7) and by PCA groupings of the total lipid levels, volcano plot analysis on individual lipid data revealed very few (<5) lipids that were altered with Sob-AM2 treatment in the brain or spinal cord (Fig. S7).

### Serum Lipidomics

Changes in serum lipid levels have potential therapeutic applications as peripheral biomarkers of CNS disease. To determine if any serum lipids were altered during demyelination, serum samples were collected and analyzed from the groups at weeks 6, 12, and 18. In the total lipid analysis, only one significant change was observed; LPC was reduced by 17% at week 12 (Figs. 9A and S8). In addition, PCA score plots discriminated between demyelination and remyelination in the serum at week 12, but not at week 6 or 18 (Fig. 9B and S9). However, no changes to individual lipids were significant in the volcano plot analysis (Fig. S10). The reduction in total serum LPC should be further validated in future studies, however, there is precedent in the literature for alterations to lyso-lipids in inflammatory demyelinating disease. LPC can be converted to lysophosphatidic acid (LPA) by the action of the enzyme autotaxin,^26^ which is known to be upregulated in multiple sclerosis.^27^ Elevations in LPA have also been observed in serum and cerebrospinal fluid samples from people with multiple sclerosis.^28^ LPA was not measured in this study, but the observed reduction in LPC correlates with previous findings and supports the further investigation of LPC/LPA/autotaxin as peripheral biomarkers for demyelinating neurological diseases.

**Figure 9.**
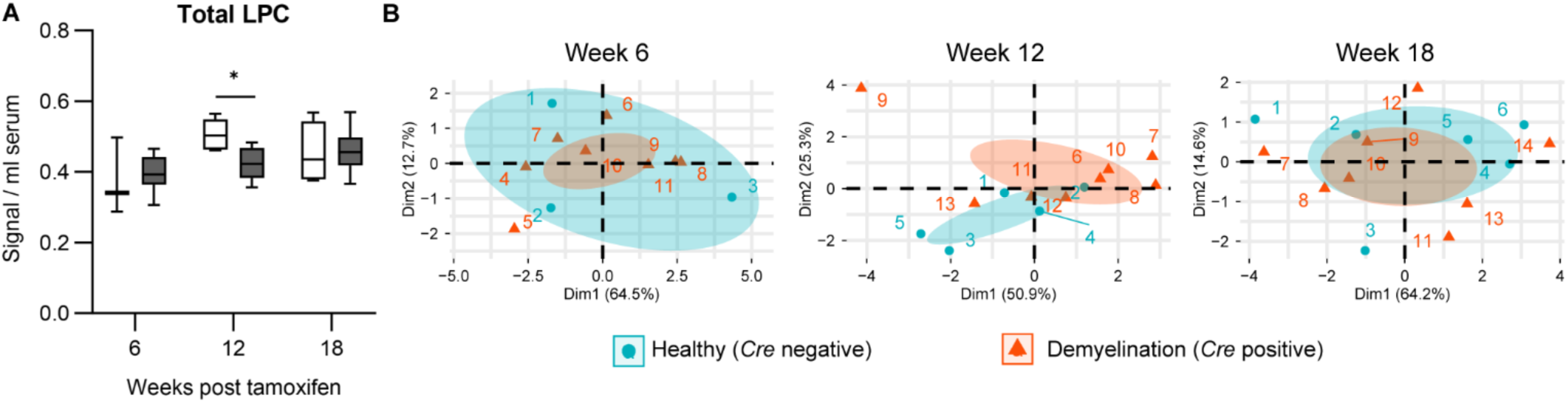
Serum lysophosphatidylcholine (LPC) is reduced at week 12 post-tamoxifen. (A) Total LPC levels at 6, 12, and 18 weeks post-tamoxifen are shown. The data are plotted as box and whisker plots with the bars representing the minimum and maximum values. Statistical analysis was performed with multiple t-tests comparing the different genotypes with and without Sob-AM2 treatment at each timepoint using a Holm-Šídák correction for multiple comparisons. (B) Principal component analysis of serum samples demonstrated no clustering of healthy (*Cre* negative) and demyelinated (*Cre* positive samples) at weeks 6 and 18. There is more separation of the two groups at week 12, which corresponds to peak demyelination in the brain.

Finally, we analyzed how treatment with Sob-AM2 affected the serum lipidome. Sob-AM2 treatment led to many more changes in the serum lipidome (Figs. S11-S13) as compared to the brain and spinal cord. Sob-AM2 is a thyroid hormone agonist, and thyroid hormone is a global regulator of lipid homeostasis in the periphery.^29^ Therefore, it is not surprising that Sob-AM2 would have greater effects in serum. A recent study examined the serum lipidomes of patients with hyper- and hypothyroidism and they identified lipids related to glycerophospholipid metabolism as potential biomarkers associated with thyroid dysfunction.^30^ We observed that Sob-AM2 treatment primarily altered the levels of the PC, ePC, LPC, and PI classes in both the total lipid analysis (Figs. S11-S12) and in the individual lipid volcano plots (Fig. S13 and Table S13). Most changes observed with Sob-AM2 treatment are observed in both *Cre* negative (healthy) and *Cre* positive (demyelination) samples, suggesting that these changes are likely not directly related to the therapeutic action of Sob-AM2 in the CNS as a remyelinating drug.

## CONCLUSION

This study represents the first longitudinal study that measures the lipidomes of demyelination and remyelination in multiple CNS tissues (brain and spinal cord) and in the periphery (serum). The brain and spinal cord tissues have distinct lipidomic profiles consistent with robust remyelination in the brain and impaired remyelination in the spinal cord. In contrast, the serum showed very few alterations with demyelination. The differences between brain and spinal cord were observed through PCA score plots using total lipid levels and through volcano plot analysis of individual lipids. Early stages of demyelination were marked by increased total levels of ePS and ePC and by increases in lipid species containing unsaturated fatty acids. Chronic demyelination, in contrast, was associated with persistent reductions in several lipid classes including PE and PS. Treatment with Sob-AM2, a drug known to promote remyelination, normalized the levels of PE and PS in the brain, suggesting that these lipids may be potential biomarkers for tracking demyelination and remyelination in the CNS. Future studies will focus on elucidating the biological mechanisms underlying lipid alterations that were observed in this study with the goal of revealing how lipids are regulated during demyelination and remyelination.

## ASSOCIATED CONTENT

### Supporting Information

Supplementary figures contain additional information: Figures S1, S2, S7, S9, and S12: PCA biplots related to Figures 4, 5, 8, and 9; Figure S3: PCA on samples classified by mass spectrometry sample order; Figure S4: Fold change values for individual lipids that change in the brain or spinal cord; Figure S5: Total lipid levels for spinal cord tissues treated with and without SobAM2 at weeks 12 and 18 post-tamoxifen; Figure S7: Volcano plots for brain and spinal cord tissues treated with and without SobAM2 at weeks 12 and 18 post-tamoxifen; Figure S8: Total lipid levels for serum from *Cre* negative and *Cre* positive samples at weeks 6, 12, and 18 post-tamoxifen; Figure S10: Volcano plots for serum from *Cre* negative and *Cre* positive samples at weeks 6, 12, and 18 post-tamoxifen; Figure S11: Total lipid levels for serum treated with and without SobAM2 at weeks 12 and 18 post-tamoxifen; Figure S13: Volcano plots for serum treated with and without SobAM2 at weeks 12 and 18 post-tamoxifen (PDF).

Supplementary tables provide the following additional information: Table S1: Mass spectrometry instrumentation parameters and internal standard information; Tables S2-S4: Lipid levels for brain, spinal cord, and serum; Tables S5-S7: Statistical analysis on total lipid levels for brain, spinal cord, and serum; Tables S8-S10: Statistical analysis on individual lipid levels for brain, spinal cord, and serum for volcano plot preparation; Tables S11-S13: Lists of significant individual lipids identified in volcano plots for brain, spinal cord, and serum (ZIP).

Supplementary code for generating PCA and volcano plot figures, tables, and statistics (ZIP). AUTHOR INFORMATION

## Author Contributions

M.D.H. designed the experiments. H.K., J.M.W., N.D.S.M., and E.H.C. performed the in vivo experiments. N.D.S.M. and R.B. carried out the lipid extractions and sample preparation. H.K., M.P.E., and M.D.H. analyzed the data. M.D.H., N.D.S.M., and H.K. wrote the manuscript. All authors have given approval to the final version of the manuscript.

## Supporting information

Supplemental Codes

## ACKNOWLEDGMENT

The lipid analyses described in this work were performed at the Kansas Lipidomics Research Center Analytical Laboratory. Instrument acquisition and lipidomics method development were supported by the National Science Foundation (including support from the Major Research Instrumentation program; most recent award DBI-1726527), K-IDeA Networks of Biomedical Research Excellence (INBRE) of National Institute of Health (P20GM103418), USDA National Institute of Food and Agriculture (Hatch/Multi-State project 1013013), and Kansas State University.

## ABBREVIATIONS

PC, phosphatidylcholine; ePC, ether-linked PC; LPC, lyso-PC; PE, phosphatidylethanolamine; ePE, ether-linked PE; LPE, lyso-PE; PS, phosphatidylserine; ePS, ether-linked PS; PA, phosphatidic acid; PI, phosphatidylinositol; SM, sphingomyelin; PCA, principal component analysis; CV, coefficient of variation; OPC, oligodendrocyte precursor cell; CNS, central nervous system.

## REFERENCES

1. Poitelon, Y.; Kopec, A. M.; Belin, S., Myelin fat facts: an overview of lipids and fatty acid metabolism. Cells 2020, 9 (4).

2. Lloyd, A. F.; Miron, V. E., The pro-remyelination properties of microglia in the central nervous system. Nat Rev Neurol 2019, 15 (8), 447–458.

3. Zhou, C. H.; Xue, S. S.; Xue, F.; Liu, L.; Liu, J. C.; Ma, Q. R.; Qin, J. H.; Tan, Q. R.; Wang, H. N.; Peng, Z. W., The impact of quetiapine on the brain lipidome in a cuprizone-induced mouse model of schizophrenia. Biomed Pharmacother 2020, 131, 110707.

4. Bergholt, M. S.; Serio, A.; McKenzie, J. S.; Boyd, A.; Soares, R. F.; Tillner, J.; Chiappini, C.; Wu, V.; Dannhorn, A.; Takats, Z.; Williams, A.; Stevens, M. M., Correlated heterospectral lipidomics for biomolecular profiling of remyelination in multiple sclerosis. ACS Cent Sci 2018, 4 (1), 39–51.

5. Trepanier, M. O.; Hildebrand, K. D.; Nyamoya, S. D.; Amor, S.; Bazinet, R. P.; Kipp, M., Phosphatidylcholine 36:1 concentration decreases along with demyelination in the cuprizone animal model and in post-mortem multiple sclerosis brain tissue. J Neurochem 2018, 145 (6), 504–515.

6. Zirngibl, M.; Assinck, P.; Sizov, A.; Caprariello, A. V.; Plemel, J. R., Oligodendrocyte death and myelin loss in the cuprizone model: an updated overview of the intrinsic and extrinsic causes of cuprizone demyelination. Mol Neurodegener 2022, 17 (1), 34.

7. Hartley, M. D.; Banerji, T.; Tagge, I. J.; Kirkemo, L. L.; Chaudhary, P.; Calkins, E.; Galipeau, D.; Shokat, M. D.; DeBell, M. J.; Van Leuven, S.; Miller, H.; Marracci, G.; Pocius, E.; Banerji, T.; Ferrara, S. J.; Meinig, J. M.; Emery, B.; Bourdette, D.; Scanlan, T. S., Myelin repair stimulated by CNS-selective thyroid hormone action. JCI Insight 2019, 4 (8).

8. Koenning, M.; Jackson, S.; Hay, C. M.; Faux, C.; Kilpatrick, T. J.; Willingham, M.; Emery, B., Myelin gene regulatory factor is required for maintenance of myelin and mature oligodendrocyte identity in the adult CNS. J. Neurosci. 2012, 32 (36), 12528–42.

9. Emery, B.; Agalliu, D.; Cahoy, J. D.; Watkins, T. A.; Dugas, J. C.; Mulinyawe, S. B.; Ibrahim, A.; Ligon, K. L.; Rowitch, D. H.; Barres, B. A., Myelin gene regulatory factor is a critical transcriptional regulator required for CNS myelination. Cell 2009, 138 (1), 172–85.

10. Ferrara, S. J.; Meinig, J. M.; Placzek, A. T.; Banerji, T.; McTigue, P.; Hartley, M. D.; Sanford-Crane, H. S.; Banerji, T.; Bourdette, D.; Scanlan, T. S., Ester-to-amide rearrangement of ethanolamine-derived prodrugs of sobetirome with increased blood-brain barrier penetration. Bioorg. Med. Chem. 2017, 25 (10), 2743–2753.

11. Meinig, J. M.; Ferrara, S. J.; Banerji, T.; Banerji, T.; Sanford-Crane, H. S.; Bourdette, D.; Scanlan, T. S., Targeting fatty-acid amide hydrolase with prodrugs for CNS-selective therapy. ACS Chem. Neurosci. 2017.

12. Placzek, A. T.; Ferrara, S. J.; Hartley, M. D.; Sanford-Crane, H. S.; Meinig, J. M.; Scanlan, T. S., Sobetirome prodrug esters with enhanced blood-brain barrier permeability. Bioorg. Med. Chem. 2016, 24 (22), 5842–5854.

13. Doerflinger, N. H.; Macklin, W. B.; Popko, B., Inducible site-specific recombination in myelinating cells. Genesis 2003, 35 (1), 63–72.

14. Bligh, E. G.; Dyer, W. J., A rapid method of total lipid extraction and purification. Can J Biochem Physiol 1959, 37 (8), 911–7.

15. Shiva, S.; Vu, H. S.; Roth, M. R.; Zhou, Z.; Marepally, S. R.; Nune, D. S.; Lushington, G. H.; Visvanathan, M.; Welti, R., Lipidomic analysis of plant membrane lipids by direct infusion tandem mass spectrometry. Methods Mol Biol 2013, 1009, 79–91.

16. R_Core_Team A language and environment for statistical computing, R Foundation for Statistical Computing: 2023.

17. A., K.; F.,M. factoextra: Extract and visualize the results of multivariate data analyses, R package version 1.0.7; 2020.

18. Shuken, S. R.; McNerney, M. W., Costs and benefits of popular P-value correction methods in three models of quantitative omic experiments. Anal Chem 2023, 95 (5), 2732–2740.

19. Shuken, S. R. permFDP: Rejection threshold correction using permutation-based FDP estimation, R package version 0.1.0; 2022.

20. Blighe, K.; Rana, S.; Lewis, M. EnhancedVolcano: Publication-ready volcano plots with enhanced colouring and labeling, R package version 1.13.2; 2023.

21. Schmitt, S.; Castelvetri, L. C.; Simons, M., Metabolism and functions of lipids in myelin. Biochim Biophys Acta 2015, 1851 (8), 999–1005.

22. Broniec, A.; Klosinski, R.; Pawlak, A.; Wrona-Krol, M.; Thompson, D.; Sarna, T., Interactions of plasmalogens and their diacyl analogs with singlet oxygen in selected model systems. Free Radic Biol Med 2011, 50 (7), 892–8.

23. Leuti, A.; Maccarrone, M.; Chiurchiu, V., Proresolving Lipid Mediators: Endogenous Modulators of Oxidative Stress. Oxid Med Cell Longev 2019, 2019, 8107265.

24. Brennan, E.; Kantharidis, P.; Cooper, M. E.; Godson, C., Pro-resolving lipid mediators: regulators of inflammation, metabolism and kidney function. Nat Rev Nephrol 2021, 17 (11), 725–739.

25. Zhou, C.; Cai, M.; Wang, Y.; Wu, W.; Yin, Y.; Wang, X.; Hu, G.; Wang, H.; Tan, Q.; Peng, Z., The effects of repetitive transcranial magnetic stimulation on cognitive impairment and the brain lipidome in a cuprizone-induced mouse model of demyelination. Front Neurosci 2021, 15, 706786.

26. Valdes-Rives, S. A.; Gonzalez-Arenas, A., Autotaxin-lysophosphatidic acid: from inflammation to cancer development. Mediators Inflamm 2017, 2017, 9173090.

27. Zahednasab, H.; Balood, M.; Harirchian, M. H.; Mesbah-Namin, S. A.; Rahimian, N.; Siroos, B., Increased autotaxin activity in multiple sclerosis. J Neuroimmunol 2014, 273 (1-2), 120–3.

28. Jiang, D.; Ju, W.; Wu, X.; Zhan, X., Elevated lysophosphatidic acid levels in the serum and cerebrospinal fluid in patients with multiple sclerosis: therapeutic response and clinical implication. Neurol Res 2018, 40 (5), 335–339.

29. Scanlan, T. S., Sobetirome: a case history of bench-to-clinic drug discovery and development. Heart Fail. Rev. 2010, 15 (2), 177–82.

30. Dong, H.; Zhou, W.; Yan, X.; Zhao, H.; Zhao, H.; Jiao, Y.; Sun, G.; Li, Y.; Zhang, Z., Serum lipidomic analysis reveals biomarkers and metabolic pathways of thyroid dysfunction. ACS Omega 2023, 8 (11), 10355–10364.

