## Supplemental Figures 1-13 for "Lipidomic analysis reveals differences in the extent of remyelination in the brain and spinal cord"

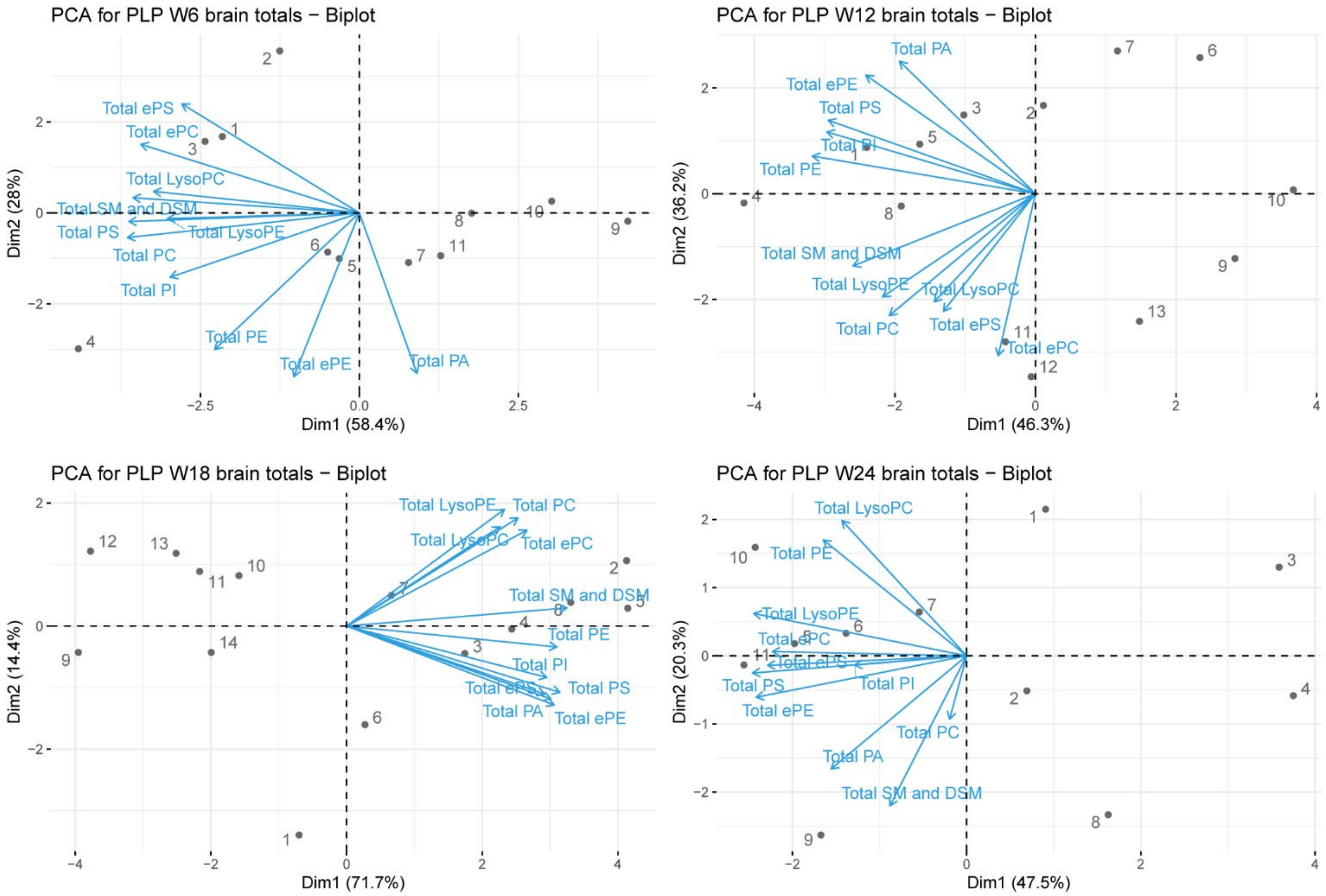


**Figure S1:** PCA biplots of brain samples illustrate lipid class trends, variability, and contribution to each principal component (PC) through arrows. PCA was performed on each timepoint (6, 12, 18, and 24 weeks post-tamoxifen). The total levels for each lipid class were used in this analysis (Table S2). Arrows may point in similar (positive correlation), opposing (negative correlation), or perpendicular (no correlation) directions, with vertical arrows contributing most to PC 1, and horizontal arrows contributing most to PC 2. The length of the arrow represents the variability of a lipid class sum, where longer arrows indicate that PC 1 and 2 contain more variability of that lipid class sum than other PCs.

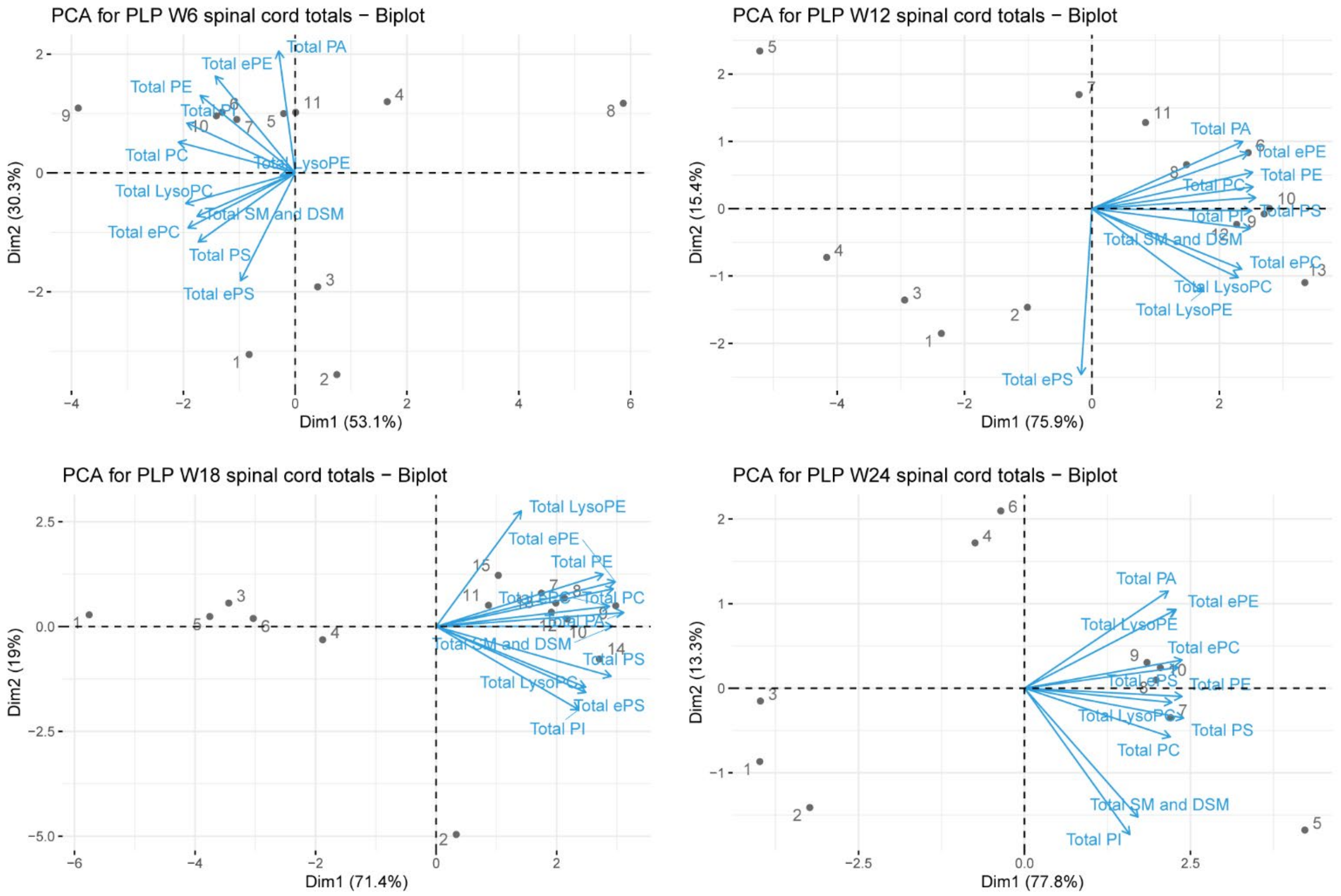


**Figure S2:** PCA biplots of spinal cord samples illustrate lipid class trends, variability, and contribution to each principal component (PC) through arrows. PCA was performed on each timepoint (6, 12, 18, and 24 weeks post-tamoxifen). The total levels for each lipid class were used in this analysis (Table S3). Arrows may point in similar (positive correlation), opposing (negative correlation), or perpendicular (no correlation) directions, with vertical arrows contributing most to PC 1, and horizontal arrows contributing most to PC 2. The length of the arrow represents the variability of a lipid class sum, where longer arrows indicate that PC 1 and 2 contain more variability of that lipid class sum than other PCs.

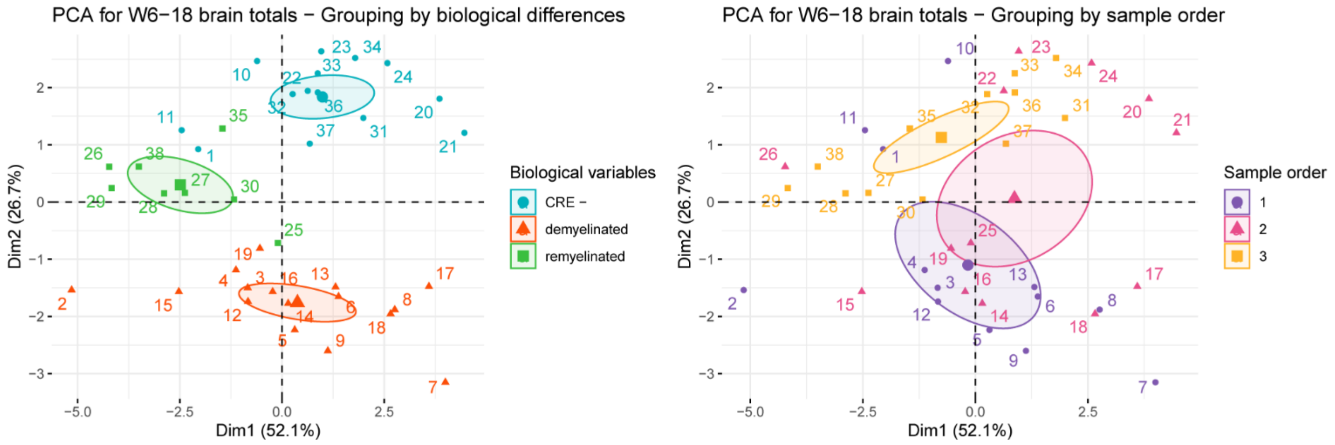


**Figure S3**: Principal component analysis (PCA) discriminates based on biological differences, but not on sample order during analysis. The brain samples were grouped into three groups either by sample order (right) or by phenotype (left). Three main phenotypes were used: (1) Healthy (*Cre* negative, weeks 6-18); (2) Demyelination (*Cre* positive, weeks 6 and 12), and (3) Remyelination (*Cre* positive, week 18). The total levels for each lipid class were used in this analysis (Tables S2). When the groups are divided by phenotype, they show distinct clustering and PCA discriminates between the groups. In striking contrast, PCA analysis of the same samples clustered by sample order showed much greater overlap between groups. These results suggest that the PCA clustering is based on biological differences and not based on mass spectrometry sample order.

| Lipid species | m/z | Brain | | | |
|---|---|---|---|---|---|
| | | Week 6 | Week 12 | Week 18 | Week 24 |
| PA(38:3) | 744.5 | - | 0.505 | - | - |
| PA(36:4) | 714.5 | - | 0.626 | 0.580 | - |
| PI(38:1) | 910.6 | - | - | - | 0.406 |
| ePC(40:4) | 824.6 | 1.463 | - | - | - |

| Lipid species | m/z | Spinal Cord | | | |
|---|---|---|---|---|---|
| | | Week 6 | Week 12 | Week 18 | Week 24 |
| ePC(32:0) | 720.6 | - | 1.892 | - | - |
| ePC(32:2) | 716.6 | 4.043 | 2.679 | - | - |
| ePC(34:1) | 746.6 | - | - | - | 0.679 |
| ePC(36:1) | 774.6 | - | - | - | 0.650 |
| ePC(38:1) | 802.7 | - | - | - | 0.585 |
| ePC(38:3) | 798.6 | - | 0.331 | 0.149 | 0.193 |
| ePC(40:3) | 826.7 | - | - | - | 0.212 |
| ePC(40:5) | 822.6 | - | 0.702 | - | - |
| ePC(40:6) | 820.6 | - | 1.536 | - | - |
| ePE(36:5) | 724.5 | - | 1.422 | - | - |
| ePE(38:4) | 754.6 | - | - | - | 0.626 |
| ePE(38:5) | 752.6 | - | - | - | 0.700 |
| ePE(40:3) | 784.6 | 0.445 | 0.131 | 0.120 | 0.138 |
| ePE(40:4) | 782.6 | - | 0.619 | 0.466 | - |
| ePE(40:5) | 780.6 | - | 0.392 | 0.480 | 0.509 |
| ePE(40:6) | 778.6 | - | 0.707 | 0.616 | 0.652 |
| ePS(34:1) | 748.5 | - | 1.467 | - | 0.450 |
| ePS(36:1) | 776.6 | 2.132 | - | - | - |
| ePS(38:4) | 798.6 | - | 16.143 | - | - |
| ePS(40:4) | 826.6 | - | 4.070 | - | - |
| LPC(18:0) | 524.4 | - | 0.675 | - | 0.647 |
| LPC(18:1) | 522.3 | - | 0.605 | - | - |
| LPC(20:4) | 544.3 | - | - | 2.491 | - |
| LPC(22:6) | 568.3 | - | 0.676 | - | - |
| LPE(16:0) | 454.3 | - | 0.578 | 0.607 | - |
| PA(32:0) | 666.5 | - | - | - | 0.117 |
| PA(40:6) | 766.5 | - | - | - | 2.324 |
| PC(32:1) | 732.5 | 1.873 | 2.330 | 1.921 | - |
| PC(36:1) | 788.6 | 0.655 | 0.482 | 0.599 | - |
| PC(36:2) | 786.6 | 0.637 | 0.469 | 0.522 | 0.621 |
| PC(36:4) | 782.6 | 1.471 | 1.504 | - | - |
| PC(38:2) | 814.6 | - | - | - | 0.140 |
| PC(40:4) | 838.6 | - | 0.650 | 0.668 | - |
| PC(40:8) | 830.6 | - | 0.555 | 0.633 | - |
| PC(42:3) | 868.7 | - | 0.319 | 0.601 | - |

| | | | | | |
|---|---|---|---|---|---|
| PC(42:5) | 864.6 | - | - | - | 0.526 |
| PC(42:7) | 860.6 | 0.647 | 0.471 | 0.476 | 0.545 |
| PC(42:8) | 858.6 | 0.611 | 0.567 | 0.566 | - |
| PC(44:2) | 898.7 | - | 0.291 | 0.382 | 0.403 |
| PE(32:1) | 690.5 | - | 1.655 | - | - |
| PE(34:1) | 718.5 | - | 0.675 | 0.687 | - |
| PE(38:3) | 770.6 | - | - | 0.680 | - |
| PE(38:5) | 766.5 | - | 0.662 | - | - |
| PE(40:4) | 796.6 | 0.587 | 0.579 | 0.662 | - |
| PE(40:8) | 788.5 | - | 0.345 | 0.420 | 0.401 |
| PE(42:2) | 828.6 | - | 0.255 | 0.443 | - |
| PE(42:5) | 822.6 | - | 0.371 | 0.261 | 0.518 |
| PE(42:8) | 816.5 | - | - | 0.626 | - |
| PE(42:9) | 814.5 | - | - | - | 0.483 |
| PE(42:10) | 812.5 | - | - | 0.650 | - |
| PI(38:2) | 908.6 | - | 0.338 | - | - |
| PI(38:5) | 902.5 | 0.501 | 0.508 | 0.553 | 0.542 |
| PI(40:7) | 926.5 | - | 0.625 | - | - |
| PS(34:0) | 764.5 | - | - | - | 0.137 |
| PS(34:1) | 762.5 | 3.934 | 2.487 | - | - |
| PS(34:2) | 760.5 | - | 3.433 | 2.784 | - |
| PS(36:2) | 788.5 | - | 0.577 | 0.565 | 0.681 |
| PS(36:3) | 786.5 | - | 1.887 | - | - |
| PS(40:3) | 842.6 | - | 0.616 | 0.476 | 0.587 |
| PS(44:10) | 884.5 | - | 0.670 | 0.654 | 0.693 |
| SM(18:0) | 731.6 | - | 0.703 | 0.638 | - |
| SM(22:1) | 785.6 | - | 0.193 | 0.244 | 0.295 |
| SM(24:0) | 815.7 | - | 0.494 | 0.663 | - |

**Figure S4**: Fold change values for all individual lipids that change significantly *either* in brain *or* the spinal cord as determined by volcano plot analyses.

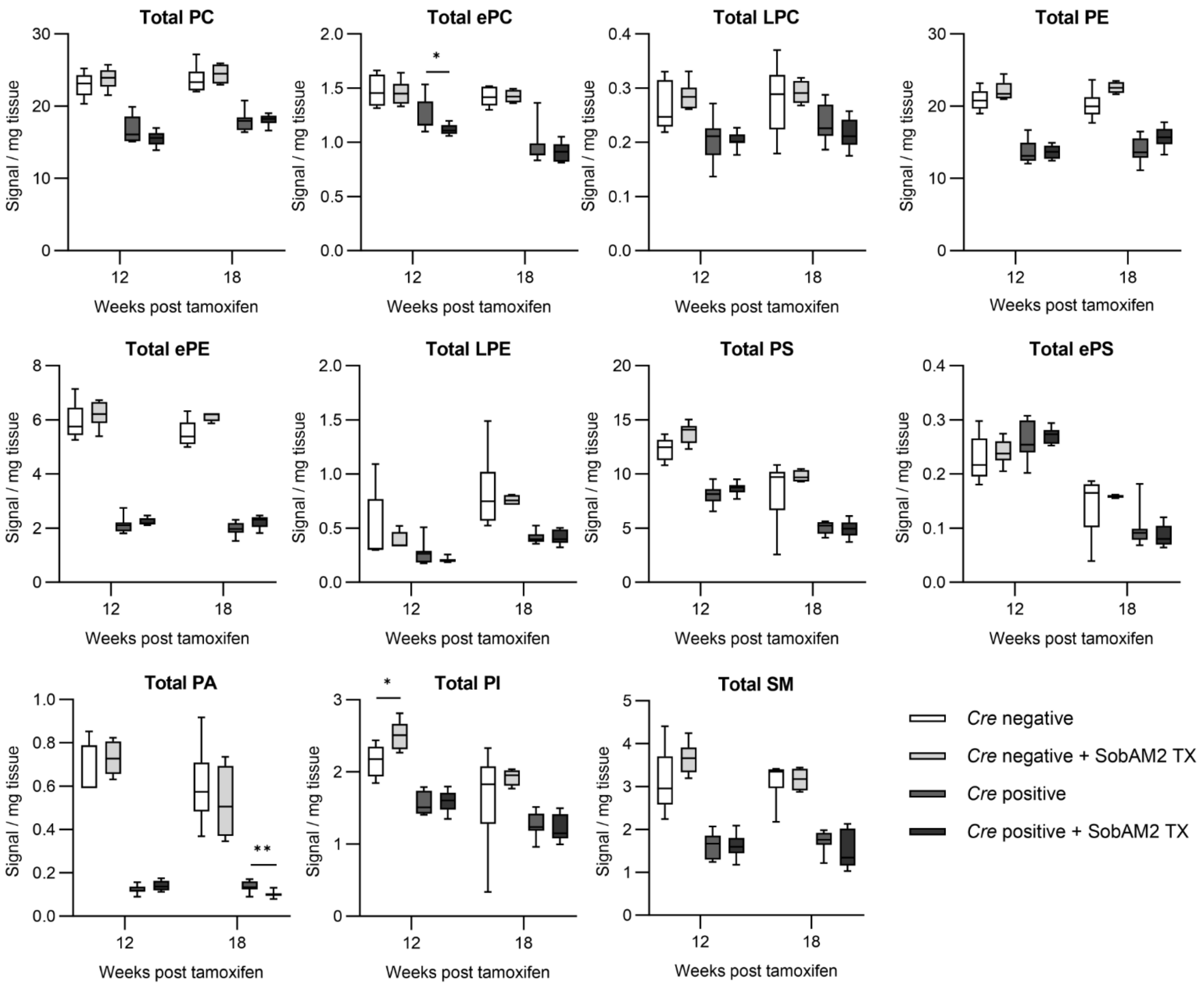


**Figure S5***:* Total lipid levels in spinal cord tissue at 12- and 18-weeks post-tamoxifen treatment with Sob-AM2 treatment (TX). The total lipid levels for 11 major classes of lipids are shown. All individual lipids were measured for each class and the values were summed for this figure. The data are plotted as box and whisker plots with the bars representing the minimum and maximum values. Statistical analysis was performed with multiple t-tests comparing the different genotypes with and without Sob-AM2 treatment at each timepoint using a Holm-Šídák correction for multiple comparisons. The sample numbers are the following (*Cre* negative: 12 weeks n = 5, 18 weeks n = 6; *Cre* negative + SobAM2: 12 weeks n = 6, 18 weeks n = 5; *Cre* positive: 12 weeks n = 8, 18 weeks n = 9; *Cre* positive + SobAM2: 12 weeks n = 8, 18 weeks n = 9). Abbreviations: PC, phosphatidylcholine; ePC, ether-linked PC; LPC, lyso-PC; PE, phosphatidylethanolamine; ePE, ether-linked PE; LPE, lyso-PE; PS, phosphatidylserine; ePS, ether-linked PS; PA, phosphatidic acid; PI, phosphatidylinositol; SM, sphingomyelin.

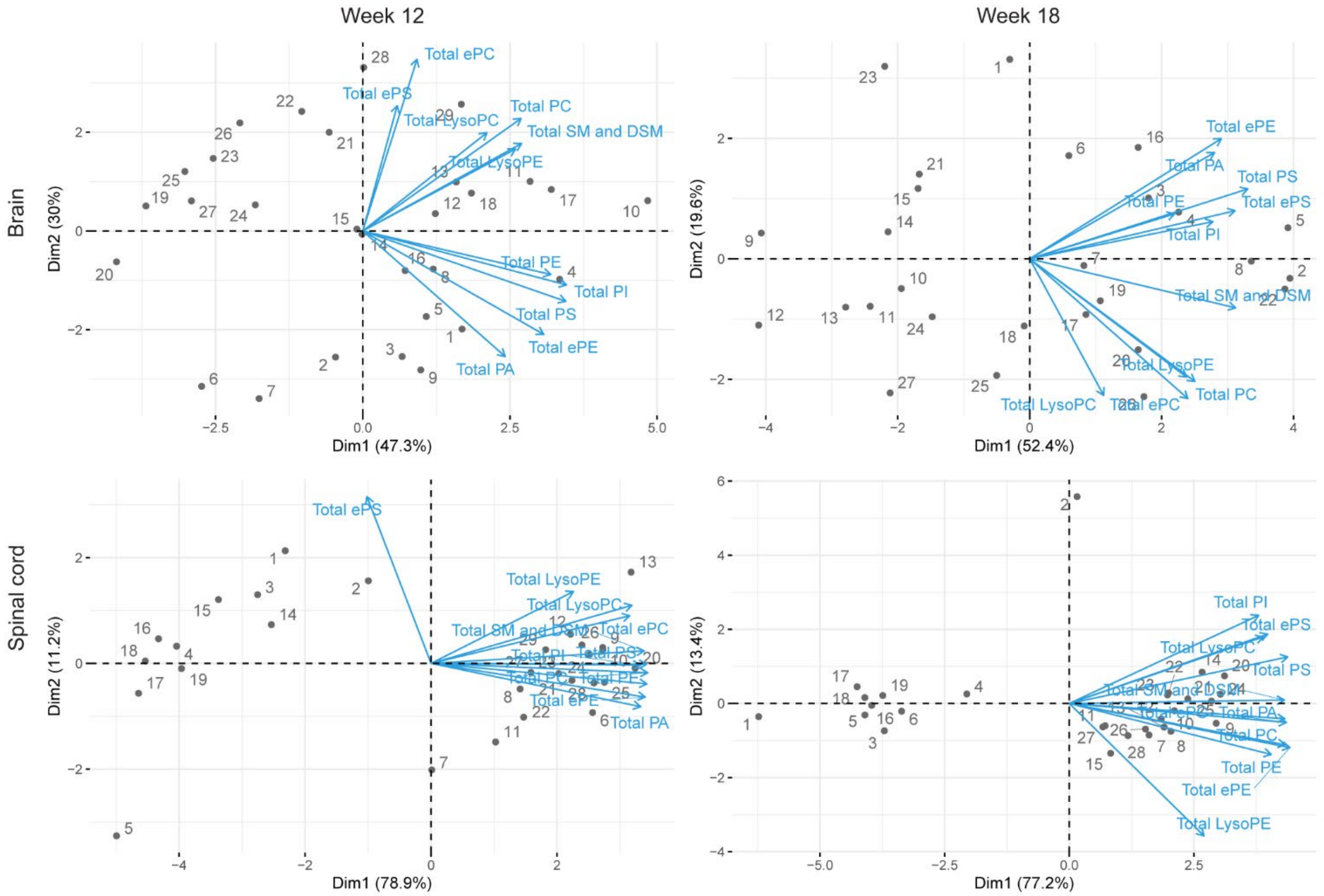


**Figure S6**: PCA biplots illustrate lipid class trends, variability, and contribution to each principal component (PC) through arrows. PCA was performed on each timepoint (12 and 18 weeks post-tamoxifen) in both tissues (brain and spinal cord). The samples were clustered into four groups based on genotype (*Cre* negative and *Cre* positive) and treatment (control and Sob-AM2).The total levels for each lipid class were used in this analysis (Tables S2 and S3). Arrows may point in similar (positive correlation), opposing (negative correlation), or perpendicular (no correlation) directions, with vertical arrows contributing most to PC 1, and horizontal arrows contributing most to PC 2. The length of the arrow represents the variability of a lipid class sum, where longer arrows indicate that PC 1 and 2 contain more variability of that lipid class sum than other PCs.

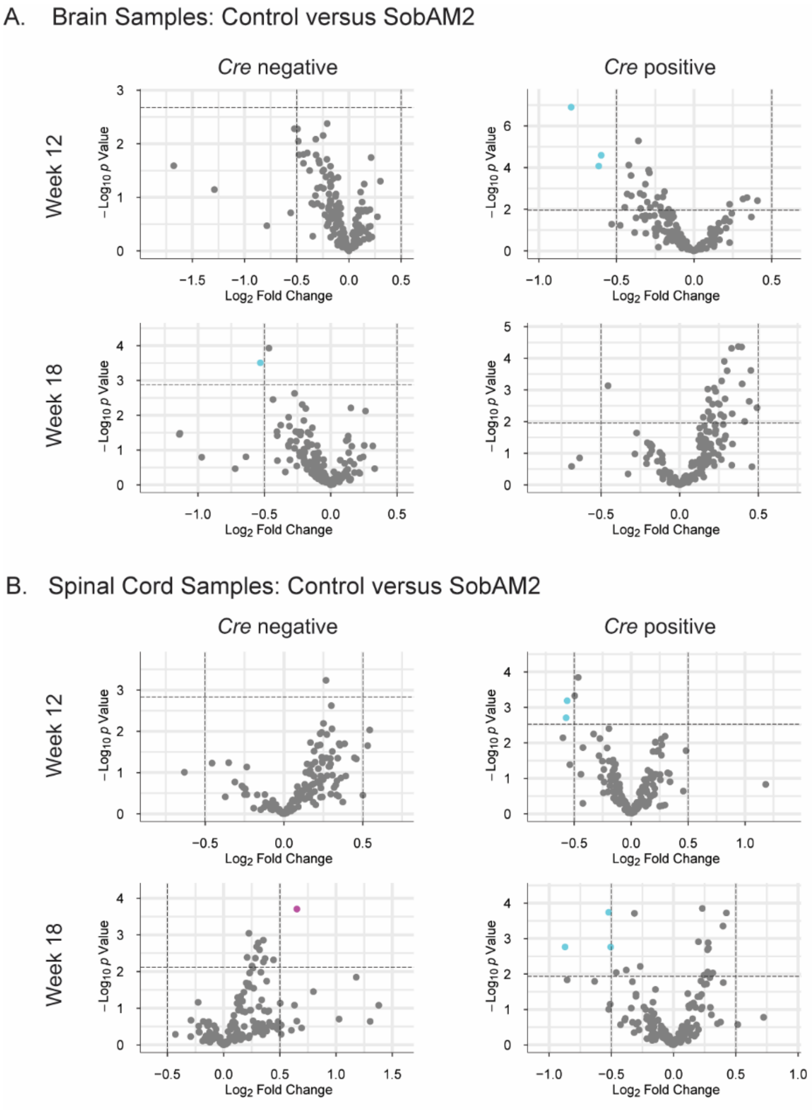


**Figure S7**: Volcano plot analyses of individual lipid changes with SobAM2 treatment show very few significant changes in the brain (A) and spinal cord (B). All lipids for each comparison that met the covariance threshold were included in this analysis (Tables S2-S3). Log2(Fold Change) is plotted on the x-axis and the threshold was set at 0.5. -Log10(P-value) is plotted on the y-axis. The threshold was plotted using a permutation-based false discovery estimation method, and the individual thresholds are listed in Tables S8-S9. Light cyan dots are individual lipids that are decreasing, and magenta dots are lipids that are increasing.

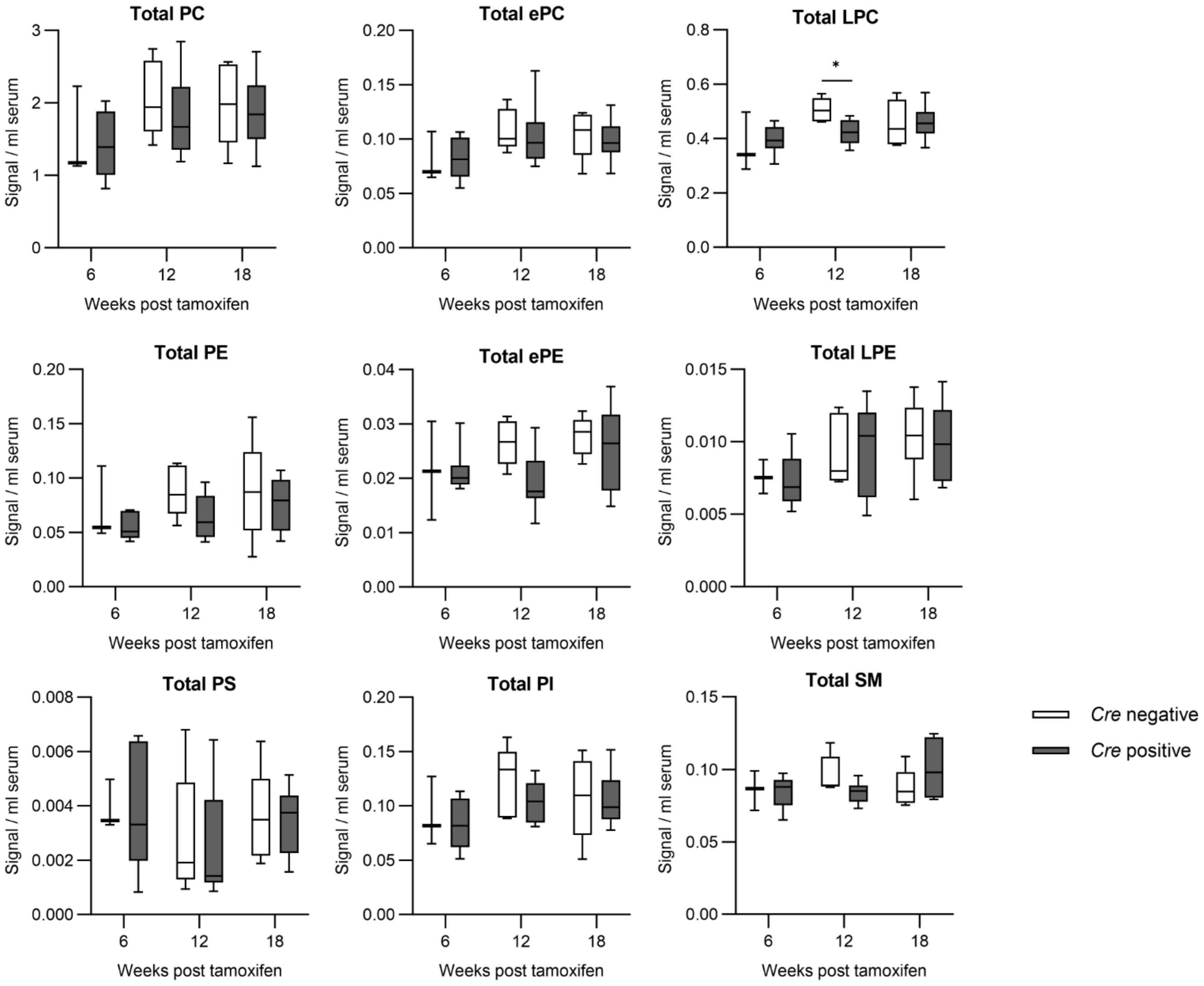


**Figure S8**: Total lipid levels in serum at 6-18 weeks post-tamoxifen treatment. The total lipid levels for the 9 major classes of lipids that met the coefficient of variation threshold (Table S4) are shown. All individual lipids were measured for each class and the values were summed for this figure. The data are plotted as box and whisker plots with the bars representing the minimum and maximum values. Statistical analysis was performed with multiple t-tests comparing *Cre* negative and *Cre* positive at each timepoint using a Holm-Šídák correction for multiple comparisons. The sample numbers are the following (Cre negative: 6 weeks n = 3, 12 weeks n = 5, 18 weeks n = 6; Cre positive: 6 weeks n = 8, 12 weeks n = 8, 18 weeks n = 8). Abbreviations: PC, phosphatidylcholine; ePC, ether-linked PC; LPC, lyso-PC; PE, phosphatidylethanolamine; ePE, ether-linked PE; LPE, lyso-PE; PS, phosphatidylserine; PI, phosphatidylinositol; SM, sphingomyelin.

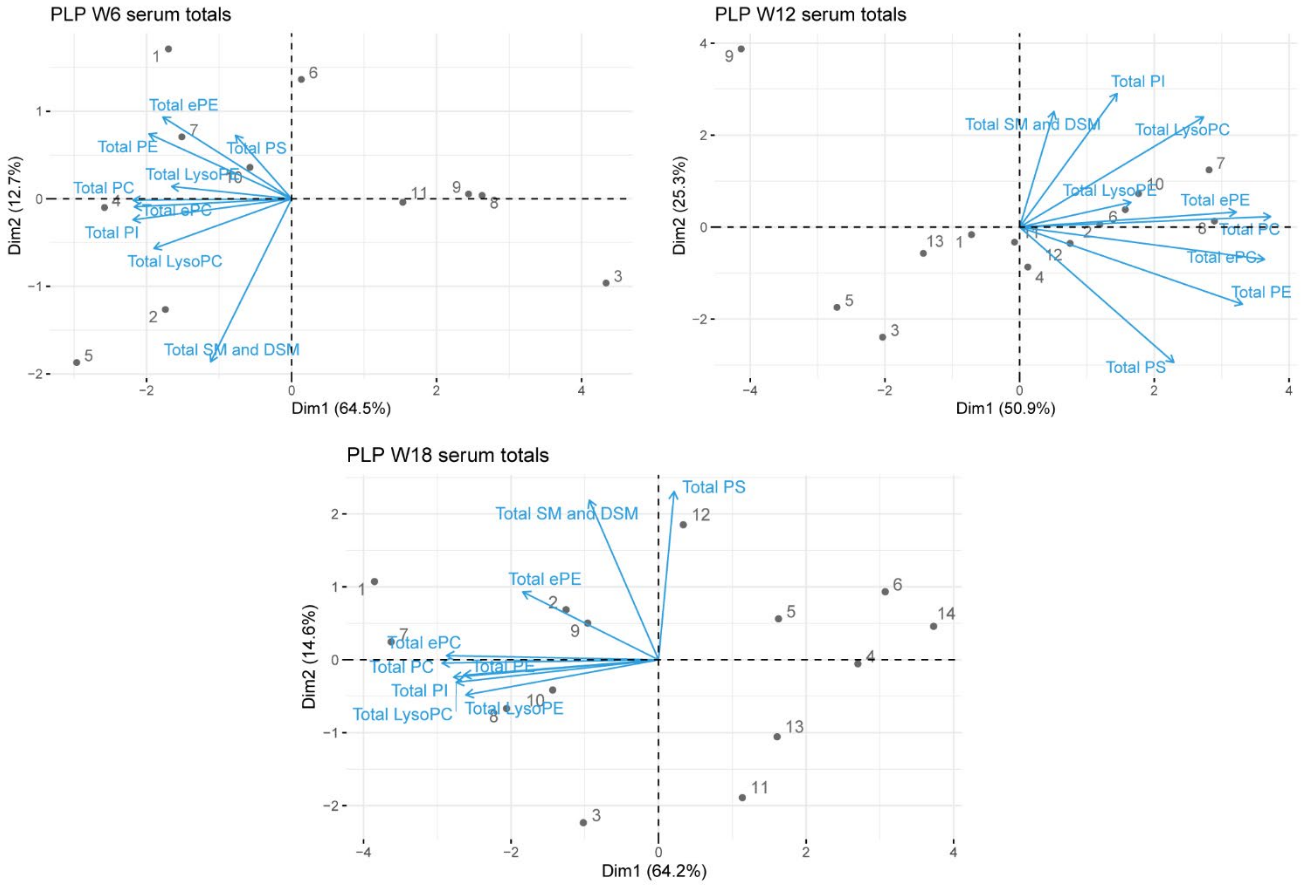


**Figure S10**: PCA biplots of serum samples illustrate lipid class trends, variability, and contribution to each principal component (PC) through arrows. PCA was performed on each timepoint (6, 12, and 18 weeks post-tamoxifen). The total levels for each lipid class were used in this analysis (Table S4). Arrows may point in similar (positive correlation), opposing (negative correlation), or perpendicular (no correlation) directions, with vertical arrows contributing most to PC 1, and horizontal arrows contributing most to PC 2. The length of the arrow represents the variability of a lipid class sum, where longer arrows indicate that PC 1 and 2 contain more variability of that lipid class sum than other PCs.

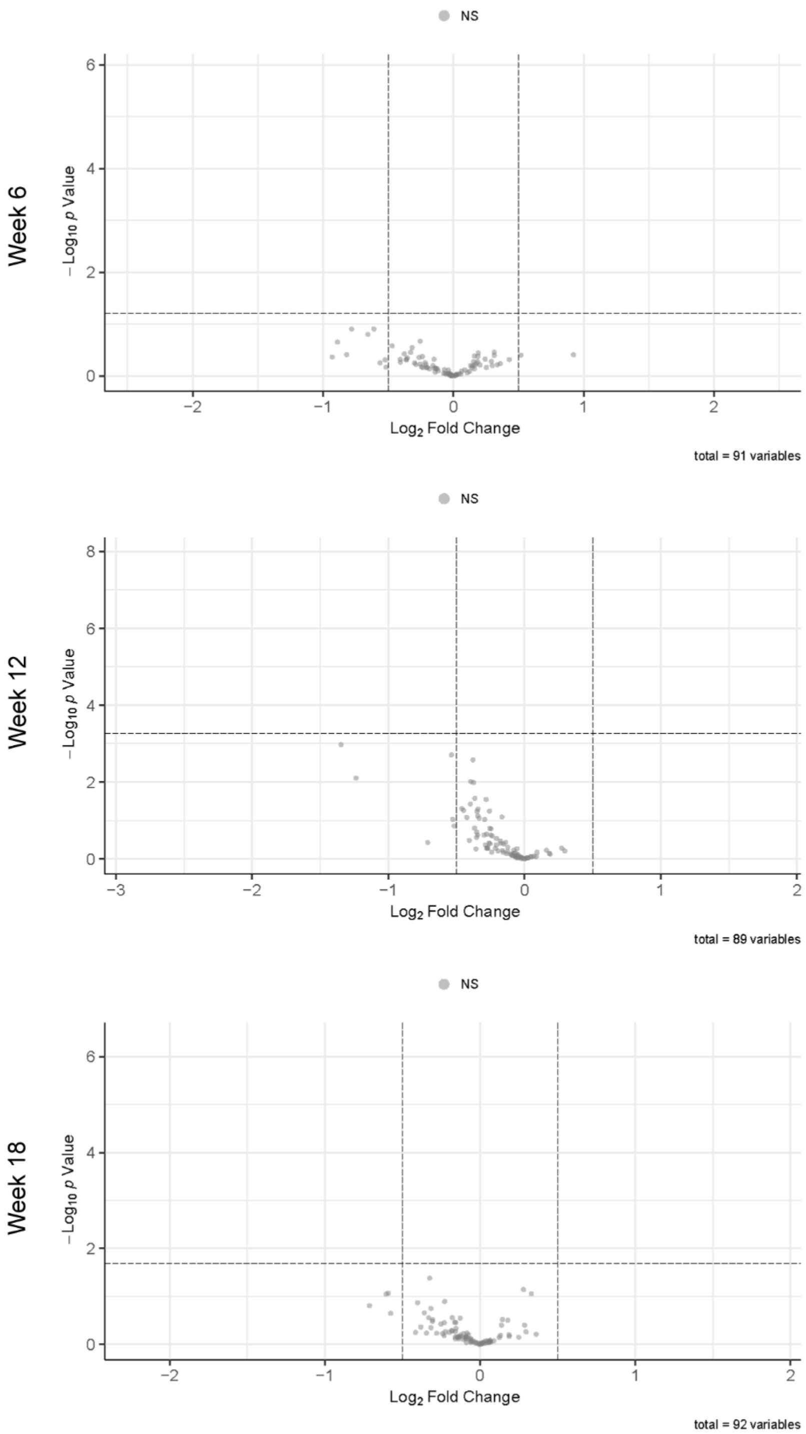


**Figure S10**: No individual lipids in serum were identified by volcano plot analyses of week 6-18 comparing healthy to demyelination samples. All lipids for each comparison that met the coefficient of variation threshold were included in this analysis. Log2(Fold Change) is plotted on the x-axis and the threshold was set at 0.5. The -Log10(P-value) is plotted on the y-axis and the threshold was plotted using a permutation-based false discovery estimation method, and the individual thresholds are listed in Table S10.

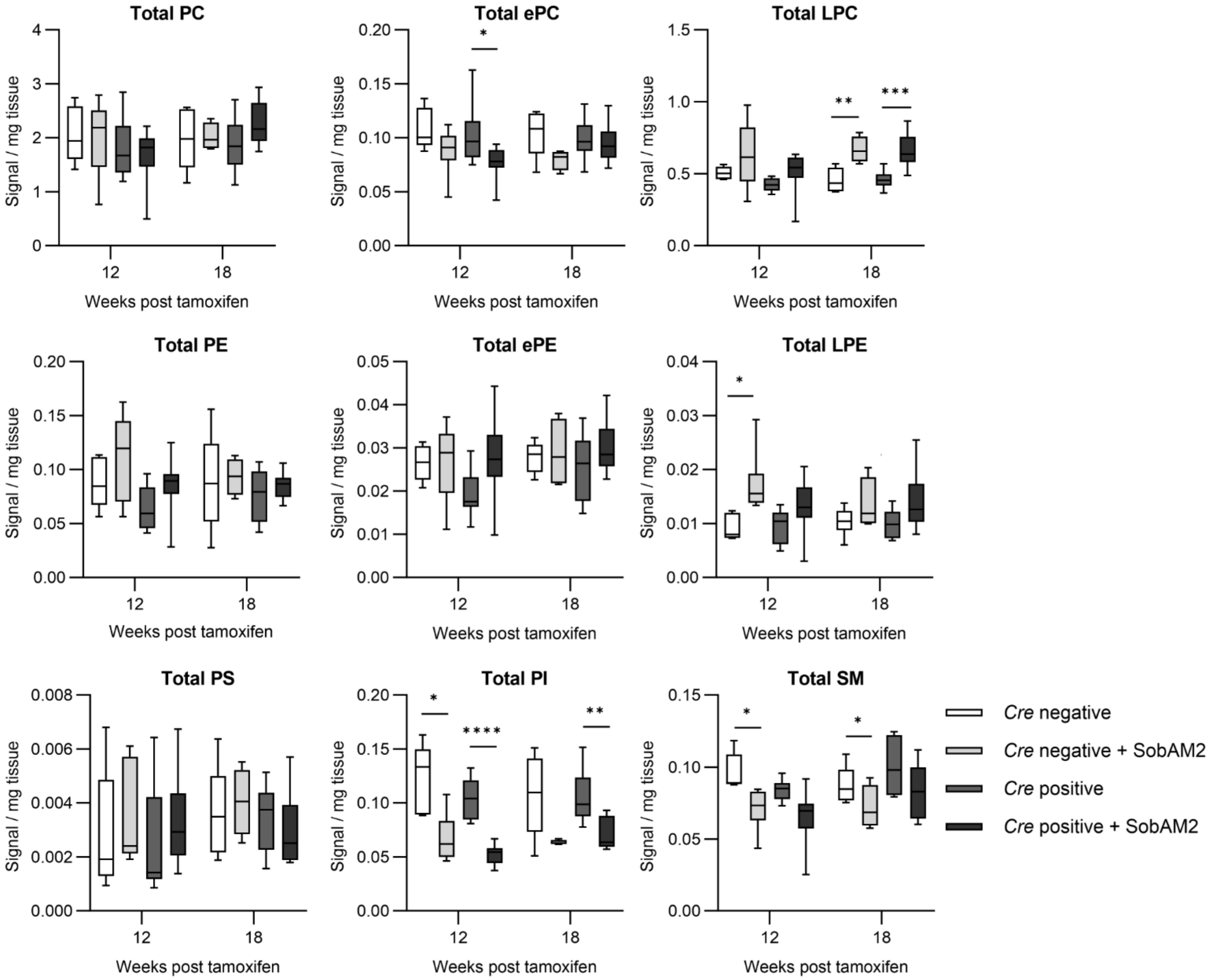


**Figure S11***:* Total lipid levels in serum at 12- and 18-weeks post-tamoxifen treatment with Sob-AM2 treatment (TX), which is remyelinating drug. The total lipid levels for the 9 major classes of lipids that met the coefficient of variation threshold (Table S4) are shown. All individual lipids were measured for each class and the values were summed for this figure. The data are plotted as box and whisker plots with the bars representing the minimum and maximum values. Statistical analysis was performed with multiple t-tests comparing the different genotypes with and without Sob-AM2 treatment at each timepoint using a Holm-Šídák correction for multiple comparisons. The sample numbers are the following (*Cre* negative: 12 weeks n = 5, 18 weeks n = 6; *Cre* negative + SobAM2: 12 weeks n = 6, 18 weeks n = 5; *Cre* positive: 12 weeks n = 8, 18 weeks n = 9; *Cre* positive + SobAM2: 12 weeks n = 8, 18 weeks n = 9). Abbreviations: PC, phosphatidylcholine; ePC, ether-linked PC; LPC, lyso-PC; PE, phosphatidylethanolamine; ePE, ether-linked PE; LPE, lyso-PE; PS, phosphatidylserine; PI, phosphatidylinositol; SM, sphingomyelin.

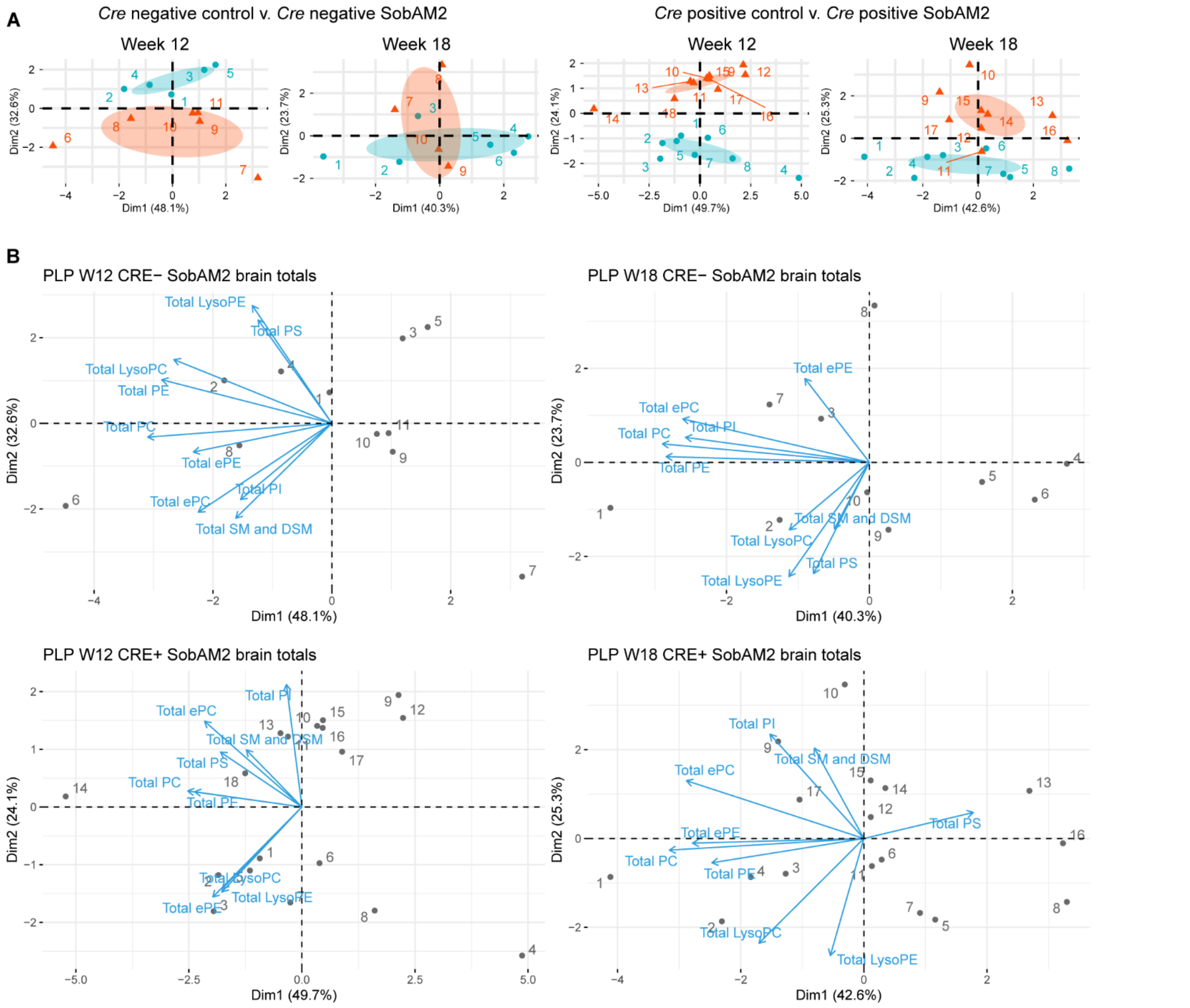


**Figure S12**: PCA plots and related biplots analyzing serum at 12- and 18-weeks post-tamoxifen treatment with and without Sob-AM2 treatment (TX). (**A**) PCA was performed on each timepoint (12 and 18 weeks post-tamoxifen) comparing the lipidomes of *Cre* negative (healthy) mice to *Cre* negative mice treated with Sob-AM2 or the lipidomes of Cre positive (demyelination) mice to Cre positive mice treated with Sob-AM2. The total levels for each lipid class were used in this analysis (Tables S4). (B) Biplots used to generate the PCA groupings shown in part A. Arrows may point in similar (positive correlation), opposing (negative correlation), or perpendicular (no correlation) directions, with vertical arrows contributing most to PC 1, and horizontal arrows contributing most to PC 2. The length of the arrow represents the variability of a lipid class sum, where longer arrows indicate that PC 1 and 2 contain more variability of that lipid class sum than other PCs.

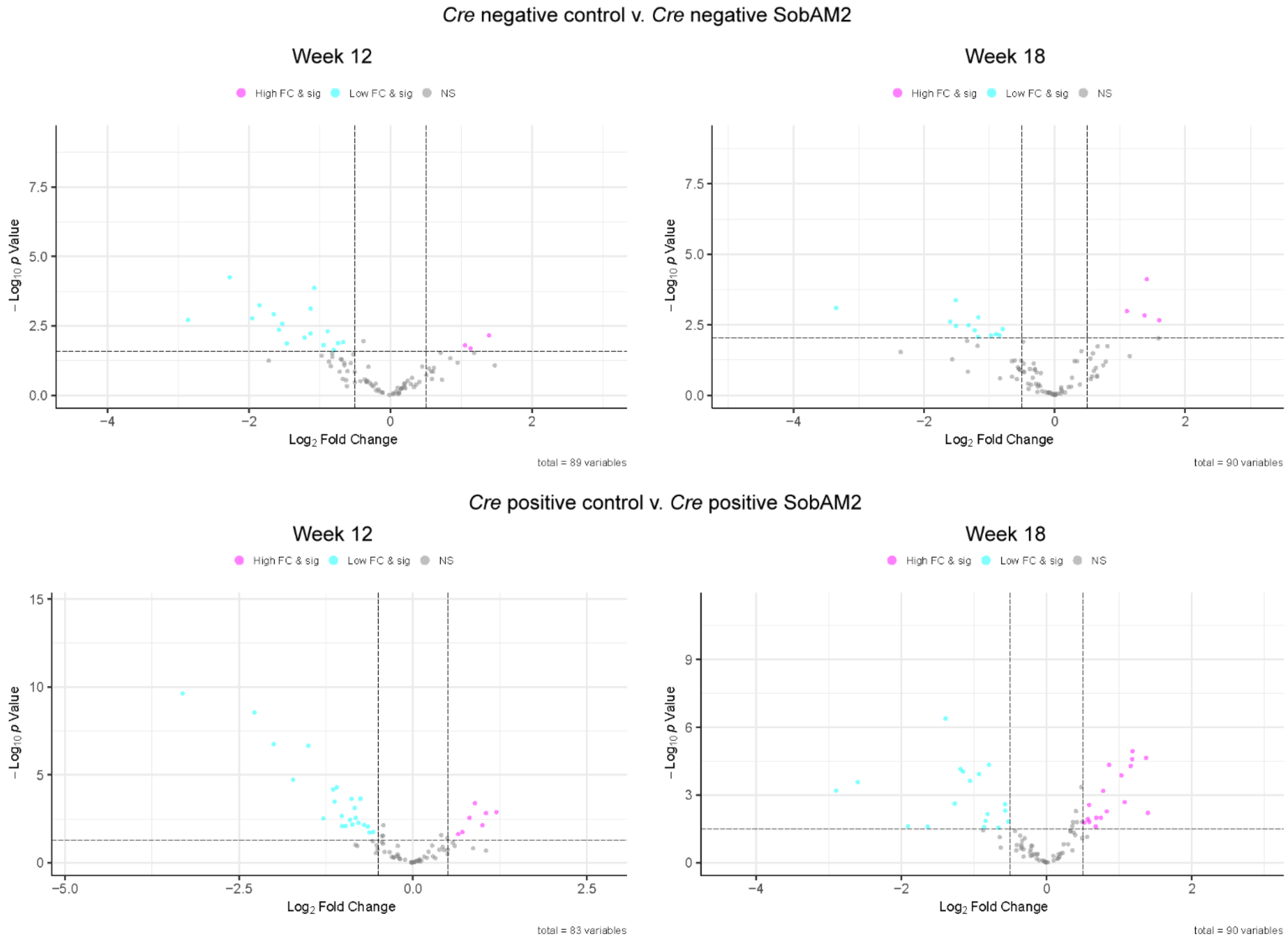


**Figure S13**: Volcano plot analyses of individual lipid changes in serum with SobAM2 treatment demonstrate that Sob-AM2 alters lipid levels both in healthy and demyelinating mice. All lipids for each comparison that met the covariance threshold were included in this analysis (Tables S2-S3). Log2(Fold Change) is plotted on the x-axis and the threshold was set at 0.5. The -Log10(P-value) is plotted on the y-axis. The threshold was plotted using a permutation-based false discovery estimation method, and the individual thresholds are listed in Tables S10. Light cyan dots are individual lipids that are decreasing, and magenta dots are lipids that are increasing.
