## Supplemental Codes for "Lipidomic analysis reveals differences in the extent of remyelination in the brain and spinal cord": Data_Cleaning.R

```
# Cleaning datasets for PCA

library(dplyr)
library(readxl)
library(tidyverse)
library(data.table)
library(janitor)

## PLP Brain 6 - 18 ####

Week6_18_Brain <- read_excel("all-lipid-totals-Hartley-70brain (edited)- removed 3 samples.xlsx",
                             sheet="8signal-per-mg-drytissue-wt ave", col_names = FALSE)

colnames(Week6_18_Brain) <- Week6_18_Brain[3,]

merger <- read_excel("all-lipid-totals-Hartley-70brain (edited)- removed 3 samples.xlsx",
                     sheet="Additional sample info", range = "A19:G100", .name_repair = "universal", )
merger <- drop_na(merger,)

### Total ########
# To add labels for merging
CleanData_6B_Total <- Week6_18_Brain %>% select(Mass, CoV, contains("Sample"))
supplemental_6B_T <- CleanData_6B_Total[1,]

#Filter for only total values and CoV <0.3
CleanData_6B_Total <- CleanData_6B_Total %>% filter(grepl('Total', Mass))
CleanData_6B_Total$CoV <- as.numeric(CleanData_6B_Total$CoV)
CleanData_6B_Total <- CleanData_6B_Total %>% filter(CoV < 0.3)

# Drop final row 
CleanData_6B_Total <- subset(CleanData_6B_Total, Mass != "Total")

# Bind supplemental for column names, and drop columns
Merge_6B_Total <- rbind(CleanData_6B_Total, supplemental_6B_T) %>% select(-CoV)

#### Transpose & Merge####
Transpose_6B_Total <- transpose(Merge_6B_Total)
# Fix column and row names
Transpose_6B_Total <- Transpose_6B_Total %>% row_to_names(row_number = 1)
Transpose_6B_Total <- mutate_all(Transpose_6B_Total, function(x) as.numeric(as.character(x)))

# Joining dataframe with another through a matching "key" variable.
names(Transpose_6B_Total)[13] <- "Sample.number"
Final_6B_Total <- Transpose_6B_Total %>% left_join(merger,by = "Sample.number" )

# Combine 
Final_6B_Total$CRE.and.Treatment <- paste(Final_6B_Total$genotype,Final_6B_Total$Treatment,sep="/")
Final_6B_Total <- Final_6B_Total %>% filter(!grepl('260-3943|260-3944|261-3947', Sample.name))

# drop columns
Final_6B_Total_CE <- Final_6B_Total %>% select(-"Sample.name",-"dry.extracted.tissue.wt",-"genotype",-"Treatment",-"Dry.lipid.wt")
Final_6B_Total_noCE <- Final_6B_Total %>% select(-"Total CholEster",-"Sample.name",-"dry.extracted.tissue.wt",-"genotype",-"Treatment",-"Dry.lipid.wt")

write_csv(Final_6B_Total_CE, "Final_6B_Total.csv")
write_csv(Final_6B_Total_noCE, "Final_6B_Total_noCE.csv")
# read with rownames(df) <- df$...1

### Individual ####

# To add labels for merging
CleanData_6B_All <- Week6_18_Brain %>% select(Mass, CoV, "Compound Name",contains("Sample")) 
supplemental_6B_A <- CleanData_6B_All[1,]
#Filter out total values
CleanData_6B_All <- CleanData_6B_All %>% filter(grepl(':', CleanData_6B_All$`Compound Name`))
CleanData_6B_All <- drop_na(CleanData_6B_All,)

# Remove all rows with CoV <0.3
CleanData_6B_All$CoV <- as.numeric(CleanData_6B_All$CoV)
CleanData_6B_All <- CleanData_6B_All %>% filter(CoV < 0.3) 

# Bind supplemental labels for merging
rowname <- CleanData_6B_All$"Compound Name"
#CleanData_All <- CleanData_All %>% select(-"Compound Name")
Merge_6B_All <- rbind(CleanData_6B_All, supplemental_6B_A) 
Merge_6B_All$'Compound Name' <- Merge_6B_All$'Compound Name' %>% replace_na('Sample.number')

# Drop CoV column
Merge_6B_All <- Merge_6B_All %>% select(-CoV,-Mass)

# Fix row index
#rowname <- append(rowname, "Sample.number")
rownames(Merge_6B_All) <- Merge_6B_All$'Compound Name'

#### Transpose & Merge ####
Transpose_6B_All <- transpose(Merge_6B_All)
# Fix column and row names
rownames(Transpose_6B_All) <- colnames(Merge_6B_All)
Transpose_6B_All <- Transpose_6B_All %>% row_to_names(row_number = 1)

# Merge dataframes for CRE status through a matching "key" variable.
Transpose_6B_All <- mutate_all(Transpose_6B_All, function(x) as.numeric(as.character(x)))
Final_6B_All <- Transpose_6B_All %>% left_join(merger,by = "Sample.number" )
Final_6B_All$CRE.and.Treatment <- paste(Final_6B_All$genotype,Final_6B_All$Treatment,sep="/")
Final_6B_All <- Final_6B_All %>% filter(!grepl('260-3943|260-3944|261-3947', Sample.name))

Final_6B_All_CE <- Final_6B_All %>% select(-"Sample.name",-"dry.extracted.tissue.wt",-"genotype",-"Treatment",-"Dry.lipid.wt")
Final_6B_All_noCE <- Final_6B_All %>% select(-contains("CE"),-"Sample.name",-"dry.extracted.tissue.wt",-"genotype",-"Treatment",-"Dry.lipid.wt")

write_csv(Final_6B_All_CE, "Final_6B_All.csv")
write_csv(Final_6B_All_noCE, "Final_6B_All_noCE.csv")
# read with rownames(df) <- df$...1

## PLP Spinal 6 - 18 ####

Week6_18_Spinal <- read_excel("all-lipid-totals-71HartleySpinal101220-v20.2 (1).xlsx",
                              sheet="8signal-per-mg-tissue wt ave", col_names = FALSE)

colnames(Week6_18_Spinal) <- Week6_18_Spinal[3,]

merger <- read_excel("all-lipid-totals-71HartleySpinal101220-v20.2 (1).xlsx",
                     sheet="KLRC data", range = "A34:G106", .name_repair = "universal", )

merger <- drop_na(merger,)

### Total ####

# To add labels for merging
CleanData_6S_Total <- Week6_18_Spinal %>% select(Mass, CoV, contains("sample"))
supplemental_6S_T <- CleanData_6S_Total[2,]

#Filter for only total values and CoV <0.3
CleanData_6S_Total <- CleanData_6S_Total %>% filter(grepl('Total', Mass))
CleanData_6S_Total$CoV <- as.numeric(CleanData_6S_Total$CoV)
CleanData_6S_Total <- CleanData_6S_Total %>% filter(CoV < 0.3)

# Drop final row 
CleanData_6S_Total <- subset(CleanData_6S_Total, Mass != "Total")

# Bind supplemental for column names, and drop columns
Merge_6S_Total <- rbind(CleanData_6S_Total, supplemental_6S_T) %>% select(-CoV)
Merge_6S_Total$Mass <- Merge_6S_Total$Mass %>% replace_na('...1')

#### Transpose & Merger ####

Transpose_6S_Total <- transpose(Merge_6S_Total)
Transpose_6S_Total <- Transpose_6S_Total %>% row_to_names(row_number = 1)
#Transpose_6S_Total <- mutate_all(Transpose_6S_Total, function(x) as.numeric(as.character(x)))

# Joining dataframe with another through a matching "key" variable.
Final_6S_Total <- Transpose_6S_Total %>% left_join(merger,by = "...1" )

# Combine 
Final_6S_Total$CRE.and.Treatment <- paste(Final_6S_Total$genotype,Final_6S_Total$Treatment,sep="/")
Final_6S_Total <- Final_6S_Total %>% filter(!grepl('260-3943|260-3944|261-3947', Sample.name))

Final_6S_Total_CE <- Final_6S_Total %>% select(-"Sample.name",-"dry.extracted.tissue.wt",-"genotype",-"Treatment",-"Dry.lipid.wt")
Final_6S_Total_noCE <- Final_6S_Total %>% select(-"Total CholEster",-"Sample.name",-"dry.extracted.tissue.wt",-"genotype",-"Treatment",-"Dry.lipid.wt")

write_csv(Final_6S_Total_CE, "Final_6S_Total.csv")
write_csv(Final_6S_Total_noCE, "Final_6S_Total_noCE.csv")
# read with rownames(df) <- df$...1


### Individual ####

# To add labels for merging
CleanData_6S_All <- Week6_18_Spinal %>% select(Mass, CoV, "Compound Name",contains("sample")) 
supplemental_6S_A <- CleanData_6S_All[2,]
#Filter out total values
CleanData_6S_All <- CleanData_6S_All %>% filter(grepl(':', CleanData_6S_All$`Compound Name`))
CleanData_6S_All <- drop_na(CleanData_6S_All,)

# Remove all rows with CoV <0.3
CleanData_6S_All$CoV <- as.numeric(CleanData_6S_All$CoV)
CleanData_6S_All <- CleanData_6S_All %>% filter(CoV < 0.3) 

# Bind supplemental labels for merging
Merge_6S_All <- rbind(CleanData_6S_All, supplemental_6S_A) 
Merge_6S_All$'Compound Name' <- Merge_6S_All$'Compound Name' %>% replace_na('...1')

# Drop CoV column
Merge_6S_All <- Merge_6S_All %>% select(-CoV,-Mass)

# Fix row index
rownames(Merge_6S_All) <- Merge_6S_All$'Compound Name'

#### Transpose & Merge ####
Transpose_6S_All <- transpose(Merge_6S_All)
# Fix column and row names
rownames(Transpose_6S_All) <- colnames(Merge_6S_All)
Transpose_6S_All <- Transpose_6S_All %>% row_to_names(row_number = 1)

# Merge dataframes for CRE status through a matching "key" variable.
#Transpose_6S_All <- mutate_all(Transpose_6S_All, function(x) as.numeric(as.character(x)))
Final_6S_All <- Transpose_6S_All %>% left_join(merger,by = "...1" )
Final_6S_All$CRE.and.Treatment <- paste(Final_6S_All$genotype,Final_6S_All$Treatment,sep="/")
Final_6S_All <- Final_6S_All %>% filter(!grepl('260-3943|260-3944|261-3947', Sample.name))

Final_6S_All_CE <- Final_6S_All %>% select(-"Sample.name",-"dry.extracted.tissue.wt",-"genotype",-"Treatment",-"Dry.lipid.wt")
Final_6S_All_noCE <- Final_6S_All %>% select(-contains("CE"),-"Sample.name",-"dry.extracted.tissue.wt",-"genotype",-"Treatment",-"Dry.lipid.wt")

write_csv(Final_6S_All_CE, "Final_6S_All.csv")
write_csv(Final_6S_All_noCE, "Final_6S_All_noCE.csv")
# read with rownames(df) <- df$...1

## PLP Serum 6 - 18 ####

Week6_18_Serum <- read_excel("all-lipid-totals-70Hartley012420-v19.1-serum-wks 6,12,18.xlsx",
                             sheet="signal-per-ul ave", col_names = FALSE)

colnames(Week6_18_Serum) <- Week6_18_Serum[3,]

merger <- read_excel("all-lipid-totals-70Hartley012420-v19.1-serum-wks 6,12,18.xlsx",
                     sheet="KLRC data", range = "A34:F104", .name_repair = "universal", )
merger <- drop_na(merger,)

### Total ########
# To add labels for merging
CleanData_6Srm_Total <- Week6_18_Serum %>% select(Mass, CoV, contains("Sample"))
supplemental_6Srm_T <- CleanData_6Srm_Total[2,]

#Filter for only total values and CoV <0.3
CleanData_6Srm_Total <- CleanData_6Srm_Total %>% filter(grepl('Total ', Mass))
CleanData_6Srm_Total$CoV <- as.numeric(CleanData_6Srm_Total$CoV)
CleanData_6Srm_Total <- CleanData_6Srm_Total %>% filter(CoV < 0.3)

# Drop final row 
#CleanData_6Srm_Total <- subset(CleanData_6Srm_Total, Mass != "Total")

# Bind supplemental for column names, and drop columns
Merge_6Srm_Total <- rbind(CleanData_6Srm_Total, supplemental_6Srm_T) %>% select(-CoV)

#### Transpose & Merge####
Transpose_6Srm_Total <- transpose(Merge_6Srm_Total)
# Fix column and row names
Transpose_6Srm_Total <- Transpose_6Srm_Total %>% row_to_names(row_number = 1)
names(Transpose_6Srm_Total)[11] <- "Sample.name"
#Transpose_6Srm_Total <- mutate_all(Transpose_6Srm_Total, function(x) as.numeric(as.character(x)))
variables <- c('Total LysoPC', 'Total PC','Total SM and DSM','Total ePC','Total LysoPE','Total PE',
               'Total ePE','Total PI','Total PS','Total CholEster')
Transpose_6Srm_Total <- Transpose_6Srm_Total %>% mutate_at(variables, as.numeric)
# Joining dataframe with another through a matching "key" variable.
Final_6Srm_Total <- Transpose_6Srm_Total %>% left_join(merger,by = "Sample.name" )

# Combine 
Final_6Srm_Total$CRE.and.Treatment <- paste(Final_6Srm_Total$genotype,Final_6Srm_Total$Treatment,sep="/")
Final_6Srm_Total <- Final_6Srm_Total %>% filter(!grepl('260-3943|260-3944|261-3947', Sample.name))

# drop columns
Final_6Srm_Total_CE <- Final_6Srm_Total %>% select(-"Sample.name",-"Units.",-"genotype",-"Treatment")
Final_6Srm_Total_noCE <- Final_6Srm_Total %>% select(-"Total CholEster",-"Units.",-"Sample.name",-"genotype",-"Treatment")

write_csv(Final_6Srm_Total_CE, "Final_6Srm_Total.csv")
write_csv(Final_6Srm_Total_noCE, "Final_6Srm_Total_noCE.csv")
# read with rownames(df) <- df$...1

### Individual ####

# To add labels for merging
CleanData_6Srm_All <- Week6_18_Serum %>% select(Mass, CoV, "Compound Name",contains("Sample")) 
supplemental_6Srm_A <- CleanData_6Srm_All[2,]
#Filter out total values
CleanData_6Srm_All <- CleanData_6Srm_All %>% filter(grepl(':', CleanData_6Srm_All$`Compound Name`))
CleanData_6Srm_All <- drop_na(CleanData_6Srm_All,)

# Remove all rows with CoV <0.3
CleanData_6Srm_All$CoV <- as.numeric(CleanData_6Srm_All$CoV)
CleanData_6Srm_All <- CleanData_6Srm_All %>% filter(CoV < 0.3) 

# Bind supplemental labels for merging
rowname <- CleanData_6Srm_All$"Compound Name"
#CleanData_All <- CleanData_All %>% select(-"Compound Name")
Merge_6Srm_All <- rbind(CleanData_6Srm_All, supplemental_6Srm_A) 
Merge_6Srm_All$'Compound Name' <- Merge_6Srm_All$'Compound Name' %>% replace_na('Sample.name')

# Drop CoV column
Merge_6Srm_All <- Merge_6Srm_All %>% select(-CoV,-Mass)

# Fix row index
#rowname <- append(rowname, "Sample.number")
rownames(Merge_6Srm_All) <- Merge_6Srm_All$'Compound Name'

#### Transpose & Merge ####
Transpose_6Srm_All <- transpose(Merge_6Srm_All)
# Fix column and row names
rownames(Transpose_6Srm_All) <- colnames(Merge_6Srm_All)
Transpose_6Srm_All <- Transpose_6Srm_All %>% row_to_names(row_number = 1)
# Merge dataframes for CRE status through a matching "key" variable.
Transpose_6Srm_All <- Transpose_6Srm_All %>% mutate_at(rowname, as.numeric)
Final_6Srm_All <- Transpose_6Srm_All %>% left_join(merger,by = "Sample.name" )
Final_6Srm_All$CRE.and.Treatment <- paste(Final_6Srm_All$genotype,Final_6Srm_All$Treatment,sep="/")
Final_6Srm_All <- Final_6Srm_All %>% filter(!grepl('260-3943|260-3944|261-3947', Sample.name))

Final_6Srm_All_CE <- Final_6Srm_All %>% select(-"Sample.name",-"genotype",-"Treatment",-"Units.")
Final_6Srm_All_noCE <- Final_6Srm_All %>% select(-contains("CE"),-"Sample.name",-"genotype",-"Treatment",-"Units.")

write_csv(Final_6Srm_All_CE, "Final_6Srm_All.csv")
write_csv(Final_6Srm_All_noCE, "Final_6Srm_All_noCE.csv")
# read with rownames(df) <- df$...1


## PLP Brain 24 ####

Week24_Brain <- read_excel("all-lipid-totals-Hartley-26brainMyelin-MRR062321.xlsx",
                           sheet="7B signal-per-mg-dry tiss ave", col_names = FALSE)

colnames(Week24_Brain) <- Week24_Brain[3,]

merger <- read_excel("all-lipid-totals-Hartley-26brainMyelin-MRR062321.xlsx",
                     sheet="KLRC data", range = "A34:F48", .name_repair = "universal", )
merger <- drop_na(merger,)

### Total ########
# To add labels for merging
CleanData_24B_Total <- Week24_Brain %>% select(Mass, CoV, contains("sample"))
supplemental_24B_T <- CleanData_24B_Total[2,]

#Filter for only total values and CoV <0.3
CleanData_24B_Total <- CleanData_24B_Total %>% filter(grepl('Total', Mass))
CleanData_24B_Total$CoV <- as.numeric(CleanData_24B_Total$CoV)
CleanData_24B_Total <- CleanData_24B_Total %>% filter(CoV < 0.3)

# Drop final row 
CleanData_24B_Total <- subset(CleanData_24B_Total, Mass != "Total")

# Bind supplemental for column names, and drop columns
Merge_24B_Total <- rbind(CleanData_24B_Total, supplemental_24B_T) %>% select(-CoV)
Merge_24B_Total$Mass <- Merge_24B_Total$Mass %>% replace_na('...1')

#### Transpose & Merger ####

Transpose_24B_Total <- transpose(Merge_24B_Total)
Transpose_24B_Total <- Transpose_24B_Total %>% row_to_names(row_number = 1)
#Transpose_6S_Total <- mutate_all(Transpose_6S_Total, function(x) as.numeric(as.character(x)))

# Joining dataframe with another through a matching "key" variable.
Final_24B_Total <- Transpose_24B_Total %>% left_join(merger,by = "...1" )

# Combine 
Final_24B_Total$CRE.and.Treatment <- paste(Final_24B_Total$genotype,Final_24B_Total$Treatment,sep="/")
Final_24B_Total <- Final_24B_Total %>% filter(!grepl('140-653|198-3662', Sample.name))


Final_24B_Total_CE <- Final_24B_Total %>% select(-"Sample.name",-"dry.extracted.tissue.wt",-"genotype",-"Treatment")
Final_24B_Total_noCE <- Final_24B_Total %>% select(-"Total CholEster",-"Sample.name",-"dry.extracted.tissue.wt",-"genotype",-"Treatment")

write_csv(Final_24B_Total_CE, "Final_24B_Total.csv")
write_csv(Final_24B_Total_noCE, "Final_24B_Total_noCE.csv")
# read with rownames(df) <- df$...1

### Individual ####

# To add labels for merging
CleanData_24B_All <- Week24_Brain %>% select(Mass, CoV, "Compound Name",contains("sample")) 
supplemental_24B_A <- CleanData_24B_All[2,]
#Filter out total values
CleanData_24B_All <- CleanData_24B_All %>% filter(grepl(':', CleanData_24B_All$`Compound Name`))
CleanData_24B_All <- drop_na(CleanData_24B_All,)

# Remove all rows with CoV <0.3
CleanData_24B_All$CoV <- as.numeric(CleanData_24B_All$CoV)
CleanData_24B_All <- CleanData_24B_All %>% filter(CoV < 0.3) 

# Bind supplemental labels for merging
Merge_24B_All <- rbind(CleanData_24B_All, supplemental_24B_A) 
Merge_24B_All$'Compound Name' <- Merge_24B_All$'Compound Name' %>% replace_na('...1')

# Drop CoV column
Merge_24B_All <- Merge_24B_All %>% select(-CoV,-Mass)

# Fix row index
rownames(Merge_24B_All) <- Merge_24B_All$'Compound Name'

#### Transpose & Merge ####

Transpose_24B_All <- transpose(Merge_24B_All)
# Fix column and row names
rownames(Transpose_24B_All) <- colnames(Merge_24B_All)
Transpose_24B_All <- Transpose_24B_All %>% row_to_names(row_number = 1)

# Merge dataframes for CRE status through a matching "key" variable.
#Transpose_24B_All <- mutate_all(Transpose_24B_All, function(x) as.numeric(as.character(x)))
Final_24B_All <- Transpose_24B_All %>% left_join(merger,by = "...1" )
Final_24B_All$CRE.and.Treatment <- paste(Final_24B_All$genotype,Final_24B_All$Treatment,sep="/")
Final_24B_All <- Final_24B_All %>% filter(!grepl('140-653|198-3662', Sample.name))


Final_24B_All_CE <- Final_24B_All %>% select(-"Sample.name",-"dry.extracted.tissue.wt",-"genotype",-"Treatment")
Final_24B_All_noCE <- Final_24B_All %>% select(-contains("CE"),-"Sample.name",-"dry.extracted.tissue.wt",-"genotype",-"Treatment")

write_csv(Final_24B_All_CE, "Final_24B_All.csv")
write_csv(Final_24B_All_noCE, "Final_24B_All_noCE.csv")
# read with rownames(df) <- df$...1


## PLP Spinal 24 ####

Week24_Spinal <- read_excel("all-lipid-totals-Hartley-22spinal112020-2-v20.2-normalization_included.xlsx",
                           sheet="7b norm-sig-p-mg-tissue-wt ave", col_names = FALSE)

colnames(Week24_Spinal) <- Week24_Spinal[3,]
# remove 140-653 & 198-3662 sample from the data set

merger <- read_excel("all-lipid-totals-Hartley-22spinal112020-2-v20.2-normalization_included.xlsx",
                     sheet="KLRC data", range = "A34:F57", .name_repair = "universal", )
merger <- drop_na(merger,)

### Total ########
# To add labels for merging
#CleanData_24S_Total <- Week24_Spinal %>% select(-contains(c("ave","stdev")),-21)
CleanData_24S_Total <- Week24_Spinal %>% select(Mass, Cov, contains("sample"),-21)
supplemental_24S_T <- CleanData_24S_Total[2,]

#Filter for only total values and CoV <0.3
CleanData_24S_Total <- CleanData_24S_Total %>% filter(grepl('Total', Mass))
CleanData_24S_Total$Cov <- as.numeric(CleanData_24S_Total$Cov)
CleanData_24S_Total <- CleanData_24S_Total %>% filter(Cov < 0.3)

# Drop final row 
CleanData_24S_Total <- subset(CleanData_24S_Total, Mass != "Total")

# Bind supplemental for column names, and drop columns
Merge_24S_Total <- rbind(CleanData_24S_Total, supplemental_24S_T) %>% select(-Cov)
Merge_24S_Total$Mass <- Merge_24S_Total$Mass %>% replace_na('...1')

#### Transpose & Merger ####

Transpose_24S_Total <- transpose(Merge_24S_Total)
Transpose_24S_Total <- Transpose_24S_Total %>% row_to_names(row_number = 1)
#Transpose_24S_Total <- mutate_all(Transpose_24S_Total, function(x) as.numeric(as.character(x)))

# Joining dataframe with another through a matching "key" variable.
Final_24S_Total <- Transpose_24S_Total %>% left_join(merger,by = "...1" )
Final_24S_Total <- Final_24S_Total %>% filter(grepl('wk 24', ...1))

# Combine 
Final_24S_Total$CRE.and.Treatment <- paste(Final_24S_Total$genotype,Final_24S_Total$Treatment,sep="/")
# remove sample 79 as well that has too many values (only applies for spinal cords)
Final_24S_Total <- Final_24S_Total %>% filter(!grepl('140-653|198-3662|197-3659', Sample.name))

Final_24S_Total_CE <- Final_24S_Total %>% select(-"Sample.name",-"dry.extracted.tissue.wt",-"genotype",-"Treatment")
Final_24S_Total_noCE <- Final_24S_Total %>% select(-"Total CholEster",-"Sample.name",-"dry.extracted.tissue.wt",-"genotype",-"Treatment")

write_csv(Final_24S_Total_CE, "Final_24S_Total.csv")
write_csv(Final_24S_Total_noCE, "Final_24S_Total_noCE.csv")
# read with rownames(df) <- df$...1


### Individual ####

# To add labels for merging
#CleanData_24S_All <- Week24_Spinal %>% select(-contains(c("ave","stdev")),-21)
CleanData_24S_All <- Week24_Spinal %>% select(Mass, Cov, "Compound Name", contains("sample"),-21)
#CleanData_24S_All <- CleanData_24S_All %>% select(Mass, Cov, "Compound Name",contains("sample")) 
supplemental_24S_A <- CleanData_24S_All[2,]
#Filter out total values
CleanData_24S_All <- CleanData_24S_All %>% filter(grepl(':', CleanData_24S_All$`Compound Name`))
CleanData_24S_All <- drop_na(CleanData_24S_All,)

# Remove all rows with Cov <0.3
CleanData_24S_All$Cov <- as.numeric(CleanData_24S_All$Cov)
CleanData_24S_All <- CleanData_24S_All %>% filter(Cov < 0.3) 

# Bind supplemental labels for merging
Merge_24S_All <- rbind(CleanData_24S_All, supplemental_24S_A) 
Merge_24S_All$'Compound Name' <- Merge_24S_All$'Compound Name' %>% replace_na('...1')

# Drop Cov column
Merge_24S_All <- Merge_24S_All %>% select(-Cov,-Mass)

# Fix row index
rownames(Merge_24S_All) <- Merge_24S_All$'Compound Name'

#### Transpose & Merge ####
Transpose_24S_All <- transpose(Merge_24S_All)
# Fix column and row names
rownames(Transpose_24S_All) <- colnames(Merge_24S_All)
Transpose_24S_All <- Transpose_24S_All %>% row_to_names(row_number = 1)

# Merge dataframes for CRE status through a matching "key" variable.
#Transpose_24S_All <- mutate_all(Transpose_24S_All, function(x) as.numeric(as.character(x)))
Final_24S_All <- Transpose_24S_All %>% left_join(merger,by = "...1" )
Final_24S_All <- Final_24S_All %>% filter(grepl('wk 24', ...1))

Final_24S_All$CRE.and.Treatment <- paste(Final_24S_All$genotype,Final_24S_All$Treatment,sep="/")
# remove sample 79 as well that has too many values (only applies for spinal cords)
Final_24S_All <- Final_24S_All %>% filter(!grepl('140-653|198-3662|197-3659', Sample.name))


Final_24S_All_CE <- Final_24S_All %>% select(-"Sample.name",-"dry.extracted.tissue.wt",-"genotype",-"Treatment")
Final_24S_All_noCE <- Final_24S_All %>% select(-contains("CE"),-"Sample.name",-"dry.extracted.tissue.wt",-"genotype",-"Treatment")

write_csv(Final_24S_All_CE, "Final_24S_All.csv")
write_csv(Final_24S_All_noCE, "Final_24S_All_noCE.csv")
# read with rownames(df) <- df$...1


## Myelin Isolation ####

Week24_Myelin<- read_excel("~/Library/CloudStorage/OneDrive-TheUniversityofKansas/Hartley Lab Material/2022/PCA/all-lipid-totals-Hartley-26brainMyelin-MRR062321.xlsx",
                           sheet="7 signal-per-mg-protein ave", col_names = FALSE)

colnames(Week24_Myelin) <- Week24_Myelin[3,]

merger <- read_excel("~/Library/CloudStorage/OneDrive-TheUniversityofKansas/Hartley Lab Material/2022/PCA/all-lipid-totals-Hartley-26brainMyelin-MRR062321.xlsx",
                     sheet="KLRC data", range = "A49:F63", .name_repair = "universal", )
merger <- merger %>% select(-4)
merger <- drop_na(merger,)

### Total ########
# To add labels for merging
CleanData_24M_Total <- Week24_Myelin %>% select(Mass, CoV, contains("SampleM"))
supplemental_24M_T <- CleanData_24M_Total[2,]

#Filter for only total values and CoV <0.3
CleanData_24M_Total <- CleanData_24M_Total %>% filter(grepl('Total', Mass))
CleanData_24M_Total$CoV <- as.numeric(CleanData_24M_Total$CoV)
CleanData_24M_Total <- CleanData_24M_Total %>% filter(CoV < 0.3)

# Drop final row 
CleanData_24M_Total <- subset(CleanData_24M_Total, Mass != "Total")

# Bind supplemental for column names, and drop columns
Merge_24M_Total <- rbind(CleanData_24M_Total, supplemental_24M_T) %>% select(-CoV)
Merge_24M_Total$Mass <- Merge_24M_Total$Mass %>% replace_na('...1')

#### Transpose & Merger ####

Transpose_24M_Total <- transpose(Merge_24M_Total)
Transpose_24M_Total <- Transpose_24M_Total %>% row_to_names(row_number = 1)
#Transpose_6S_Total <- mutate_all(Transpose_6S_Total, function(x) as.numeric(as.character(x)))

# Joining dataframe with another through a matching "key" variable.
Final_24M_Total <- Transpose_24M_Total %>% left_join(merger,by = "...1" )

# Combine 
Final_24M_Total$CRE.and.Treatment <- paste(Final_24M_Total$genotype,Final_24M_Total$Treatment,sep="/")
Final_24M_Total <- Final_24M_Total %>% filter(!grepl('140-653|198-3662', Sample.name))

Final_24M_Total_CE <- Final_24M_Total %>% select(-"Sample.name",-"genotype",-"Treatment")
Final_24M_Total_noCE <- Final_24M_Total %>% select(-"Total CholEster",-"Sample.name",-"genotype",-"Treatment")

write_csv(Final_24M_Total_CE, "Final_24M_Total.csv")
write_csv(Final_24M_Total_noCE, "Final_24M_Total_noCE.csv")
# read with rownames(df) <- df$...1

### Individual ####

# To add labels for merging
CleanData_24M_All <- Week24_Myelin %>% select(Mass, CoV, "Compound Name",contains("sampleM")) 
supplemental_24M_A <- CleanData_24M_All[2,]
#Filter out total values
CleanData_24M_All <- CleanData_24M_All %>% filter(grepl(':', CleanData_24M_All$`Compound Name`))
CleanData_24M_All <- drop_na(CleanData_24M_All,)

# Remove all rows with CoV <0.3
CleanData_24M_All$CoV <- as.numeric(CleanData_24M_All$CoV)
CleanData_24M_All <- CleanData_24M_All %>% filter(CoV < 0.3) 

# Bind supplemental labels for merging
Merge_24M_All <- rbind(CleanData_24M_All, supplemental_24M_A) 
Merge_24M_All$'Compound Name' <- Merge_24M_All$'Compound Name' %>% replace_na('...1')

# Drop CoV column
Merge_24M_All <- Merge_24M_All %>% select(-CoV,-Mass)

# Fix row index
rownames(Merge_24M_All) <- Merge_24M_All$'Compound Name'

#### Transpose & Merge ####

Transpose_24M_All <- transpose(Merge_24M_All)
# Fix column and row names
rownames(Transpose_24M_All) <- colnames(Merge_24M_All)
Transpose_24M_All <- Transpose_24M_All %>% row_to_names(row_number = 1)

# Merge dataframes for CRE status through a matching "key" variable.
#Transpose_24M_All <- mutate_all(Transpose_24M_All, function(x) as.numeric(as.character(x)))
Final_24M_All <- Transpose_24M_All %>% left_join(merger,by = "...1" )
Final_24M_All$CRE.and.Treatment <- paste(Final_24M_All$genotype,Final_24M_All$Treatment,sep="/")

Final_24M_All <- Final_24M_All %>% filter(!grepl('140-653|198-3662', Sample.name))

Final_24M_All_CE <- Final_24M_All %>% select(-"Sample.name",-"genotype",-"Treatment")
Final_24M_All_noCE <- Final_24M_All %>% select(-contains("CE"),-"Sample.name",-"genotype",-"Treatment")

write_csv(Final_24M_All_CE, "Final_24M_All.csv")
write_csv(Final_24M_All_noCE, "Final_24M_All_noCE.csv")
# read with rownames(df) <- df$...1

## Sox10 Brain 6 - 9 ####

SoxWeek6_18_Brain <- read_excel("~/Library/CloudStorage/OneDrive-TheUniversityofKansas/Hartley Lab Material/2022/PCA/all-lipid-totals-Hartley-BRAIN102322+CE-2-v21.1 (1).xlsx",
                             sheet="8 signal per mg tissue wt ave", col_names = FALSE)

colnames(SoxWeek6_18_Brain) <- SoxWeek6_18_Brain[3,]

merger <- read_excel("~/Library/CloudStorage/OneDrive-TheUniversityofKansas/Hartley Lab Material/2022/PCA/all-lipid-totals-Hartley-BRAIN102322+CE-2-v21.1 (1).xlsx",
                     sheet="KLRC data", range = "A34:F80", .name_repair = "universal", )
merger <- drop_na(merger,)

### Total ########
# To add labels for merging
CleanData_6B_Total <- SoxWeek6_18_Brain %>% select(Mass, CoV, contains("B_"))
supplemental_6B_T <- CleanData_6B_Total[2,]

#Filter for only total values and CoV <0.3
CleanData_6B_Total <- CleanData_6B_Total %>% filter(grepl('Total', Mass))
CleanData_6B_Total$CoV <- as.numeric(CleanData_6B_Total$CoV)
CleanData_6B_Total <- CleanData_6B_Total %>% filter(CoV < 0.3)

# Drop final row 
CleanData_6B_Total <- subset(CleanData_6B_Total, Mass != "Total")

# Bind supplemental for column names, and drop columns
Merge_6B_Total <- rbind(CleanData_6B_Total, supplemental_6B_T) %>% select(-CoV)

#### Transpose & Merge####
Transpose_6B_Total <- transpose(Merge_6B_Total)

# Fix column and row names
Transpose_6B_Total <- Transpose_6B_Total %>% row_to_names(row_number = 1)
names(Transpose_6B_Total)[14] <- "...1"
Transpose_6B_Total <- Transpose_6B_Total %>% mutate_at(c('Total LysoPC', 'Total PC', 'Total SM and DSM', 'Total ePC', 'Total LysoPE', 'Total PE', 
                   'Total ePE', 'Total PI', 'Total PS', 'Total ePS', 'Total PA', 'Total PG', 'Total CholEsters'), as.numeric)
#Transpose_6B_Total <- mutate_all(Transpose_6B_Total, function(x) as.numeric(as.character(x)))

# Joining dataframe with another through a matching "key" variable.
#names(Transpose_6B_Total)[13] <- "Sample.number"
Final_6B_Total <- Transpose_6B_Total %>% left_join(merger,by = "...1" )

# Combine 
Final_6B_Total$CRE.and.Treatment <- paste(Final_6B_Total$genotype,Final_6B_Total$Treatment,sep="/")
Final_6B_Total <- Final_6B_Total %>% filter(!grepl('86|87', Sample.number))

# drop columns
Final_6B_Total_CE <- Final_6B_Total %>% select(-"Sample.name",-"dry.extracted.tissue.wt",-"genotype",-"Treatment")
Final_6B_Total_noCE <- Final_6B_Total %>% select(-"Total CholEsters",-"Sample.name",-"dry.extracted.tissue.wt",-"genotype",-"Treatment")

write_csv(Final_6B_Total_CE, "Sox10_Final_6B_Total.csv")
write_csv(Final_6B_Total_noCE, "Sox10_Final_6B_Total_noCE.csv")
# read with rownames(df) <- df$...1

### Individual ####

# To add labels for merging
CleanData_6B_All <- SoxWeek6_18_Brain %>% select(Mass, CoV, "Compound Name", contains("B_")) 
supplemental_6B_A <- CleanData_6B_All[2,]

#Filter out total values
CleanData_6B_All <- CleanData_6B_All %>% filter(grepl(':', CleanData_6B_All$`Compound Name`))
CleanData_6B_All <- drop_na(CleanData_6B_All,)

# Remove all rows with CoV <0.3
CleanData_6B_All$CoV <- as.numeric(CleanData_6B_All$CoV)
CleanData_6B_All <- CleanData_6B_All %>% filter(CoV < 0.3) 

# Bind supplemental labels for merging
rowname <- CleanData_6B_All$"Compound Name"
#CleanData_All <- CleanData_All %>% select(-"Compound Name")
Merge_6B_All <- rbind(CleanData_6B_All, supplemental_6B_A) 
Merge_6B_All$'Compound Name' <- Merge_6B_All$'Compound Name' %>% replace_na('...1')

# Drop CoV column
Merge_6B_All <- Merge_6B_All %>% select(-CoV,-Mass)

# Fix row index
#rowname <- append(rowname, "Sample.number")
rownames(Merge_6B_All) <- Merge_6B_All$'Compound Name'

#### Transpose & Merge ####
Transpose_6B_All <- transpose(Merge_6B_All)
# Fix column and row names
rownames(Transpose_6B_All) <- colnames(Merge_6B_All)
Transpose_6B_All <- Transpose_6B_All %>% row_to_names(row_number = 1)

# Merge dataframes for CRE status through a matching "key" variable.
#Transpose_6B_All <- mutate_all(Transpose_6B_All, function(x) as.numeric(as.character(x)))
Transpose_6B_All <- Transpose_6B_All %>% mutate_at((1:170), as.numeric)
str(Transpose_6B_All$...1)
Final_6B_All <- Transpose_6B_All %>% left_join(merger,by = "...1" )
Final_6B_All$CRE.and.Treatment <- paste(Final_6B_All$genotype,Final_6B_All$Treatment,sep="/")
Final_6B_All <- Final_6B_All %>% filter(!grepl('86|87', Sample.number))

Final_6B_All_CE <- Final_6B_All %>% select(-"Sample.name",-"dry.extracted.tissue.wt",-"genotype",-"Treatment")
Final_6B_All_noCE <- Final_6B_All %>% select(-(157:170),-"Sample.name",-"dry.extracted.tissue.wt",-"genotype",-"Treatment")

write_csv(Final_6B_All_CE, "Sox10_Final_6B_All.csv")
write_csv(Final_6B_All_noCE, "Sox10_Final_6B_All_noCE.csv")
# read with rownames(df) <- df$...1

## Sox10 Spinal 6 - 9 ####

SoxWeek6_18_Spinal <- read_excel("all-lipid-totals-Hartley-SPINAL102222+CE-2-v21.1 (1).xlsx",
                              sheet="8 signal per mg tissue wt ave", col_names = FALSE)

colnames(SoxWeek6_18_Spinal) <- SoxWeek6_18_Spinal[3,]

merger <- read_excel("~/Library/CloudStorage/OneDrive-TheUniversityofKansas/Hartley Lab Material/2022/PCA/all-lipid-totals-Hartley-SPINAL102222+CE-2-v21.1 (1).xlsx",
                     sheet="KLRC data", range = "A34:G80", .name_repair = "universal", )

merger <- drop_na(merger,)

### Total ####

# To add labels for merging
CleanData_6S_Total <- SoxWeek6_18_Spinal %>% select(Mass, CoV, contains("S_"))
supplemental_6S_T <- CleanData_6S_Total[2,]

#Filter for only total values and CoV <0.3
CleanData_6S_Total <- CleanData_6S_Total %>% filter(grepl('Total', Mass))
CleanData_6S_Total$CoV <- as.numeric(CleanData_6S_Total$CoV)
CleanData_6S_Total <- CleanData_6S_Total %>% filter(CoV < 0.3)

# Drop final row 
CleanData_6S_Total <- subset(CleanData_6S_Total, Mass != "Total")

# Bind supplemental for column names, and drop columns
Merge_6S_Total <- rbind(CleanData_6S_Total, supplemental_6S_T) %>% select(-CoV)
Merge_6S_Total$Mass <- Merge_6S_Total$Mass %>% replace_na('...1')

#### Transpose & Merger ####

Transpose_6S_Total <- transpose(Merge_6S_Total)
Transpose_6S_Total <- Transpose_6S_Total %>% row_to_names(row_number = 1)
#Transpose_6S_Total <- mutate_all(Transpose_6S_Total, function(x) as.numeric(as.character(x)))

# Joining dataframe with another through a matching "key" variable.
Final_6S_Total <- Transpose_6S_Total %>% left_join(merger,by = "...1" )

# Combine 
Final_6S_Total$CRE.and.Treatment <- paste(Final_6S_Total$genotype,Final_6S_Total$Treatment,sep="/")
Final_6S_Total <- Final_6S_Total %>% filter(!grepl('86|87', Sample.number))
Final_6S_Total_CE <- Final_6S_Total %>% select(-"Sample.name",-"dry.extracted.tissue.wt",-"genotype",-"Treatment",-"Dry.lipid.wt")
Final_6S_Total_noCE <- Final_6S_Total %>% select(-"Total CholEsters",-"Sample.name",-"dry.extracted.tissue.wt",-"genotype",-"Treatment",-"Dry.lipid.wt")

write_csv(Final_6S_Total_CE, "Sox10_Final_6S_Total.csv")
write_csv(Final_6S_Total_noCE, "Sox10_Final_6S_Total_noCE.csv")
# read with rownames(df) <- df$...1


### Individual ####

# To add labels for merging
CleanData_6S_All <- SoxWeek6_18_Spinal %>% select(Mass, CoV, "Compound Name",contains("S_")) 
supplemental_6S_A <- CleanData_6S_All[2,]
#Filter out total values
CleanData_6S_All <- CleanData_6S_All %>% filter(grepl(':', CleanData_6S_All$`Compound Name`))
CleanData_6S_All <- drop_na(CleanData_6S_All,)

# Remove all rows with CoV <0.3
CleanData_6S_All$CoV <- as.numeric(CleanData_6S_All$CoV)
CleanData_6S_All <- CleanData_6S_All %>% filter(CoV < 0.3) 

# Bind supplemental labels for merging
Merge_6S_All <- rbind(CleanData_6S_All, supplemental_6S_A) 
Merge_6S_All$'Compound Name' <- Merge_6S_All$'Compound Name' %>% replace_na('...1')

# Drop CoV column
Merge_6S_All <- Merge_6S_All %>% select(-CoV,-Mass)

# Fix row index
rownames(Merge_6S_All) <- Merge_6S_All$'Compound Name'

#### Transpose & Merge ####
Transpose_6S_All <- transpose(Merge_6S_All)
# Fix column and row names
rownames(Transpose_6S_All) <- colnames(Merge_6S_All)
Transpose_6S_All <- Transpose_6S_All %>% row_to_names(row_number = 1)

# Merge dataframes for CRE status through a matching "key" variable.
#Transpose_6S_All <- mutate_all(Transpose_6S_All, function(x) as.numeric(as.character(x)))
Final_6S_All <- Transpose_6S_All %>% left_join(merger,by = "...1" )
Final_6S_All$CRE.and.Treatment <- paste(Final_6S_All$genotype,Final_6S_All$Treatment,sep="/")
Final_6S_All <- Final_6S_All %>% filter(!grepl('86|87', Sample.number))

Final_6S_All_CE <- Final_6S_All %>% select(-"Sample.name",-"dry.extracted.tissue.wt",-"genotype",-"Treatment",-"Dry.lipid.wt")
Final_6S_All_noCE <- Final_6S_All %>% select(-(161:171),-"Sample.name",-"dry.extracted.tissue.wt",-"genotype",-"Treatment",-"Dry.lipid.wt")

write_csv(Final_6S_All_CE, "Sox10_Final_6S_All.csv")
write_csv(Final_6S_All_noCE, "Sox10_Final_6S_All_noCE.csv")
# read with rownames(df) <- df$...1
```
