## Supplemental Codes for "Lipidomic analysis reveals differences in the extent of remyelination in the brain and spinal cord": Multi_timepoint_PCA.R

```
library(dplyr)
library(readxl)
library(tidyverse)
library(igraph)
library(ggplot2)    
library(data.table)
library(factoextra)
library("corrplot")

Brain_6to18 <- read_csv("Final_6B_Total_noCE.csv", name_repair = unique)
rownames(Brain_6to18) <- Brain_6to18$...1
Brain_6to18 <- Brain_6to18 %>% filter(grepl('Control', CRE.and.Treatment))

Spinal_6to18 <- read_csv("Final_6S_Total_noCE_w_LPE.csv", name_repair = unique)
rownames(Spinal_6to18) <- Spinal_6to18$...1
Spinal_6to18 <- Spinal_6to18 %>% filter(grepl('Control', CRE.and.Treatment))

##  Revised sample order plots 4/3/2023 #####

Grouping <- read_csv("new_6-18_grouping.csv")

# Merge grouping information with data
Brain_6to18 <-merge(x=Brain_6to18,y=Grouping,by='Sample.number',all.x=TRUE)
# Get rid of extra ...1 column
Brain_6to18 <-Brain_6to18[-15]

# Remove extra characters from ...1 column to use for categorizing/grouping
Brain_6to18$...1 <- str_sub(Brain_6to18$...1, 1, str_length(Brain_6to18$...1)-3)
Brain_6to18 <- Brain_6to18 %>% mutate(across('...1', str_replace, 'wk 6', 'wk 06'))
Brain_6to18$CRE.Treatment.Week <- paste(Brain_6to18$CRE.and.Treatment,Brain_6to18$...1,sep="/")

PCA_Brain <- Brain_6to18 %>% select(contains("Total"))
PCA_Brain <- mutate_all(PCA_Brain, function(x) as.numeric(as.character(x)))

PrincipleComponent_Brain <- prcomp(PCA_Brain, scale = TRUE)
PrincipleComponent_Brain$rotation <- -1*PrincipleComponent_Brain$rotation
summary(PrincipleComponent_Brain)
# 78.79% in 2 PC

names(PrincipleComponent_Brain)
PrincipleComponent_Brain$center
PrincipleComponent_Brain$scale

### Grouping by 3 groups - CRE neg, demyelinated, remyelinated ####
# Grouping PCA visual of CRE +/-
groups <- as.factor(Brain_6to18$Cre_group)
fviz_pca_ind(PrincipleComponent_Brain,
             col.ind = groups, palette = c("#00AFBB", "#FC4E07","#3ABF3F"), addEllipses = TRUE, ellipse.type = "confidence",
             legend.title = "Biological variables",
             repel = TRUE, title = "PCA for W6-18 brain totals - Grouping by biological differences")

### Grouping by 3 sample order groups ####
groups <- as.factor(Brain_6to18$Sample_group)
fviz_pca_ind(PrincipleComponent_Brain,
             col.ind = groups, palette = c("#7F58AF", "#E84D8A","#FEB326"), addEllipses = TRUE, ellipse.type = "confidence",
             legend.title = "Sample order",
             repel = TRUE, title = "PCA for W6-18 brain totals - Grouping by sample order")
# "#7F58AF", "#64C5EB", "#E84D8A", "#FEB326"

##  Spinal Grouping #####

Grouping <- read_csv("new_6-18_grouping_SC.csv")

# Merge grouping information with data
Spinal_6to18 <-merge(x=Spinal_6to18,y=Grouping,by='Sample.number',all.x=TRUE)
# Get rid of extra ...1 column
Spinal_6to18 <-Spinal_6to18[-15]

# Remove extra characters from ...1 column to use for categorizing/grouping
Spinal_6to18$...1 <- str_sub(Spinal_6to18$...1, 1, str_length(Spinal_6to18$...1)-3)
Spinal_6to18 <- Spinal_6to18 %>% mutate(across('...1', str_replace, 'wk 6', 'wk 06'))
Spinal_6to18$CRE.Treatment.Week <- paste(Spinal_6to18$CRE.and.Treatment,Spinal_6to18$...1,sep="/")

PCA_Spinal <- Spinal_6to18 %>% select(contains("Total"))
PCA_Spinal <- mutate_all(PCA_Spinal, function(x) as.numeric(as.character(x)))

PrincipleComponent_Spinal <- prcomp(PCA_Spinal, scale = TRUE)
PrincipleComponent_Spinal$rotation <- -1*PrincipleComponent_Spinal$rotation
summary(PrincipleComponent_Spinal)
# 83.51% in 2 PC

names(PrincipleComponent_Spinal)
PrincipleComponent_Spinal$center
PrincipleComponent_Spinal$scale

### Spinal -Grouping by 3 groups - CRE neg, demyelinated, remyelinated ####
# Grouping PCA visual of CRE +/-
groups <- as.factor(Spinal_6to18$Cre_group)
fviz_pca_ind(PrincipleComponent_Spinal,
             col.ind = groups, palette = c("#00AFBB", "#FC4E07","#3ABF3F"), addEllipses = TRUE, ellipse.type = "confidence",
             legend.title = "Biological variables",
             repel = TRUE, title = "PCA for W6-18 spinal cord totals - Grouping by \nbiological differences")

### Spinal - Grouping by 3 sample order groups ####
groups <- as.factor(Spinal_6to18$Sample_group)
fviz_pca_ind(PrincipleComponent_Spinal,
             col.ind = groups, palette = c("#7F58AF", "#E84D8A","#FEB326"), addEllipses = TRUE, ellipse.type = "confidence",
             legend.title = "Sample order",
             repel = TRUE, title = "PCA for W6-18 spinal cord totals - Grouping by sample order")
# "#7F58AF", "#64C5EB", "#E84D8A", "#FEB326"
```
