## Supplemental Codes for "Lipidomic analysis reveals differences in the extent of remyelination in the brain and spinal cord": PCA_Final.R

```
# To-Do ####

library(dplyr)
library(readxl)
library(tidyverse)
library(igraph)
library(ggplot2)    
library(data.table)
library(factoextra)
library("corrplot")

# NO CE in any data set

T6_18B <- read_csv("Final_6B_Total_noCE.csv", col_names = TRUE, name_repair = unique)
T6_18S <- read_csv("Final_6S_Total_noCE_w_LPE.csv", col_names = TRUE, name_repair = unique)

## T6 = W6 CRE+ totals vs W6 CRE- totals ####

T6B <- T6_18B %>% filter(grepl('wk 6', ...1)) %>% filter(grepl('Control', CRE.and.Treatment))
T6S <- T6_18S %>% filter(grepl('wk 6', ...1)) %>% filter(grepl('Control', CRE.and.Treatment))

### PCA for T6B ####
# Must drop not numerical values, aka : sample.number, ...1, cre.and.treatment
PCA_T6B <- T6B %>% select(contains("Total"))
PCA_T6B <- mutate_all(PCA_T6B, function(x) as.numeric(as.character(x)))

PrincipleComponent_T6B <- prcomp(PCA_T6B, scale. = TRUE)
PrincipleComponent_T6B$rotation <- -1*PrincipleComponent_T6B$rotation
summary(PrincipleComponent_T6B)
# 86.48% in 2 PC

## Biplot of PCA
pdf("Supplemental/Biplot_T6B_PLP.pdf", width = 6, height = 4)
fviz_pca_biplot(PrincipleComponent_T6B, repel = TRUE,
                col.var = "#2E9FDF",
                col.ind = "#696969", title = "PLP W6 brain totals")
dev.off()

## Grouping PCA visual of CRE +/-
pdf("Adjusted_PCA/Grouping_T6B_PLP.pdf", width = 1.75, height = 1.17)
groups <- as.factor(T6B$CRE.and.Treatment)
fviz_pca_ind(PrincipleComponent_T6B,
             col.ind = groups, palette = c("#00AFBB", "#FC4E07"), addEllipses = TRUE, ellipse.type = "confidence",
             legend.title = NULL, pointsize = 1, repel = TRUE, title = NULL, labelsize = 2, 
             ellipse.alpha = 0.3, ellipse.border.remove = TRUE, mean.point = FALSE
) + theme(legend.position = "none",
          text = element_text(size = 0),
          axis.text.x = element_text(face="plain", color="black", size=5, angle=0),
          axis.text.y = element_text(face="plain", color="black", size=5, angle=0),
          axis.ticks.length=unit(.05, "cm"), axis.ticks=element_line(linewidth=0.2),
          axis.title.x = element_text(family = "sans", size = 5, margin=margin(1,0,0,0)), 
          axis.title.y = element_text(family = "sans", size = 5, margin=margin(0,1,0,0)),
          panel.grid.major = element_line(linewidth = 0.5), 
          panel.grid.minor = element_line(linewidth = 0.5),
          plot.margin = margin(4,4,4,4)
)
dev.off()

### PCA for T6S ####
# Must drop not numerical values, aka : sample.number, ...1, cre.and.treatment
PCA_T6S <- T6S %>% select(contains("Total"))
PCA_T6S <- mutate_all(PCA_T6S, function(x) as.numeric(as.character(x)))

PrincipleComponent_T6S <- prcomp(PCA_T6S, scale. = TRUE)
PrincipleComponent_T6S$rotation <- -1*PrincipleComponent_T6S$rotation
summary(PrincipleComponent_T6S)
# 91.64% in 2 PC (without LPE)
# 83.42% w LPE

## Biplot of PCA
pdf("Supplemental/Biplot_T6S_PLP.pdf", width = 6, height = 4)
fviz_pca_biplot(PrincipleComponent_T6S, repel = TRUE,
                col.var = "#2E9FDF",
                col.ind = "#696969", title = "PLP W6 spinal cord totals")
dev.off()

## Grouping PCA visual of CRE +/-
pdf("Adjusted_PCA/Grouping_T6S_PLP.pdf", width = 1.75, height = 1.17)
groups <- as.factor(T6S$CRE.and.Treatment)
fviz_pca_ind(PrincipleComponent_T6S,
             col.ind = groups, palette = c("#00AFBB", "#FC4E07"), addEllipses = TRUE, ellipse.type = "confidence",
             legend.title = NULL, pointsize = 1, repel = TRUE, title = NULL, labelsize = 2, 
             ellipse.alpha = 0.3, ellipse.border.remove = TRUE, mean.point = FALSE
) + theme(legend.position = "none",
          text = element_text(size = 0),
          axis.text.x = element_text(face="plain", color="black", size=5, angle=0),
          axis.text.y = element_text(face="plain", color="black", size=5, angle=0),
          axis.ticks.length=unit(.05, "cm"), axis.ticks=element_line(linewidth=0.2),
          axis.title.x = element_text(family = "sans", size = 5, margin=margin(1,0,0,0)), 
          axis.title.y = element_text(family = "sans", size = 5, margin=margin(0,1,0,0)),
          panel.grid.major = element_line(linewidth = 0.5), 
          panel.grid.minor = element_line(linewidth = 0.5),
          plot.margin = margin(4,4,4,4)
)
dev.off()

## T12 = W12 CRE+ totals vs W12 CRE- totals ####

T12B <- T6_18B %>% filter(grepl('wk 12', ...1)) %>% filter(grepl('Control', CRE.and.Treatment))
T12S <- T6_18S %>% filter(grepl('wk 12', ...1)) %>% filter(grepl('Control', CRE.and.Treatment))

### PCA for T12B ####

# Must drop not numerical values, aka : sample.number, ...1, cre.and.treatment
PCA_T12B <- T12B %>% select(contains("Total"))
PCA_T12B <- mutate_all(PCA_T12B, function(x) as.numeric(as.character(x)))

PrincipleComponent_T12B <- prcomp(PCA_T12B, scale. = TRUE)
PrincipleComponent_T12B$rotation <- -1*PrincipleComponent_T12B$rotation
summary(PrincipleComponent_T12B)

# 82.53% in 2 PC

## Biplot of PCA
pdf("Supplemental/Biplot_T12B_PLP.pdf", width = 6, height = 4)
fviz_pca_biplot(PrincipleComponent_T12B, repel = TRUE,
                col.var = "#2E9FDF",
                col.ind = "#696969", title = "PLP W12 brain totals")
dev.off()

## Grouping PCA visual of CRE +/-
pdf("Adjusted_PCA/Grouping_T12B_PLP.pdf", width = 1.75, height = 1.17)
groups <- as.factor(T12B$CRE.and.Treatment)
fviz_pca_ind(PrincipleComponent_T12B,
             col.ind = groups, palette = c("#00AFBB", "#FC4E07"), addEllipses = TRUE, ellipse.type = "confidence",
             legend.title = NULL, pointsize = 1, repel = TRUE, title = NULL, labelsize = 2, 
             ellipse.alpha = 0.3, ellipse.border.remove = TRUE, mean.point = FALSE
) + theme(legend.position = "none",
          text = element_text(size = 0),
          axis.text.x = element_text(face="plain", color="black", size=5, angle=0),
          axis.text.y = element_text(face="plain", color="black", size=5, angle=0),
          axis.ticks.length=unit(.05, "cm"), axis.ticks=element_line(linewidth=0.2),
          axis.title.x = element_text(family = "sans", size = 5, margin=margin(1,0,0,0)), 
          axis.title.y = element_text(family = "sans", size = 5, margin=margin(0,1,0,0)),
          panel.grid.major = element_line(linewidth = 0.5), 
          panel.grid.minor = element_line(linewidth = 0.5),
          plot.margin = margin(4,4,4,4)
)
dev.off()

### PCA for T12S ####

# Must drop not numerical values, aka : sample.number, ...1, cre.and.treatment
PCA_T12S <- T12S %>% select(contains("Total"))
PCA_T12S <- mutate_all(PCA_T12S, function(x) as.numeric(as.character(x)))

PrincipleComponent_T12S <- prcomp(PCA_T12S, scale. = TRUE)
PrincipleComponent_T12S$rotation <- -1*PrincipleComponent_T12S$rotation
summary(PrincipleComponent_T12S)
# 94.63% in 2 PC (without LPE)
# 91.29% w LPE

## Biplot of PCA
pdf("Supplemental/Biplot_T12S_PLP.pdf", width = 6, height = 4)
fviz_pca_biplot(PrincipleComponent_T12S, repel = TRUE,
                col.var = "#2E9FDF",
                col.ind = "#696969", title = "PLP W12 spinal cord totals")
dev.off()

## Grouping PCA visual of CRE +/-
pdf("Adjusted_PCA/Grouping_T12S_PLP.pdf", width = 1.75, height = 1.17)
groups <- as.factor(T12S$CRE.and.Treatment)
fviz_pca_ind(PrincipleComponent_T12S,
             col.ind = groups, palette = c("#00AFBB", "#FC4E07"), addEllipses = TRUE, ellipse.type = "confidence",
             legend.title = NULL, pointsize = 1, repel = TRUE, title = NULL, labelsize = 2, 
             ellipse.alpha = 0.3, ellipse.border.remove = TRUE, mean.point = FALSE
) + theme(legend.position = "none",
          text = element_text(size = 0),
          axis.text.x = element_text(face="plain", color="black", size=5, angle=0),
          axis.text.y = element_text(face="plain", color="black", size=5, angle=0),
          axis.ticks.length=unit(.05, "cm"), axis.ticks=element_line(linewidth=0.2),
          axis.title.x = element_text(family = "sans", size = 5, margin=margin(1,0,0,0)), 
          axis.title.y = element_text(family = "sans", size = 5, margin=margin(0,1,0,0)),
          panel.grid.major = element_line(linewidth = 0.5), 
          panel.grid.minor = element_line(linewidth = 0.5),
          plot.margin = margin(4,4,4,4)
)
dev.off()

## T18 = W18 CRE+ totals vs W18 CRE- totals ####

T18B <- T6_18B %>% filter(grepl('wk 18', ...1)) %>% filter(grepl('Control', CRE.and.Treatment))
T18S <- T6_18S %>% filter(grepl('wk 18', ...1)) %>% filter(grepl('Control', CRE.and.Treatment))

### PCA for T18B ####

# Must drop not numerical values, aka : sample.number, ...1, cre.and.treatment
PCA_T18B <- T18B %>% select(contains("Total"))
PCA_T18B <- mutate_all(PCA_T18B, function(x) as.numeric(as.character(x)))

PrincipleComponent_T18B <- prcomp(PCA_T18B, scale. = TRUE)
PrincipleComponent_T18B$rotation <- -1*PrincipleComponent_T18B$rotation
summary(PrincipleComponent_T18B)

# 86.11% in 2 PC

## Biplot of PCA
pdf("Supplemental/Biplot_T18B_PLP.pdf", width = 6, height = 4)
fviz_pca_biplot(PrincipleComponent_T18B, repel = TRUE,
                col.var = "#2E9FDF",
                col.ind = "#696969", title = "PLP W18 brain totals")
dev.off()

## Grouping PCA visual of CRE +/-
pdf("Adjusted_PCA/Grouping_T18B_PLP.pdf", width = 1.75, height = 1.17)
groups <- as.factor(T18B$CRE.and.Treatment)
fviz_pca_ind(PrincipleComponent_T18B,
             col.ind = groups, palette = c("#00AFBB", "#FC4E07"), addEllipses = TRUE, ellipse.type = "confidence",
             legend.title = NULL, pointsize = 1, repel = TRUE, title = NULL, labelsize = 2, 
             ellipse.alpha = 0.3, ellipse.border.remove = TRUE, mean.point = FALSE
) + theme(legend.position = "none",
          text = element_text(size = 0),
          axis.text.x = element_text(face="plain", color="black", size=5, angle=0),
          axis.text.y = element_text(face="plain", color="black", size=5, angle=0),
          axis.ticks.length=unit(.05, "cm"), axis.ticks=element_line(linewidth=0.2),
          axis.title.x = element_text(family = "sans", size = 5, margin=margin(1,0,0,0)), 
          axis.title.y = element_text(family = "sans", size = 5, margin=margin(0,1,0,0)),
          panel.grid.major = element_line(linewidth = 0.5), 
          panel.grid.minor = element_line(linewidth = 0.5),
          plot.margin = margin(4,4,4,4)
)
dev.off()

### PCA for T18S ####

# Must drop not numerical values, aka : sample.number, ...1, cre.and.treatment
PCA_T18S <- T18S %>% select(contains("Total"))
PCA_T18S <- mutate_all(PCA_T18S, function(x) as.numeric(as.character(x)))

PrincipleComponent_T18S <- prcomp(PCA_T18S, scale. = TRUE)
PrincipleComponent_T18S$rotation <- -1*PrincipleComponent_T18S$rotation
summary(PrincipleComponent_T18S)
# 90.43% in 2 PC (without LPE)
# 90.40% w LPE

## Biplot of PCA
pdf("Supplemental/Biplot_T18S_PLP.pdf", width = 6, height = 4)
fviz_pca_biplot(PrincipleComponent_T18S, repel = TRUE,
                col.var = "#2E9FDF",
                col.ind = "#696969", title = "PLP W18 spinal cord totals")
dev.off()

## Grouping PCA visual of CRE +/-
pdf("Adjusted_PCA/Grouping_T18S_PLP.pdf", width = 1.75, height = 1.17)
groups <- as.factor(T18S$CRE.and.Treatment)
fviz_pca_ind(PrincipleComponent_T18S,
             col.ind = groups, palette = c("#00AFBB", "#FC4E07"), addEllipses = TRUE, ellipse.type = "confidence",
             legend.title = NULL, pointsize = 1, repel = TRUE, title = NULL, labelsize = 2, 
             ellipse.alpha = 0.3, ellipse.border.remove = TRUE, mean.point = FALSE
) + theme(legend.position = "none",
          text = element_text(size = 0),
          axis.text.x = element_text(face="plain", color="black", size=5, angle=0),
          axis.text.y = element_text(face="plain", color="black", size=5, angle=0),
          axis.ticks.length=unit(.05, "cm"), axis.ticks=element_line(linewidth=0.2),
          axis.title.x = element_text(family = "sans", size = 5, margin=margin(1,0,0,0)), 
          axis.title.y = element_text(family = "sans", size = 5, margin=margin(0,1,0,0)),
          panel.grid.major = element_line(linewidth = 0.5), 
          panel.grid.minor = element_line(linewidth = 0.5),
          plot.margin = margin(4,4,4,4)
)
dev.off()

## T24 = W24 CRE+ totals vs W18 CRE- totals ####

T24B <- read_csv("Final_24B_Total_noCE.csv", col_names = TRUE, name_repair = unique)
T24B <- T24B %>% filter(grepl('wk 24', ...1)) %>% filter(grepl('Control', CRE.and.Treatment))

T24S <- read_csv("Final_24S_Total_noCE_w_LPE.csv", col_names = TRUE, name_repair = unique)
T24S <- T24S %>% filter(grepl('wk 24', ...1)) %>% filter(grepl('Control', CRE.and.Treatment))

### PCA for T24B ####
# Must drop not numerical values, aka : sample.number, ...1, cre.and.treatment
PCA_T24B <- T24B %>% select(contains("Total"))
PCA_T24B <- mutate_all(PCA_T24B, function(x) as.numeric(as.character(x)))

PrincipleComponent_T24B <- prcomp(PCA_T24B, scale. = TRUE)
PrincipleComponent_T24B$rotation <- -1*PrincipleComponent_T24B$rotation
summary(PrincipleComponent_T24B)

# 67.82% in 2 PC

## Biplot of PCA
pdf("Supplemental/Biplot_T24B_PLP.pdf", width = 6, height = 4)
fviz_pca_biplot(PrincipleComponent_T24B, repel = TRUE,
                col.var = "#2E9FDF",
                col.ind = "#696969", title = "PLP W24 brain totals")
dev.off()

## Grouping PCA visual of CRE +/-
pdf("Adjusted_PCA/Grouping_T24B_PLP.pdf", width = 1.75, height = 1.17)
groups <- as.factor(T24B$CRE.and.Treatment)
fviz_pca_ind(PrincipleComponent_T24B,
             col.ind = groups, palette = c("#00AFBB", "#FC4E07"), addEllipses = TRUE, ellipse.type = "confidence",
             legend.title = NULL, pointsize = 1, repel = TRUE, title = NULL, labelsize = 2, 
             ellipse.alpha = 0.3, ellipse.border.remove = TRUE, mean.point = FALSE
) + theme(legend.position = "none",
          text = element_text(size = 0),
          axis.text.x = element_text(face="plain", color="black", size=5, angle=0),
          axis.text.y = element_text(face="plain", color="black", size=5, angle=0),
          axis.ticks.length=unit(.05, "cm"), axis.ticks=element_line(linewidth=0.2),
          axis.title.x = element_text(family = "sans", size = 5, margin=margin(1,0,0,0)), 
          axis.title.y = element_text(family = "sans", size = 5, margin=margin(0,1,0,0)),
          panel.grid.major = element_line(linewidth = 0.5), 
          panel.grid.minor = element_line(linewidth = 0.5),
          plot.margin = margin(4,4,4,4)
)
dev.off()


### PCA for T24S ####
# Must drop not numerical values, aka : sample.number, ...1, cre.and.treatment
PCA_T24S <- T24S %>% select(contains("Total"))
PCA_T24S <- mutate_all(PCA_T24S, function(x) as.numeric(as.character(x)))

PrincipleComponent_T24S <- prcomp(PCA_T24S, scale. = TRUE)
PrincipleComponent_T24S$rotation <- -1*PrincipleComponent_T24S$rotation
summary(PrincipleComponent_T24S)
# 90.67% in 2 PC (without LPE)
# 91.16% w LPE

## Biplot of PCA
pdf("Supplemental/Biplot_T24S_PLP.pdf", width = 6, height = 4)
fviz_pca_biplot(PrincipleComponent_T24S, repel = TRUE,
                col.var = "#2E9FDF",
                col.ind = "#696969", title = "PLP W24 spinal cord totals")
dev.off()

## Grouping PCA visual of CRE +/-
pdf("Adjusted_PCA/Grouping_T24S_PLP.pdf", width = 1.75, height = 1.17)
groups <- as.factor(T24S$CRE.and.Treatment)
fviz_pca_ind(PrincipleComponent_T24S,
             col.ind = groups, palette = c("#00AFBB", "#FC4E07"), addEllipses = TRUE, ellipse.type = "confidence",
             legend.title = NULL, pointsize = 1, repel = TRUE, title = NULL, labelsize = 2, 
             ellipse.alpha = 0.3, ellipse.border.remove = TRUE, mean.point = FALSE
) + theme(legend.position = "none",
          text = element_text(size = 0),
          axis.text.x = element_text(face="plain", color="black", size=5, angle=0),
          axis.text.y = element_text(face="plain", color="black", size=5, angle=0),
          axis.ticks.length=unit(.05, "cm"), axis.ticks=element_line(linewidth=0.2),
          axis.title.x = element_text(family = "sans", size = 5, margin=margin(1,0,0,0)), 
          axis.title.y = element_text(family = "sans", size = 5, margin=margin(0,1,0,0)),
          panel.grid.major = element_line(linewidth = 0.5), 
          panel.grid.minor = element_line(linewidth = 0.5),
          plot.margin = margin(4,4,4,4)
)
dev.off()

# SobAM2 Brain ####

## T12 w SobAM2 = W12 CRE+ totals vs W12 CRE+ w SobAM2 totals ####

T6_18B <- read_csv("Final_6B_Total_noCE.csv", col_names = TRUE, name_repair = unique)
T12B_SobAM2_CrePos <- T6_18B %>% filter(grepl('wk 12', ...1)) %>% filter(!grepl('-', CRE.and.Treatment))

### PCA for T12B_SobAM2_CrePos ####
# Must drop not numerical values, aka : sample.number, ...1, cre.and.treatment
PCA_T12B_SobAM2_CrePos <- T12B_SobAM2_CrePos %>% select(contains("Total"))
PCA_T12B_SobAM2_CrePos <- mutate_all(PCA_T12B_SobAM2_CrePos, function(x) as.numeric(as.character(x)))

PrincipleComponent_T12B_SobAM2_CrePos <- prcomp(PCA_T12B_SobAM2_CrePos, scale. = TRUE)
PrincipleComponent_T12B_SobAM2_CrePos$rotation <- -1*PrincipleComponent_T12B_SobAM2_CrePos$rotation
summary(PrincipleComponent_T12B_SobAM2_CrePos)

# 79.04% in 2 PC

## Biplot of PCA
pdf("Supplemental/Biplot_T12B_SobAM2_CrePos_PLP.pdf", width = 6, height = 4)
fviz_pca_biplot(PrincipleComponent_T12B_SobAM2_CrePos, repel = TRUE,
                col.var = "#2E9FDF",
                col.ind = "#696969", title = "PLP W12 CRE+ SobAM2 brain totals")
dev.off()

## Grouping PCA visual of CRE +/-
pdf("Adjusted_PCA/Grouping_T12B_SobAM2_CrePos_PLP.pdf", width = 1.75, height = 1.17)
groups <- as.factor(T12B_SobAM2_CrePos$CRE.and.Treatment)
fviz_pca_ind(PrincipleComponent_T12B_SobAM2_CrePos,
             col.ind = groups, palette = c("#00AFBB", "#FC4E07"), addEllipses = TRUE, ellipse.type = "confidence",
             legend.title = NULL, pointsize = 1, repel = TRUE, title = NULL, labelsize = 2, 
             ellipse.alpha = 0.3, ellipse.border.remove = TRUE, mean.point = FALSE
) + theme(legend.position = "none",
          text = element_text(size = 0),
          axis.text.x = element_text(face="plain", color="black", size=5, angle=0),
          axis.text.y = element_text(face="plain", color="black", size=5, angle=0),
          axis.ticks.length=unit(.05, "cm"), axis.ticks=element_line(linewidth=0.2),
          axis.title.x = element_text(family = "sans", size = 5, margin=margin(1,0,0,0)), 
          axis.title.y = element_text(family = "sans", size = 5, margin=margin(0,1,0,0)),
          panel.grid.major = element_line(linewidth = 0.5), 
          panel.grid.minor = element_line(linewidth = 0.5),
          plot.margin = margin(4,4,4,4)
)
dev.off()


## T12 w SobAM2 = W12 CRE- totals vs W12 CRE- w SobAM2 totals ####

T12B_SobAM2_CreNeg <- T6_18B %>% filter(grepl('wk 12', ...1)) %>% filter(grepl('-', CRE.and.Treatment))

### PCA for T12B_SobAM2_CreNeg ####
# Must drop not numerical values, aka : sample.number, ...1, cre.and.treatment
PCA_T12B_SobAM2_CreNeg <- T12B_SobAM2_CreNeg %>% select(contains("Total"))
PCA_T12B_SobAM2_CreNeg <- mutate_all(PCA_T12B_SobAM2_CreNeg, function(x) as.numeric(as.character(x)))

PrincipleComponent_T12B_SobAM2_CreNeg <- prcomp(PCA_T12B_SobAM2_CreNeg, scale. = TRUE)
PrincipleComponent_T12B_SobAM2_CreNeg$rotation <- -1*PrincipleComponent_T12B_SobAM2_CreNeg$rotation
summary(PrincipleComponent_T12B_SobAM2_CreNeg)

# 73.62% in 2 PC

## Biplot of PCA
pdf("Supplemental/Biplot_T12B_SobAM2_CreNeg_PLP.pdf", width = 6, height = 4)
fviz_pca_biplot(PrincipleComponent_T12B_SobAM2_CreNeg, repel = TRUE,
                col.var = "#2E9FDF",
                col.ind = "#696969", title = "PLP W12 CRE- SobAM2 brain totals")
dev.off()

## Grouping PCA visual of CRE +/-
pdf("Adjusted_PCA/Grouping_T12B_SobAM2_CreNeg_PLP.pdf", width = 1.75, height = 1.17)
groups <- as.factor(T12B_SobAM2_CreNeg$CRE.and.Treatment)
fviz_pca_ind(PrincipleComponent_T12B_SobAM2_CreNeg,
             col.ind = groups, palette = c("#00AFBB", "#FC4E07"), addEllipses = TRUE, ellipse.type = "confidence",
             legend.title = NULL, pointsize = 1, repel = TRUE, title = NULL, labelsize = 2, 
             ellipse.alpha = 0.3, ellipse.border.remove = TRUE, mean.point = FALSE
) + theme(legend.position = "none",
          text = element_text(size = 0),
          axis.text.x = element_text(face="plain", color="black", size=5, angle=0),
          axis.text.y = element_text(face="plain", color="black", size=5, angle=0),
          axis.ticks.length=unit(.05, "cm"), axis.ticks=element_line(linewidth=0.2),
          axis.title.x = element_text(family = "sans", size = 5, margin=margin(1,0,0,0)), 
          axis.title.y = element_text(family = "sans", size = 5, margin=margin(0,1,0,0)),
          panel.grid.major = element_line(linewidth = 0.5), 
          panel.grid.minor = element_line(linewidth = 0.5),
          plot.margin = margin(4,4,4,4)
)
dev.off()

## T18 w SobAM2 = W18 CRE+ totals vs W18 CRE+ w SobAM2 totals ####

T18B_SobAM2_CrePos <- T6_18B %>% filter(grepl('wk 18', ...1)) %>% filter(!grepl('-', CRE.and.Treatment))

### PCA for T18B_SobAM2_CrePos ####
# Must drop not numerical values, aka : sample.number, ...1, cre.and.treatment
PCA_T18B_SobAM2_CrePos <- T18B_SobAM2_CrePos %>% select(contains("Total"))
PCA_T18B_SobAM2_CrePos <- mutate_all(PCA_T18B_SobAM2_CrePos, function(x) as.numeric(as.character(x)))

PrincipleComponent_T18B_SobAM2_CrePos <- prcomp(PCA_T18B_SobAM2_CrePos, scale. = TRUE)
PrincipleComponent_T18B_SobAM2_CrePos$rotation <- -1*PrincipleComponent_T18B_SobAM2_CrePos$rotation
summary(PrincipleComponent_T18B_SobAM2_CrePos)

# 76.16% in 2 PC

## Biplot of PCA
pdf("Supplemental/Biplot_T18B_SobAM2_CrePos_PLP.pdf", width = 6, height = 4)
fviz_pca_biplot(PrincipleComponent_T18B_SobAM2_CrePos, repel = TRUE,
                col.var = "#2E9FDF",
                col.ind = "#696969", title = "PLP W18 CRE+ SobAM2 brain totals")
dev.off()

## Grouping PCA visual of CRE +/-
pdf("Adjusted_PCA/Grouping_T18B_SobAM2_CrePos_PLP.pdf", width = 1.75, height = 1.17)
groups <- as.factor(T18B_SobAM2_CrePos$CRE.and.Treatment)
fviz_pca_ind(PrincipleComponent_T18B_SobAM2_CrePos,
             col.ind = groups, palette = c("#00AFBB", "#FC4E07"), addEllipses = TRUE, ellipse.type = "confidence",
             legend.title = NULL, pointsize = 1, repel = TRUE, title = NULL, labelsize = 2, 
             ellipse.alpha = 0.3, ellipse.border.remove = TRUE, mean.point = FALSE
) + theme(legend.position = "none",
          text = element_text(size = 0),
          axis.text.x = element_text(face="plain", color="black", size=5, angle=0),
          axis.text.y = element_text(face="plain", color="black", size=5, angle=0),
          axis.ticks.length=unit(.05, "cm"), axis.ticks=element_line(linewidth=0.2),
          axis.title.x = element_text(family = "sans", size = 5, margin=margin(1,0,0,0)), 
          axis.title.y = element_text(family = "sans", size = 5, margin=margin(0,1,0,0)),
          panel.grid.major = element_line(linewidth = 0.5), 
          panel.grid.minor = element_line(linewidth = 0.5),
          plot.margin = margin(4,4,4,4)
)
dev.off()

## T18 w SobAM2 = W18 CRE- totals vs W18 CRE- w SobAM2 totals ####

T18B_SobAM2_CreNeg <- T6_18B %>% filter(grepl('wk 18', ...1)) %>% filter(grepl('-', CRE.and.Treatment))

### PCA for T18B_SobAM2_CreNeg ####
# Must drop not numerical values, aka : sample.number, ...1, cre.and.treatment
PCA_T18B_SobAM2_CreNeg <- T18B_SobAM2_CreNeg %>% select(contains("Total"))
PCA_T18B_SobAM2_CreNeg <- mutate_all(PCA_T18B_SobAM2_CreNeg, function(x) as.numeric(as.character(x)))

PrincipleComponent_T18B_SobAM2_CreNeg <- prcomp(PCA_T18B_SobAM2_CreNeg, scale. = TRUE)
PrincipleComponent_T18B_SobAM2_CreNeg$rotation <- -1*PrincipleComponent_T18B_SobAM2_CreNeg$rotation
summary(PrincipleComponent_T18B_SobAM2_CreNeg)

#70.85% in 2 PC

## Biplot of PCA
pdf("Supplemental/Biplot_T18B_SobAM2_CreNeg_PLP.pdf", width = 6, height = 4)
fviz_pca_biplot(PrincipleComponent_T18B_SobAM2_CreNeg, repel = TRUE,
                col.var = "#2E9FDF",
                col.ind = "#696969", title = "PLP W18 CRE- SobAM2 brain totals")
dev.off()

## Grouping PCA visual of CRE +/-
pdf("Adjusted_PCA/Grouping_T18B_SobAM2_CreNeg_PLP.pdf", width = 1.75, height = 1.17)
groups <- as.factor(T18B_SobAM2_CreNeg$CRE.and.Treatment)
fviz_pca_ind(PrincipleComponent_T18B_SobAM2_CreNeg,
             col.ind = groups, palette = c("#00AFBB", "#FC4E07"), addEllipses = TRUE, ellipse.type = "confidence",
             legend.title = NULL, pointsize = 1, repel = TRUE, title = NULL, labelsize = 2, 
             ellipse.alpha = 0.3, ellipse.border.remove = TRUE, mean.point = FALSE
) + theme(legend.position = "none",
          text = element_text(size = 0),
          axis.text.x = element_text(face="plain", color="black", size=5, angle=0),
          axis.text.y = element_text(face="plain", color="black", size=5, angle=0),
          axis.ticks.length=unit(.05, "cm"), axis.ticks=element_line(linewidth=0.2),
          axis.title.x = element_text(family = "sans", size = 5, margin=margin(1,0,0,0)), 
          axis.title.y = element_text(family = "sans", size = 5, margin=margin(0,1,0,0)),
          panel.grid.major = element_line(linewidth = 0.5), 
          panel.grid.minor = element_line(linewidth = 0.5),
          plot.margin = margin(4,4,4,4)
)
dev.off()

# SobAM2 Spinal ####

T6_18S <- read_csv("Final_6S_Total_noCE_w_LPE.csv", col_names = TRUE, name_repair = unique)

## T12 w SobAM2 = W12 CRE+ totals vs W12 CRE+ w SobAM2 totals ####

T12S_SobAM2_CrePos <- T6_18S %>% filter(grepl('wk 12', ...1)) %>% filter(!grepl('-', CRE.and.Treatment))

### PCA for T12S_SobAM2_CrePos ####
# Must drop not numerical values, aka : sample.number, ...1, cre.and.treatment
PCA_T12S_SobAM2_CrePos <- T12S_SobAM2_CrePos %>% select(contains("Total"))
PCA_T12S_SobAM2_CrePos <- mutate_all(PCA_T12S_SobAM2_CrePos, function(x) as.numeric(as.character(x)))

PrincipleComponent_T12S_SobAM2_CrePos <- prcomp(PCA_T12S_SobAM2_CrePos, scale. = TRUE)
PrincipleComponent_T12S_SobAM2_CrePos$rotation <- -1*PrincipleComponent_T12S_SobAM2_CrePos$rotation
summary(PrincipleComponent_T12S_SobAM2_CrePos)

# 76.28% in 2 PC

## Biplot of PCA
pdf("Supplemental/Biplot_T12S_SobAM2_CrePos_PLP.pdf", width = 6, height = 4)
fviz_pca_biplot(PrincipleComponent_T12S_SobAM2_CrePos, repel = TRUE,
                col.var = "#2E9FDF",
                col.ind = "#696969", title = "PLP W12 CRE+ SobAM2 spinal cord totals")
dev.off()

## Grouping PCA visual of CRE +/-
pdf("Adjusted_PCA/Grouping_T12S_SobAM2_CrePos_PLP.pdf", width = 1.75, height = 1.17)
groups <- as.factor(T12S_SobAM2_CrePos$CRE.and.Treatment)
fviz_pca_ind(PrincipleComponent_T12S_SobAM2_CrePos,
             col.ind = groups, palette = c("#00AFBB", "#FC4E07"), addEllipses = TRUE, ellipse.type = "confidence",
             legend.title = NULL, pointsize = 1, repel = TRUE, title = NULL, labelsize = 2, 
             ellipse.alpha = 0.3, ellipse.border.remove = TRUE, mean.point = FALSE
) + theme(legend.position = "none",
          text = element_text(size = 0),
          axis.text.x = element_text(face="plain", color="black", size=5, angle=0),
          axis.text.y = element_text(face="plain", color="black", size=5, angle=0),
          axis.ticks.length=unit(.05, "cm"), axis.ticks=element_line(linewidth=0.2),
          axis.title.x = element_text(family = "sans", size = 5, margin=margin(1,0,0,0)), 
          axis.title.y = element_text(family = "sans", size = 5, margin=margin(0,1,0,0)),
          panel.grid.major = element_line(linewidth = 0.5), 
          panel.grid.minor = element_line(linewidth = 0.5),
          plot.margin = margin(4,4,4,4)
)
dev.off()


## T12 w SobAM2 = W12 CRE- totals vs W12 CRE- w SobAM2 totals ####

T12S_SobAM2_CreNeg <- T6_18S %>% filter(grepl('wk 12', ...1)) %>% filter(grepl('-', CRE.and.Treatment))

### PCA for T12S_SobAM2_CreNeg ####
# Must drop not numerical values, aka : sample.number, ...1, cre.and.treatment
PCA_T12S_SobAM2_CreNeg <- T12S_SobAM2_CreNeg %>% select(contains("Total"))
PCA_T12S_SobAM2_CreNeg <- mutate_all(PCA_T12S_SobAM2_CreNeg, function(x) as.numeric(as.character(x)))

PrincipleComponent_T12S_SobAM2_CreNeg <- prcomp(PCA_T12S_SobAM2_CreNeg, scale. = TRUE)
PrincipleComponent_T12S_SobAM2_CreNeg$rotation <- -1*PrincipleComponent_T12S_SobAM2_CreNeg$rotation
summary(PrincipleComponent_T12S_SobAM2_CreNeg)

# 78.26% in 2 PC

## Biplot of PCA
pdf("Supplemental/Biplot_T12S_SobAM2_CreNeg_PLP.pdf", width = 6, height = 4)
fviz_pca_biplot(PrincipleComponent_T12S_SobAM2_CreNeg, repel = TRUE,
                col.var = "#2E9FDF",
                col.ind = "#696969", title = "PLP W12 CRE- SobAM2 spinal cord totals")
dev.off()

## Grouping PCA visual of CRE +/-
pdf("Adjusted_PCA/Grouping_T12S_SobAM2_CreNeg_PLP.pdf", width = 1.75, height = 1.17)
groups <- as.factor(T12S_SobAM2_CreNeg$CRE.and.Treatment)
fviz_pca_ind(PrincipleComponent_T12S_SobAM2_CreNeg,
             col.ind = groups, palette = c("#00AFBB", "#FC4E07"), addEllipses = TRUE, ellipse.type = "confidence",
             legend.title = NULL, pointsize = 1, repel = TRUE, title = NULL, labelsize = 2, 
             ellipse.alpha = 0.3, ellipse.border.remove = TRUE, mean.point = FALSE
) + theme(legend.position = "none",
          text = element_text(size = 0),
          axis.text.x = element_text(face="plain", color="black", size=5, angle=0),
          axis.text.y = element_text(face="plain", color="black", size=5, angle=0),
          axis.ticks.length=unit(.05, "cm"), axis.ticks=element_line(linewidth=0.2),
          axis.title.x = element_text(family = "sans", size = 5, margin=margin(1,0,0,0)), 
          axis.title.y = element_text(family = "sans", size = 5, margin=margin(0,1,0,0)),
          panel.grid.major = element_line(linewidth = 0.5), 
          panel.grid.minor = element_line(linewidth = 0.5),
          plot.margin = margin(4,4,4,4)
)
dev.off()

## T18 w SobAM2 = W18 CRE+ totals vs W18 CRE+ w SobAM2 totals ####

T18S_SobAM2_CrePos <- T6_18S %>% filter(grepl('wk 18', ...1)) %>% filter(!grepl('-', CRE.and.Treatment))

### PCA for T18S_SobAM2_CrePos ####
# Must drop not numerical values, aka : sample.number, ...1, cre.and.treatment
PCA_T18S_SobAM2_CrePos <- T18S_SobAM2_CrePos %>% select(contains("Total"))
PCA_T18S_SobAM2_CrePos <- mutate_all(PCA_T18S_SobAM2_CrePos, function(x) as.numeric(as.character(x)))

PrincipleComponent_T18S_SobAM2_CrePos <- prcomp(PCA_T18S_SobAM2_CrePos, scale. = TRUE)
PrincipleComponent_T18S_SobAM2_CrePos$rotation <- -1*PrincipleComponent_T18S_SobAM2_CrePos$rotation
summary(PrincipleComponent_T18S_SobAM2_CrePos)

# 62.13% in 2 PC

## Biplot of PCA
pdf("Supplemental/Biplot_T18S_SobAM2_CrePos_PLP.pdf", width = 6, height = 4)
fviz_pca_biplot(PrincipleComponent_T18S_SobAM2_CrePos, repel = TRUE,
                col.var = "#2E9FDF",
                col.ind = "#696969", title = "PLP W18 CRE+ SobAM2 spinal cord totals")
dev.off()

## Grouping PCA visual of CRE +/-
pdf("Adjusted_PCA/Grouping_T18S_SobAM2_CrePos_PLP.pdf", width = 1.75, height = 1.17)
groups <- as.factor(T18S_SobAM2_CrePos$CRE.and.Treatment)
fviz_pca_ind(PrincipleComponent_T18S_SobAM2_CrePos,
             col.ind = groups, palette = c("#00AFBB", "#FC4E07"), addEllipses = TRUE, ellipse.type = "confidence",
             legend.title = NULL, pointsize = 1, repel = TRUE, title = NULL, labelsize = 2, 
             ellipse.alpha = 0.3, ellipse.border.remove = TRUE, mean.point = FALSE
) + theme(legend.position = "none",
          text = element_text(size = 0),
          axis.text.x = element_text(face="plain", color="black", size=5, angle=0),
          axis.text.y = element_text(face="plain", color="black", size=5, angle=0),
          axis.ticks.length=unit(.05, "cm"), axis.ticks=element_line(linewidth=0.2),
          axis.title.x = element_text(family = "sans", size = 5, margin=margin(1,0,0,0)), 
          axis.title.y = element_text(family = "sans", size = 5, margin=margin(0,1,0,0)),
          panel.grid.major = element_line(linewidth = 0.5), 
          panel.grid.minor = element_line(linewidth = 0.5),
          plot.margin = margin(4,4,4,4)
)
dev.off()

## T18 w SobAM2 = W18 CRE- totals vs W18 CRE- w SobAM2 totals ####

T18S_SobAM2_CreNeg <- T6_18S %>% filter(grepl('wk 18', ...1)) %>% filter(grepl('-', CRE.and.Treatment))

### PCA for T18S_SobAM2_CreNeg ####
# Must drop not numerical values, aka : sample.number, ...1, cre.and.treatment
PCA_T18S_SobAM2_CreNeg <- T18S_SobAM2_CreNeg %>% select(contains("Total"))
PCA_T18S_SobAM2_CreNeg <- mutate_all(PCA_T18S_SobAM2_CreNeg, function(x) as.numeric(as.character(x)))

PrincipleComponent_T18S_SobAM2_CreNeg <- prcomp(PCA_T18S_SobAM2_CreNeg, scale. = TRUE)
PrincipleComponent_T18S_SobAM2_CreNeg$rotation <- -1*PrincipleComponent_T18S_SobAM2_CreNeg$rotation
summary(PrincipleComponent_T18S_SobAM2_CreNeg)

#82.05% in 2 PC

## Biplot of PCA
pdf("Supplemental/Biplot_T18S_SobAM2_CreNeg_PLP.pdf", width = 6, height = 4)
fviz_pca_biplot(PrincipleComponent_T18S_SobAM2_CreNeg, repel = TRUE,
                col.var = "#2E9FDF",
                col.ind = "#696969", title = "PLP W18 CRE- SobAM2 spinal cord totals")
dev.off()

## Grouping PCA visual of CRE +/-
pdf("Adjusted_PCA/Grouping_T18S_SobAM2_CreNeg_PLP.pdf", width = 1.75, height = 1.17)
groups <- as.factor(T18S_SobAM2_CreNeg$CRE.and.Treatment)
fviz_pca_ind(PrincipleComponent_T18S_SobAM2_CreNeg,
             col.ind = groups, palette = c("#00AFBB", "#FC4E07"), addEllipses = TRUE, ellipse.type = "confidence",
             legend.title = NULL, pointsize = 1, repel = TRUE, title = NULL, labelsize = 2, 
             ellipse.alpha = 0.3, ellipse.border.remove = TRUE, mean.point = FALSE
) + theme(legend.position = "none",
          text = element_text(size = 0),
          axis.text.x = element_text(face="plain", color="black", size=5, angle=0),
          axis.text.y = element_text(face="plain", color="black", size=5, angle=0),
          axis.ticks.length=unit(.05, "cm"), axis.ticks=element_line(linewidth=0.2),
          axis.title.x = element_text(family = "sans", size = 5, margin=margin(1,0,0,0)), 
          axis.title.y = element_text(family = "sans", size = 5, margin=margin(0,1,0,0)),
          panel.grid.major = element_line(linewidth = 0.5), 
          panel.grid.minor = element_line(linewidth = 0.5),
          plot.margin = margin(4,4,4,4)
)
dev.off()


# PCA Serum ####

T6_18Srm <- read_csv("Final_6Srm_Total_noCE.csv", col_names = TRUE, name_repair = unique)

## T6 = W6 CRE+ totals vs W6 CRE- totals ####

T6Srm <- T6_18Srm %>% filter(grepl('wk 6', ...1)) %>% filter(grepl('Control', CRE.and.Treatment))

### PCA for T6Srm ####
# Must drop not numerical values, aka : sample.number, ...1, cre.and.treatment
PCA_T6Srm <- T6Srm %>% select(contains("Total"))
PCA_T6Srm <- mutate_all(PCA_T6Srm, function(x) as.numeric(as.character(x)))

PrincipleComponent_T6Srm <- prcomp(PCA_T6Srm, scale. = TRUE)
PrincipleComponent_T6Srm$rotation <- -1*PrincipleComponent_T6Srm$rotation
summary(PrincipleComponent_T6Srm)

# 77.19% in 2 PC

## Biplot of PCA
pdf("Serum/Biplot_T6Srm_PLP.pdf", width = 6, height = 4)
fviz_pca_biplot(PrincipleComponent_T6Srm, repel = TRUE,
                col.var = "#2E9FDF",
                col.ind = "#696969", title = "PLP W6 serum totals")
dev.off()

## Grouping PCA visual of CRE +/-
pdf("Serum/Grouping_T6Srm_PLP.pdf", width = 1.75, height = 1.17)
groups <- as.factor(T6Srm$CRE.and.Treatment)
fviz_pca_ind(PrincipleComponent_T6Srm,
             col.ind = groups, palette = c("#00AFBB", "#FC4E07"), addEllipses = TRUE, ellipse.type = "confidence",
             legend.title = NULL, pointsize = 1, repel = TRUE, title = NULL, labelsize = 2, 
             ellipse.alpha = 0.3, ellipse.border.remove = TRUE, mean.point = FALSE
) + theme(legend.position = "none",
          text = element_text(size = 0),
          axis.text.x = element_text(face="plain", color="black", size=5, angle=0),
          axis.text.y = element_text(face="plain", color="black", size=5, angle=0),
          axis.ticks.length=unit(.05, "cm"), axis.ticks=element_line(linewidth=0.2),
          axis.title.x = element_text(family = "sans", size = 5, margin=margin(1,0,0,0)), 
          axis.title.y = element_text(family = "sans", size = 5, margin=margin(0,1,0,0)),
          panel.grid.major = element_line(linewidth = 0.5), 
          panel.grid.minor = element_line(linewidth = 0.5),
          plot.margin = margin(4,4,4,4)
)
dev.off()

## T12 = W12 CRE+ totals vs W12 CRE- totals ####

T12Srm <- T6_18Srm %>% filter(grepl('wk 12', ...1)) %>% filter(grepl('Control', CRE.and.Treatment))

### PCA for T12Srm ####
# Must drop not numerical values, aka : sample.number, ...1, cre.and.treatment
PCA_T12Srm <- T12Srm %>% select(contains("Total"))
PCA_T12Srm <- mutate_all(PCA_T12Srm, function(x) as.numeric(as.character(x)))

PrincipleComponent_T12Srm <- prcomp(PCA_T12Srm, scale. = TRUE)
PrincipleComponent_T12Srm$rotation <- -1*PrincipleComponent_T12Srm$rotation
summary(PrincipleComponent_T12Srm)

# 76.25% in 2 PC

## Biplot of PCA
pdf("Serum/Biplot_T12Srm_PLP.pdf", width = 6, height = 4)
fviz_pca_biplot(PrincipleComponent_T12Srm, repel = TRUE,
                col.var = "#2E9FDF",
                col.ind = "#696969", title = "PLP W12 serum totals")
dev.off()

## Grouping PCA visual of CRE +/-
pdf("Serum/Grouping_T12Srm_PLP.pdf", width = 1.75, height = 1.17)
groups <- as.factor(T12Srm$CRE.and.Treatment)
fviz_pca_ind(PrincipleComponent_T12Srm,
             col.ind = groups, palette = c("#00AFBB", "#FC4E07"), addEllipses = TRUE, ellipse.type = "confidence",
             legend.title = NULL, pointsize = 1, repel = TRUE, title = NULL, labelsize = 2, 
             ellipse.alpha = 0.3, ellipse.border.remove = TRUE, mean.point = FALSE
) + theme(legend.position = "none",
          text = element_text(size = 0),
          axis.text.x = element_text(face="plain", color="black", size=5, angle=0),
          axis.text.y = element_text(face="plain", color="black", size=5, angle=0),
          axis.ticks.length=unit(.05, "cm"), axis.ticks=element_line(linewidth=0.2),
          axis.title.x = element_text(family = "sans", size = 5, margin=margin(1,0,0,0)), 
          axis.title.y = element_text(family = "sans", size = 5, margin=margin(0,1,0,0)),
          panel.grid.major = element_line(linewidth = 0.5), 
          panel.grid.minor = element_line(linewidth = 0.5),
          plot.margin = margin(4,4,4,4)
)
dev.off()

## T18 = W18 CRE+ totals vs W18 CRE- totals ####

T18Srm <- T6_18Srm %>% filter(grepl('wk 18', ...1)) %>% filter(grepl('Control', CRE.and.Treatment))

### PCA for T18Srm ####
# Must drop not numerical values, aka : sample.number, ...1, cre.and.treatment
PCA_T18Srm <- T18Srm %>% select(contains("Total"))
PCA_T18Srm <- mutate_all(PCA_T18Srm, function(x) as.numeric(as.character(x)))

PrincipleComponent_T18Srm <- prcomp(PCA_T18Srm, scale. = TRUE)
PrincipleComponent_T18Srm$rotation <- -1*PrincipleComponent_T18Srm$rotation
summary(PrincipleComponent_T18Srm)

# 78.74% in 2 PC

## Biplot of PCA
pdf("Serum/Biplot_T18Srm_PLP.pdf", width = 6, height = 4)
fviz_pca_biplot(PrincipleComponent_T18Srm, repel = TRUE,
                col.var = "#2E9FDF",
                col.ind = "#696969", title = "PLP W18 serum totals")
dev.off()

## Grouping PCA visual of CRE +/-
pdf("Serum/Grouping_T18Srm_PLP.pdf", width = 1.75, height = 1.17)
groups <- as.factor(T18Srm$CRE.and.Treatment)
fviz_pca_ind(PrincipleComponent_T18Srm,
             col.ind = groups, palette = c("#00AFBB", "#FC4E07"), addEllipses = TRUE, ellipse.type = "confidence",
             legend.title = NULL, pointsize = 1, repel = TRUE, title = NULL, labelsize = 2, 
             ellipse.alpha = 0.3, ellipse.border.remove = TRUE, mean.point = FALSE
) + theme(legend.position = "none",
          text = element_text(size = 0),
          axis.text.x = element_text(face="plain", color="black", size=5, angle=0),
          axis.text.y = element_text(face="plain", color="black", size=5, angle=0),
          axis.ticks.length=unit(.05, "cm"), axis.ticks=element_line(linewidth=0.2),
          axis.title.x = element_text(family = "sans", size = 5, margin=margin(1,0,0,0)), 
          axis.title.y = element_text(family = "sans", size = 5, margin=margin(0,1,0,0)),
          panel.grid.major = element_line(linewidth = 0.5), 
          panel.grid.minor = element_line(linewidth = 0.5),
          plot.margin = margin(4,4,4,4)
)
dev.off()

## T12 w SobAM2 = W12 CRE+ totals vs W12 CRE+ w SobAM2 totals ####

T12Srm_SobAM2_CrePos <- T6_18Srm %>% filter(grepl('wk 12', ...1)) %>% filter(!grepl('-', CRE.and.Treatment))

### PCA for T12Srm_SobAM2_CrePos ####
# Must drop not numerical values, aka : sample.number, ...1, cre.and.treatment
PCA_T12Srm_SobAM2_CrePos <- T12Srm_SobAM2_CrePos %>% select(contains("Total"))
PCA_T12Srm_SobAM2_CrePos <- mutate_all(PCA_T12Srm_SobAM2_CrePos, function(x) as.numeric(as.character(x)))

PrincipleComponent_T12Srm_SobAM2_CrePos <- prcomp(PCA_T12Srm_SobAM2_CrePos, scale. = TRUE)
PrincipleComponent_T12Srm_SobAM2_CrePos$rotation <- -1*PrincipleComponent_T12Srm_SobAM2_CrePos$rotation
summary(PrincipleComponent_T12Srm_SobAM2_CrePos)

# 73.80% in 2 PC

## Biplot of PCA
pdf("Serum/Biplot_T12Srm_SobAM2_CrePos_PLP.pdf", width = 6, height = 4)
fviz_pca_biplot(PrincipleComponent_T12Srm_SobAM2_CrePos, repel = TRUE,
                col.var = "#2E9FDF",
                col.ind = "#696969", title = "PLP W12 CRE+ SobAM2 brain totals")
dev.off()

## Grouping PCA visual of CRE +/-
pdf("Serum/Grouping_T12Srm_SobAM2_CrePos_PLP.pdf", width = 1.75, height = 1.17)
groups <- as.factor(T12Srm_SobAM2_CrePos$CRE.and.Treatment)
fviz_pca_ind(PrincipleComponent_T12Srm_SobAM2_CrePos,
             col.ind = groups, palette = c("#00AFBB", "#FC4E07"), addEllipses = TRUE, ellipse.type = "confidence",
             legend.title = NULL, pointsize = 1, repel = TRUE, title = NULL, labelsize = 2, 
             ellipse.alpha = 0.3, ellipse.border.remove = TRUE, mean.point = FALSE
) + theme(legend.position = "none",
          text = element_text(size = 0),
          axis.text.x = element_text(face="plain", color="black", size=5, angle=0),
          axis.text.y = element_text(face="plain", color="black", size=5, angle=0),
          axis.ticks.length=unit(.05, "cm"), axis.ticks=element_line(linewidth=0.2),
          axis.title.x = element_text(family = "sans", size = 5, margin=margin(1,0,0,0)), 
          axis.title.y = element_text(family = "sans", size = 5, margin=margin(0,1,0,0)),
          panel.grid.major = element_line(linewidth = 0.5), 
          panel.grid.minor = element_line(linewidth = 0.5),
          plot.margin = margin(4,4,4,4)
)
dev.off()

## T12 w SobAM2 = W12 CRE- totals vs W12 CRE- w SobAM2 totals ####

T12Srm_SobAM2_CreNeg <- T6_18Srm %>% filter(grepl('wk 12', ...1)) %>% filter(grepl('-', CRE.and.Treatment))

### PCA for T12Srm_SobAM2_CreNeg ####
# Must drop not numerical values, aka : sample.number, ...1, cre.and.treatment
PCA_T12Srm_SobAM2_CreNeg <- T12Srm_SobAM2_CreNeg %>% select(contains("Total"))
PCA_T12Srm_SobAM2_CreNeg <- mutate_all(PCA_T12Srm_SobAM2_CreNeg, function(x) as.numeric(as.character(x)))

PrincipleComponent_T12Srm_SobAM2_CreNeg <- prcomp(PCA_T12Srm_SobAM2_CreNeg, scale. = TRUE)
PrincipleComponent_T12Srm_SobAM2_CreNeg$rotation <- -1*PrincipleComponent_T12Srm_SobAM2_CreNeg$rotation
summary(PrincipleComponent_T12Srm_SobAM2_CreNeg)

# 80.66% in 2 PC

## Biplot of PCA
pdf("Serum/Biplot_T12Srm_SobAM2_CreNeg_PLP.pdf", width = 6, height = 4)
fviz_pca_biplot(PrincipleComponent_T12Srm_SobAM2_CreNeg, repel = TRUE,
                col.var = "#2E9FDF",
                col.ind = "#696969", title = "PLP W12 CRE- SobAM2 brain totals")
dev.off()

## Grouping PCA visual of CRE +/-
pdf("Serum/Grouping_T12Srm_SobAM2_CreNeg_PLP.pdf", width = 1.75, height = 1.17)
groups <- as.factor(T12Srm_SobAM2_CreNeg$CRE.and.Treatment)
fviz_pca_ind(PrincipleComponent_T12Srm_SobAM2_CreNeg,
             col.ind = groups, palette = c("#00AFBB", "#FC4E07"), addEllipses = TRUE, ellipse.type = "confidence",
             legend.title = NULL, pointsize = 1, repel = TRUE, title = NULL, labelsize = 2, 
             ellipse.alpha = 0.3, ellipse.border.remove = TRUE, mean.point = FALSE
) + theme(legend.position = "none",
          text = element_text(size = 0),
          axis.text.x = element_text(face="plain", color="black", size=5, angle=0),
          axis.text.y = element_text(face="plain", color="black", size=5, angle=0),
          axis.ticks.length=unit(.05, "cm"), axis.ticks=element_line(linewidth=0.2),
          axis.title.x = element_text(family = "sans", size = 5, margin=margin(1,0,0,0)), 
          axis.title.y = element_text(family = "sans", size = 5, margin=margin(0,1,0,0)),
          panel.grid.major = element_line(linewidth = 0.5), 
          panel.grid.minor = element_line(linewidth = 0.5),
          plot.margin = margin(4,4,4,4)
)
dev.off()

## T18 w SobAM2 = W18 CRE+ totals vs W18 CRE+ w SobAM2 totals ####

T18Srm_SobAM2_CrePos <- T6_18Srm %>% filter(grepl('wk 18', ...1)) %>% filter(!grepl('-', CRE.and.Treatment))

### PCA for T18Srm_SobAM2_CrePos ####
# Must drop not numerical values, aka : sample.number, ...1, cre.and.treatment
PCA_T18Srm_SobAM2_CrePos <- T18Srm_SobAM2_CrePos %>% select(contains("Total"))
PCA_T18Srm_SobAM2_CrePos <- mutate_all(PCA_T18Srm_SobAM2_CrePos, function(x) as.numeric(as.character(x)))

PrincipleComponent_T18Srm_SobAM2_CrePos <- prcomp(PCA_T18Srm_SobAM2_CrePos, scale. = TRUE)
PrincipleComponent_T18Srm_SobAM2_CrePos$rotation <- -1*PrincipleComponent_T18Srm_SobAM2_CrePos$rotation
summary(PrincipleComponent_T18Srm_SobAM2_CrePos)

# 67.84% in 2 PC

## Biplot of PCA
pdf("Serum/Biplot_T18Srm_SobAM2_CrePos_PLP.pdf", width = 6, height = 4)
fviz_pca_biplot(PrincipleComponent_T18Srm_SobAM2_CrePos, repel = TRUE,
                col.var = "#2E9FDF",
                col.ind = "#696969", title = "PLP W18 CRE+ SobAM2 brain totals")
dev.off()

## Grouping PCA visual of CRE +/-
pdf("Serum/Grouping_T18Srm_SobAM2_CrePos_PLP.pdf", width = 1.75, height = 1.17)
groups <- as.factor(T18Srm_SobAM2_CrePos$CRE.and.Treatment)
fviz_pca_ind(PrincipleComponent_T18Srm_SobAM2_CrePos,
             col.ind = groups, palette = c("#00AFBB", "#FC4E07"), addEllipses = TRUE, ellipse.type = "confidence",
             legend.title = NULL, pointsize = 1, repel = TRUE, title = NULL, labelsize = 2, 
             ellipse.alpha = 0.3, ellipse.border.remove = TRUE, mean.point = FALSE
) + theme(legend.position = "none",
          text = element_text(size = 0),
          axis.text.x = element_text(face="plain", color="black", size=5, angle=0),
          axis.text.y = element_text(face="plain", color="black", size=5, angle=0),
          axis.ticks.length=unit(.05, "cm"), axis.ticks=element_line(linewidth=0.2),
          axis.title.x = element_text(family = "sans", size = 5, margin=margin(1,0,0,0)), 
          axis.title.y = element_text(family = "sans", size = 5, margin=margin(0,1,0,0)),
          panel.grid.major = element_line(linewidth = 0.5), 
          panel.grid.minor = element_line(linewidth = 0.5),
          plot.margin = margin(4,4,4,4)
)
dev.off()

## T18 w SobAM2 = W18 CRE- totals vs W18 CRE- w SobAM2 totals ####

T18Srm_SobAM2_CreNeg <- T6_18Srm %>% filter(grepl('wk 18', ...1)) %>% filter(grepl('-', CRE.and.Treatment))

### PCA for T18Srm_SobAM2_CreNeg ####
# Must drop not numerical values, aka : sample.number, ...1, cre.and.treatment
PCA_T18Srm_SobAM2_CreNeg <- T18Srm_SobAM2_CreNeg %>% select(contains("Total"))
PCA_T18Srm_SobAM2_CreNeg <- mutate_all(PCA_T18Srm_SobAM2_CreNeg, function(x) as.numeric(as.character(x)))

PrincipleComponent_T18Srm_SobAM2_CreNeg <- prcomp(PCA_T18Srm_SobAM2_CreNeg, scale. = TRUE)
PrincipleComponent_T18Srm_SobAM2_CreNeg$rotation <- -1*PrincipleComponent_T18Srm_SobAM2_CreNeg$rotation
summary(PrincipleComponent_T18Srm_SobAM2_CreNeg)

# 64.06% in 2 PC

## Biplot of PCA
pdf("Serum/Biplot_T18Srm_SobAM2_CreNeg_PLP.pdf", width = 6, height = 4)
fviz_pca_biplot(PrincipleComponent_T18Srm_SobAM2_CreNeg, repel = TRUE,
                col.var = "#2E9FDF",
                col.ind = "#696969", title = "PLP W18 CRE- SobAM2 brain totals")
dev.off()

## Grouping PCA visual of CRE +/-
pdf("Serum/Grouping_T18Srm_SobAM2_CreNeg_PLP.pdf", width = 1.75, height = 1.17)
groups <- as.factor(T18Srm_SobAM2_CreNeg$CRE.and.Treatment)
fviz_pca_ind(PrincipleComponent_T18Srm_SobAM2_CreNeg,
             col.ind = groups, palette = c("#00AFBB", "#FC4E07"), addEllipses = TRUE, ellipse.type = "confidence",
             legend.title = NULL, pointsize = 1, repel = TRUE, title = NULL, labelsize = 2, 
             ellipse.alpha = 0.3, ellipse.border.remove = TRUE, mean.point = FALSE
) + theme(legend.position = "none",
          text = element_text(size = 0),
          axis.text.x = element_text(face="plain", color="black", size=5, angle=0),
          axis.text.y = element_text(face="plain", color="black", size=5, angle=0),
          axis.ticks.length=unit(.05, "cm"), axis.ticks=element_line(linewidth=0.2),
          axis.title.x = element_text(family = "sans", size = 5, margin=margin(1,0,0,0)), 
          axis.title.y = element_text(family = "sans", size = 5, margin=margin(0,1,0,0)),
          panel.grid.major = element_line(linewidth = 0.5), 
          panel.grid.minor = element_line(linewidth = 0.5),
          plot.margin = margin(4,4,4,4)
)
dev.off()
```
