## Supplemental Codes for "Lipidomic analysis reveals differences in the extent of remyelination in the brain and spinal cord": SobAM2_AllGroups_PCA.R

```
# PCA plots for SobAM2 effects on brain and spinal cord

# Library

library(dplyr)
library(readxl)
library(tidyverse)
library(igraph)
library(ggplot2)    
library(data.table)
library(factoextra)

# Load data

T12_18B <- read_csv("Final_6B_Total_noCE.csv", col_names = TRUE, name_repair = unique)
rownames(T12_18B) <- T12_18B$...1
T12B <- T12_18B %>% filter(grepl('wk 12', ...1)) 
T18B <- T12_18B %>% filter(grepl('wk 18', ...1)) 

T12_18S <- read_csv("Final_6S_Total_noCE_w_LPE.csv", col_names = TRUE, name_repair = unique)
rownames(T12_18S) <- T12_18S$...1
T12S <- T12_18S %>% filter(grepl('wk 12', ...1)) 
T18S <- T12_18S %>% filter(grepl('wk 18', ...1)) 

# Plot PCAs

# W12 Brain ####

# Drop non-numerical values, aka : sample.number, ...1, cre.and.treatment
PCA_T12B <- T12B %>% select(contains("Total"))

PrincipleComponent_T12B <- prcomp(PCA_T12B, scale. = TRUE)
PrincipleComponent_T12B$rotation <- -1*PrincipleComponent_T12B$rotation
summary(PrincipleComponent_T12B)

## Biplot of PCA
pdf("Biplot_T12B_all_PLP.pdf", width = 6, height = 4)
fviz_pca_biplot(PrincipleComponent_T12B, repel = TRUE,
                col.var = "#2E9FDF",
                col.ind = "#696969", title = NULL)
dev.off()

## Grouping PCA visual of all groups in W12
pdf("Grouping_T12B_all_PLP.pdf", width = 6, height = 4)
groups <- as.factor(T12B$CRE.and.Treatment)
fviz_pca_ind(PrincipleComponent_T12B,
             col.ind = groups, palette = c("#00AFBB", "#FC4E07", "#33B631","#6B409F"), addEllipses = TRUE, ellipse.type = "confidence",
             repel = TRUE, title = NULL
) + theme (legend.position = "none", plot.title = element_blank())

dev.off()

# W12 Spinal Cord ####

# Drop non-numerical values, aka : sample.number, ...1, cre.and.treatment
PCA_T12S <- T12S %>% select(contains("Total"))

PrincipleComponent_T12S <- prcomp(PCA_T12S, scale. = TRUE)
PrincipleComponent_T12S$rotation <- -1*PrincipleComponent_T12S$rotation
summary(PrincipleComponent_T12S)

## Biplot of PCA
pdf("Biplot_T12S_all_PLP.pdf", width = 6, height = 4)
fviz_pca_biplot(PrincipleComponent_T12S, repel = TRUE,
                col.var = "#2E9FDF",
                col.ind = "#696969", title = NULL)
dev.off()

## Grouping PCA visual of all groups in W12
pdf("Grouping_T12S_all_PLP.pdf", width = 6, height = 4)
groups <- as.factor(T12S$CRE.and.Treatment)
fviz_pca_ind(PrincipleComponent_T12S,
             col.ind = groups, palette = c("#00AFBB", "#FC4E07", "#33B631","#6B409F"), addEllipses = TRUE, ellipse.type = "confidence",
             repel = TRUE, title = NULL
) + theme (legend.position = "none", plot.title = element_blank())

dev.off()

# W18 Brain ####
PCA_T18B <- T18B %>% select(contains("Total"))

PrincipleComponent_T18B <- prcomp(PCA_T18B, scale. = TRUE)
PrincipleComponent_T18B$rotation <- -1*PrincipleComponent_T18B$rotation
summary(PrincipleComponent_T18B)

## Biplot of PCA
pdf("Biplot_T18B_all_PLP.pdf", width = 6, height = 4)
fviz_pca_biplot(PrincipleComponent_T18B, repel = TRUE,
                col.var = "#2E9FDF",
                col.ind = "#696969", title = NULL)
dev.off()

## Grouping PCA visual of all groups in W18
pdf("Grouping_T18B_all_PLP.pdf", width = 6, height = 4)
groups <- as.factor(T18B$CRE.and.Treatment)
fviz_pca_ind(PrincipleComponent_T18B,
             col.ind = groups, palette = c("#00AFBB", "#FC4E07", "#33B631","#6B409F"), addEllipses = TRUE, ellipse.type = "confidence",
             repel = TRUE, title = NULL
) + theme (legend.position = "none", plot.title = element_blank())

dev.off()

# W18 Spinal Cord ####
PCA_T18S <- T18S %>% select(contains("Total"))

PrincipleComponent_T18S <- prcomp(PCA_T18S, scale. = TRUE)
PrincipleComponent_T18S$rotation <- -1*PrincipleComponent_T18S$rotation
summary(PrincipleComponent_T18S)

## Biplot of PCA
pdf("Biplot_T18S_all_PLP.pdf", width = 6, height = 4)
fviz_pca_biplot(PrincipleComponent_T18S, repel = TRUE,
                col.var = "#2E9FDF",
                col.ind = "#696969", title = NULL)
dev.off()

## Grouping PCA visual of all groups in W18
pdf("Grouping_T18S_all_PLP.pdf", width = 6, height = 4)
groups <- as.factor(T18S$CRE.and.Treatment)
fviz_pca_ind(PrincipleComponent_T18S,
             col.ind = groups, palette = c("#00AFBB", "#FC4E07", "#33B631","#6B409F"), addEllipses = TRUE, ellipse.type = "confidence",
             repel = TRUE, title = NULL
) + theme (legend.position = "none", plot.title = element_blank())

dev.off()
```
