## Supplemental Codes for "Lipidomic analysis reveals differences in the extent of remyelination in the brain and spinal cord": VP_serum.R

```
# Volcano Plots

# Load library #####
library(dplyr)
library(readxl)
library(tidyverse)
library(data.table)
library(igraph)
library(ggplot2)
library(countdata)
library(gtools)
library(devtools)
library(permFDP) #install_github('steven-shuken/permFDP')
library(EnhancedVolcano) #install_github('kevinblighe/EnhancedVolcano')
library(plotrix)
#library(ggrastr)

A6_18_Srm <- read_csv("Final_6Srm_All_noCE.csv", col_names = TRUE, name_repair = unique)

# Functions ####
mean.sem <- function(x) {
  c(mean = mean(x), sem = sd(x)/sqrt(length(x)))
}


# PLP W6 Serum ####

A6Srm <- A6_18_Srm %>% filter(grepl('wk 6', ...1)) %>% filter(grepl('Control', CRE.and.Treatment))
A6Srm_no0 <- A6Srm %>% select(-"Sample.number", -"...1")

# For no 0 df, drop PC(40:3), PC(44:12) and PE(40:5)
# https://stackoverflow.com/questions/72096913/remove-columns-that-have-one-zero-value
print(select(A6Srm_no0, where(~any(. == 0))))
A6Srm_no0 <- Filter(function(x) !any(x %in% 0), A6Srm_no0) 
log_A6Srm_no0 = log2(A6Srm_no0[,-ncol(A6Srm_no0)]) 
log_A6Srm_no0 <- cbind(log_A6Srm_no0, 'CRE.and.Treatment' = A6Srm$CRE.and.Treatment)

# Fix column names & remove extra rows
ttest_A6Srm_no0 <- transpose(log_A6Srm_no0, keep.names = 'Lipid species')
ttest_A6Srm_no0 <- ttest_A6Srm_no0[-nrow(ttest_A6Srm_no0),]

samples <- A6Srm$Sample.number
names <- c('Lipid species', samples)
colnames(ttest_A6Srm_no0) = c(names)

# Get p values from t test 
ttest_A6Srm_no0$"pvalue" <- unlist(lapply(1:nrow(ttest_A6Srm_no0), function(i) t.test((as.numeric(ttest_A6Srm_no0[i,2:4])), as.numeric(ttest_A6Srm_no0[i,5:12]))[c("p.value")]))

# calculate average values & sem for each species using CRE and treatment as factor
mean.sem_A6Srm_no0 <- data.table(log_A6Srm_no0)[, lapply(.SD, mean.sem), by=CRE.and.Treatment]
mean.sem_A6Srm_no0 <- transpose(mean.sem_A6Srm_no0[,2:(ncol(mean.sem_A6Srm_no0))], keep.names = "Lipid species")
# Make sure the order of columns is correct, some df will have experimental first
CRE_factor <- c("CRE-/control avg","CRE-/control sem","CRE+/control avg","CRE+/control sem")
names <- c('Lipid species', CRE_factor)
colnames(mean.sem_A6Srm_no0) = c(names)

# add means to main df
#VP_A6Srm_no0 <- cbind(ttest_A6Srm_no0, mean.sem_A6Srm_no0)
VP_A6Srm_no0 <- merge(ttest_A6Srm_no0, mean.sem_A6Srm_no0, by = 'Lipid species')
# Calculate fold change
VP_A6Srm_no0$"Log2FC" <- VP_A6Srm_no0$'CRE+/control avg' - VP_A6Srm_no0$'CRE-/control avg'

#save df 
write_csv(VP_A6Srm_no0, "Serum/A6Srm_no0_Volcano_plot.csv")

## permFDP ####

VP_A6Srm_no0 <- read_csv("Serum/A6Srm_no0_Volcano_plot.csv")

pVals = VP_A6Srm_no0$pvalue
threshold = 0.05
myDesign = c(1,1,1,2,2,2,2,2,2,2,2)
intOnly = VP_A6Srm_no0 %>% select(-c("Lipid species","pvalue","CRE-/control avg","CRE-/control sem","CRE+/control avg","CRE+/control sem","Log2FC"))
intOnly <- mutate_all(intOnly, function(x) as.numeric(as.character(x))) 
corrThreshold_500 = permFDP::permFDP.adjust.threshold(pVals, threshold, myDesign, intOnly, 500)
corrThreshold_1000 = permFDP::permFDP.adjust.threshold(pVals, threshold, myDesign, intOnly, 1000)
corrThreshold_3000 = permFDP::permFDP.adjust.threshold(pVals, threshold, myDesign, intOnly, 3000)
corrThreshold_10000 = permFDP::permFDP.adjust.threshold(pVals, threshold, myDesign, intOnly, 10000)

print(corrThreshold_3000, digits = 20)
columns <- c("Timepoint", "Threshold")
rows <- c("A6Srm","A12Srm","A18Srm","A12Srm_SobAM2_CrePos","A12Srm_SobAM2_CreNeg",
          "A18Srm_SobAM2_CrePos","A18Srm_SobAM2_CreNeg")
FDPthreshold = data.frame(matrix(nrow = length(rows), ncol = length(columns))) 
colnames(FDPthreshold) = columns
FDPthreshold$Timepoint <- rows
FDPthreshold[1,2] <- corrThreshold_3000
#corrThreshold_3000 = 0.062171252216525522971

## EnhancedVolcano ####
pdf("Serum/VP_A6Srm_no0.pdf", width = 11.69, height = 8.27)

# Lipid species, p values, log2 FCs
toptable <- VP_A6Srm_no0[, c(1,ncol(VP_A6Srm_no0)-5,ncol(VP_A6Srm_no0))]
lab = toptable$"Lipid species"
x = "Log2FC"
y = "pvalue"
pCutoff = corrThreshold_3000
FCcutoff = 0.5
title = "PLP serum samples"
subtitle = bquote(italic('Week 6 with no 0 values, no CE, 3000 iterations, 11 mice'))

keyvals.colour <- ifelse(
  toptable$Log2FC > 0.5 & toptable$pvalue < pCutoff, '#FF00FF',
  ifelse(toptable$Log2FC <(-0.5) & toptable$pvalue < pCutoff, '#00FFFF',
         '#808080'))
# Gray - #A8B6CC - #DAECCF
# Magenta - #F05039 - #F32A50
# Cyan - #0090FF - #0882E0

keyvals.colour[is.na(keyvals.colour)] <- '#808080'
names(keyvals.colour)[keyvals.colour == '#FF00FF'] <- 'High FC & sig'
names(keyvals.colour)[keyvals.colour == '#808080'] <- 'NS'
names(keyvals.colour)[keyvals.colour == '#00FFFF'] <- 'Low FC & sig'

EnhancedVolcano(toptable, lab, x, y, selectLab = rownames(toptable)[which(names(keyvals.colour) %in% c('High', 'Low'))],
                xlab = bquote(~Log[2] ~ "Fold Change"),
                ylab = bquote(~-Log[10] ~ italic(p) ~ Value), colCustom = keyvals.colour,
                title = title, subtitle = subtitle, pCutoff = pCutoff, pCutoffCol = y, FCcutoff = FCcutoff,labSize = 5,
                drawConnectors = FALSE
)
dev.off()

# PLP W12 Serum ####

A12Srm <- A6_18_Srm %>% filter(grepl('wk 12', ...1)) %>% filter(grepl('Control', CRE.and.Treatment))
A12Srm_no0 <- A12Srm %>% select(-"Sample.number", -"...1")

# For no 0 df, drop PC(40:3), PC(42:11), PC(44:12), PE(40:5), and ePE(36.5)
# https://stackoverflow.com/questions/72096913/remove-columns-that-have-one-zero-value
print(select(A12Srm_no0, where(~any(. == 0))))
A12Srm_no0 <- Filter(function(x) !any(x %in% 0), A12Srm_no0) 
log_A12Srm_no0 = log2(A12Srm_no0[,-ncol(A12Srm_no0)]) 
log_A12Srm_no0 <- cbind(log_A12Srm_no0, 'CRE.and.Treatment' = A12Srm$CRE.and.Treatment)

# Fix column names & remove extra rows
ttest_A12Srm_no0 <- transpose(log_A12Srm_no0, keep.names = 'Lipid species')
ttest_A12Srm_no0 <- ttest_A12Srm_no0[-nrow(ttest_A12Srm_no0),]

samples <- A12Srm$Sample.number
names <- c('Lipid species', samples)
colnames(ttest_A12Srm_no0) = c(names)

# Get p values from t test 
ttest_A12Srm_no0$"pvalue" <- unlist(lapply(1:nrow(ttest_A12Srm_no0), function(i) t.test((as.numeric(ttest_A12Srm_no0[i,2:6])), as.numeric(ttest_A12Srm_no0[i,7:14]))[c("p.value")]))

# calculate average values & sem for each species using CRE and treatment as factor
mean.sem_A12Srm_no0 <- data.table(log_A12Srm_no0)[, lapply(.SD, mean.sem), by=CRE.and.Treatment]
mean.sem_A12Srm_no0 <- transpose(mean.sem_A12Srm_no0[,2:(ncol(mean.sem_A12Srm_no0))], keep.names = "Lipid species")
# Make sure the order of columns is correct, some df will have experimental first
CRE_factor <- c("CRE-/control avg","CRE-/control sem","CRE+/control avg","CRE+/control sem")
names <- c('Lipid species', CRE_factor)
colnames(mean.sem_A12Srm_no0) = c(names)

# add means to main df
#VP_A12Srm_no0 <- cbind(ttest_A12Srm_no0, mean.sem_A12Srm_no0)
VP_A12Srm_no0 <- merge(ttest_A12Srm_no0, mean.sem_A12Srm_no0, by = 'Lipid species')
# Calculate fold change
VP_A12Srm_no0$"Log2FC" <- VP_A12Srm_no0$'CRE+/control avg' - VP_A12Srm_no0$'CRE-/control avg'

#save df 
write_csv(VP_A12Srm_no0, "Serum/A12Srm_no0_Volcano_plot.csv")

## permFDP ####

VP_A12Srm_no0 <- read_csv("Serum/A12Srm_no0_Volcano_plot.csv")

pVals = VP_A12Srm_no0$pvalue
threshold = 0.05
myDesign = c(1,1,1,1,1,2,2,2,2,2,2,2,2)
intOnly = VP_A12Srm_no0 %>% select(-c("Lipid species","pvalue","CRE-/control avg","CRE-/control sem","CRE+/control avg","CRE+/control sem","Log2FC"))
intOnly <- mutate_all(intOnly, function(x) as.numeric(as.character(x))) 
corrThreshold_500 = permFDP::permFDP.adjust.threshold(pVals, threshold, myDesign, intOnly, 500)
corrThreshold_1000 = permFDP::permFDP.adjust.threshold(pVals, threshold, myDesign, intOnly, 1000)
corrThreshold_3000 = permFDP::permFDP.adjust.threshold(pVals, threshold, myDesign, intOnly, 3000)
corrThreshold_10000 = permFDP::permFDP.adjust.threshold(pVals, threshold, myDesign, intOnly, 10000)

print(corrThreshold_3000, digits = 20)
FDPthreshold[2,2] <- corrThreshold_3000
#corrThreshold_3000 = 0.00053779780440070502293

## EnhancedVolcano ####
pdf("Serum/VP_A12Srm_no0.pdf", width = 11.69, height = 8.27)

# Lipid species, p values, log2 FCs
toptable <- VP_A12Srm_no0[, c(1,ncol(VP_A12Srm_no0)-5,ncol(VP_A12Srm_no0))]
lab = toptable$"Lipid species"
x = "Log2FC"
y = "pvalue"
pCutoff = corrThreshold_3000
FCcutoff = 0.5
title = "PLP serum samples"
subtitle = bquote(italic('Week 12 with no 0 values, no CE, 3000 iterations, 13 mice'))

keyvals.colour <- ifelse(
  toptable$Log2FC > 0.5 & toptable$pvalue < pCutoff, '#FF00FF',
  ifelse(toptable$Log2FC <(-0.5) & toptable$pvalue < pCutoff, '#00FFFF',
         '#808080'))
# Gray - #A8B6CC - #DAECCF
# Magenta - #F05039 - #F32A50
# Cyan - #0090FF - #0882E0

keyvals.colour[is.na(keyvals.colour)] <- '#808080'
names(keyvals.colour)[keyvals.colour == '#FF00FF'] <- 'High FC & sig'
names(keyvals.colour)[keyvals.colour == '#808080'] <- 'NS'
names(keyvals.colour)[keyvals.colour == '#00FFFF'] <- 'Low FC & sig'

EnhancedVolcano(toptable, lab, x, y, selectLab = rownames(toptable)[which(names(keyvals.colour) %in% c('High', 'Low'))],
                xlab = bquote(~Log[2] ~ "Fold Change"),
                ylab = bquote(~-Log[10] ~ italic(p) ~ Value), colCustom = keyvals.colour,
                title = title, subtitle = subtitle, pCutoff = pCutoff, pCutoffCol = y, FCcutoff = FCcutoff,labSize = 5,
                drawConnectors = FALSE
)
dev.off()

# PLP W18 Serum ####

A18Srm <- A6_18_Srm %>% filter(grepl('wk 18', ...1)) %>% filter(grepl('Control', CRE.and.Treatment))
A18Srm_no0 <- A18Srm %>% select(-"Sample.number", -"...1")

# For no 0 df, drop PC(36:0) and PC(44:12)
# https://stackoverflow.com/questions/72096913/remove-columns-that-have-one-zero-value
print(select(A18Srm_no0, where(~any(. == 0))))
A18Srm_no0 <- Filter(function(x) !any(x %in% 0), A18Srm_no0) 
log_A18Srm_no0 = log2(A18Srm_no0[,-ncol(A18Srm_no0)]) 
log_A18Srm_no0 <- cbind(log_A18Srm_no0, 'CRE.and.Treatment' = A18Srm$CRE.and.Treatment)

# Fix column names & remove extra rows
ttest_A18Srm_no0 <- transpose(log_A18Srm_no0, keep.names = 'Lipid species')
ttest_A18Srm_no0 <- ttest_A18Srm_no0[-nrow(ttest_A18Srm_no0),]

samples <- A18Srm$Sample.number
names <- c('Lipid species', samples)
colnames(ttest_A18Srm_no0) = c(names)

# Get p values from t test 
ttest_A18Srm_no0$"pvalue" <- unlist(lapply(1:nrow(ttest_A18Srm_no0), function(i) t.test((as.numeric(ttest_A18Srm_no0[i,2:7])), as.numeric(ttest_A18Srm_no0[i,8:15]))[c("p.value")]))

# calculate average values & sem for each species using CRE and treatment as factor
mean.sem_A18Srm_no0 <- data.table(log_A18Srm_no0)[, lapply(.SD, mean.sem), by=CRE.and.Treatment]
mean.sem_A18Srm_no0 <- transpose(mean.sem_A18Srm_no0[,2:(ncol(mean.sem_A18Srm_no0))], keep.names = "Lipid species")
# Make sure the order of columns is correct, some df will have experimental first
CRE_factor <- c("CRE-/control avg","CRE-/control sem","CRE+/control avg","CRE+/control sem")
names <- c('Lipid species', CRE_factor)
colnames(mean.sem_A18Srm_no0) = c(names)

# add means to main df
#VP_A18Srm_no0 <- cbind(ttest_A18Srm_no0, mean.sem_A18Srm_no0)
VP_A18Srm_no0 <- merge(ttest_A18Srm_no0, mean.sem_A18Srm_no0, by = 'Lipid species')
# Calculate fold change
VP_A18Srm_no0$"Log2FC" <- VP_A18Srm_no0$'CRE+/control avg' - VP_A18Srm_no0$'CRE-/control avg'

#save df 
write_csv(VP_A18Srm_no0, "Serum/A18Srm_no0_Volcano_plot.csv")

## permFDP ####

VP_A18Srm_no0 <- read_csv("Serum/A18Srm_no0_Volcano_plot.csv")

pVals = VP_A18Srm_no0$pvalue
threshold = 0.05
myDesign = c(1,1,1,1,1,1,2,2,2,2,2,2,2,2)
intOnly = VP_A18Srm_no0 %>% select(-c("Lipid species","pvalue","CRE-/control avg","CRE-/control sem","CRE+/control avg","CRE+/control sem","Log2FC"))
intOnly <- mutate_all(intOnly, function(x) as.numeric(as.character(x))) 
corrThreshold_500 = permFDP::permFDP.adjust.threshold(pVals, threshold, myDesign, intOnly, 500)
corrThreshold_1000 = permFDP::permFDP.adjust.threshold(pVals, threshold, myDesign, intOnly, 1000)
corrThreshold_3000 = permFDP::permFDP.adjust.threshold(pVals, threshold, myDesign, intOnly, 3000)
corrThreshold_10000 = permFDP::permFDP.adjust.threshold(pVals, threshold, myDesign, intOnly, 10000)

print(corrThreshold_3000, digits = 20)
FDPthreshold[3,2] <- corrThreshold_3000
#corrThreshold_3000 = 0.020799368247483664141

## EnhancedVolcano ####
pdf("Serum/VP_A18Srm_no0.pdf", width = 11.69, height = 8.27)

# Lipid species, p values, log2 FCs
toptable <- VP_A18Srm_no0[, c(1,ncol(VP_A18Srm_no0)-5,ncol(VP_A18Srm_no0))]
lab = toptable$"Lipid species"
x = "Log2FC"
y = "pvalue"
pCutoff = corrThreshold_3000
FCcutoff = 0.5
title = "PLP serum samples"
subtitle = bquote(italic('Week 18 with no 0 values, no CE, 3000 iterations, 14 mice'))

keyvals.colour <- ifelse(
  toptable$Log2FC > 0.5 & toptable$pvalue < pCutoff, '#FF00FF',
  ifelse(toptable$Log2FC <(-0.5) & toptable$pvalue < pCutoff, '#00FFFF',
         '#808080'))
# Gray - #A8B6CC - #DAECCF
# Magenta - #F05039 - #F32A50
# Cyan - #0090FF - #0882E0

keyvals.colour[is.na(keyvals.colour)] <- '#808080'
names(keyvals.colour)[keyvals.colour == '#FF00FF'] <- 'High FC & sig'
names(keyvals.colour)[keyvals.colour == '#808080'] <- 'NS'
names(keyvals.colour)[keyvals.colour == '#00FFFF'] <- 'Low FC & sig'

EnhancedVolcano(toptable, lab, x, y, selectLab = rownames(toptable)[which(names(keyvals.colour) %in% c('High', 'Low'))],
                xlab = bquote(~Log[2] ~ "Fold Change"),
                ylab = bquote(~-Log[10] ~ italic(p) ~ Value), colCustom = keyvals.colour,
                title = title, subtitle = subtitle, pCutoff = pCutoff, pCutoffCol = y, FCcutoff = FCcutoff,labSize = 5,
                drawConnectors = FALSE
)
dev.off()

write_csv(FDPthreshold,"Serum/FDP_Threshold_Srm.csv")
FDPthreshold <- read_csv("Serum/FDP_Threshold_Srm.csv")

# PLP W12 Serum SobAM2 CrePos ####

A12Srm_SobAM2_CrePos <- A6_18_Srm %>% filter(grepl('wk 12', ...1)) %>% filter(!grepl('-', CRE.and.Treatment))
A12Srm_SobAM2_CrePos_no0 <- A12Srm_SobAM2_CrePos %>% select(-"Sample.number", -"...1")

# For no 0 df, drop PC(36:6), PC(38:6), PC(40:3), PC(42:11), PC(44:12), ePC(40:2), ePE(36.5),
# ePE(38:5), PI(34:1), PI(36:1), and PI(40:5)
# https://stackoverflow.com/questions/72096913/remove-columns-that-have-one-zero-value
print(select(A12Srm_SobAM2_CrePos_no0, where(~any(. == 0))))
A12Srm_SobAM2_CrePos_no0 <- Filter(function(x) !any(x %in% 0), A12Srm_SobAM2_CrePos_no0) 
log_A12Srm_SobAM2_CrePos_no0 = log2(A12Srm_SobAM2_CrePos_no0[,-ncol(A12Srm_SobAM2_CrePos_no0)]) 
log_A12Srm_SobAM2_CrePos_no0 <- cbind(log_A12Srm_SobAM2_CrePos_no0, 'CRE.and.Treatment' = A12Srm_SobAM2_CrePos_no0$CRE.and.Treatment)

# Fix column names & remove extra rows
ttest_A12Srm_SobAM2_CrePos_no0 <- transpose(log_A12Srm_SobAM2_CrePos_no0, keep.names = 'Lipid species')
ttest_A12Srm_SobAM2_CrePos_no0 <- ttest_A12Srm_SobAM2_CrePos_no0[-nrow(ttest_A12Srm_SobAM2_CrePos_no0),]

samples <- A12Srm_SobAM2_CrePos$Sample.number
names <- c('Lipid species', samples)
colnames(ttest_A12Srm_SobAM2_CrePos_no0) = c(names)

# Get p values from t test 
ttest_A12Srm_SobAM2_CrePos_no0$"pvalue" <- unlist(lapply(1:nrow(ttest_A12Srm_SobAM2_CrePos_no0), function(i) t.test((as.numeric(ttest_A12Srm_SobAM2_CrePos_no0[i,2:9])), as.numeric(ttest_A12Srm_SobAM2_CrePos_no0[i,10:19]))[c("p.value")]))

# calculate average values & sem for each species using CRE and treatment as factor
mean.sem_A12Srm_SobAM2_CrePos_no0 <- data.table(log_A12Srm_SobAM2_CrePos_no0)[, lapply(.SD, mean.sem), by=CRE.and.Treatment]
mean.sem_A12Srm_SobAM2_CrePos_no0 <- transpose(mean.sem_A12Srm_SobAM2_CrePos_no0[,2:(ncol(mean.sem_A12Srm_SobAM2_CrePos_no0))], keep.names = "Lipid species")
# Make sure the order of columns is correct, some df will have experimental first
CRE_factor <- c("CRE+/control avg","CRE+/control sem","CRE+/SobAM2 avg","CRE+/SobAM2 sem")
names <- c('Lipid species', CRE_factor)
colnames(mean.sem_A12Srm_SobAM2_CrePos_no0) = c(names)

# add means to main df
#VP_A12Srm_SobAM2_CrePos_no0 <- cbind(ttest_A12Srm_SobAM2_CrePos_no0, mean.sem_A12Srm_SobAM2_CrePos_no0)
VP_A12Srm_SobAM2_CrePos_no0 <- merge(ttest_A12Srm_SobAM2_CrePos_no0, mean.sem_A12Srm_SobAM2_CrePos_no0, by = 'Lipid species')
# Calculate fold change
VP_A12Srm_SobAM2_CrePos_no0$"Log2FC" <- VP_A12Srm_SobAM2_CrePos_no0$'CRE+/SobAM2 avg' - VP_A12Srm_SobAM2_CrePos_no0$'CRE+/control avg'

#save df 
write_csv(VP_A12Srm_SobAM2_CrePos_no0, "Serum/A12Srm_SobAM2_CrePos_no0_Volcano_plot.csv")

## permFDP ####

VP_A12Srm_SobAM2_CrePos_no0 <- read_csv("Serum/A12Srm_SobAM2_CrePos_no0_Volcano_plot.csv")

pVals = VP_A12Srm_SobAM2_CrePos_no0$pvalue
threshold = 0.05
myDesign = c(1,1,1,1,1,1,1,1,2,2,2,2,2,2,2,2,2,2)
intOnly = VP_A12Srm_SobAM2_CrePos_no0 %>% select(-c("Lipid species","pvalue","CRE+/control avg","CRE+/control sem","CRE+/SobAM2 avg","CRE+/SobAM2 sem","Log2FC"))
intOnly <- mutate_all(intOnly, function(x) as.numeric(as.character(x))) 
corrThreshold_500 = permFDP::permFDP.adjust.threshold(pVals, threshold, myDesign, intOnly, 500)
corrThreshold_1000 = permFDP::permFDP.adjust.threshold(pVals, threshold, myDesign, intOnly, 1000)
corrThreshold_3000 = permFDP::permFDP.adjust.threshold(pVals, threshold, myDesign, intOnly, 3000)
corrThreshold_10000 = permFDP::permFDP.adjust.threshold(pVals, threshold, myDesign, intOnly, 10000)

print(corrThreshold_3000, digits = 20)
FDPthreshold[4,2] <- corrThreshold_3000
#corrThreshold_3000 = 0.04887480829396512183

## EnhancedVolcano ####
pdf("Serum/VP_A12Srm_SobAM2_CrePos_no0.pdf", width = 11.69, height = 8.27)

# Lipid species, p values, log2 FCs
toptable <- VP_A12Srm_SobAM2_CrePos_no0[, c(1,ncol(VP_A12Srm_SobAM2_CrePos_no0)-5,ncol(VP_A12Srm_SobAM2_CrePos_no0))]
lab = toptable$"Lipid species"
x = "Log2FC"
y = "pvalue"
pCutoff = corrThreshold_3000
FCcutoff = 0.5
title = "PLP serum samples w Cre+ SobAM2"
subtitle = bquote(italic('Week 12 with no 0 values, no CE, 3000 iterations, 18 mice'))

keyvals.colour <- ifelse(
  toptable$Log2FC > 0.5 & toptable$pvalue < pCutoff, '#FF00FF',
  ifelse(toptable$Log2FC <(-0.5) & toptable$pvalue < pCutoff, '#00FFFF',
         '#808080'))
# Gray - #A8B6CC - #DAECCF
# Magenta - #F05039 - #F32A50
# Cyan - #0090FF - #0882E0

keyvals.colour[is.na(keyvals.colour)] <- '#808080'
names(keyvals.colour)[keyvals.colour == '#FF00FF'] <- 'High FC & sig'
names(keyvals.colour)[keyvals.colour == '#808080'] <- 'NS'
names(keyvals.colour)[keyvals.colour == '#00FFFF'] <- 'Low FC & sig'

EnhancedVolcano(toptable, lab, x, y, selectLab = rownames(toptable)[which(names(keyvals.colour) %in% c('High', 'Low'))],
                xlab = bquote(~Log[2] ~ "Fold Change"),
                ylab = bquote(~-Log[10] ~ italic(p) ~ Value), colCustom = keyvals.colour,
                title = title, subtitle = subtitle, pCutoff = pCutoff, pCutoffCol = y, FCcutoff = FCcutoff,labSize = 5,
                drawConnectors = FALSE
)
dev.off()

# PLP W12 Serum SobAM2 CreNeg ####

A12Srm_SobAM2_CreNeg <- A6_18_Srm %>% filter(grepl('wk 12', ...1)) %>% filter(grepl('-', CRE.and.Treatment))
A12Srm_SobAM2_CreNeg_no0 <- A12Srm_SobAM2_CreNeg %>% select(-"Sample.number", -"...1")

# For no 0 df, drop PC(36:0), PC(42:11), PC(44:12), PE(40:5), and PI(40:5)
# https://stackoverflow.com/questions/72096913/remove-columns-that-have-one-zero-value
print(select(A12Srm_SobAM2_CreNeg_no0, where(~any(. == 0))))
A12Srm_SobAM2_CreNeg_no0 <- Filter(function(x) !any(x %in% 0), A12Srm_SobAM2_CreNeg_no0) 
log_A12Srm_SobAM2_CreNeg_no0 = log2(A12Srm_SobAM2_CreNeg_no0[,-ncol(A12Srm_SobAM2_CreNeg_no0)]) 
log_A12Srm_SobAM2_CreNeg_no0 <- cbind(log_A12Srm_SobAM2_CreNeg_no0, 'CRE.and.Treatment' = A12Srm_SobAM2_CreNeg$CRE.and.Treatment)

# Fix column names & remove extra rows
ttest_A12Srm_SobAM2_CreNeg_no0 <- transpose(log_A12Srm_SobAM2_CreNeg_no0, keep.names = 'Lipid species')
ttest_A12Srm_SobAM2_CreNeg_no0 <- ttest_A12Srm_SobAM2_CreNeg_no0[-nrow(ttest_A12Srm_SobAM2_CreNeg_no0),]

samples <- A12Srm_SobAM2_CreNeg$Sample.number
names <- c('Lipid species', samples)
colnames(ttest_A12Srm_SobAM2_CreNeg_no0) = c(names)

# Get p values from t test 
ttest_A12Srm_SobAM2_CreNeg_no0$"pvalue" <- unlist(lapply(1:nrow(ttest_A12Srm_SobAM2_CreNeg_no0), function(i) t.test((as.numeric(ttest_A12Srm_SobAM2_CreNeg_no0[i,2:6])), as.numeric(ttest_A12Srm_SobAM2_CreNeg_no0[i,7:12]))[c("p.value")]))

# calculate average values & sem for each species using CRE and treatment as factor
mean.sem_A12Srm_SobAM2_CreNeg_no0 <- data.table(log_A12Srm_SobAM2_CreNeg_no0)[, lapply(.SD, mean.sem), by=CRE.and.Treatment]
mean.sem_A12Srm_SobAM2_CreNeg_no0 <- transpose(mean.sem_A12Srm_SobAM2_CreNeg_no0[,2:(ncol(mean.sem_A12Srm_SobAM2_CreNeg_no0))], keep.names = "Lipid species")
# Make sure the order of columns is correct, some df will have experimental first
CRE_factor <- c("CRE-/control avg","CRE-/control sem","CRE-/SobAM2 avg","CRE-/SobAM2 sem")
names <- c('Lipid species', CRE_factor)
colnames(mean.sem_A12Srm_SobAM2_CreNeg_no0) = c(names)

# add means to main df
#VP_A12Srm_SobAM2_CreNeg_no0 <- cbind(ttest_A12Srm_SobAM2_CreNeg_no0, mean.sem_A12Srm_SobAM2_CreNeg_no0)
VP_A12Srm_SobAM2_CreNeg_no0 <- merge(ttest_A12Srm_SobAM2_CreNeg_no0, mean.sem_A12Srm_SobAM2_CreNeg_no0, by = 'Lipid species')
# Calculate fold change
VP_A12Srm_SobAM2_CreNeg_no0$"Log2FC" <- VP_A12Srm_SobAM2_CreNeg_no0$'CRE-/SobAM2 avg' - VP_A12Srm_SobAM2_CreNeg_no0$'CRE-/control avg'

#save df 
write_csv(VP_A12Srm_SobAM2_CreNeg_no0, "Serum/A12Srm_SobAM2_CreNeg_no0_Volcano_plot.csv")

## permFDP ####

VP_A12Srm_SobAM2_CreNeg_no0 <- read_csv("Serum/A12Srm_SobAM2_CreNeg_no0_Volcano_plot.csv")

pVals = VP_A12Srm_SobAM2_CreNeg_no0$pvalue
threshold = 0.05
myDesign = c(1,1,1,1,1,2,2,2,2,2,2)
intOnly = VP_A12Srm_SobAM2_CreNeg_no0 %>% select(-c("Lipid species","pvalue","CRE-/control avg","CRE-/control sem","CRE-/SobAM2 avg","CRE-/SobAM2 sem","Log2FC"))
intOnly <- mutate_all(intOnly, function(x) as.numeric(as.character(x))) 
corrThreshold_500 = permFDP::permFDP.adjust.threshold(pVals, threshold, myDesign, intOnly, 500)
corrThreshold_1000 = permFDP::permFDP.adjust.threshold(pVals, threshold, myDesign, intOnly, 1000)
corrThreshold_3000 = permFDP::permFDP.adjust.threshold(pVals, threshold, myDesign, intOnly, 3000)
corrThreshold_10000 = permFDP::permFDP.adjust.threshold(pVals, threshold, myDesign, intOnly, 10000)

print(corrThreshold_3000, digits = 20)
FDPthreshold[5,2] <- corrThreshold_3000
#corrThreshold_3000 = 0.026423699995021976417

## EnhancedVolcano ####
pdf("Serum/VP_A12Srm_SobAM2_CreNeg_no0.pdf", width = 11.69, height = 8.27)

# Lipid species, p values, log2 FCs
toptable <- VP_A12Srm_SobAM2_CreNeg_no0[, c(1,ncol(VP_A12Srm_SobAM2_CreNeg_no0)-5,ncol(VP_A12Srm_SobAM2_CreNeg_no0))]
lab = toptable$"Lipid species"
x = "Log2FC"
y = "pvalue"
pCutoff = corrThreshold_3000
FCcutoff = 0.5
title = "PLP serum samples w Cre- SobAM2"
subtitle = bquote(italic('Week 12 with no 0 values, no CE, 3000 iterations, 11 mice'))

keyvals.colour <- ifelse(
  toptable$Log2FC > 0.5 & toptable$pvalue < pCutoff, '#FF00FF',
  ifelse(toptable$Log2FC <(-0.5) & toptable$pvalue < pCutoff, '#00FFFF',
         '#808080'))
# Gray - #A8B6CC - #DAECCF
# Magenta - #F05039 - #F32A50
# Cyan - #0090FF - #0882E0

keyvals.colour[is.na(keyvals.colour)] <- '#808080'
names(keyvals.colour)[keyvals.colour == '#FF00FF'] <- 'High FC & sig'
names(keyvals.colour)[keyvals.colour == '#808080'] <- 'NS'
names(keyvals.colour)[keyvals.colour == '#00FFFF'] <- 'Low FC & sig'

EnhancedVolcano(toptable, lab, x, y, selectLab = rownames(toptable)[which(names(keyvals.colour) %in% c('High', 'Low'))],
                xlab = bquote(~Log[2] ~ "Fold Change"),
                ylab = bquote(~-Log[10] ~ italic(p) ~ Value), colCustom = keyvals.colour,
                title = title, subtitle = subtitle, pCutoff = pCutoff, pCutoffCol = y, FCcutoff = FCcutoff,labSize = 5,
                drawConnectors = FALSE
)
dev.off()

# PLP W18 Serum SobAM2 CrePos ####

A18Srm_SobAM2_CrePos <- A6_18_Srm %>% filter(grepl('wk 18', ...1)) %>% filter(!grepl('-', CRE.and.Treatment))
A18Srm_SobAM2_CrePos_no0 <- A18Srm_SobAM2_CrePos %>% select(-"Sample.number", -"...1")

# For no 0 df, drop PC(36:0), PC(44:12), PE(40:5), and ePE(36:5)
# https://stackoverflow.com/questions/72096913/remove-columns-that-have-one-zero-value
print(select(A18Srm_SobAM2_CrePos_no0, where(~any(. == 0))))
A18Srm_SobAM2_CrePos_no0 <- Filter(function(x) !any(x %in% 0), A18Srm_SobAM2_CrePos_no0) 
log_A18Srm_SobAM2_CrePos_no0 = log2(A18Srm_SobAM2_CrePos_no0[,-ncol(A18Srm_SobAM2_CrePos_no0)]) 
log_A18Srm_SobAM2_CrePos_no0 <- cbind(log_A18Srm_SobAM2_CrePos_no0, 'CRE.and.Treatment' = A18Srm_SobAM2_CrePos_no0$CRE.and.Treatment)

# Fix column names & remove extra rows
ttest_A18Srm_SobAM2_CrePos_no0 <- transpose(log_A18Srm_SobAM2_CrePos_no0, keep.names = 'Lipid species')
ttest_A18Srm_SobAM2_CrePos_no0 <- ttest_A18Srm_SobAM2_CrePos_no0[-nrow(ttest_A18Srm_SobAM2_CrePos_no0),]

samples <- A18Srm_SobAM2_CrePos$Sample.number
names <- c('Lipid species', samples)
colnames(ttest_A18Srm_SobAM2_CrePos_no0) = c(names)

# Get p values from t test 
ttest_A18Srm_SobAM2_CrePos_no0$"pvalue" <- unlist(lapply(1:nrow(ttest_A18Srm_SobAM2_CrePos_no0), function(i) t.test((as.numeric(ttest_A18Srm_SobAM2_CrePos_no0[i,2:9])), as.numeric(ttest_A18Srm_SobAM2_CrePos_no0[i,10:18]))[c("p.value")]))

# calculate average values & sem for each species using CRE and treatment as factor
mean.sem_A18Srm_SobAM2_CrePos_no0 <- data.table(log_A18Srm_SobAM2_CrePos_no0)[, lapply(.SD, mean.sem), by=CRE.and.Treatment]
mean.sem_A18Srm_SobAM2_CrePos_no0 <- transpose(mean.sem_A18Srm_SobAM2_CrePos_no0[,2:(ncol(mean.sem_A18Srm_SobAM2_CrePos_no0))], keep.names = "Lipid species")
# Make sure the order of columns is correct, some df will have experimental first
CRE_factor <- c("CRE+/control avg","CRE+/control sem","CRE+/SobAM2 avg","CRE+/SobAM2 sem")
names <- c('Lipid species', CRE_factor)
colnames(mean.sem_A18Srm_SobAM2_CrePos_no0) = c(names)

# add means to main df
#VP_A18Srm_SobAM2_CrePos_no0 <- cbind(ttest_A18Srm_SobAM2_CrePos_no0, mean.sem_A18Srm_SobAM2_CrePos_no0)
VP_A18Srm_SobAM2_CrePos_no0 <- merge(ttest_A18Srm_SobAM2_CrePos_no0, mean.sem_A18Srm_SobAM2_CrePos_no0, by = 'Lipid species')
# Calculate fold change
VP_A18Srm_SobAM2_CrePos_no0$"Log2FC" <- VP_A18Srm_SobAM2_CrePos_no0$'CRE+/SobAM2 avg' - VP_A18Srm_SobAM2_CrePos_no0$'CRE+/control avg'

#save df 
write_csv(VP_A18Srm_SobAM2_CrePos_no0, "Serum/A18Srm_SobAM2_CrePos_no0_Volcano_plot.csv")

## permFDP ####

VP_A18Srm_SobAM2_CrePos_no0 <- read_csv("Serum/A18Srm_SobAM2_CrePos_no0_Volcano_plot.csv")

pVals = VP_A18Srm_SobAM2_CrePos_no0$pvalue
threshold = 0.05
myDesign = c(1,1,1,1,1,1,1,1,2,2,2,2,2,2,2,2,2)
intOnly = VP_A18Srm_SobAM2_CrePos_no0 %>% select(-c("Lipid species","pvalue","CRE+/control avg","CRE+/control sem","CRE+/SobAM2 avg","CRE+/SobAM2 sem","Log2FC"))
intOnly <- mutate_all(intOnly, function(x) as.numeric(as.character(x))) 
corrThreshold_500 = permFDP::permFDP.adjust.threshold(pVals, threshold, myDesign, intOnly, 500)
corrThreshold_1000 = permFDP::permFDP.adjust.threshold(pVals, threshold, myDesign, intOnly, 1000)
corrThreshold_3000 = permFDP::permFDP.adjust.threshold(pVals, threshold, myDesign, intOnly, 3000)
corrThreshold_10000 = permFDP::permFDP.adjust.threshold(pVals, threshold, myDesign, intOnly, 10000)

print(corrThreshold_3000, digits = 20)
FDPthreshold[6,2] <- corrThreshold_3000
#corrThreshold_3000 = 0.032390326926054612

## EnhancedVolcano ####
pdf("Serum/VP_A18Srm_SobAM2_CrePos_no0.pdf", width = 11.69, height = 8.27)

# Lipid species, p values, log2 FCs
toptable <- VP_A18Srm_SobAM2_CrePos_no0[, c(1,ncol(VP_A18Srm_SobAM2_CrePos_no0)-5,ncol(VP_A18Srm_SobAM2_CrePos_no0))]
lab = toptable$"Lipid species"
x = "Log2FC"
y = "pvalue"
pCutoff = corrThreshold_3000
FCcutoff = 0.5
title = "PLP serum samples w Cre+ SobAM2"
subtitle = bquote(italic('Week 18 with no 0 values, no CE, 3000 iterations, 17 mice'))

keyvals.colour <- ifelse(
  toptable$Log2FC > 0.5 & toptable$pvalue < pCutoff, '#FF00FF',
  ifelse(toptable$Log2FC <(-0.5) & toptable$pvalue < pCutoff, '#00FFFF',
         '#808080'))
# Gray - #A8B6CC - #DAECCF
# Magenta - #F05039 - #F32A50
# Cyan - #0090FF - #0882E0

keyvals.colour[is.na(keyvals.colour)] <- '#808080'
names(keyvals.colour)[keyvals.colour == '#FF00FF'] <- 'High FC & sig'
names(keyvals.colour)[keyvals.colour == '#808080'] <- 'NS'
names(keyvals.colour)[keyvals.colour == '#00FFFF'] <- 'Low FC & sig'

EnhancedVolcano(toptable, lab, x, y, selectLab = rownames(toptable)[which(names(keyvals.colour) %in% c('High', 'Low'))],
                xlab = bquote(~Log[2] ~ "Fold Change"),
                ylab = bquote(~-Log[10] ~ italic(p) ~ Value), colCustom = keyvals.colour,
                title = title, subtitle = subtitle, pCutoff = pCutoff, pCutoffCol = y, FCcutoff = FCcutoff,labSize = 5,
                drawConnectors = FALSE
)
dev.off()

# PLP W18 Serum SobAM2 CreNeg ####

A18Srm_SobAM2_CreNeg <- A6_18_Srm %>% filter(grepl('wk 18', ...1)) %>% filter(grepl('-', CRE.and.Treatment))
A18Srm_SobAM2_CreNeg_no0 <- A18Srm_SobAM2_CreNeg %>% select(-"Sample.number", -"...1")

# For no 0 df, drop PC(36:0), PC(40:2), PC(44:12), and ePE(36:4)
# https://stackoverflow.com/questions/72096913/remove-columns-that-have-one-zero-value
print(select(A18Srm_SobAM2_CreNeg_no0, where(~any(. == 0))))
A18Srm_SobAM2_CreNeg_no0 <- Filter(function(x) !any(x %in% 0), A18Srm_SobAM2_CreNeg_no0) 
log_A18Srm_SobAM2_CreNeg_no0 = log2(A18Srm_SobAM2_CreNeg_no0[,-ncol(A18Srm_SobAM2_CreNeg_no0)]) 
log_A18Srm_SobAM2_CreNeg_no0 <- cbind(log_A18Srm_SobAM2_CreNeg_no0, 'CRE.and.Treatment' = A18Srm_SobAM2_CreNeg$CRE.and.Treatment)

# Fix column names & remove extra rows
ttest_A18Srm_SobAM2_CreNeg_no0 <- transpose(log_A18Srm_SobAM2_CreNeg_no0, keep.names = 'Lipid species')
ttest_A18Srm_SobAM2_CreNeg_no0 <- ttest_A18Srm_SobAM2_CreNeg_no0[-nrow(ttest_A18Srm_SobAM2_CreNeg_no0),]

samples <- A18Srm_SobAM2_CreNeg$Sample.number
names <- c('Lipid species', samples)
colnames(ttest_A18Srm_SobAM2_CreNeg_no0) = c(names)

# Get p values from t test 
ttest_A18Srm_SobAM2_CreNeg_no0$"pvalue" <- unlist(lapply(1:nrow(ttest_A18Srm_SobAM2_CreNeg_no0), function(i) t.test((as.numeric(ttest_A18Srm_SobAM2_CreNeg_no0[i,2:7])), as.numeric(ttest_A18Srm_SobAM2_CreNeg_no0[i,8:11]))[c("p.value")]))

# calculate average values & sem for each species using CRE and treatment as factor
mean.sem_A18Srm_SobAM2_CreNeg_no0 <- data.table(log_A18Srm_SobAM2_CreNeg_no0)[, lapply(.SD, mean.sem), by=CRE.and.Treatment]
mean.sem_A18Srm_SobAM2_CreNeg_no0 <- transpose(mean.sem_A18Srm_SobAM2_CreNeg_no0[,2:(ncol(mean.sem_A18Srm_SobAM2_CreNeg_no0))], keep.names = "Lipid species")
# Make sure the order of columns is correct, some df will have experimental first
CRE_factor <- c("CRE-/control avg","CRE-/control sem","CRE-/SobAM2 avg","CRE-/SobAM2 sem")
names <- c('Lipid species', CRE_factor)
colnames(mean.sem_A18Srm_SobAM2_CreNeg_no0) = c(names)

# add means to main df
#VP_A18Srm_SobAM2_CreNeg_no0 <- cbind(ttest_A18Srm_SobAM2_CreNeg_no0, mean.sem_A18Srm_SobAM2_CreNeg_no0)
VP_A18Srm_SobAM2_CreNeg_no0 <- merge(ttest_A18Srm_SobAM2_CreNeg_no0, mean.sem_A18Srm_SobAM2_CreNeg_no0, by = 'Lipid species')
# Calculate fold change
VP_A18Srm_SobAM2_CreNeg_no0$"Log2FC" <- VP_A18Srm_SobAM2_CreNeg_no0$'CRE-/SobAM2 avg' - VP_A18Srm_SobAM2_CreNeg_no0$'CRE-/control avg'

#save df 
write_csv(VP_A18Srm_SobAM2_CreNeg_no0, "Serum/A18Srm_SobAM2_CreNeg_no0_Volcano_plot.csv")

## permFDP ####

VP_A18Srm_SobAM2_CreNeg_no0 <- read_csv("Serum/A18Srm_SobAM2_CreNeg_no0_Volcano_plot.csv")

pVals = VP_A18Srm_SobAM2_CreNeg_no0$pvalue
threshold = 0.05
myDesign = c(1,1,1,1,1,1,2,2,2,2)
intOnly = VP_A18Srm_SobAM2_CreNeg_no0 %>% select(-c("Lipid species","pvalue","CRE-/control avg","CRE-/control sem","CRE-/SobAM2 avg","CRE-/SobAM2 sem","Log2FC"))
intOnly <- mutate_all(intOnly, function(x) as.numeric(as.character(x))) 
corrThreshold_500 = permFDP::permFDP.adjust.threshold(pVals, threshold, myDesign, intOnly, 500)
corrThreshold_1000 = permFDP::permFDP.adjust.threshold(pVals, threshold, myDesign, intOnly, 1000)
corrThreshold_3000 = permFDP::permFDP.adjust.threshold(pVals, threshold, myDesign, intOnly, 3000)
corrThreshold_10000 = permFDP::permFDP.adjust.threshold(pVals, threshold, myDesign, intOnly, 10000)

print(corrThreshold_3000, digits = 20)
FDPthreshold[7,2] <- corrThreshold_3000
#corrThreshold_3000 = 0.0090122461590888235106

## EnhancedVolcano ####
pdf("Serum/VP_A18Srm_SobAM2_CreNeg_no0.pdf", width = 11.69, height = 8.27)

# Lipid species, p values, log2 FCs
toptable <- VP_A18Srm_SobAM2_CreNeg_no0[, c(1,ncol(VP_A18Srm_SobAM2_CreNeg_no0)-5,ncol(VP_A18Srm_SobAM2_CreNeg_no0))]
lab = toptable$"Lipid species"
x = "Log2FC"
y = "pvalue"
pCutoff = corrThreshold_3000
FCcutoff = 0.5
title = "PLP serum samples w Cre- SobAM2"
subtitle = bquote(italic('Week 18 with no 0 values, no CE, 3000 iterations, 10 mice'))

keyvals.colour <- ifelse(
  toptable$Log2FC > 0.5 & toptable$pvalue < pCutoff, '#FF00FF',
  ifelse(toptable$Log2FC <(-0.5) & toptable$pvalue < pCutoff, '#00FFFF',
         '#808080'))
# Gray - #A8B6CC - #DAECCF
# Magenta - #F05039 - #F32A50
# Cyan - #0090FF - #0882E0

keyvals.colour[is.na(keyvals.colour)] <- '#808080'
names(keyvals.colour)[keyvals.colour == '#FF00FF'] <- 'High FC & sig'
names(keyvals.colour)[keyvals.colour == '#808080'] <- 'NS'
names(keyvals.colour)[keyvals.colour == '#00FFFF'] <- 'Low FC & sig'

EnhancedVolcano(toptable, lab, x, y, selectLab = rownames(toptable)[which(names(keyvals.colour) %in% c('High', 'Low'))],
                xlab = bquote(~Log[2] ~ "Fold Change"),
                ylab = bquote(~-Log[10] ~ italic(p) ~ Value), colCustom = keyvals.colour,
                title = title, subtitle = subtitle, pCutoff = pCutoff, pCutoffCol = y, FCcutoff = FCcutoff,labSize = 5,
                drawConnectors = FALSE
)
dev.off()

write_csv(FDPthreshold,"Serum/FDP_Threshold_Srm.csv")
```
