## Supplemental Codes for "Lipidomic analysis reveals differences in the extent of remyelination in the brain and spinal cord": VP_table_serum.R

```
# This code will generate tables from volcano plot results
# Specifically for serum

# Library ####

library(openxlsx)
library(readr)
library(dplyr)

# Functions ####

merge_sheets <- function(df, keyvals, control_num, experiment_num){
  new_df <- select(df, all_of(keyvals))
  new_df$" -log10_pvalue" <- -log10(new_df$"pvalue")
  new_df$"FC" <- 2^new_df$"Log2FC"
  colnames(new_df)[2] <- paste(colnames(new_df[2])," n=",control_num, sep = "")
  colnames(new_df)[4] <- paste(colnames(new_df[4])," n=",experiment_num, sep = "")
  return(new_df)
}

lipid_levels <- function(df, threshold_ref, timepoint, filter_var){
  new_df <- select(df, all_of(filter_var))
  new_df <- filter(new_df, Log2FC <(-0.5) | Log2FC >0.5 ,
                   ` -log10_pvalue` > -log10(threshold_ref[timepoint]))
  new_df$Expression <- ifelse(new_df$Log2FC > 0.5, 'Increased', 'Decreased')
  return(new_df)
}

lipid_level_count <- function(df_list, count_df) {
  week = 1
  for (df in df_list){
    up = 0
    down = 0
    for (r in 1:nrow(df_list[[week]])){
      if (nrow(df_list[[week]]) == 0) {
        count_df[week,2] = 0
        count_df[week,3] = 0
      }
      else {
        if (df_list[[week]][r,4] == "Increased"){
          count_df[week,2] = up + 1
          up = up + 1
        }
        else if (df_list[[week]][r,4] == "Decreased"){
          count_df[week,3] = down + 1
          down = down + 1
        }
      }
    }
    count_df[is.na(count_df)] <- 0
    week = week + 1
  }
  return(count_df)
}

# Load VP data ####
A6Srm_PLP <- read_csv("Serum/Tables_VP/A6Srm_no0_Volcano_plot.csv")
A12Srm_PLP <- read_csv("Serum/Tables_VP/A12Srm_no0_Volcano_plot.csv")
A18Srm_PLP <- read_csv("Serum/Tables_VP/A18Srm_no0_Volcano_plot.csv")
A12Srm_SobAM2_CrePos_PLP <- read_csv("Serum/Tables_VP/A12Srm_SobAM2_CrePos_no0_Volcano_plot.csv")
A12Srm_SobAM2_CreNeg_PLP <- read_csv("Serum/Tables_VP/A12Srm_SobAM2_CreNeg_no0_Volcano_plot.csv")
A18Srm_SobAM2_CrePos_PLP <- read_csv("Serum/Tables_VP/A18Srm_SobAM2_CrePos_no0_Volcano_plot.csv")
A18Srm_SobAM2_CreNeg_PLP <- read_csv("Serum/Tables_VP/A18Srm_SobAM2_CreNeg_no0_Volcano_plot.csv")

## Keyvalue df ####

keyvals <- c("Lipid species","CRE-/control avg","CRE-/control sem","CRE+/control avg","CRE+/control sem","Log2FC","pvalue")
keyvals_SobAM2_CrePos <- c("Lipid species","CRE+/control avg","CRE+/control sem","CRE+/SobAM2 avg","CRE+/SobAM2 sem","Log2FC","pvalue")
keyvals_SobAM2_CreNeg <- c("Lipid species","CRE-/control avg","CRE-/control sem","CRE-/SobAM2 avg","CRE-/SobAM2 sem","Log2FC","pvalue")

A6Srm_keyvals <- merge_sheets(A6Srm_PLP, keyvals, 3, 8)
A12Srm_keyvals <- merge_sheets(A12Srm_PLP, keyvals, 5, 8)
A18Srm_keyvals <- merge_sheets(A18Srm_PLP, keyvals, 6, 8)

A12Srm_SobAM2_CrePos_keyvals <- merge_sheets(A12Srm_SobAM2_CrePos_PLP, keyvals_SobAM2_CrePos, 8, 10)
A12Srm_SobAM2_CreNeg_keyvals <- merge_sheets(A12Srm_SobAM2_CreNeg_PLP, keyvals_SobAM2_CreNeg, 5, 6)
A18Srm_SobAM2_CrePos_keyvals <- merge_sheets(A18Srm_SobAM2_CrePos_PLP, keyvals_SobAM2_CrePos, 8, 9)
A18Srm_SobAM2_CreNeg_keyvals <- merge_sheets(A18Srm_SobAM2_CreNeg_PLP, keyvals_SobAM2_CreNeg, 6, 4)

require(openxlsx)

list_serum <- list("A6Srm_n11" = A6Srm_keyvals, "A12Srm_n13" = A12Srm_keyvals,
                   "A18Srm_n14" = A18Srm_keyvals, "A12Srm_SobAM2_CrePos_n18" = A12Srm_SobAM2_CrePos_keyvals,
                   "A12Srm_SobAM2_CreNeg_n11" = A12Srm_SobAM2_CreNeg_keyvals, 
                   "A18Srm_SobAM2_CrePos_n17" = A18Srm_SobAM2_CrePos_keyvals,
                   "A18Srm_SobAM2_CreNeg_n10" = A18Srm_SobAM2_CreNeg_keyvals)
write.xlsx(list_serum, file = "Serum/Serum_PLP_keyvals.xlsx")

## lipid regulation - cleaning ####

FDPthreshold <- read_csv("Serum/Tables_VP/FDP_Threshold_Srm.csv")
filter <- c("Lipid species","Log2FC"," -log10_pvalue")

A6Srm_exp <- lipid_levels(A6Srm_keyvals, FDPthreshold$Threshold, 1, filter)
A12Srm_exp <- lipid_levels(A12Srm_keyvals, FDPthreshold$Threshold, 2, filter)
A18Srm_exp <- lipid_levels(A18Srm_keyvals, FDPthreshold$Threshold, 3, filter)
A12Srm_SobAM2_CrePos_exp <- lipid_levels(A12Srm_SobAM2_CrePos_keyvals, FDPthreshold$Threshold, 4, filter)
A12Srm_SobAM2_CreNeg_exp <- lipid_levels(A12Srm_SobAM2_CreNeg_keyvals, FDPthreshold$Threshold, 5, filter)
A18Srm_SobAM2_CrePos_exp <- lipid_levels(A18Srm_SobAM2_CrePos_keyvals, FDPthreshold$Threshold, 6, filter)
A18Srm_SobAM2_CreNeg_exp <- lipid_levels(A18Srm_SobAM2_CreNeg_keyvals, FDPthreshold$Threshold, 7, filter)

require(openxlsx)
list_serum2 <- list("A6B_n11" = A6Srm_exp, "A12B_n13" = A12Srm_exp, "A18B_n14" = A18Srm_exp,
                   "A12Srm_SobAM2_CrePos_n18" = A12Srm_SobAM2_CrePos_exp, 
                   "A12Srm_SobAM2_CreNeg_n11" = A12Srm_SobAM2_CreNeg_exp,
                   "A18Srm_SobAM2_CrePos_n17" = A18Srm_SobAM2_CrePos_exp, 
                   "A18Srm_SobAM2_CreNeg_n10" = A18Srm_SobAM2_CreNeg_exp
                   )
write.xlsx(list_serum2, file = "Serum/Serum_PLP_regulation.xlsx")

## lipid level count ####

columns <- c("Timepoint", "Increased", "Decreased")
rows <- c("A6Srm","A12Srm","A18Srm","A12Srm_SobAM2_CrePos","A12Srm_SobAM2_CreNeg","A18Srm_SobAM2_CrePos","A18Srm_SobAM2_CreNeg")
count = data.frame(matrix(nrow = length(rows), ncol = length(columns))) 
colnames(count) = columns
count$Timepoint <- rows

PLP_list <- list("A6Srm" = A6Srm_exp,"A12Srm" = A12Srm_exp,"A18Srm" = A18Srm_exp,
                 "A12Srm_SobAM2_CrePos" = A12Srm_SobAM2_CrePos_exp, "A12Srm_SobAM2_CreNeg" = A12Srm_SobAM2_CreNeg_exp,
                 "A18Srm_SobAM2_CrePos" = A18Srm_SobAM2_CrePos_exp, "A18Srm_SobAM2_CreNeg" = A18Srm_SobAM2_CreNeg_exp
)

lipid_count <- lipid_level_count(PLP_list, count)

write_csv(lipid_count, "Serum/VP_expression_count_Srm.csv")
```
