## Supplemental Codes for "Lipidomic analysis reveals differences in the extent of remyelination in the brain and spinal cord": Volcano_Plot_Tables.R

```
# VP tables and lipid levels

# Library ####

library(openxlsx)
library(readr)
library(dplyr)

# Functions ####

merge_sheets <- function(df, keyvals, control_num, experiment_num){
  new_df <- select(df, all_of(keyvals))
  new_df$" -log10_pvalue" <- -log10(new_df$"pvalue")
  new_df$"FC" <- 2^new_df$"Log2FC"
  colnames(new_df)[2] <- paste(colnames(new_df[2])," n=",control_num, sep = "")
  colnames(new_df)[4] <- paste(colnames(new_df[4])," n=",experiment_num, sep = "")
  return(new_df)
}

lipid_levels <- function(df, threshold_ref, timepoint, filter_var){
  new_df <- select(df, all_of(filter_var))
  new_df <- filter(new_df, Log2FC <(-0.5) | Log2FC >0.5 ,
                   ` -log10_pvalue` > -log10(threshold_ref[timepoint]))
  new_df$Expression <- ifelse(new_df$Log2FC > 0.5, 'Increased', 'Decreased')
  return(new_df)
}

lipid_level_count <- function(df_list, count_df) {
  week = 1
  for (df in df_list){
    up = 0
    down = 0
    for (r in 1:nrow(df_list[[week]])){
      if (nrow(df_list[[week]]) == 0) {
        count_df[week,2] = 0
        count_df[week,3] = 0
      }
      else {
        if (df_list[[week]][r,4] == "Increased"){
          count_df[week,2] = up + 1
          up = up + 1
        }
        else if (df_list[[week]][r,4] == "Decreased"){
          count_df[week,3] = down + 1
          down = down + 1
        }
      }
    }
    count_df[is.na(count_df)] <- 0
    week = week + 1
  }
  return(count_df)
}

# Load VP data ####
df_A6B_PLP <- read_csv("VP_csv/A6B_no0_Volcano_plot.csv")
df_A6S_PLP <- read_csv("VP_csv/A6S_no0_Volcano_plot.csv")
df_A12B_PLP <- read_csv("VP_csv/A12B_no0_Volcano_plot.csv")
df_A12S_PLP <- read_csv("VP_csv/A12S_no0_Volcano_plot.csv")
df_A18B_PLP <- read_csv("VP_csv/A18B_no0_Volcano_plot.csv")
df_A18S_PLP <- read_csv("VP_csv/A18S_no0_Volcano_plot.csv")
df_A24B_PLP <- read_csv("VP_csv/A24B_no0_Volcano_plot.csv")
df_A24S_PLP <- read_csv("VP_csv/A24S_no0_Volcano_plot.csv")
df_A12B_SobAM2_CrePos_PLP <- read_csv("VP_csv/A12B_SobAM2_CrePos_no0_Volcano_plot.csv")
df_A12S_SobAM2_CrePos_PLP <- read_csv("VP_csv/A12S_SobAM2_CrePos_no0_Volcano_plot.csv")
df_A12B_SobAM2_CreNeg_PLP <- read_csv("VP_csv/A12B_SobAM2_CreNeg_no0_Volcano_plot.csv")
df_A12S_SobAM2_CreNeg_PLP <- read_csv("VP_csv/A12S_SobAM2_CreNeg_no0_Volcano_plot.csv")
df_A18B_SobAM2_CrePos_PLP <- read_csv("VP_csv/A18B_SobAM2_CrePos_no0_Volcano_plot.csv")
df_A18S_SobAM2_CrePos_PLP <- read_csv("VP_csv/A18S_SobAM2_CrePos_no0_Volcano_plot.csv")
df_A18B_SobAM2_CreNeg_PLP <- read_csv("VP_csv/A18B_SobAM2_CreNeg_no0_Volcano_plot.csv")
df_A18S_SobAM2_CreNeg_PLP <- read_csv("VP_csv/A18S_SobAM2_CreNeg_no0_Volcano_plot.csv")

## Keyvalue df ####

keyvals <- c("Lipid species","CRE-/control avg","CRE-/control sem","CRE+/control avg","CRE+/control sem","Log2FC","pvalue")
keyvals_CrePos_SobAM2 <- c("Lipid species","CRE+/control avg","CRE+/control sem","CRE+/SobAM2 avg","CRE+/SobAM2 sem","Log2FC","pvalue")
keyvals_CreNeg_SobAM2 <- c("Lipid species","CRE-/control avg","CRE-/control sem","CRE-/SobAM2 avg","CRE-/SobAM2 sem","Log2FC","pvalue")

A6B_keyvals <- merge_sheets(df_A6B_PLP, keyvals, 3, 8)
A6S_keyvals <- merge_sheets(df_A6S_PLP, keyvals, 3, 8)
A12B_keyvals <- merge_sheets(df_A12B_PLP, keyvals, 5, 8)
A12S_keyvals <- merge_sheets(df_A12S_PLP, keyvals, 5, 8)
A18B_keyvals <- merge_sheets(df_A18B_PLP, keyvals, 6, 8)
A18S_keyvals <- merge_sheets(df_A18S_PLP, keyvals, 6, 9)
A24B_keyvals <- merge_sheets(df_A24B_PLP, keyvals, 4, 7)
A24S_keyvals <- merge_sheets(df_A24S_PLP, keyvals, 3, 7)

A12B_SobAM2_CrePos_keyvals <- merge_sheets(df_A12B_SobAM2_CrePos_PLP, keyvals_CrePos_SobAM2, 8, 10)
A12S_SobAM2_CrePos_keyvals <- merge_sheets(df_A12S_SobAM2_CrePos_PLP, keyvals_CrePos_SobAM2, 8, 10)
A12B_SobAM2_CreNeg_keyvals <- merge_sheets(df_A12B_SobAM2_CreNeg_PLP, keyvals_CreNeg_SobAM2, 5, 6)
A12S_SobAM2_CreNeg_keyvals <- merge_sheets(df_A12S_SobAM2_CreNeg_PLP, keyvals_CreNeg_SobAM2, 5, 6)
A18B_SobAM2_CrePos_keyvals <- merge_sheets(df_A18B_SobAM2_CrePos_PLP, keyvals_CrePos_SobAM2, 8, 9)
A18S_SobAM2_CrePos_keyvals <- merge_sheets(df_A18S_SobAM2_CrePos_PLP, keyvals_CrePos_SobAM2, 9, 9)
A18B_SobAM2_CreNeg_keyvals <- merge_sheets(df_A18B_SobAM2_CreNeg_PLP, keyvals_CreNeg_SobAM2, 6, 4)
A18S_SobAM2_CreNeg_keyvals <- merge_sheets(df_A18S_SobAM2_CreNeg_PLP, keyvals_CreNeg_SobAM2, 6, 4)

require(openxlsx)
list_brain <- list("A6B_n11" = A6B_keyvals, "A12B_n13" = A12B_keyvals,
                   "A18B_n14" = A18B_keyvals, "A24B_n11" = A24B_keyvals,
                   "A12B_SobAM2_CrePos_n18" = A12B_SobAM2_CrePos_keyvals, "A12B_SobAM2_CreNeg_n11" = A12B_SobAM2_CreNeg_keyvals,
                   "A18B_SobAM2_CrePos_n17" = A18B_SobAM2_CrePos_keyvals, "A18B_SobAM2_CreNeg_n10" = A18B_SobAM2_CreNeg_keyvals)
write.xlsx(list_brain, file = "FinalVP/Brain_PLP_keyvals.xlsx")

list_spinal <- list("A6S_n11" = A6S_keyvals, "A12S_n13" = A12S_keyvals,
                    "A18S_n15" = A18S_keyvals, "A24S_n10" = A24S_keyvals,
                    "A12S_SobAM2_CrePos_n18" = A12S_SobAM2_CrePos_keyvals, "A12S_SobAM2_CreNeg_n11" = A12S_SobAM2_CreNeg_keyvals,
                    "A18S_SobAM2_CrePos_n18" = A18S_SobAM2_CrePos_keyvals, "A18S_SobAM2_CreNeg_n10" = A18S_SobAM2_CreNeg_keyvals)
write.xlsx(list_spinal, file = "FinalVP/Spinal_PLP_keyvals.xlsx")

## lipid regulation - cleaning ####

FDPthreshold <- read_csv("FDP_Threshold.csv")
filter <- c("Lipid species","Log2FC"," -log10_pvalue")

A6B_exp <- lipid_levels(A6B_keyvals, FDPthreshold$Threshold, 1, filter)
A6S_exp <- lipid_levels(A6S_keyvals, FDPthreshold$Threshold, 2, filter)
A12B_exp <- lipid_levels(A12B_keyvals, FDPthreshold$Threshold, 3, filter)
A12S_exp <- lipid_levels(A12S_keyvals, FDPthreshold$Threshold, 4, filter)
A18B_exp <- lipid_levels(A18B_keyvals, FDPthreshold$Threshold, 5, filter)
A18S_exp <- lipid_levels(A18S_keyvals, FDPthreshold$Threshold, 6, filter)
A24B_exp <- lipid_levels(A24B_keyvals, FDPthreshold$Threshold, 7, filter)
A24S_exp <- lipid_levels(A24S_keyvals, FDPthreshold$Threshold, 8, filter)
A12B_SobAM2_CrePos_exp <- lipid_levels(A12B_SobAM2_CrePos_keyvals, FDPthreshold$Threshold, 9, filter)
A12S_SobAM2_CrePos_exp <- lipid_levels(A12S_SobAM2_CrePos_keyvals, FDPthreshold$Threshold, 10, filter)
A12B_SobAM2_CreNeg_exp <- lipid_levels(A12B_SobAM2_CreNeg_keyvals, FDPthreshold$Threshold, 11, filter)
A12S_SobAM2_CreNeg_exp <- lipid_levels(A12S_SobAM2_CreNeg_keyvals, FDPthreshold$Threshold, 12, filter)
A18B_SobAM2_CrePos_exp <- lipid_levels(A18B_SobAM2_CrePos_keyvals, FDPthreshold$Threshold, 13, filter)
A18S_SobAM2_CrePos_exp <- lipid_levels(A18S_SobAM2_CrePos_keyvals, FDPthreshold$Threshold, 14, filter)
A18B_SobAM2_CreNeg_exp <- lipid_levels(A18B_SobAM2_CreNeg_keyvals, FDPthreshold$Threshold, 15, filter)
A18S_SobAM2_CreNeg_exp <- lipid_levels(A18S_SobAM2_CreNeg_keyvals, FDPthreshold$Threshold, 16, filter)

require(openxlsx)
list_brain <- list("A6B_n11" = A6B_exp, "A12B_n13" = A12B_exp,
                   "A18B_n14" = A18B_exp, "A24B_n11" = A24B_exp,
                   "A12B_SobAM2_CrePos_n18" = A12B_SobAM2_CrePos_exp, "A12B_SobAM2_CreNeg_n11" = A12B_SobAM2_CreNeg_exp,
                   "A18B_SobAM2_CrePos_n17" = A18B_SobAM2_CrePos_exp, "A18B_SobAM2_CreNeg_n10" = A18B_SobAM2_CreNeg_exp)
write.xlsx(list_brain, file = "FinalVP/Brain_PLP_regulation.xlsx")

list_spinal <- list("A6S_n11" = A6S_exp, "A12S_n13" = A12S_exp,
                    "A18S_n15" = A18S_exp, "A24S_n10" = A24S_exp,
                    "A12S_SobAM2_CrePos_n18" = A12S_SobAM2_CrePos_exp, "A12S_SobAM2_CreNeg_n11" = A12S_SobAM2_CreNeg_exp,
                    "A18S_SobAM2_CrePos_n18" = A18S_SobAM2_CrePos_exp, "A18S_SobAM2_CreNeg_n10" = A18S_SobAM2_CreNeg_exp)
write.xlsx(list_spinal, file = "FinalVP/Spinal_PLP_regulation.xlsx")

## lipid level count ####

columns <- c("Timepoint", "Increased", "Decreased")
rows <- c("A6B","A6S","A12B","A12S","A18B","A18S", "A24B", "A24S",
          "A12B_SobAM2_CrePos","A12S_SobAM2_CrePos","A12B_SobAM2_CreNeg","A12S_SobAM2_CreNeg",
          "A18B_SobAM2_CrePos","A18S_SobAM2_CrePos","A18B_SobAM2_CreNeg","A18S_SobAM2_CreNeg")
count = data.frame(matrix(nrow = length(rows), ncol = length(columns))) 
colnames(count) = columns
count$Timepoint <- rows

PLP_list <- list("A6B" = A6B_exp, "A6S" = A6S_exp,"A12B" = A12B_exp, "A12S" = A12S_exp,
                 "A18B" = A18B_exp, "A18S" = A18S_exp,"A24B" = A24B_exp, "A24S" = A24S_exp,
                 "A12B_SobAM2_CrePos" = A12B_SobAM2_CrePos_exp, "A12S_SobAM2_CrePos" = A12S_SobAM2_CrePos_exp,
                 "A12B_SobAM2_CreNeg" = A12B_SobAM2_CreNeg_exp, "A12S_SobAM2_CreNeg" = A12S_SobAM2_CreNeg_exp,
                 "A18B_SobAM2_CrePos" = A18B_SobAM2_CrePos_exp, "A18S_SobAM2_CrePos" = A18S_SobAM2_CrePos_exp,
                 "A18B_SobAM2_CreNeg" = A18B_SobAM2_CreNeg_exp, "A18S_SobAM2_CreNeg" = A18S_SobAM2_CreNeg_exp
)

lipid_count <- lipid_level_count(PLP_list, count)

write_csv(lipid_count, "FinalVP/VP_expression_count.csv")
```
