## Supplemental Codes for "Lipidomic analysis reveals differences in the extent of remyelination in the brain and spinal cord": VolPlot_Final.R

```
# Volcano Plots

# To-Do ####

# Re-run all the volcano plots using refreshed data from corrected Data_Cleaning.R script

# Load library #####
library(dplyr)
library(readxl)
library(tidyverse)
library(data.table)
library(igraph)
library(ggplot2)
library(countdata)
library(gtools)
library(devtools)
library(permFDP) #install_github('steven-shuken/permFDP')
library(EnhancedVolcano) #install_github('kevinblighe/EnhancedVolcano')
library(plotrix)
library(ggrastr)

A6_18_B <- read_csv("Final_6B_All_noCE.csv", col_names = TRUE, name_repair = unique)
A6_18_S <- read_csv("Final_6S_All_noCE.csv", col_names = TRUE, name_repair = unique)

# Functions ####
mean.sem <- function(x) {
  c(mean = mean(x), sem = sd(x)/sqrt(length(x)))
}

# PLP W6 Brain ####

A6B <- A6_18_B %>% filter(grepl('wk 6', ...1)) %>% filter(grepl('Control', CRE.and.Treatment))
A6B_no0 <- A6B %>% select(-"Sample.number", -"...1")

# For no 0 df, drop SM(16:1), PE(44:9) and PA(38:3)
# https://stackoverflow.com/questions/72096913/remove-columns-that-have-one-zero-value
print(select(A6B_no0, where(~any(. == 0))))
A6B_no0 <- Filter(function(x) !any(x %in% 0), A6B_no0) 
log_A6B_no0 = log2(A6B_no0[,-ncol(A6B_no0)]) 
log_A6B_no0 <- cbind(log_A6B_no0, 'CRE.and.Treatment' = A6B$CRE.and.Treatment)

# Fix column names & remove extra rows
ttest_A6B_no0 <- transpose(log_A6B_no0, keep.names = 'Lipid species')
ttest_A6B_no0 <- ttest_A6B_no0[-nrow(ttest_A6B_no0),]

samples <- A6B$Sample.number
names <- c('Lipid species', samples)
colnames(ttest_A6B_no0) = c(names)

# Get p values from t test 
ttest_A6B_no0$"pvalue" <- unlist(lapply(1:nrow(ttest_A6B_no0), function(i) t.test((as.numeric(ttest_A6B_no0[i,2:4])), as.numeric(ttest_A6B_no0[i,5:12]))[c("p.value")]))

# calculate average values & sem for each species using CRE and treatment as factor
mean.sem_A6B_no0 <- data.table(log_A6B_no0)[, lapply(.SD, mean.sem), by=CRE.and.Treatment]
mean.sem_A6B_no0 <- transpose(mean.sem_A6B_no0[,2:(ncol(mean.sem_A6B_no0))], keep.names = "Lipid species")
# Make sure the order of columns is correct, some df will have experimental first
CRE_factor <- c("CRE-/control avg","CRE-/control sem","CRE+/control avg","CRE+/control sem")
names <- c('Lipid species', CRE_factor)
colnames(mean.sem_A6B_no0) = c(names)

# add means to main df
#VP_A6B_no0 <- cbind(ttest_A6B_no0, mean.sem_A6B_no0)
VP_A6B_no0 <- merge(ttest_A6B_no0, mean.sem_A6B_no0, by = 'Lipid species')
# Calculate fold change
VP_A6B_no0$"Log2FC" <- VP_A6B_no0$'CRE+/control avg' - VP_A6B_no0$'CRE-/control avg'

#save df 
write_csv(VP_A6B_no0, "VP_csv/A6B_no0_Volcano_plot.csv")

## permFDP ####

VP_A6B_no0 <- read_csv("VP_csv/A6B_no0_Volcano_plot.csv")

pVals = VP_A6B_no0$pvalue
threshold = 0.05
myDesign = c(1,1,1,2,2,2,2,2,2,2,2)
intOnly = VP_A6B_no0 %>% select(-c("Lipid species","pvalue","CRE-/control avg","CRE-/control sem","CRE+/control avg","CRE+/control sem","Log2FC"))
intOnly <- mutate_all(intOnly, function(x) as.numeric(as.character(x))) 
corrThreshold_500 = permFDP::permFDP.adjust.threshold(pVals, threshold, myDesign, intOnly, 500)
corrThreshold_1000 = permFDP::permFDP.adjust.threshold(pVals, threshold, myDesign, intOnly, 1000)
corrThreshold_3000 = permFDP::permFDP.adjust.threshold(pVals, threshold, myDesign, intOnly, 3000)
corrThreshold_10000 = permFDP::permFDP.adjust.threshold(pVals, threshold, myDesign, intOnly, 10000)

print(corrThreshold_3000, digits = 20)
columns <- c("Timepoint", "Threshold")
rows <- c("A6B","A6S","A12B","A12S","A18B","A18S", "A24B", "A24S",
          "A12B_SobAM2_CrePos","A12S_SobAM2_CrePos","A12B_SobAM2_CreNeg","A12S_SobAM2_CreNeg",
          "A18B_SobAM2_CrePos","A18S_SobAM2_CrePos","A18B_SobAM2_CreNeg","A18S_SobAM2_CreNeg")
FDPthreshold = data.frame(matrix(nrow = length(rows), ncol = length(columns))) 
colnames(FDPthreshold) = columns
FDPthreshold$Timepoint <- rows
FDPthreshold[1,2] <- corrThreshold_3000
#corrThreshold_3000 = 0.0019010953080093998847

## EnhancedVolcano ####
pdf("FinalVP/VP_A6B_no0.pdf", width = 1.75, height = 1.24)

# Lipid species, p values, log2 FCs
toptable <- VP_A6B_no0[, c(1,ncol(VP_A6B_no0)-5,ncol(VP_A6B_no0))]
lab = toptable$"Lipid species"
x = "Log2FC"
y = "pvalue"
pCutoff = corrThreshold_3000
FCcutoff = 0.5
#title = "PLP brain samples"
#subtitle = bquote(italic('Week 6 with no 0 values, 3000 iterations, 11 mice'))

keyvals.colour <- ifelse(
  toptable$Log2FC > 0.5 & toptable$pvalue < pCutoff, '#FF00FF',
  ifelse(toptable$Log2FC <(-0.5) & toptable$pvalue < pCutoff, '#00FFFF',
         '#808080'))
# Gray - #A8B6CC - #DAECCF
# Magenta - #F05039 - #F32A50
# Cyan - #0090FF - #0882E0

keyvals.colour[is.na(keyvals.colour)] <- '#808080'
names(keyvals.colour)[keyvals.colour == '#FF00FF'] <- 'High FC & sig'
names(keyvals.colour)[keyvals.colour == '#808080'] <- 'NS'
names(keyvals.colour)[keyvals.colour == '#00FFFF'] <- 'Low FC & sig'

EnhancedVolcano(toptable, lab, x, y, selectLab = rownames(toptable)[which(names(keyvals.colour) %in% c('High', 'Low'))],
                xlab = bquote(~Log[2] ~ "Fold Change"),legendPosition = 'none',
                ylab = bquote(~-Log[10] ~ italic(p) ~ Value), colCustom = keyvals.colour,
                title = NULL, subtitle = NULL, pCutoff = pCutoff, pCutoffCol = y, FCcutoff = FCcutoff,
                drawConnectors = FALSE, xlim = c(min(toptable[[x]], na.rm = TRUE) - 0.2, max(toptable[[x]], na.rm = TRUE) + 0.2),
                ylim = c(0, max(-log10(toptable[[y]]), na.rm = TRUE) + 0.5), colAlpha = 1,
                cutoffLineWidth = 0.1, pointSize = 0.5,axisLabSize = 5, captionLabSize = 3, borderWidth = 0.1,
                vlineWidth = 0.1, hlineWidth = 0.1, caption = NULL
) + theme(axis.text.x = element_text(face="plain", color="black", size=5, angle=0),
          axis.text.y = element_text(face="plain", color="black", size=5, angle=0),
          axis.ticks.length=unit(.05, "cm"), axis.ticks=element_line(linewidth=0.2),
          axis.title.x = element_text(family = "sans", size = 5, margin=margin(1,0,0,0)), 
          axis.title.y = element_text(family = "sans", size = 5, margin=margin(0,1,0,0)),
          panel.grid.major = element_line(linewidth = 0.5), 
          panel.grid.minor = element_line(linewidth = 0.3),
          plot.margin = margin(4,4,4,4)
)

dev.off()

# PLP W6 Spinal ####

A6S <- A6_18_S %>% filter(grepl('wk 6', ...1)) %>% filter(grepl('Control', CRE.and.Treatment))
A6S_no0 <- A6S %>% select(-"Sample.number", -"...1")

# For no 0 df, drop ePC(38:6) and PS(32:1)
# https://stackoverflow.com/questions/72096913/remove-columns-that-have-one-zero-value
print(select(A6S_no0, where(~any(. == 0))))
A6S_no0 <- Filter(function(x) !any(x %in% 0), A6S_no0) 
log_A6S_no0 = log2(A6S_no0[,-ncol(A6S_no0)]) 
log_A6S_no0 <- cbind(log_A6S_no0, 'CRE.and.Treatment' = A6S$CRE.and.Treatment)

# Fix column names & remove extra rows
ttest_A6S_no0 <- transpose(log_A6S_no0, keep.names = 'Lipid species')
ttest_A6S_no0 <- ttest_A6S_no0[-nrow(ttest_A6S_no0),]

samples <- A6S$Sample.number
names <- c('Lipid species', samples)
colnames(ttest_A6S_no0) = c(names)

# Get p values from t test 
ttest_A6S_no0$"pvalue" <- unlist(lapply(1:nrow(ttest_A6S_no0), function(i) t.test((as.numeric(ttest_A6S_no0[i,2:4])), as.numeric(ttest_A6S_no0[i,5:12]))[c("p.value")]))

# calculate average values & sem for each species using CRE and treatment as factor
mean.sem_A6S_no0 <- data.table(log_A6S_no0)[, lapply(.SD, mean.sem), by=CRE.and.Treatment]
mean.sem_A6S_no0 <- transpose(mean.sem_A6S_no0[,2:(ncol(mean.sem_A6S_no0))], keep.names = "Lipid species")
# Make sure the order of columns is correct, some df will have experimental first
CRE_factor <- c("CRE-/control avg","CRE-/control sem","CRE+/control avg","CRE+/control sem")
names <- c('Lipid species', CRE_factor)
colnames(mean.sem_A6S_no0) = c(names)

# add means to main df
#VP_A6S_no0 <- cbind(ttest_A6S_no0, mean.sem_A6S_no0)
VP_A6S_no0 <- merge(ttest_A6S_no0, mean.sem_A6S_no0, by = 'Lipid species')
# Calculate fold change
VP_A6S_no0$"Log2FC" <- VP_A6S_no0$'CRE+/control avg' - VP_A6S_no0$'CRE-/control avg'

#save df 
write_csv(VP_A6S_no0, "VP_csv/A6S_no0_Volcano_plot.csv")

## permFDP ####

VP_A6S_no0 <- read_csv("VP_csv/A6S_no0_Volcano_plot.csv")

pVals = VP_A6S_no0$pvalue
threshold = 0.05
myDesign = c(1,1,1,2,2,2,2,2,2,2,2)
intOnly = VP_A6S_no0 %>% select(-c("Lipid species","pvalue","CRE-/control avg","CRE-/control sem","CRE+/control avg","CRE+/control sem","Log2FC"))
intOnly <- mutate_all(intOnly, function(x) as.numeric(as.character(x))) 
corrThreshold_500 = permFDP::permFDP.adjust.threshold(pVals, threshold, myDesign, intOnly, 500)
corrThreshold_1000 = permFDP::permFDP.adjust.threshold(pVals, threshold, myDesign, intOnly, 1000)
corrThreshold_3000 = permFDP::permFDP.adjust.threshold(pVals, threshold, myDesign, intOnly, 3000)
corrThreshold_10000 = permFDP::permFDP.adjust.threshold(pVals, threshold, myDesign, intOnly, 10000)

print(corrThreshold_3000, digits = 20)
FDPthreshold[2,2] <- corrThreshold_3000
#corrThreshold_3000 = 0.007261606023270235008

## EnhancedVolcano ####

pdf("FinalVP/VP_A6S_no0.pdf",  width = 1.75, height = 1.24)

# Lipid species, p values, log2 FCs
toptable <- VP_A6S_no0[, c(1,ncol(VP_A6S_no0)-5,ncol(VP_A6S_no0))]
lab = toptable$"Lipid species"
x = "Log2FC"
y = "pvalue"
pCutoff = corrThreshold_3000
FCcutoff = 0.5
#title = "PLP spinal cord samples"
#subtitle = bquote(italic('Week 6 with no 0 values, 3000 iterations, 11 mice'))

keyvals.colour <- ifelse(
  toptable$Log2FC > 0.5 & toptable$pvalue < pCutoff, '#FF00FF',
  ifelse(toptable$Log2FC <(-0.5) & toptable$pvalue < pCutoff, '#00FFFF',
         '#808080'))

keyvals.colour[is.na(keyvals.colour)] <- '#808080'
names(keyvals.colour)[keyvals.colour == '#FF00FF'] <- 'High FC & sig'
names(keyvals.colour)[keyvals.colour == '#808080'] <- 'NS'
names(keyvals.colour)[keyvals.colour == '#00FFFF'] <- 'Low FC & sig'


EnhancedVolcano(toptable, lab, x, y, selectLab = rownames(toptable)[which(names(keyvals.colour) %in% c('High', 'Low'))],
                xlab = bquote(~Log[2] ~ "Fold Change"),legendPosition = 'none',
                ylab = bquote(~-Log[10] ~ italic(p) ~ Value), colCustom = keyvals.colour,
                title = NULL, subtitle = NULL, pCutoff = pCutoff, pCutoffCol = y, FCcutoff = FCcutoff,
                drawConnectors = FALSE, xlim = c(min(toptable[[x]], na.rm = TRUE) - 0.2, max(toptable[[x]], na.rm = TRUE) + 0.2),
                ylim = c(0, max(-log10(toptable[[y]]), na.rm = TRUE) + 0.5), colAlpha = 1,
                cutoffLineWidth = 0.1, pointSize = 0.5,axisLabSize = 5, captionLabSize = 3, borderWidth = 0.1,
                vlineWidth = 0.1, hlineWidth = 0.1, caption = NULL
) + theme(axis.text.x = element_text(face="plain", color="black", size=5, angle=0),
          axis.text.y = element_text(face="plain", color="black", size=5, angle=0),
          axis.ticks.length=unit(.05, "cm"), axis.ticks=element_line(linewidth=0.2),
          axis.title.x = element_text(family = "sans", size = 5, margin=margin(1,0,0,0)), 
          axis.title.y = element_text(family = "sans", size = 5, margin=margin(0,1,0,0)),
          panel.grid.major = element_line(linewidth = 0.5), 
          panel.grid.minor = element_line(linewidth = 0.3),
          plot.margin = margin(4,4,4,4)
)

dev.off()

# PLP W12 Brain ####

A12B <- A6_18_B %>% filter(grepl('wk 12', ...1)) %>% filter(grepl('Control', CRE.and.Treatment))
A12B_no0 <- A12B %>% select(-"Sample.number", -"...1")

# For no 0 df, drop PS(32:1) and ePS(38:4)
# https://stackoverflow.com/questions/72096913/remove-columns-that-have-one-zero-value
print(select(A12B_no0, where(~any(. == 0))))
A12B_no0 <- Filter(function(x) !any(x %in% 0), A12B_no0) 
log_A12B_no0 = log2(A12B_no0[,-ncol(A12B_no0)]) 
log_A12B_no0 <- cbind(log_A12B_no0, 'CRE.and.Treatment' = A12B$CRE.and.Treatment)

# Fix column names & remove extra rows
ttest_A12B_no0 <- transpose(log_A12B_no0, keep.names = 'Lipid species')
ttest_A12B_no0 <- ttest_A12B_no0[-nrow(ttest_A12B_no0),]

samples <- A12B$Sample.number
names <- c('Lipid species', samples)
colnames(ttest_A12B_no0) = c(names)

# Get p values from t test 
ttest_A12B_no0$"pvalue" <- unlist(lapply(1:nrow(ttest_A12B_no0), function(i) t.test((as.numeric(ttest_A12B_no0[i,10:14])), as.numeric(ttest_A12B_no0[i,2:9]))[c("p.value")]))

# calculate average values & sem for each species using CRE and treatment as factor
mean.sem_A12B_no0 <- data.table(log_A12B_no0)[, lapply(.SD, mean.sem), by=CRE.and.Treatment]
mean.sem_A12B_no0 <- transpose(mean.sem_A12B_no0[,2:(ncol(mean.sem_A12B_no0))], keep.names = "Lipid species")
# Make sure the order of columns is correct, some df will have experimental first
CRE_factor <- c("CRE+/control avg","CRE+/control sem","CRE-/control avg","CRE-/control sem")
names <- c('Lipid species', CRE_factor)
colnames(mean.sem_A12B_no0) = c(names)

# add means to main df
#VP_A12B_no0 <- cbind(ttest_A12B_no0, mean.sem_A12B_no0)
VP_A12B_no0 <- merge(ttest_A12B_no0, mean.sem_A12B_no0, by = 'Lipid species')
# Calculate fold change
VP_A12B_no0$"Log2FC" <- VP_A12B_no0$'CRE+/control avg' - VP_A12B_no0$'CRE-/control avg'

#save df 
write_csv(VP_A12B_no0, "VP_csv/A12B_no0_Volcano_plot.csv")

## permFDP ####

VP_A12B_no0 <- read_csv("VP_csv/A12B_no0_Volcano_plot.csv")

pVals = VP_A12B_no0$pvalue
threshold = 0.05
myDesign = c(2,2,2,2,2,2,2,2,1,1,1,1,1)
intOnly = VP_A12B_no0 %>% select(-c("Lipid species","pvalue","CRE-/control avg","CRE-/control sem","CRE+/control avg","CRE+/control sem","Log2FC"))
intOnly <- mutate_all(intOnly, function(x) as.numeric(as.character(x))) 
corrThreshold_500 = permFDP::permFDP.adjust.threshold(pVals, threshold, myDesign, intOnly, 500)
corrThreshold_1000 = permFDP::permFDP.adjust.threshold(pVals, threshold, myDesign, intOnly, 1000)
corrThreshold_3000 = permFDP::permFDP.adjust.threshold(pVals, threshold, myDesign, intOnly, 3000)
corrThreshold_10000 = permFDP::permFDP.adjust.threshold(pVals, threshold, myDesign, intOnly, 10000)

print(corrThreshold_3000, digits = 20)
FDPthreshold[3,2] <- corrThreshold_3000
#corrThreshold_3000 = 0.019324139678623689087
## EnhancedVolcano ####

pdf("FinalVP/VP_A12B_no0.pdf", width = 1.75, height = 1.24)

# Lipid species, p values, log2 FCs
toptable <- VP_A12B_no0[, c(1,ncol(VP_A12B_no0)-5,ncol(VP_A12B_no0))]
lab = toptable$"Lipid species"
x = "Log2FC"
y = "pvalue"
pCutoff = corrThreshold_3000
FCcutoff = 0.5
#title = "PLP brain samples"
#subtitle = bquote(italic('Week 12 with no 0 values, 3000 iterations, 13 mice'))

keyvals.colour <- ifelse(
  toptable$Log2FC > 0.5 & toptable$pvalue < pCutoff, '#FF00FF',
  ifelse(toptable$Log2FC <(-0.5) & toptable$pvalue < pCutoff, '#00FFFF',
         '#808080'))
# Gray - #A8B6CC - #DAECCF
# Magenta - #F05039 - #F32A50
# Cyan - #0090FF - #0882E0

keyvals.colour[is.na(keyvals.colour)] <- '#808080'
names(keyvals.colour)[keyvals.colour == '#FF00FF'] <- 'High FC & sig'
names(keyvals.colour)[keyvals.colour == '#808080'] <- 'NS'
names(keyvals.colour)[keyvals.colour == '#00FFFF'] <- 'Low FC & sig'

EnhancedVolcano(toptable, lab, x, y, selectLab = rownames(toptable)[which(names(keyvals.colour) %in% c('High', 'Low'))],
                xlab = bquote(~Log[2] ~ "Fold Change"),legendPosition = 'none',
                ylab = bquote(~-Log[10] ~ italic(p) ~ Value), colCustom = keyvals.colour,
                title = NULL, subtitle = NULL, pCutoff = pCutoff, pCutoffCol = y, FCcutoff = FCcutoff,
                drawConnectors = FALSE, xlim = c(min(toptable[[x]], na.rm = TRUE) - 0.2, max(toptable[[x]], na.rm = TRUE) + 0.2),
                ylim = c(0, max(-log10(toptable[[y]]), na.rm = TRUE) + 0.5), colAlpha = 1,
                cutoffLineWidth = 0.1, pointSize = 0.5,axisLabSize = 5, captionLabSize = 3, borderWidth = 0.1,
                vlineWidth = 0.1, hlineWidth = 0.1, caption = NULL
) + theme(axis.text.x = element_text(face="plain", color="black", size=5, angle=0),
          axis.text.y = element_text(face="plain", color="black", size=5, angle=0),
          axis.ticks.length=unit(.05, "cm"), axis.ticks=element_line(linewidth=0.2),
          axis.title.x = element_text(family = "sans", size = 5, margin=margin(1,0,0,0)), 
          axis.title.y = element_text(family = "sans", size = 5, margin=margin(0,1,0,0)),
          panel.grid.major = element_line(linewidth = 0.5), 
          panel.grid.minor = element_line(linewidth = 0.3),
          plot.margin = margin(4,4,4,4)
)

dev.off()

# PLP W12 Spinal ####

A12S <- A6_18_S %>% filter(grepl('wk 12', ...1)) %>% filter(grepl('Control', CRE.and.Treatment))
A12S_no0 <- A12S %>% select(-"Sample.number", -"...1")

# For no 0 df, drop PC(40:3), PE(40:5) and PS(32:1)
# https://stackoverflow.com/questions/72096913/remove-columns-that-have-one-zero-value
print(select(A12S_no0, where(~any(. == 0))))
A12S_no0 <- Filter(function(x) !any(x %in% 0), A12S_no0) 
log_A12S_no0 = log2(A12S_no0[,-ncol(A12S_no0)]) 
log_A12S_no0 <- cbind(log_A12S_no0, 'CRE.and.Treatment' = A12S$CRE.and.Treatment)

# Fix column names & remove extra rows
ttest_A12S_no0 <- transpose(log_A12S_no0, keep.names = 'Lipid species')
ttest_A12S_no0 <- ttest_A12S_no0[-nrow(ttest_A12S_no0),]

samples <- A12S$Sample.number
names <- c('Lipid species', samples)
colnames(ttest_A12S_no0) = c(names)

# Get p values from t test 
ttest_A12S_no0$"pvalue" <- unlist(lapply(1:nrow(ttest_A12S_no0), function(i) t.test((as.numeric(ttest_A12S_no0[i,2:6])), as.numeric(ttest_A12S_no0[i,7:14]))[c("p.value")]))

# calculate average values & sem for each species using CRE and treatment as factor
mean.sem_A12S_no0 <- data.table(log_A12S_no0)[, lapply(.SD, mean.sem), by=CRE.and.Treatment]
mean.sem_A12S_no0 <- transpose(mean.sem_A12S_no0[,2:(ncol(mean.sem_A12S_no0))], keep.names = "Lipid species")
# Make sure the order of columns is correct, some df will have experimental first
CRE_factor <- c("CRE-/control avg","CRE-/control sem","CRE+/control avg","CRE+/control sem")
names <- c('Lipid species', CRE_factor)
colnames(mean.sem_A12S_no0) = c(names)

# add means to main df
#VP_A12S_no0 <- cbind(ttest_A12S_no0, mean.sem_A12S_no0)
VP_A12S_no0 <- merge(ttest_A12S_no0, mean.sem_A12S_no0, by = 'Lipid species')
# Calculate fold change
VP_A12S_no0$"Log2FC" <- VP_A12S_no0$'CRE+/control avg' - VP_A12S_no0$'CRE-/control avg'

#save df 
write_csv(VP_A12S_no0, "VP_csv/A12S_no0_Volcano_plot.csv")

## permFDP ####

VP_A12S_no0 <- read_csv("VP_csv/A12S_no0_Volcano_plot.csv")

pVals = VP_A12S_no0$pvalue
threshold = 0.05
myDesign = c(1,1,1,1,1,2,2,2,2,2,2,2,2)
intOnly = VP_A12S_no0 %>% select(-c("Lipid species","pvalue","CRE-/control avg","CRE-/control sem","CRE+/control avg","CRE+/control sem","Log2FC"))
intOnly <- mutate_all(intOnly, function(x) as.numeric(as.character(x))) 
corrThreshold_500 = permFDP::permFDP.adjust.threshold(pVals, threshold, myDesign, intOnly, 500)
corrThreshold_1000 = permFDP::permFDP.adjust.threshold(pVals, threshold, myDesign, intOnly, 1000)
corrThreshold_3000 = permFDP::permFDP.adjust.threshold(pVals, threshold, myDesign, intOnly, 3000)
corrThreshold_10000 = permFDP::permFDP.adjust.threshold(pVals, threshold, myDesign, intOnly, 10000)

print(corrThreshold_3000, digits = 20)
FDPthreshold[4,2] <- corrThreshold_3000
#corrThreshold_3000 = 0.070363141959697228511
## EnhancedVolcano ####

pdf("FinalVP/VP_A12S_no0.pdf", width = 1.75, height = 1.24)

# Lipid species, p values, log2 FCs
toptable <- VP_A12S_no0[, c(1,ncol(VP_A12S_no0)-5,ncol(VP_A12S_no0))]
lab = toptable$"Lipid species"
x = "Log2FC"
y = "pvalue"
pCutoff = corrThreshold_3000
FCcutoff = 0.5
#title = "PLP spinal cord samples"
#subtitle = bquote(italic('Week 12 with no 0 values, 3000 iterations, 13 mice'))

keyvals.colour <- ifelse(
  toptable$Log2FC > 0.5 & toptable$pvalue < pCutoff, '#FF00FF',
  ifelse(toptable$Log2FC <(-0.5) & toptable$pvalue < pCutoff, '#00FFFF',
         '#808080'))

keyvals.colour[is.na(keyvals.colour)] <- '#808080'
names(keyvals.colour)[keyvals.colour == '#FF00FF'] <- 'High FC & sig'
names(keyvals.colour)[keyvals.colour == '#808080'] <- 'NS'
names(keyvals.colour)[keyvals.colour == '#00FFFF'] <- 'Low FC & sig'

EnhancedVolcano(toptable, lab, x, y, selectLab = rownames(toptable)[which(names(keyvals.colour) %in% c('High', 'Low'))],
                xlab = bquote(~Log[2] ~ "Fold Change"),legendPosition = 'none',
                ylab = bquote(~-Log[10] ~ italic(p) ~ Value), colCustom = keyvals.colour,
                title = NULL, subtitle = NULL, pCutoff = pCutoff, pCutoffCol = y, FCcutoff = FCcutoff,
                drawConnectors = FALSE, xlim = c(min(toptable[[x]], na.rm = TRUE) - 0.2, max(toptable[[x]], na.rm = TRUE) + 0.2),
                ylim = c(0, max(-log10(toptable[[y]]), na.rm = TRUE) + 0.5), colAlpha = 1,
                cutoffLineWidth = 0.1, pointSize = 0.5,axisLabSize = 5, captionLabSize = 3, borderWidth = 0.1,
                vlineWidth = 0.1, hlineWidth = 0.1, caption = NULL
) + theme(axis.text.x = element_text(face="plain", color="black", size=5, angle=0),
          axis.text.y = element_text(face="plain", color="black", size=5, angle=0),
          axis.ticks.length=unit(.05, "cm"), axis.ticks=element_line(linewidth=0.2),
          axis.title.x = element_text(family = "sans", size = 5, margin=margin(1,0,0,0)), 
          axis.title.y = element_text(family = "sans", size = 5, margin=margin(0,1,0,0)),
          panel.grid.major = element_line(linewidth = 0.5), 
          panel.grid.minor = element_line(linewidth = 0.3),
          plot.margin = margin(4,4,4,4)
)

dev.off()


# PLP W18 Brain ####

A18B <- A6_18_B %>% filter(grepl('wk 18', ...1)) %>% filter(grepl('Control', CRE.and.Treatment))
A18B_no0 <- A18B %>% select(-"Sample.number", -"...1")

# For no 0 df, drop NONE
# https://stackoverflow.com/questions/72096913/remove-columns-that-have-one-zero-value
print(select(A18B_no0, where(~any(. == 0))))
A18B_no0 <- Filter(function(x) !any(x %in% 0), A18B_no0) 
log_A18B_no0 = log2(A18B_no0[,-ncol(A18B_no0)]) 
log_A18B_no0 <- cbind(log_A18B_no0, 'CRE.and.Treatment' = A18B$CRE.and.Treatment)

# Fix column names & remove extra rows
ttest_A18B_no0 <- transpose(log_A18B_no0, keep.names = 'Lipid species')
ttest_A18B_no0 <- ttest_A18B_no0[-nrow(ttest_A18B_no0),]

samples <- A18B$Sample.number
names <- c('Lipid species', samples)
colnames(ttest_A18B_no0) = c(names)

# Get p values from t test 
ttest_A18B_no0$"pvalue" <- unlist(lapply(1:nrow(ttest_A18B_no0), function(i) t.test((as.numeric(ttest_A18B_no0[i,10:15])), as.numeric(ttest_A18B_no0[i,2:9]))[c("p.value")]))

# calculate average values & sem for each species using CRE and treatment as factor
mean.sem_A18B_no0 <- data.table(log_A18B_no0)[, lapply(.SD, mean.sem), by=CRE.and.Treatment]
mean.sem_A18B_no0 <- transpose(mean.sem_A18B_no0[,2:(ncol(mean.sem_A18B_no0))], keep.names = "Lipid species")
# Make sure the order of columns is correct, some df will have experimental first
CRE_factor <- c("CRE+/control avg","CRE+/control sem","CRE-/control avg","CRE-/control sem")
names <- c('Lipid species', CRE_factor)
colnames(mean.sem_A18B_no0) = c(names)

# add means to main df
#VP_A18B_no0 <- cbind(ttest_A18B_no0, mean.sem_A18B_no0)
VP_A18B_no0 <- merge(ttest_A18B_no0, mean.sem_A18B_no0, by = 'Lipid species')
# Calculate fold change
VP_A18B_no0$"Log2FC" <- VP_A18B_no0$'CRE+/control avg' - VP_A18B_no0$'CRE-/control avg'

#save df 
write_csv(VP_A18B_no0, "VP_csv/A18B_no0_Volcano_plot.csv")

## permFDP ####

VP_A18B_no0 <- read_csv("VP_csv/A18B_no0_Volcano_plot.csv")

pVals = VP_A18B_no0$pvalue
threshold = 0.05
myDesign = c(2,2,2,2,2,2,2,2,1,1,1,1,1,1)
intOnly = VP_A18B_no0 %>% select(-c("Lipid species","pvalue","CRE-/control avg","CRE-/control sem","CRE+/control avg","CRE+/control sem","Log2FC"))
intOnly <- mutate_all(intOnly, function(x) as.numeric(as.character(x))) 
corrThreshold_500 = permFDP::permFDP.adjust.threshold(pVals, threshold, myDesign, intOnly, 500)
corrThreshold_1000 = permFDP::permFDP.adjust.threshold(pVals, threshold, myDesign, intOnly, 1000)
corrThreshold_3000 = permFDP::permFDP.adjust.threshold(pVals, threshold, myDesign, intOnly, 3000)
corrThreshold_10000 = permFDP::permFDP.adjust.threshold(pVals, threshold, myDesign, intOnly, 10000)

print(corrThreshold_3000, digits = 20)
FDPthreshold[5,2] <- corrThreshold_3000
#corrThreshold_3000 = 0.043547652826758480316
## EnhancedVolcano ####

pdf("FinalVP/VP_A18B_no0.pdf", width = 1.75, height = 1.24)

# Lipid species, p values, log2 FCs
toptable <- VP_A18B_no0[, c(1,ncol(VP_A18B_no0)-5,ncol(VP_A18B_no0))]
lab = toptable$"Lipid species"
x = "Log2FC"
y = "pvalue"
pCutoff = corrThreshold_3000
FCcutoff = 0.5
#title = "PLP brain samples"
#subtitle = bquote(italic('Week 18 with no 0 values, 3000 iterations, 14 mice'))

keyvals.colour <- ifelse(
  toptable$Log2FC > 0.5 & toptable$pvalue < pCutoff, '#FF00FF',
  ifelse(toptable$Log2FC <(-0.5) & toptable$pvalue < pCutoff, '#00FFFF',
         '#808080'))
# Gray - #A8B6CC - #DAECCF
# Magenta - #F05039 - #F32A50
# Cyan - #0090FF - #0882E0

keyvals.colour[is.na(keyvals.colour)] <- '#808080'
names(keyvals.colour)[keyvals.colour == '#FF00FF'] <- 'High FC & sig'
names(keyvals.colour)[keyvals.colour == '#808080'] <- 'NS'
names(keyvals.colour)[keyvals.colour == '#00FFFF'] <- 'Low FC & sig'

EnhancedVolcano(toptable, lab, x, y, selectLab = rownames(toptable)[which(names(keyvals.colour) %in% c('High', 'Low'))],
                xlab = bquote(~Log[2] ~ "Fold Change"),legendPosition = 'none',
                ylab = bquote(~-Log[10] ~ italic(p) ~ Value), colCustom = keyvals.colour,
                title = NULL, subtitle = NULL, pCutoff = pCutoff, pCutoffCol = y, FCcutoff = FCcutoff,
                drawConnectors = FALSE, xlim = c(min(toptable[[x]], na.rm = TRUE) - 0.2, max(toptable[[x]], na.rm = TRUE) + 0.2),
                ylim = c(0, max(-log10(toptable[[y]]), na.rm = TRUE) + 0.5), colAlpha = 1,
                cutoffLineWidth = 0.1, pointSize = 0.5,axisLabSize = 5, captionLabSize = 3, borderWidth = 0.1,
                vlineWidth = 0.1, hlineWidth = 0.1, caption = NULL
) + theme(axis.text.x = element_text(face="plain", color="black", size=5, angle=0),
          axis.text.y = element_text(face="plain", color="black", size=5, angle=0),
          axis.ticks.length=unit(.05, "cm"), axis.ticks=element_line(linewidth=0.2),
          axis.title.x = element_text(family = "sans", size = 5, margin=margin(1,0,0,0)), 
          axis.title.y = element_text(family = "sans", size = 5, margin=margin(0,1,0,0)),
          panel.grid.major = element_line(linewidth = 0.5), 
          panel.grid.minor = element_line(linewidth = 0.3),
          plot.margin = margin(4,4,4,4)
)

dev.off()


# PLP W18 Spinal ####

A18S <- A6_18_S %>% filter(grepl('wk 18', ...1)) %>% filter(grepl('Control', CRE.and.Treatment))
A18S_no0 <- A18S %>% select(-"Sample.number", -"...1")

# For no 0 df, drop PC(40:3), PS(32:1), ePS(38:4) and ePS(40:4)
# https://stackoverflow.com/questions/72096913/remove-columns-that-have-one-zero-value
print(select(A18S_no0, where(~any(. == 0))))
A18S_no0 <- Filter(function(x) !any(x %in% 0), A18S_no0) 
log_A18S_no0 = log2(A18S_no0[,-ncol(A18S_no0)]) 
log_A18S_no0 <- cbind(log_A18S_no0, 'CRE.and.Treatment' = A18S$CRE.and.Treatment)

# Fix column names & remove extra rows
ttest_A18S_no0 <- transpose(log_A18S_no0, keep.names = 'Lipid species')
ttest_A18S_no0 <- ttest_A18S_no0[-nrow(ttest_A18S_no0),]

samples <- A18S$Sample.number
names <- c('Lipid species', samples)
colnames(ttest_A18S_no0) = c(names)

# Get p values from t test 
ttest_A18S_no0$"pvalue" <- unlist(lapply(1:nrow(ttest_A18S_no0), function(i) t.test((as.numeric(ttest_A18S_no0[i,2:7])), as.numeric(ttest_A18S_no0[i,8:16]))[c("p.value")]))

# calculate average values & sem for each species using CRE and treatment as factor
mean.sem_A18S_no0 <- data.table(log_A18S_no0)[, lapply(.SD, mean.sem), by=CRE.and.Treatment]
mean.sem_A18S_no0 <- transpose(mean.sem_A18S_no0[,2:(ncol(mean.sem_A18S_no0))], keep.names = "Lipid species")
# Make sure the order of columns is correct, some df will have experimental first
CRE_factor <- c("CRE-/control avg","CRE-/control sem","CRE+/control avg","CRE+/control sem")
names <- c('Lipid species', CRE_factor)
colnames(mean.sem_A18S_no0) = c(names)

# add means to main df
#VP_A18S_no0 <- cbind(ttest_A18S_no0, mean.sem_A18S_no0)
VP_A18S_no0 <- merge(ttest_A18S_no0, mean.sem_A18S_no0, by = 'Lipid species')
# Calculate fold change
VP_A18S_no0$"Log2FC" <- VP_A18S_no0$'CRE+/control avg' - VP_A18S_no0$'CRE-/control avg'

#save df 
write_csv(VP_A18S_no0, "VP_csv/A18S_no0_Volcano_plot.csv")

## permFDP ####

VP_A18S_no0 <- read_csv("VP_csv/A18S_no0_Volcano_plot.csv")

pVals = VP_A18S_no0$pvalue
threshold = 0.05
myDesign = c(1,1,1,1,1,1,2,2,2,2,2,2,2,2,2)
intOnly = VP_A18S_no0 %>% select(-c("Lipid species","pvalue","CRE-/control avg","CRE-/control sem","CRE+/control avg","CRE+/control sem","Log2FC"))
intOnly <- mutate_all(intOnly, function(x) as.numeric(as.character(x))) 
corrThreshold_500 = permFDP::permFDP.adjust.threshold(pVals, threshold, myDesign, intOnly, 500)
corrThreshold_1000 = permFDP::permFDP.adjust.threshold(pVals, threshold, myDesign, intOnly, 1000)
corrThreshold_3000 = permFDP::permFDP.adjust.threshold(pVals, threshold, myDesign, intOnly, 3000)
corrThreshold_10000 = permFDP::permFDP.adjust.threshold(pVals, threshold, myDesign, intOnly, 10000)

print(corrThreshold_3000, digits = 20)
FDPthreshold[6,2] <- corrThreshold_3000
#corrThreshold_3000 = 0.096126099761683028722
## EnhancedVolcano ####

pdf("FinalVP/VP_A18S_no0.pdf", width = 1.75, height = 1.24)

# Lipid species, p values, log2 FCs
toptable <- VP_A18S_no0[, c(1,ncol(VP_A18S_no0)-5,ncol(VP_A18S_no0))]
lab = toptable$"Lipid species"
x = "Log2FC"
y = "pvalue"
pCutoff = corrThreshold_3000
FCcutoff = 0.5
#title = "PLP spinal cord samples"
#subtitle = bquote(italic('Week 18 with no 0 values, 3000 iterations, 15 mice'))

keyvals.colour <- ifelse(
  toptable$Log2FC > 0.5 & toptable$pvalue < pCutoff, '#FF00FF',
  ifelse(toptable$Log2FC <(-0.5) & toptable$pvalue < pCutoff, '#00FFFF',
         '#808080'))

keyvals.colour[is.na(keyvals.colour)] <- '#808080'
names(keyvals.colour)[keyvals.colour == '#FF00FF'] <- 'High FC & sig'
names(keyvals.colour)[keyvals.colour == '#808080'] <- 'NS'
names(keyvals.colour)[keyvals.colour == '#00FFFF'] <- 'Low FC & sig'

EnhancedVolcano(toptable, lab, x, y, selectLab = rownames(toptable)[which(names(keyvals.colour) %in% c('High', 'Low'))],
                xlab = bquote(~Log[2] ~ "Fold Change"),legendPosition = 'none',
                ylab = bquote(~-Log[10] ~ italic(p) ~ Value), colCustom = keyvals.colour,
                title = NULL, subtitle = NULL, pCutoff = pCutoff, pCutoffCol = y, FCcutoff = FCcutoff,
                drawConnectors = FALSE, xlim = c(min(toptable[[x]], na.rm = TRUE) - 0.2, max(toptable[[x]], na.rm = TRUE) + 0.2),
                ylim = c(0, max(-log10(toptable[[y]]), na.rm = TRUE) + 0.5), colAlpha = 1,
                cutoffLineWidth = 0.1, pointSize = 0.5,axisLabSize = 5, captionLabSize = 3, borderWidth = 0.1,
                vlineWidth = 0.1, hlineWidth = 0.1, caption = NULL
) + theme(axis.text.x = element_text(face="plain", color="black", size=5, angle=0),
          axis.text.y = element_text(face="plain", color="black", size=5, angle=0),
          axis.ticks.length=unit(.05, "cm"), axis.ticks=element_line(linewidth=0.2),
          axis.title.x = element_text(family = "sans", size = 5, margin=margin(1,0,0,0)), 
          axis.title.y = element_text(family = "sans", size = 5, margin=margin(0,1,0,0)),
          panel.grid.major = element_line(linewidth = 0.5), 
          panel.grid.minor = element_line(linewidth = 0.3),
          plot.margin = margin(4,4,4,4)
)

dev.off()


# PLP W24 Brain ####

A24B <- read_csv("Final_24B_All_noCE.csv", col_names = TRUE, name_repair = unique)
A24B <- A24B %>% filter(grepl('wk 24', ...1)) %>% filter(grepl('Control', CRE.and.Treatment))
A24B_no0 <- A24B %>% select(-"Sample.number", -"...1")

# For no 0 df, drop PC(36:3), PC(44:10) and ePC(38:3)
# https://stackoverflow.com/questions/72096913/remove-columns-that-have-one-zero-value
print(select(A24B_no0, where(~any(. == 0))))
A24B_no0 <- Filter(function(x) !any(x %in% 0), A24B_no0) 
log_A24B_no0 = log2(A24B_no0[,-ncol(A24B_no0)]) 
log_A24B_no0 <- cbind(log_A24B_no0, 'CRE.and.Treatment' = A24B$CRE.and.Treatment)

# Fix column names & remove extra rows
ttest_A24B_no0 <- transpose(log_A24B_no0, keep.names = 'Lipid species')
ttest_A24B_no0 <- ttest_A24B_no0[-nrow(ttest_A24B_no0),]

samples <- A24B$Sample.number
names <- c('Lipid species', samples)
colnames(ttest_A24B_no0) = c(names)

# Get p values from t test 
ttest_A24B_no0$"pvalue" <- unlist(lapply(1:nrow(ttest_A24B_no0), function(i) t.test((as.numeric(ttest_A24B_no0[i,2:5])), as.numeric(ttest_A24B_no0[i,6:12]))[c("p.value")]))

# calculate average values & sem for each species using CRE and treatment as factor
mean.sem_A24B_no0 <- data.table(log_A24B_no0)[, lapply(.SD, mean.sem), by=CRE.and.Treatment]
mean.sem_A24B_no0 <- transpose(mean.sem_A24B_no0[,2:(ncol(mean.sem_A24B_no0))], keep.names = "Lipid species")
# Make sure the order of columns is correct, some df will have experimental first
CRE_factor <- c("CRE-/control avg","CRE-/control sem","CRE+/control avg","CRE+/control sem")
names <- c('Lipid species', CRE_factor)
colnames(mean.sem_A24B_no0) = c(names)

# add means to main df
#VP_A24B_no0 <- cbind(ttest_A24B_no0, mean.sem_A24B_no0)
VP_A24B_no0 <- merge(ttest_A24B_no0, mean.sem_A24B_no0, by = 'Lipid species')
# Calculate fold change
VP_A24B_no0$"Log2FC" <- VP_A24B_no0$'CRE+/control avg' - VP_A24B_no0$'CRE-/control avg'

#save df 
write_csv(VP_A24B_no0, "VP_csv/A24B_no0_Volcano_plot.csv")

## permFDP ####

VP_A24B_no0 <- read_csv("VP_csv/A24B_no0_Volcano_plot.csv")

pVals = VP_A24B_no0$pvalue
threshold = 0.05
myDesign = c(1,1,1,1,2,2,2,2,2,2,2)
intOnly = VP_A24B_no0 %>% select(-c("Lipid species","pvalue","CRE-/control avg","CRE-/control sem","CRE+/control avg","CRE+/control sem","Log2FC"))
intOnly <- mutate_all(intOnly, function(x) as.numeric(as.character(x))) 
corrThreshold_500 = permFDP::permFDP.adjust.threshold(pVals, threshold, myDesign, intOnly, 500)
corrThreshold_1000 = permFDP::permFDP.adjust.threshold(pVals, threshold, myDesign, intOnly, 1000)
corrThreshold_3000 = permFDP::permFDP.adjust.threshold(pVals, threshold, myDesign, intOnly, 3000)
corrThreshold_10000 = permFDP::permFDP.adjust.threshold(pVals, threshold, myDesign, intOnly, 10000)

print(corrThreshold_3000, digits = 20)
FDPthreshold[7,2] <- corrThreshold_3000
#corrThreshold_3000 = 0.011484808231057349465
## EnhancedVolcano ####

pdf("FinalVP/VP_A24B_no0.pdf", width = 1.75, height = 1.24)

# Lipid species, p values, log2 FCs
toptable <- VP_A24B_no0[, c(1,ncol(VP_A24B_no0)-5,ncol(VP_A24B_no0))]
lab = toptable$"Lipid species"
x = "Log2FC"
y = "pvalue"
pCutoff = corrThreshold_3000
FCcutoff = 0.5
#title = "PLP brain samples"
#subtitle = bquote(italic('Week 24 with no 0 values, 3000 iterations, 11 mice'))

keyvals.colour <- ifelse(
  toptable$Log2FC > 0.5 & toptable$pvalue < pCutoff, '#FF00FF',
  ifelse(toptable$Log2FC <(-0.5) & toptable$pvalue < pCutoff, '#00FFFF',
         '#808080'))
# Gray - #A8B6CC - #DAECCF
# Magenta - #F05039 - #F32A50
# Cyan - #0090FF - #0882E0

keyvals.colour[is.na(keyvals.colour)] <- '#808080'
names(keyvals.colour)[keyvals.colour == '#FF00FF'] <- 'High FC & sig'
names(keyvals.colour)[keyvals.colour == '#808080'] <- 'NS'
names(keyvals.colour)[keyvals.colour == '#00FFFF'] <- 'Low FC & sig'

EnhancedVolcano(toptable, lab, x, y, selectLab = rownames(toptable)[which(names(keyvals.colour) %in% c('High', 'Low'))],
                xlab = bquote(~Log[2] ~ "Fold Change"),legendPosition = 'none',
                ylab = bquote(~-Log[10] ~ italic(p) ~ Value), colCustom = keyvals.colour,
                title = NULL, subtitle = NULL, pCutoff = pCutoff, pCutoffCol = y, FCcutoff = FCcutoff,
                drawConnectors = FALSE, xlim = c(min(toptable[[x]], na.rm = TRUE) - 0.2, max(toptable[[x]], na.rm = TRUE) + 0.2),
                ylim = c(0, max(-log10(toptable[[y]]), na.rm = TRUE) + 0.5), colAlpha = 1,
                cutoffLineWidth = 0.1, pointSize = 0.5,axisLabSize = 5, captionLabSize = 3, borderWidth = 0.1,
                vlineWidth = 0.1, hlineWidth = 0.1, caption = NULL
) + theme(axis.text.x = element_text(face="plain", color="black", size=5, angle=0),
          axis.text.y = element_text(face="plain", color="black", size=5, angle=0),
          axis.ticks.length=unit(.05, "cm"), axis.ticks=element_line(linewidth=0.2),
          axis.title.x = element_text(family = "sans", size = 5, margin=margin(1,0,0,0)), 
          axis.title.y = element_text(family = "sans", size = 5, margin=margin(0,1,0,0)),
          panel.grid.major = element_line(linewidth = 0.5), 
          panel.grid.minor = element_line(linewidth = 0.3),
          plot.margin = margin(4,4,4,4)
)

dev.off()


# PLP W24 Spinal ####

A24S <- read_csv("Final_24S_All_noCE.csv", col_names = TRUE, name_repair = unique)
A24S <- A24S %>% filter(grepl('wk 24', ...1)) %>% filter(grepl('Control', CRE.and.Treatment))
A24S_no0 <- A24S %>% select(-"Sample.number", -"...1")

# For no 0 df, drop PC(44:7), SM(22:0), ePC(40:5), PI(40:8), PS(42:11) and PS(42:6)
# https://stackoverflow.com/questions/72096913/remove-columns-that-have-one-zero-value
print(select(A24S_no0, where(~any(. == 0))))
A24S_no0 <- Filter(function(x) !any(x %in% 0), A24S_no0) 
log_A24S_no0 = log2(A24S_no0[,-ncol(A24S_no0)]) 
log_A24S_no0 <- cbind(log_A24S_no0, 'CRE.and.Treatment' = A24S$CRE.and.Treatment)

# Fix column names & remove extra rows
ttest_A24S_no0 <- transpose(log_A24S_no0, keep.names = 'Lipid species')
ttest_A24S_no0 <- ttest_A24S_no0[-nrow(ttest_A24S_no0),]

samples <- A24S$Sample.number
names <- c('Lipid species', samples)
colnames(ttest_A24S_no0) = c(names)

# Get p values from t test 
ttest_A24S_no0$"pvalue" <- unlist(lapply(1:nrow(ttest_A24S_no0), function(i) t.test((as.numeric(ttest_A24S_no0[i,2:4])), as.numeric(ttest_A24S_no0[i,5:11]))[c("p.value")]))

# calculate average values & sem for each species using CRE and treatment as factor
mean.sem_A24S_no0 <- data.table(log_A24S_no0)[, lapply(.SD, mean.sem), by=CRE.and.Treatment]
mean.sem_A24S_no0 <- transpose(mean.sem_A24S_no0[,2:(ncol(mean.sem_A24S_no0))], keep.names = "Lipid species")
# Make sure the order of columns is correct, some df will have experimental first
CRE_factor <- c("CRE-/control avg","CRE-/control sem","CRE+/control avg","CRE+/control sem")
names <- c('Lipid species', CRE_factor)
colnames(mean.sem_A24S_no0) = c(names)

# add means to main df
#VP_A24S_no0 <- cbind(ttest_A24S_no0, mean.sem_A24S_no0)
VP_A24S_no0 <- merge(ttest_A24S_no0, mean.sem_A24S_no0, by = 'Lipid species')
# Calculate fold change
VP_A24S_no0$"Log2FC" <- VP_A24S_no0$'CRE+/control avg' - VP_A24S_no0$'CRE-/control avg'

#save df 
write_csv(VP_A24S_no0, "VP_csv/A24S_no0_Volcano_plot.csv")

## permFDP ####

VP_A24S_no0 <- read_csv("VP_csv/A24S_no0_Volcano_plot.csv")

pVals = VP_A24S_no0$pvalue
threshold = 0.05
myDesign = c(1,1,1,2,2,2,2,2,2,2)
intOnly = VP_A24S_no0 %>% select(-c("Lipid species","pvalue","CRE-/control avg","CRE-/control sem","CRE+/control avg","CRE+/control sem","Log2FC"))
intOnly <- mutate_all(intOnly, function(x) as.numeric(as.character(x))) 
corrThreshold_500 = permFDP::permFDP.adjust.threshold(pVals, threshold, myDesign, intOnly, 500)
corrThreshold_1000 = permFDP::permFDP.adjust.threshold(pVals, threshold, myDesign, intOnly, 1000)
corrThreshold_3000 = permFDP::permFDP.adjust.threshold(pVals, threshold, myDesign, intOnly, 3000)
corrThreshold_10000 = permFDP::permFDP.adjust.threshold(pVals, threshold, myDesign, intOnly, 10000)

print(corrThreshold_3000, digits = 20)
FDPthreshold[8,2] <- corrThreshold_3000
#corrThreshold_3000 = 0.011659899720548269966
## EnhancedVolcano ####

pdf("FinalVP/VP_A24S_no0.pdf", width = 1.75, height = 1.24)

# Lipid species, p values, log2 FCs
toptable <- VP_A24S_no0[, c(1,ncol(VP_A24S_no0)-5,ncol(VP_A24S_no0))]
lab = toptable$"Lipid species"
x = "Log2FC"
y = "pvalue"
pCutoff = corrThreshold_3000
FCcutoff = 0.5
#title = "PLP spinal cord samples"
#subtitle = bquote(italic('Week 24 with no 0 values, 3000 iterations, 10 mice'))

keyvals.colour <- ifelse(
  toptable$Log2FC > 0.5 & toptable$pvalue < pCutoff, '#FF00FF',
  ifelse(toptable$Log2FC <(-0.5) & toptable$pvalue < pCutoff, '#00FFFF',
         '#808080'))

keyvals.colour[is.na(keyvals.colour)] <- '#808080'
names(keyvals.colour)[keyvals.colour == '#FF00FF'] <- 'High FC & sig'
names(keyvals.colour)[keyvals.colour == '#808080'] <- 'NS'
names(keyvals.colour)[keyvals.colour == '#00FFFF'] <- 'Low FC & sig'

EnhancedVolcano(toptable, lab, x, y, selectLab = rownames(toptable)[which(names(keyvals.colour) %in% c('High', 'Low'))],
                xlab = bquote(~Log[2] ~ "Fold Change"),legendPosition = 'none',
                ylab = bquote(~-Log[10] ~ italic(p) ~ Value), colCustom = keyvals.colour,
                title = NULL, subtitle = NULL, pCutoff = pCutoff, pCutoffCol = y, FCcutoff = FCcutoff,
                drawConnectors = FALSE, xlim = c(min(toptable[[x]], na.rm = TRUE) - 0.2, max(toptable[[x]], na.rm = TRUE) + 0.2),
                ylim = c(0, max(-log10(toptable[[y]]), na.rm = TRUE) + 0.5), colAlpha = 1,
                cutoffLineWidth = 0.1, pointSize = 0.5,axisLabSize = 5, captionLabSize = 3, borderWidth = 0.1,
                vlineWidth = 0.1, hlineWidth = 0.1, caption = NULL
) + theme(axis.text.x = element_text(face="plain", color="black", size=5, angle=0),
          axis.text.y = element_text(face="plain", color="black", size=5, angle=0),
          axis.ticks.length=unit(.05, "cm"), axis.ticks=element_line(linewidth=0.2),
          axis.title.x = element_text(family = "sans", size = 5, margin=margin(1,0,0,0)), 
          axis.title.y = element_text(family = "sans", size = 5, margin=margin(0,1,0,0)),
          panel.grid.major = element_line(linewidth = 0.5), 
          panel.grid.minor = element_line(linewidth = 0.3),
          plot.margin = margin(4,4,4,4)
)

dev.off()

write_csv(FDPthreshold,"FDP_Threshold.csv")
FDPthreshold <- read_csv("FDP_Threshold.csv")

# PLP W12 Brain + SobAM2 ####

A12B_SobAM2_CrePos <- A6_18_B %>% filter(grepl('wk 12', ...1)) %>% filter(!grepl('-', CRE.and.Treatment))
A12B_SobAM2_CrePos_no0 <- A12B_SobAM2_CrePos %>% select(-"Sample.number", -"...1")

# For no 0 df, drop PE(44:9), PI(38:3) and PA(38:3)
# https://stackoverflow.com/questions/72096913/remove-columns-that-have-one-zero-value
print(select(A12B_SobAM2_CrePos_no0, where(~any(. == 0))))
A12B_SobAM2_CrePos_no0 <- Filter(function(x) !any(x %in% 0), A12B_SobAM2_CrePos_no0) 
log_A12B_SobAM2_CrePos_no0 = log2(A12B_SobAM2_CrePos_no0[,-ncol(A12B_SobAM2_CrePos_no0)]) 
log_A12B_SobAM2_CrePos_no0 <- cbind(log_A12B_SobAM2_CrePos_no0, 'CRE.and.Treatment' = A12B_SobAM2_CrePos$CRE.and.Treatment)

# Fix column names & remove extra rows
ttest_A12B_SobAM2_CrePos_no0 <- transpose(log_A12B_SobAM2_CrePos_no0, keep.names = 'Lipid species')
ttest_A12B_SobAM2_CrePos_no0 <- ttest_A12B_SobAM2_CrePos_no0[-nrow(ttest_A12B_SobAM2_CrePos_no0),]

samples <- A12B_SobAM2_CrePos$Sample.number
names <- c('Lipid species', samples)
colnames(ttest_A12B_SobAM2_CrePos_no0) = c(names)

# Get p values from t test 
ttest_A12B_SobAM2_CrePos_no0$"pvalue" <- unlist(lapply(1:nrow(ttest_A12B_SobAM2_CrePos_no0), function(i) t.test((as.numeric(ttest_A12B_SobAM2_CrePos_no0[i,2:9])), as.numeric(ttest_A12B_SobAM2_CrePos_no0[i,10:19]))[c("p.value")]))

# calculate average values & sem for each species using CRE and treatment as factor
mean.sem_A12B_SobAM2_CrePos_no0 <- data.table(log_A12B_SobAM2_CrePos_no0)[, lapply(.SD, mean.sem), by=CRE.and.Treatment]
mean.sem_A12B_SobAM2_CrePos_no0 <- transpose(mean.sem_A12B_SobAM2_CrePos_no0[,2:(ncol(mean.sem_A12B_SobAM2_CrePos_no0))], keep.names = "Lipid species")
# Make sure the order of columns is correct, some df will have experimental first
CRE_factor <- c("CRE+/control avg","CRE+/control sem","CRE+/SobAM2 avg","CRE+/SobAM2 sem")
names <- c('Lipid species', CRE_factor)
colnames(mean.sem_A12B_SobAM2_CrePos_no0) = c(names)

# add means to main df
#VP_A12B_SobAM2_CrePos_no0 <- cbind(ttest_A12B_SobAM2_CrePos_no0, mean.sem_A12B_SobAM2_CrePos_no0)
VP_A12B_SobAM2_CrePos_no0 <- merge(ttest_A12B_SobAM2_CrePos_no0, mean.sem_A12B_SobAM2_CrePos_no0, by = 'Lipid species')
# Calculate fold change
VP_A12B_SobAM2_CrePos_no0$"Log2FC" <- VP_A12B_SobAM2_CrePos_no0$'CRE+/SobAM2 avg' - VP_A12B_SobAM2_CrePos_no0$'CRE+/control avg'

#save df 
write_csv(VP_A12B_SobAM2_CrePos_no0, "VP_csv/A12B_SobAM2_CrePos_no0_Volcano_plot.csv")

## permFDP ####

VP_A12B_SobAM2_CrePos_no0 <- read_csv("VP_csv/A12B_SobAM2_CrePos_no0_Volcano_plot.csv")

pVals = VP_A12B_SobAM2_CrePos_no0$pvalue
threshold = 0.05
myDesign = c(1,1,1,1,1,1,1,1,2,2,2,2,2,2,2,2,2,2)
intOnly = VP_A12B_SobAM2_CrePos_no0 %>% select(-c("Lipid species","pvalue","CRE+/control avg","CRE+/control sem","CRE+/SobAM2 avg","CRE+/SobAM2 sem","Log2FC"))
intOnly <- mutate_all(intOnly, function(x) as.numeric(as.character(x))) 
corrThreshold_500 = permFDP::permFDP.adjust.threshold(pVals, threshold, myDesign, intOnly, 500)
corrThreshold_1000 = permFDP::permFDP.adjust.threshold(pVals, threshold, myDesign, intOnly, 1000)
corrThreshold_3000 = permFDP::permFDP.adjust.threshold(pVals, threshold, myDesign, intOnly, 3000)
corrThreshold_10000 = permFDP::permFDP.adjust.threshold(pVals, threshold, myDesign, intOnly, 10000)

print(corrThreshold_3000, digits = 20)
FDPthreshold[9,2] <- corrThreshold_3000
# corrThreshold_3000 = 0.011067124829758429733
## EnhancedVolcano ####

pdf("FinalVP/VP_A12B_SobAM2_CrePos_no0.pdf", width = 1.75, height = 1.24)

# Lipid species, p values, log2 FCs
toptable <- VP_A12B_SobAM2_CrePos_no0[, c(1,ncol(VP_A12B_SobAM2_CrePos_no0)-5,ncol(VP_A12B_SobAM2_CrePos_no0))]
lab = toptable$"Lipid species"
x = "Log2FC"
y = "pvalue"
pCutoff = corrThreshold_3000
FCcutoff = 0.5

keyvals.colour <- ifelse(
  toptable$Log2FC > 0.5 & toptable$pvalue < pCutoff, '#FF00FF',
  ifelse(toptable$Log2FC <(-0.5) & toptable$pvalue < pCutoff, '#00FFFF',
         '#808080'))
# Gray - #A8B6CC - #DAECCF
# Magenta - #F05039 - #F32A50
# Cyan - #0090FF - #0882E0

keyvals.colour[is.na(keyvals.colour)] <- '#808080'
names(keyvals.colour)[keyvals.colour == '#FF00FF'] <- 'High FC & sig'
names(keyvals.colour)[keyvals.colour == '#808080'] <- 'NS'
names(keyvals.colour)[keyvals.colour == '#00FFFF'] <- 'Low FC & sig'

EnhancedVolcano(toptable, lab, x, y, selectLab = rownames(toptable)[which(names(keyvals.colour) %in% c('High', 'Low'))],
                xlab = bquote(~Log[2] ~ "Fold Change"),legendPosition = 'none',
                ylab = bquote(~-Log[10] ~ italic(p) ~ Value), colCustom = keyvals.colour,
                title = NULL, subtitle = NULL, pCutoff = pCutoff, pCutoffCol = y, FCcutoff = FCcutoff,
                drawConnectors = FALSE, xlim = c(min(toptable[[x]], na.rm = TRUE) - 0.2, max(toptable[[x]], na.rm = TRUE) + 0.2),
                ylim = c(0, max(-log10(toptable[[y]]), na.rm = TRUE) + 0.5), colAlpha = 1,
                cutoffLineWidth = 0.1, pointSize = 0.5,axisLabSize = 5, captionLabSize = 3, borderWidth = 0.1,
                vlineWidth = 0.1, hlineWidth = 0.1, caption = NULL, legendLabSize = 3, legendIconSize =1
) + theme(axis.text.x = element_text(face="plain", color="black", size=5, angle=0),
          axis.text.y = element_text(face="plain", color="black", size=5, angle=0),
          axis.ticks.length=unit(.05, "cm"), axis.ticks=element_line(linewidth=0.2),
          axis.title.x = element_text(family = "sans", size = 5, margin=margin(1,0,0,0)), 
          axis.title.y = element_text(family = "sans", size = 5, margin=margin(0,1,0,0)),
          panel.grid.major = element_line(linewidth = 0.5), 
          panel.grid.minor = element_line(linewidth = 0.3),
          plot.margin = margin(4,4,4,4),
          legend.spacing.x = unit(0.5, "cm"),
          legend.spacing.y = unit(0.5, "cm")
)

dev.off()

# PLP W12 Spinal + SobAM2 ####

A12S_SobAM2_CrePos <- A6_18_S %>% filter(grepl('wk 12', ...1)) %>% filter(!grepl('-', CRE.and.Treatment))
A12S_SobAM2_CrePos_no0 <- A12S_SobAM2_CrePos %>% select(-"Sample.number", -"...1")

# For no 0 df, drop PC(40:3), ePC(40:5) and PE(40:5)
# https://stackoverflow.com/questions/72096913/remove-columns-that-have-one-zero-value
print(select(A12S_SobAM2_CrePos_no0, where(~any(. == 0))))
A12S_SobAM2_CrePos_no0 <- Filter(function(x) !any(x %in% 0), A12S_SobAM2_CrePos_no0) 
log_A12S_SobAM2_CrePos_no0 = log2(A12S_SobAM2_CrePos_no0[,-ncol(A12S_SobAM2_CrePos_no0)]) 
log_A12S_SobAM2_CrePos_no0 <- cbind(log_A12S_SobAM2_CrePos_no0, 'CRE.and.Treatment' = A12S_SobAM2_CrePos$CRE.and.Treatment)

# Fix column names & remove extra rows
ttest_A12S_SobAM2_CrePos_no0 <- transpose(log_A12S_SobAM2_CrePos_no0, keep.names = 'Lipid species')
ttest_A12S_SobAM2_CrePos_no0 <- ttest_A12S_SobAM2_CrePos_no0[-nrow(ttest_A12S_SobAM2_CrePos_no0),]

samples <- A12S_SobAM2_CrePos$Sample.number
names <- c('Lipid species', samples)
colnames(ttest_A12S_SobAM2_CrePos_no0) = c(names)

# Get p values from t test 
ttest_A12S_SobAM2_CrePos_no0$"pvalue" <- unlist(lapply(1:nrow(ttest_A12S_SobAM2_CrePos_no0), function(i) t.test((as.numeric(ttest_A12S_SobAM2_CrePos_no0[i,2:9])), as.numeric(ttest_A12S_SobAM2_CrePos_no0[i,10:19]))[c("p.value")]))

# calculate average values & sem for each species using CRE and treatment as factor
mean.sem_A12S_SobAM2_CrePos_no0 <- data.table(log_A12S_SobAM2_CrePos_no0)[, lapply(.SD, mean.sem), by=CRE.and.Treatment]
mean.sem_A12S_SobAM2_CrePos_no0 <- transpose(mean.sem_A12S_SobAM2_CrePos_no0[,2:(ncol(mean.sem_A12S_SobAM2_CrePos_no0))], keep.names = "Lipid species")
# Make sure the order of columns is correct, some df will have experimental first
CRE_factor <- c("CRE+/control avg","CRE+/control sem","CRE+/SobAM2 avg","CRE+/SobAM2 sem")
names <- c('Lipid species', CRE_factor)
colnames(mean.sem_A12S_SobAM2_CrePos_no0) = c(names)

# add means to main df
#VP_A12S_SobAM2_CrePos_no0 <- cbind(ttest_A12S_SobAM2_CrePos_no0, mean.sem_A12S_SobAM2_CrePos_no0)
VP_A12S_SobAM2_CrePos_no0 <- merge(ttest_A12S_SobAM2_CrePos_no0, mean.sem_A12S_SobAM2_CrePos_no0, by = 'Lipid species')
# Calculate fold change
VP_A12S_SobAM2_CrePos_no0$"Log2FC" <- VP_A12S_SobAM2_CrePos_no0$'CRE+/SobAM2 avg' - VP_A12S_SobAM2_CrePos_no0$'CRE+/control avg'

#save df 
write_csv(VP_A12S_SobAM2_CrePos_no0, "VP_csv/A12S_SobAM2_CrePos_no0_Volcano_plot.csv")

## permFDP ####

VP_A12S_SobAM2_CrePos_no0 <- read_csv("VP_csv/A12S_SobAM2_CrePos_no0_Volcano_plot.csv")

pVals = VP_A12S_SobAM2_CrePos_no0$pvalue
threshold = 0.05
myDesign = c(1,1,1,1,1,1,1,1,2,2,2,2,2,2,2,2,2,2)
intOnly = VP_A12S_SobAM2_CrePos_no0 %>% select(-c("Lipid species","pvalue","CRE+/control avg","CRE+/control sem","CRE+/SobAM2 avg","CRE+/SobAM2 sem","Log2FC"))
intOnly <- mutate_all(intOnly, function(x) as.numeric(as.character(x))) 
corrThreshold_500 = permFDP::permFDP.adjust.threshold(pVals, threshold, myDesign, intOnly, 500)
corrThreshold_1000 = permFDP::permFDP.adjust.threshold(pVals, threshold, myDesign, intOnly, 1000)
corrThreshold_3000 = permFDP::permFDP.adjust.threshold(pVals, threshold, myDesign, intOnly, 3000)
corrThreshold_10000 = permFDP::permFDP.adjust.threshold(pVals, threshold, myDesign, intOnly, 10000)

print(corrThreshold_3000, digits = 20)
FDPthreshold[10,2] <- corrThreshold_3000
# corrThreshold_3000 = 0.002960860240916025031
## EnhancedVolcano ####

pdf("FinalVP/VP_A12S_SobAM2_CrePos_no0.pdf", width = 1.75, height = 1.24)

# Lipid species, p values, log2 FCs
toptable <- VP_A12S_SobAM2_CrePos_no0[, c(1,ncol(VP_A12S_SobAM2_CrePos_no0)-5,ncol(VP_A12S_SobAM2_CrePos_no0))]
lab = toptable$"Lipid species"
x = "Log2FC"
y = "pvalue"
pCutoff = corrThreshold_3000
FCcutoff = 0.5

keyvals.colour <- ifelse(
  toptable$Log2FC > 0.5 & toptable$pvalue < pCutoff, '#FF00FF',
  ifelse(toptable$Log2FC <(-0.5) & toptable$pvalue < pCutoff, '#00FFFF',
         '#808080'))
# Gray - #A8B6CC - #DAECCF
# Magenta - #F05039 - #F32A50
# Cyan - #0090FF - #0882E0

keyvals.colour[is.na(keyvals.colour)] <- '#808080'
names(keyvals.colour)[keyvals.colour == '#FF00FF'] <- 'High FC & sig'
names(keyvals.colour)[keyvals.colour == '#808080'] <- 'NS'
names(keyvals.colour)[keyvals.colour == '#00FFFF'] <- 'Low FC & sig'

EnhancedVolcano(toptable, lab, x, y, selectLab = rownames(toptable)[which(names(keyvals.colour) %in% c('High', 'Low'))],
                xlab = bquote(~Log[2] ~ "Fold Change"),legendPosition = 'none',
                ylab = bquote(~-Log[10] ~ italic(p) ~ Value), colCustom = keyvals.colour,
                title = NULL, subtitle = NULL, pCutoff = pCutoff, pCutoffCol = y, FCcutoff = FCcutoff,
                drawConnectors = FALSE, xlim = c(min(toptable[[x]], na.rm = TRUE) - 0.2, max(toptable[[x]], na.rm = TRUE) + 0.2),
                ylim = c(0, max(-log10(toptable[[y]]), na.rm = TRUE) + 0.5), colAlpha = 1,
                cutoffLineWidth = 0.1, pointSize = 0.5,axisLabSize = 5, captionLabSize = 3, borderWidth = 0.1,
                vlineWidth = 0.1, hlineWidth = 0.1, caption = NULL, legendLabSize = 3, legendIconSize =1
) + theme(axis.text.x = element_text(face="plain", color="black", size=5, angle=0),
          axis.text.y = element_text(face="plain", color="black", size=5, angle=0),
          axis.ticks.length=unit(.05, "cm"), axis.ticks=element_line(linewidth=0.2),
          axis.title.x = element_text(family = "sans", size = 5, margin=margin(1,0,0,0)), 
          axis.title.y = element_text(family = "sans", size = 5, margin=margin(0,1,0,0)),
          panel.grid.major = element_line(linewidth = 0.5), 
          panel.grid.minor = element_line(linewidth = 0.3),
          plot.margin = margin(4,4,4,4),
          legend.spacing.x = unit(0.5, "cm"),
          legend.spacing.y = unit(0.5, "cm")
)

dev.off()

# PLP W12 Brain - SobAM2 ####

A12B_SobAM2_CreNeg <- A6_18_B %>% filter(grepl('wk 12', ...1)) %>% filter(grepl('-', CRE.and.Treatment))
A12B_SobAM2_CreNeg_no0 <- A12B_SobAM2_CreNeg %>% select(-"Sample.number", -"...1")

# For no 0 df, drop ePC(38:6), PS(32:1), ePS(38:4) and PA(38:3)
# https://stackoverflow.com/questions/72096913/remove-columns-that-have-one-zero-value
print(select(A12B_SobAM2_CreNeg_no0, where(~any(. == 0))))
A12B_SobAM2_CreNeg_no0 <- Filter(function(x) !any(x %in% 0), A12B_SobAM2_CreNeg_no0) 
log_A12B_SobAM2_CreNeg_no0 = log2(A12B_SobAM2_CreNeg_no0[,-ncol(A12B_SobAM2_CreNeg_no0)]) 
log_A12B_SobAM2_CreNeg_no0 <- cbind(log_A12B_SobAM2_CreNeg_no0, 'CRE.and.Treatment' = A12B_SobAM2_CreNeg$CRE.and.Treatment)

# Fix column names & remove extra rows
ttest_A12B_SobAM2_CreNeg_no0 <- transpose(log_A12B_SobAM2_CreNeg_no0, keep.names = 'Lipid species')
ttest_A12B_SobAM2_CreNeg_no0 <- ttest_A12B_SobAM2_CreNeg_no0[-nrow(ttest_A12B_SobAM2_CreNeg_no0),]

samples <- A12B_SobAM2_CreNeg$Sample.number
names <- c('Lipid species', samples)
colnames(ttest_A12B_SobAM2_CreNeg_no0) = c(names)

# Get p values from t test 
ttest_A12B_SobAM2_CreNeg_no0$"pvalue" <- unlist(lapply(1:nrow(ttest_A12B_SobAM2_CreNeg_no0), function(i) t.test((as.numeric(ttest_A12B_SobAM2_CreNeg_no0[i,2:6])), as.numeric(ttest_A12B_SobAM2_CreNeg_no0[i,7:12]))[c("p.value")]))

# calculate average values & sem for each species using CRE and treatment as factor
mean.sem_A12B_SobAM2_CreNeg_no0 <- data.table(log_A12B_SobAM2_CreNeg_no0)[, lapply(.SD, mean.sem), by=CRE.and.Treatment]
mean.sem_A12B_SobAM2_CreNeg_no0 <- transpose(mean.sem_A12B_SobAM2_CreNeg_no0[,2:(ncol(mean.sem_A12B_SobAM2_CreNeg_no0))], keep.names = "Lipid species")
# Make sure the order of columns is correct, some df will have experimental first
CRE_factor <- c("CRE-/control avg","CRE-/control sem","CRE-/SobAM2 avg","CRE-/SobAM2 sem")
names <- c('Lipid species', CRE_factor)
colnames(mean.sem_A12B_SobAM2_CreNeg_no0) = c(names)

# add means to main df
#VP_A12B_SobAM2_CreNeg_no0 <- cbind(ttest_A12B_SobAM2_CreNeg_no0, mean.sem_A12B_SobAM2_CreNeg_no0)
VP_A12B_SobAM2_CreNeg_no0 <- merge(ttest_A12B_SobAM2_CreNeg_no0, mean.sem_A12B_SobAM2_CreNeg_no0, by = 'Lipid species')
# Calculate fold change
VP_A12B_SobAM2_CreNeg_no0$"Log2FC" <- VP_A12B_SobAM2_CreNeg_no0$'CRE-/SobAM2 avg' - VP_A12B_SobAM2_CreNeg_no0$'CRE-/control avg'

#save df 
write_csv(VP_A12B_SobAM2_CreNeg_no0, "VP_csv/A12B_SobAM2_CreNeg_no0_Volcano_plot.csv")

## permFDP ####

VP_A12B_SobAM2_CreNeg_no0 <- read_csv("VP_csv/A12B_SobAM2_CreNeg_no0_Volcano_plot.csv")

pVals = VP_A12B_SobAM2_CreNeg_no0$pvalue
threshold = 0.05
myDesign = c(1,1,1,1,1,2,2,2,2,2,2)
intOnly = VP_A12B_SobAM2_CreNeg_no0 %>% select(-c("Lipid species","pvalue","CRE-/control avg","CRE-/control sem","CRE-/SobAM2 avg","CRE-/SobAM2 sem","Log2FC"))
intOnly <- mutate_all(intOnly, function(x) as.numeric(as.character(x))) 
corrThreshold_500 = permFDP::permFDP.adjust.threshold(pVals, threshold, myDesign, intOnly, 500)
corrThreshold_1000 = permFDP::permFDP.adjust.threshold(pVals, threshold, myDesign, intOnly, 1000)
corrThreshold_3000 = permFDP::permFDP.adjust.threshold(pVals, threshold, myDesign, intOnly, 3000)
corrThreshold_10000 = permFDP::permFDP.adjust.threshold(pVals, threshold, myDesign, intOnly, 10000)

print(corrThreshold_3000, digits = 20)
FDPthreshold[11,2] <- corrThreshold_3000
# corrThreshold_3000 = 0.0020927663994541648683
## EnhancedVolcano ####

pdf("FinalVP/VP_A12B_SobAM2_CreNeg_no0.pdf", width = 1.75, height = 1.24)

# Lipid species, p values, log2 FCs
toptable <- VP_A12B_SobAM2_CreNeg_no0[, c(1,ncol(VP_A12B_SobAM2_CreNeg_no0)-5,ncol(VP_A12B_SobAM2_CreNeg_no0))]
lab = toptable$"Lipid species"
x = "Log2FC"
y = "pvalue"
pCutoff = corrThreshold_3000
FCcutoff = 0.5

keyvals.colour <- ifelse(
  toptable$Log2FC > 0.5 & toptable$pvalue < pCutoff, '#FF00FF',
  ifelse(toptable$Log2FC <(-0.5) & toptable$pvalue < pCutoff, '#00FFFF',
         '#808080'))
# Gray - #A8B6CC - #DAECCF
# Magenta - #F05039 - #F32A50
# Cyan - #0090FF - #0882E0

keyvals.colour[is.na(keyvals.colour)] <- '#808080'
names(keyvals.colour)[keyvals.colour == '#FF00FF'] <- 'High FC & sig'
names(keyvals.colour)[keyvals.colour == '#808080'] <- 'NS'
names(keyvals.colour)[keyvals.colour == '#00FFFF'] <- 'Low FC & sig'

EnhancedVolcano(toptable, lab, x, y, selectLab = rownames(toptable)[which(names(keyvals.colour) %in% c('High', 'Low'))],
                xlab = bquote(~Log[2] ~ "Fold Change"),legendPosition = 'none',
                ylab = bquote(~-Log[10] ~ italic(p) ~ Value), colCustom = keyvals.colour,
                title = NULL, subtitle = NULL, pCutoff = pCutoff, pCutoffCol = y, FCcutoff = FCcutoff,
                drawConnectors = FALSE, xlim = c(min(toptable[[x]], na.rm = TRUE) - 0.2, max(toptable[[x]], na.rm = TRUE) + 0.2),
                ylim = c(0, max(-log10(toptable[[y]]), na.rm = TRUE) + 0.5), colAlpha = 1,
                cutoffLineWidth = 0.1, pointSize = 0.5,axisLabSize = 5, captionLabSize = 3, borderWidth = 0.1,
                vlineWidth = 0.1, hlineWidth = 0.1, caption = NULL, legendLabSize = 3, legendIconSize =1
) + theme(axis.text.x = element_text(face="plain", color="black", size=5, angle=0),
          axis.text.y = element_text(face="plain", color="black", size=5, angle=0),
          axis.ticks.length=unit(.05, "cm"), axis.ticks=element_line(linewidth=0.2),
          axis.title.x = element_text(family = "sans", size = 5, margin=margin(1,0,0,0)), 
          axis.title.y = element_text(family = "sans", size = 5, margin=margin(0,1,0,0)),
          panel.grid.major = element_line(linewidth = 0.5), 
          panel.grid.minor = element_line(linewidth = 0.3),
          plot.margin = margin(4,4,4,4),
          legend.spacing.x = unit(0.5, "cm"),
          legend.spacing.y = unit(0.5, "cm")
)

dev.off()

# PLP W12 Spinal - SobAM2 ####

A12S_SobAM2_CreNeg <- A6_18_S %>% filter(grepl('wk 12', ...1)) %>% filter(grepl('-', CRE.and.Treatment))
A12S_SobAM2_CreNeg_no0 <- A12S_SobAM2_CreNeg %>% select(-"Sample.number", -"...1")

# For no 0 df, drop PI(38:3) and PS(32:1)
# https://stackoverflow.com/questions/72096913/remove-columns-that-have-one-zero-value
print(select(A12S_SobAM2_CreNeg_no0, where(~any(. == 0))))
A12S_SobAM2_CreNeg_no0 <- Filter(function(x) !any(x %in% 0), A12S_SobAM2_CreNeg_no0) 
log_A12S_SobAM2_CreNeg_no0 = log2(A12S_SobAM2_CreNeg_no0[,-ncol(A12S_SobAM2_CreNeg_no0)]) 
log_A12S_SobAM2_CreNeg_no0 <- cbind(log_A12S_SobAM2_CreNeg_no0, 'CRE.and.Treatment' = A12S_SobAM2_CreNeg$CRE.and.Treatment)

# Fix column names & remove extra rows
ttest_A12S_SobAM2_CreNeg_no0 <- transpose(log_A12S_SobAM2_CreNeg_no0, keep.names = 'Lipid species')
ttest_A12S_SobAM2_CreNeg_no0 <- ttest_A12S_SobAM2_CreNeg_no0[-nrow(ttest_A12S_SobAM2_CreNeg_no0),]

samples <- A12S_SobAM2_CreNeg$Sample.number
names <- c('Lipid species', samples)
colnames(ttest_A12S_SobAM2_CreNeg_no0) = c(names)

# Get p values from t test 
ttest_A12S_SobAM2_CreNeg_no0$"pvalue" <- unlist(lapply(1:nrow(ttest_A12S_SobAM2_CreNeg_no0), function(i) t.test((as.numeric(ttest_A12S_SobAM2_CreNeg_no0[i,2:6])), as.numeric(ttest_A12S_SobAM2_CreNeg_no0[i,7:12]))[c("p.value")]))

# calculate average values & sem for each species using CRE and treatment as factor
mean.sem_A12S_SobAM2_CreNeg_no0 <- data.table(log_A12S_SobAM2_CreNeg_no0)[, lapply(.SD, mean.sem), by=CRE.and.Treatment]
mean.sem_A12S_SobAM2_CreNeg_no0 <- transpose(mean.sem_A12S_SobAM2_CreNeg_no0[,2:(ncol(mean.sem_A12S_SobAM2_CreNeg_no0))], keep.names = "Lipid species")
# Make sure the order of columns is correct, some df will have experimental first
CRE_factor <- c("CRE-/control avg","CRE-/control sem","CRE-/SobAM2 avg","CRE-/SobAM2 sem")
names <- c('Lipid species', CRE_factor)
colnames(mean.sem_A12S_SobAM2_CreNeg_no0) = c(names)

# add means to main df
#VP_A12S_SobAM2_CreNeg_no0 <- cbind(ttest_A12S_SobAM2_CreNeg_no0, mean.sem_A12S_SobAM2_CreNeg_no0)
VP_A12S_SobAM2_CreNeg_no0 <- merge(ttest_A12S_SobAM2_CreNeg_no0, mean.sem_A12S_SobAM2_CreNeg_no0, by = 'Lipid species')
# Calculate fold change
VP_A12S_SobAM2_CreNeg_no0$"Log2FC" <- VP_A12S_SobAM2_CreNeg_no0$'CRE-/SobAM2 avg' - VP_A12S_SobAM2_CreNeg_no0$'CRE-/control avg'

#save df 
write_csv(VP_A12S_SobAM2_CreNeg_no0, "VP_csv/A12S_SobAM2_CreNeg_no0_Volcano_plot.csv")

## permFDP ####

VP_A12S_SobAM2_CreNeg_no0 <- read_csv("VP_csv/A12S_SobAM2_CreNeg_no0_Volcano_plot.csv")

pVals = VP_A12S_SobAM2_CreNeg_no0$pvalue
threshold = 0.05
myDesign = c(1,1,1,1,1,2,2,2,2,2,2)
intOnly = VP_A12S_SobAM2_CreNeg_no0 %>% select(-c("Lipid species","pvalue","CRE-/control avg","CRE-/control sem","CRE-/SobAM2 avg","CRE-/SobAM2 sem","Log2FC"))
intOnly <- mutate_all(intOnly, function(x) as.numeric(as.character(x))) 
corrThreshold_500 = permFDP::permFDP.adjust.threshold(pVals, threshold, myDesign, intOnly, 500)
corrThreshold_1000 = permFDP::permFDP.adjust.threshold(pVals, threshold, myDesign, intOnly, 1000)
corrThreshold_3000 = permFDP::permFDP.adjust.threshold(pVals, threshold, myDesign, intOnly, 3000)
corrThreshold_10000 = permFDP::permFDP.adjust.threshold(pVals, threshold, myDesign, intOnly, 10000)

print(corrThreshold_3000, digits = 20)
FDPthreshold[12,2] <- corrThreshold_3000
# corrThreshold_3000 = 0.0014826874448812020837
## EnhancedVolcano ####

pdf("FinalVP/VP_A12S_SobAM2_CreNeg_no0.pdf", width = 1.75, height = 1.24)

# Lipid species, p values, log2 FCs
toptable <- VP_A12S_SobAM2_CreNeg_no0[, c(1,ncol(VP_A12S_SobAM2_CreNeg_no0)-5,ncol(VP_A12S_SobAM2_CreNeg_no0))]
lab = toptable$"Lipid species"
x = "Log2FC"
y = "pvalue"
pCutoff = corrThreshold_3000
FCcutoff = 0.5

keyvals.colour <- ifelse(
  toptable$Log2FC > 0.5 & toptable$pvalue < pCutoff, '#FF00FF',
  ifelse(toptable$Log2FC <(-0.5) & toptable$pvalue < pCutoff, '#00FFFF',
         '#808080'))
# Gray - #A8B6CC - #DAECCF
# Magenta - #F05039 - #F32A50
# Cyan - #0090FF - #0882E0

keyvals.colour[is.na(keyvals.colour)] <- '#808080'
names(keyvals.colour)[keyvals.colour == '#FF00FF'] <- 'High FC & sig'
names(keyvals.colour)[keyvals.colour == '#808080'] <- 'NS'
names(keyvals.colour)[keyvals.colour == '#00FFFF'] <- 'Low FC & sig'

EnhancedVolcano(toptable, lab, x, y, selectLab = rownames(toptable)[which(names(keyvals.colour) %in% c('High', 'Low'))],
                xlab = bquote(~Log[2] ~ "Fold Change"),legendPosition = 'none',
                ylab = bquote(~-Log[10] ~ italic(p) ~ Value), colCustom = keyvals.colour,
                title = NULL, subtitle = NULL, pCutoff = pCutoff, pCutoffCol = y, FCcutoff = FCcutoff,
                drawConnectors = FALSE, xlim = c(min(toptable[[x]], na.rm = TRUE) - 0.2, max(toptable[[x]], na.rm = TRUE) + 0.2),
                ylim = c(0, max(-log10(toptable[[y]]), na.rm = TRUE) + 0.5), colAlpha = 1,
                cutoffLineWidth = 0.1, pointSize = 0.5,axisLabSize = 5, captionLabSize = 3, borderWidth = 0.1,
                vlineWidth = 0.1, hlineWidth = 0.1, caption = NULL, legendLabSize = 3, legendIconSize =1
) + theme(axis.text.x = element_text(face="plain", color="black", size=5, angle=0),
          axis.text.y = element_text(face="plain", color="black", size=5, angle=0),
          axis.ticks.length=unit(.05, "cm"), axis.ticks=element_line(linewidth=0.2),
          axis.title.x = element_text(family = "sans", size = 5, margin=margin(1,0,0,0)), 
          axis.title.y = element_text(family = "sans", size = 5, margin=margin(0,1,0,0)),
          panel.grid.major = element_line(linewidth = 0.5), 
          panel.grid.minor = element_line(linewidth = 0.3),
          plot.margin = margin(4,4,4,4),
          legend.spacing.x = unit(0.5, "cm"),
          legend.spacing.y = unit(0.5, "cm")
)

dev.off()

write_csv(FDPthreshold,"FDP_Threshold.csv")
FDPthreshold <- read_csv("FDP_Threshold.csv")

# PLP W18 Brain + SobAM2 ####

A18B_SobAM2_CrePos <- A6_18_B %>% filter(grepl('wk 18', ...1)) %>% filter(!grepl('-', CRE.and.Treatment))
A18B_SobAM2_CrePos_no0 <- A18B_SobAM2_CrePos %>% select(-"Sample.number", -"...1")

# For no 0 df, drop PE(36:3) and PI(38:3)
# https://stackoverflow.com/questions/72096913/remove-columns-that-have-one-zero-value
print(select(A18B_SobAM2_CrePos_no0, where(~any(. == 0))))
A18B_SobAM2_CrePos_no0 <- Filter(function(x) !any(x %in% 0), A18B_SobAM2_CrePos_no0) 
log_A18B_SobAM2_CrePos_no0 = log2(A18B_SobAM2_CrePos_no0[,-ncol(A18B_SobAM2_CrePos_no0)]) 
log_A18B_SobAM2_CrePos_no0 <- cbind(log_A18B_SobAM2_CrePos_no0, 'CRE.and.Treatment' = A18B_SobAM2_CrePos$CRE.and.Treatment)

# Fix column names & remove extra rows
ttest_A18B_SobAM2_CrePos_no0 <- transpose(log_A18B_SobAM2_CrePos_no0, keep.names = 'Lipid species')
ttest_A18B_SobAM2_CrePos_no0 <- ttest_A18B_SobAM2_CrePos_no0[-nrow(ttest_A18B_SobAM2_CrePos_no0),]

samples <- A18B_SobAM2_CrePos$Sample.number
names <- c('Lipid species', samples)
colnames(ttest_A18B_SobAM2_CrePos_no0) = c(names)

# Get p values from t test 
ttest_A18B_SobAM2_CrePos_no0$"pvalue" <- unlist(lapply(1:nrow(ttest_A18B_SobAM2_CrePos_no0), function(i) t.test((as.numeric(ttest_A18B_SobAM2_CrePos_no0[i,2:9])), as.numeric(ttest_A18B_SobAM2_CrePos_no0[i,10:18]))[c("p.value")]))

# calculate average values & sem for each species using CRE and treatment as factor
mean.sem_A18B_SobAM2_CrePos_no0 <- data.table(log_A18B_SobAM2_CrePos_no0)[, lapply(.SD, mean.sem), by=CRE.and.Treatment]
mean.sem_A18B_SobAM2_CrePos_no0 <- transpose(mean.sem_A18B_SobAM2_CrePos_no0[,2:(ncol(mean.sem_A18B_SobAM2_CrePos_no0))], keep.names = "Lipid species")
# Make sure the order of columns is correct, some df will have experimental first
CRE_factor <- c("CRE+/control avg","CRE+/control sem","CRE+/SobAM2 avg","CRE+/SobAM2 sem")
names <- c('Lipid species', CRE_factor)
colnames(mean.sem_A18B_SobAM2_CrePos_no0) = c(names)

# add means to main df
#VP_A18B_SobAM2_CrePos_no0 <- cbind(ttest_A18B_SobAM2_CrePos_no0, mean.sem_A18B_SobAM2_CrePos_no0)
VP_A18B_SobAM2_CrePos_no0 <- merge(ttest_A18B_SobAM2_CrePos_no0, mean.sem_A18B_SobAM2_CrePos_no0, by = 'Lipid species')
# Calculate fold change
VP_A18B_SobAM2_CrePos_no0$"Log2FC" <- VP_A18B_SobAM2_CrePos_no0$'CRE+/SobAM2 avg' - VP_A18B_SobAM2_CrePos_no0$'CRE+/control avg'

#save df 
write_csv(VP_A18B_SobAM2_CrePos_no0, "VP_csv/A18B_SobAM2_CrePos_no0_Volcano_plot.csv")

## permFDP ####

VP_A18B_SobAM2_CrePos_no0 <- read_csv("VP_csv/A18B_SobAM2_CrePos_no0_Volcano_plot.csv")

pVals = VP_A18B_SobAM2_CrePos_no0$pvalue
threshold = 0.05
myDesign = c(1,1,1,1,1,1,1,1,2,2,2,2,2,2,2,2,2)
intOnly = VP_A18B_SobAM2_CrePos_no0 %>% select(-c("Lipid species","pvalue","CRE+/control avg","CRE+/control sem","CRE+/SobAM2 avg","CRE+/SobAM2 sem","Log2FC"))
intOnly <- mutate_all(intOnly, function(x) as.numeric(as.character(x))) 
corrThreshold_500 = permFDP::permFDP.adjust.threshold(pVals, threshold, myDesign, intOnly, 500)
corrThreshold_1000 = permFDP::permFDP.adjust.threshold(pVals, threshold, myDesign, intOnly, 1000)
corrThreshold_3000 = permFDP::permFDP.adjust.threshold(pVals, threshold, myDesign, intOnly, 3000)
corrThreshold_10000 = permFDP::permFDP.adjust.threshold(pVals, threshold, myDesign, intOnly, 10000)

print(corrThreshold_3000, digits = 20)
FDPthreshold[13,2] <- corrThreshold_3000
# corrThreshold_3000 = 0.011078706616200067331
## EnhancedVolcano ####

pdf("FinalVP/VP_A18B_SobAM2_CrePos_no0.pdf", width = 1.75, height = 1.24)

# Lipid species, p values, log2 FCs
toptable <- VP_A18B_SobAM2_CrePos_no0[, c(1,ncol(VP_A18B_SobAM2_CrePos_no0)-5,ncol(VP_A18B_SobAM2_CrePos_no0))]
lab = toptable$"Lipid species"
x = "Log2FC"
y = "pvalue"
pCutoff = corrThreshold_3000
FCcutoff = 0.5

keyvals.colour <- ifelse(
  toptable$Log2FC > 0.5 & toptable$pvalue < pCutoff, '#FF00FF',
  ifelse(toptable$Log2FC <(-0.5) & toptable$pvalue < pCutoff, '#00FFFF',
         '#808080'))
# Gray - #A8B6CC - #DAECCF
# Magenta - #F05039 - #F32A50
# Cyan - #0090FF - #0882E0

keyvals.colour[is.na(keyvals.colour)] <- '#808080'
names(keyvals.colour)[keyvals.colour == '#FF00FF'] <- 'High FC & sig'
names(keyvals.colour)[keyvals.colour == '#808080'] <- 'NS'
names(keyvals.colour)[keyvals.colour == '#00FFFF'] <- 'Low FC & sig'

EnhancedVolcano(toptable, lab, x, y, selectLab = rownames(toptable)[which(names(keyvals.colour) %in% c('High', 'Low'))],
                xlab = bquote(~Log[2] ~ "Fold Change"),legendPosition = 'none',
                ylab = bquote(~-Log[10] ~ italic(p) ~ Value), colCustom = keyvals.colour,
                title = NULL, subtitle = NULL, pCutoff = pCutoff, pCutoffCol = y, FCcutoff = FCcutoff,
                drawConnectors = FALSE, xlim = c(min(toptable[[x]], na.rm = TRUE) - 0.2, max(toptable[[x]], na.rm = TRUE) + 0.2),
                ylim = c(0, max(-log10(toptable[[y]]), na.rm = TRUE) + 0.5), colAlpha = 1,
                cutoffLineWidth = 0.1, pointSize = 0.5,axisLabSize = 5, captionLabSize = 3, borderWidth = 0.1,
                vlineWidth = 0.1, hlineWidth = 0.1, caption = NULL, legendLabSize = 3, legendIconSize =1
) + theme(axis.text.x = element_text(face="plain", color="black", size=5, angle=0),
          axis.text.y = element_text(face="plain", color="black", size=5, angle=0),
          axis.ticks.length=unit(.05, "cm"), axis.ticks=element_line(linewidth=0.2),
          axis.title.x = element_text(family = "sans", size = 5, margin=margin(1,0,0,0)), 
          axis.title.y = element_text(family = "sans", size = 5, margin=margin(0,1,0,0)),
          panel.grid.major = element_line(linewidth = 0.5), 
          panel.grid.minor = element_line(linewidth = 0.3),
          plot.margin = margin(4,4,4,4),
          legend.spacing.x = unit(0.5, "cm"),
          legend.spacing.y = unit(0.5, "cm")
)

dev.off()

# PLP W18 Spinal + SobAM2 ####

A18S_SobAM2_CrePos <- A6_18_S %>% filter(grepl('wk 18', ...1)) %>% filter(!grepl('-', CRE.and.Treatment))
A18S_SobAM2_CrePos_no0 <- A18S_SobAM2_CrePos %>% select(-"Sample.number", -"...1")

# For no 0 df, drop PC(40:3), ePC(38:3) and PS(32:1)
# https://stackoverflow.com/questions/72096913/remove-columns-that-have-one-zero-value
print(select(A18S_SobAM2_CrePos_no0, where(~any(. == 0))))
A18S_SobAM2_CrePos_no0 <- Filter(function(x) !any(x %in% 0), A18S_SobAM2_CrePos_no0) 
log_A18S_SobAM2_CrePos_no0 = log2(A18S_SobAM2_CrePos_no0[,-ncol(A18S_SobAM2_CrePos_no0)]) 
log_A18S_SobAM2_CrePos_no0 <- cbind(log_A18S_SobAM2_CrePos_no0, 'CRE.and.Treatment' = A18S_SobAM2_CrePos$CRE.and.Treatment)

# Fix column names & remove extra rows
ttest_A18S_SobAM2_CrePos_no0 <- transpose(log_A18S_SobAM2_CrePos_no0, keep.names = 'Lipid species')
ttest_A18S_SobAM2_CrePos_no0 <- ttest_A18S_SobAM2_CrePos_no0[-nrow(ttest_A18S_SobAM2_CrePos_no0),]

samples <- A18S_SobAM2_CrePos$Sample.number
names <- c('Lipid species', samples)
colnames(ttest_A18S_SobAM2_CrePos_no0) = c(names)

# Get p values from t test 
ttest_A18S_SobAM2_CrePos_no0$"pvalue" <- unlist(lapply(1:nrow(ttest_A18S_SobAM2_CrePos_no0), function(i) t.test((as.numeric(ttest_A18S_SobAM2_CrePos_no0[i,2:10])), as.numeric(ttest_A18S_SobAM2_CrePos_no0[i,11:19]))[c("p.value")]))

# calculate average values & sem for each species using CRE and treatment as factor
mean.sem_A18S_SobAM2_CrePos_no0 <- data.table(log_A18S_SobAM2_CrePos_no0)[, lapply(.SD, mean.sem), by=CRE.and.Treatment]
mean.sem_A18S_SobAM2_CrePos_no0 <- transpose(mean.sem_A18S_SobAM2_CrePos_no0[,2:(ncol(mean.sem_A18S_SobAM2_CrePos_no0))], keep.names = "Lipid species")
# Make sure the order of columns is correct, some df will have experimental first
CRE_factor <- c("CRE+/control avg","CRE+/control sem","CRE+/SobAM2 avg","CRE+/SobAM2 sem")
names <- c('Lipid species', CRE_factor)
colnames(mean.sem_A18S_SobAM2_CrePos_no0) = c(names)

# add means to main df
#VP_A18S_SobAM2_CrePos_no0 <- cbind(ttest_A18S_SobAM2_CrePos_no0, mean.sem_A18S_SobAM2_CrePos_no0)
VP_A18S_SobAM2_CrePos_no0 <- merge(ttest_A18S_SobAM2_CrePos_no0, mean.sem_A18S_SobAM2_CrePos_no0, by = 'Lipid species')
# Calculate fold change
VP_A18S_SobAM2_CrePos_no0$"Log2FC" <- VP_A18S_SobAM2_CrePos_no0$'CRE+/SobAM2 avg' - VP_A18S_SobAM2_CrePos_no0$'CRE+/control avg'

#save df 
write_csv(VP_A18S_SobAM2_CrePos_no0, "VP_csv/A18S_SobAM2_CrePos_no0_Volcano_plot.csv")

## permFDP ####

VP_A18S_SobAM2_CrePos_no0 <- read_csv("VP_csv/A18S_SobAM2_CrePos_no0_Volcano_plot.csv")

pVals = VP_A18S_SobAM2_CrePos_no0$pvalue
threshold = 0.05
myDesign = c(1,1,1,1,1,1,1,1,1,2,2,2,2,2,2,2,2,2)
intOnly = VP_A18S_SobAM2_CrePos_no0 %>% select(-c("Lipid species","pvalue","CRE+/control avg","CRE+/control sem","CRE+/SobAM2 avg","CRE+/SobAM2 sem","Log2FC"))
intOnly <- mutate_all(intOnly, function(x) as.numeric(as.character(x))) 
corrThreshold_500 = permFDP::permFDP.adjust.threshold(pVals, threshold, myDesign, intOnly, 500)
corrThreshold_1000 = permFDP::permFDP.adjust.threshold(pVals, threshold, myDesign, intOnly, 1000)
corrThreshold_3000 = permFDP::permFDP.adjust.threshold(pVals, threshold, myDesign, intOnly, 3000)
corrThreshold_10000 = permFDP::permFDP.adjust.threshold(pVals, threshold, myDesign, intOnly, 10000)

print(corrThreshold_3000, digits = 20)
FDPthreshold[14,2] <- corrThreshold_3000
# corrThreshold_3000 = 0.011718513795776697856
## EnhancedVolcano ####

pdf("FinalVP/VP_A18S_SobAM2_CrePos_no0.pdf", width = 1.75, height = 1.24)

# Lipid species, p values, log2 FCs
toptable <- VP_A18S_SobAM2_CrePos_no0[, c(1,ncol(VP_A18S_SobAM2_CrePos_no0)-5,ncol(VP_A18S_SobAM2_CrePos_no0))]
lab = toptable$"Lipid species"
x = "Log2FC"
y = "pvalue"
pCutoff = corrThreshold_3000
FCcutoff = 0.5

keyvals.colour <- ifelse(
  toptable$Log2FC > 0.5 & toptable$pvalue < pCutoff, '#FF00FF',
  ifelse(toptable$Log2FC <(-0.5) & toptable$pvalue < pCutoff, '#00FFFF',
         '#808080'))
# Gray - #A8B6CC - #DAECCF
# Magenta - #F05039 - #F32A50
# Cyan - #0090FF - #0882E0

keyvals.colour[is.na(keyvals.colour)] <- '#808080'
names(keyvals.colour)[keyvals.colour == '#FF00FF'] <- 'High FC & sig'
names(keyvals.colour)[keyvals.colour == '#808080'] <- 'NS'
names(keyvals.colour)[keyvals.colour == '#00FFFF'] <- 'Low FC & sig'

EnhancedVolcano(toptable, lab, x, y, selectLab = rownames(toptable)[which(names(keyvals.colour) %in% c('High', 'Low'))],
                xlab = bquote(~Log[2] ~ "Fold Change"),legendPosition = 'none',
                ylab = bquote(~-Log[10] ~ italic(p) ~ Value), colCustom = keyvals.colour,
                title = NULL, subtitle = NULL, pCutoff = pCutoff, pCutoffCol = y, FCcutoff = FCcutoff,
                drawConnectors = FALSE, xlim = c(min(toptable[[x]], na.rm = TRUE) - 0.2, max(toptable[[x]], na.rm = TRUE) + 0.2),
                ylim = c(0, max(-log10(toptable[[y]]), na.rm = TRUE) + 0.5), colAlpha = 1,
                cutoffLineWidth = 0.1, pointSize = 0.5,axisLabSize = 5, captionLabSize = 3, borderWidth = 0.1,
                vlineWidth = 0.1, hlineWidth = 0.1, caption = NULL, legendLabSize = 3, legendIconSize =1
) + theme(axis.text.x = element_text(face="plain", color="black", size=5, angle=0),
          axis.text.y = element_text(face="plain", color="black", size=5, angle=0),
          axis.ticks.length=unit(.05, "cm"), axis.ticks=element_line(linewidth=0.2),
          axis.title.x = element_text(family = "sans", size = 5, margin=margin(1,0,0,0)), 
          axis.title.y = element_text(family = "sans", size = 5, margin=margin(0,1,0,0)),
          panel.grid.major = element_line(linewidth = 0.5), 
          panel.grid.minor = element_line(linewidth = 0.3),
          plot.margin = margin(4,4,4,4),
          legend.spacing.x = unit(0.5, "cm"),
          legend.spacing.y = unit(0.5, "cm")
)

dev.off()

# PLP W18 Brain - SobAM2 ####

A18B_SobAM2_CreNeg <- A6_18_B %>% filter(grepl('wk 18', ...1)) %>% filter(grepl('-', CRE.and.Treatment))
A18B_SobAM2_CreNeg_no0 <- A18B_SobAM2_CreNeg %>% select(-"Sample.number", -"...1")

# For no 0 df, drop ePC(38:6) and PI(38:3)
# https://stackoverflow.com/questions/72096913/remove-columns-that-have-one-zero-value
print(select(A18B_SobAM2_CreNeg_no0, where(~any(. == 0))))
A18B_SobAM2_CreNeg_no0 <- Filter(function(x) !any(x %in% 0), A18B_SobAM2_CreNeg_no0) 
log_A18B_SobAM2_CreNeg_no0 = log2(A18B_SobAM2_CreNeg_no0[,-ncol(A18B_SobAM2_CreNeg_no0)]) 
log_A18B_SobAM2_CreNeg_no0 <- cbind(log_A18B_SobAM2_CreNeg_no0, 'CRE.and.Treatment' = A18B_SobAM2_CreNeg$CRE.and.Treatment)

# Fix column names & remove extra rows
ttest_A18B_SobAM2_CreNeg_no0 <- transpose(log_A18B_SobAM2_CreNeg_no0, keep.names = 'Lipid species')
ttest_A18B_SobAM2_CreNeg_no0 <- ttest_A18B_SobAM2_CreNeg_no0[-nrow(ttest_A18B_SobAM2_CreNeg_no0),]

samples <- A18B_SobAM2_CreNeg$Sample.number
names <- c('Lipid species', samples)
colnames(ttest_A18B_SobAM2_CreNeg_no0) = c(names)

# Get p values from t test 
ttest_A18B_SobAM2_CreNeg_no0$"pvalue" <- unlist(lapply(1:nrow(ttest_A18B_SobAM2_CreNeg_no0), function(i) t.test((as.numeric(ttest_A18B_SobAM2_CreNeg_no0[i,2:7])), as.numeric(ttest_A18B_SobAM2_CreNeg_no0[i,8:11]))[c("p.value")]))

# calculate average values & sem for each species using CRE and treatment as factor
mean.sem_A18B_SobAM2_CreNeg_no0 <- data.table(log_A18B_SobAM2_CreNeg_no0)[, lapply(.SD, mean.sem), by=CRE.and.Treatment]
mean.sem_A18B_SobAM2_CreNeg_no0 <- transpose(mean.sem_A18B_SobAM2_CreNeg_no0[,2:(ncol(mean.sem_A18B_SobAM2_CreNeg_no0))], keep.names = "Lipid species")
# Make sure the order of columns is correct, some df will have experimental first
CRE_factor <- c("CRE-/control avg","CRE-/control sem","CRE-/SobAM2 avg","CRE-/SobAM2 sem")
names <- c('Lipid species', CRE_factor)
colnames(mean.sem_A18B_SobAM2_CreNeg_no0) = c(names)

# add means to main df
#VP_A18B_SobAM2_CreNeg_no0 <- cbind(ttest_A18B_SobAM2_CreNeg_no0, mean.sem_A18B_SobAM2_CreNeg_no0)
VP_A18B_SobAM2_CreNeg_no0 <- merge(ttest_A18B_SobAM2_CreNeg_no0, mean.sem_A18B_SobAM2_CreNeg_no0, by = 'Lipid species')
# Calculate fold change
VP_A18B_SobAM2_CreNeg_no0$"Log2FC" <- VP_A18B_SobAM2_CreNeg_no0$'CRE-/SobAM2 avg' - VP_A18B_SobAM2_CreNeg_no0$'CRE-/control avg'

#save df 
write_csv(VP_A18B_SobAM2_CreNeg_no0, "VP_csv/A18B_SobAM2_CreNeg_no0_Volcano_plot.csv")

## permFDP ####

VP_A18B_SobAM2_CreNeg_no0 <- read_csv("VP_csv/A18B_SobAM2_CreNeg_no0_Volcano_plot.csv")

pVals = VP_A18B_SobAM2_CreNeg_no0$pvalue
threshold = 0.05
myDesign = c(1,1,1,1,1,1,2,2,2,2)
intOnly = VP_A18B_SobAM2_CreNeg_no0 %>% select(-c("Lipid species","pvalue","CRE-/control avg","CRE-/control sem","CRE-/SobAM2 avg","CRE-/SobAM2 sem","Log2FC"))
intOnly <- mutate_all(intOnly, function(x) as.numeric(as.character(x))) 
corrThreshold_500 = permFDP::permFDP.adjust.threshold(pVals, threshold, myDesign, intOnly, 500)
corrThreshold_1000 = permFDP::permFDP.adjust.threshold(pVals, threshold, myDesign, intOnly, 1000)
corrThreshold_3000 = permFDP::permFDP.adjust.threshold(pVals, threshold, myDesign, intOnly, 3000)
corrThreshold_10000 = permFDP::permFDP.adjust.threshold(pVals, threshold, myDesign, intOnly, 10000)

print(corrThreshold_3000, digits = 20)
FDPthreshold[15,2] <- corrThreshold_3000
# corrThreshold_3000 = 0.0013333255077575658157
## EnhancedVolcano ####

pdf("FinalVP/VP_A18B_SobAM2_CreNeg_no0.pdf", width = 1.75, height = 1.24)

# Lipid species, p values, log2 FCs
toptable <- VP_A18B_SobAM2_CreNeg_no0[, c(1,ncol(VP_A18B_SobAM2_CreNeg_no0)-5,ncol(VP_A18B_SobAM2_CreNeg_no0))]
lab = toptable$"Lipid species"
x = "Log2FC"
y = "pvalue"
pCutoff = corrThreshold_3000
FCcutoff = 0.5

keyvals.colour <- ifelse(
  toptable$Log2FC > 0.5 & toptable$pvalue < pCutoff, '#FF00FF',
  ifelse(toptable$Log2FC <(-0.5) & toptable$pvalue < pCutoff, '#00FFFF',
         '#808080'))
# Gray - #A8B6CC - #DAECCF
# Magenta - #F05039 - #F32A50
# Cyan - #0090FF - #0882E0

keyvals.colour[is.na(keyvals.colour)] <- '#808080'
names(keyvals.colour)[keyvals.colour == '#FF00FF'] <- 'High FC & sig'
names(keyvals.colour)[keyvals.colour == '#808080'] <- 'NS'
names(keyvals.colour)[keyvals.colour == '#00FFFF'] <- 'Low FC & sig'

EnhancedVolcano(toptable, lab, x, y, selectLab = rownames(toptable)[which(names(keyvals.colour) %in% c('High', 'Low'))],
                xlab = bquote(~Log[2] ~ "Fold Change"),legendPosition = 'none',
                ylab = bquote(~-Log[10] ~ italic(p) ~ Value), colCustom = keyvals.colour,
                title = NULL, subtitle = NULL, pCutoff = pCutoff, pCutoffCol = y, FCcutoff = FCcutoff,
                drawConnectors = FALSE, xlim = c(min(toptable[[x]], na.rm = TRUE) - 0.2, max(toptable[[x]], na.rm = TRUE) + 0.2),
                ylim = c(0, max(-log10(toptable[[y]]), na.rm = TRUE) + 0.5), colAlpha = 1,
                cutoffLineWidth = 0.1, pointSize = 0.5,axisLabSize = 5, captionLabSize = 3, borderWidth = 0.1,
                vlineWidth = 0.1, hlineWidth = 0.1, caption = NULL, legendLabSize = 3, legendIconSize =1
) + theme(axis.text.x = element_text(face="plain", color="black", size=5, angle=0),
          axis.text.y = element_text(face="plain", color="black", size=5, angle=0),
          axis.ticks.length=unit(.05, "cm"), axis.ticks=element_line(linewidth=0.2),
          axis.title.x = element_text(family = "sans", size = 5, margin=margin(1,0,0,0)), 
          axis.title.y = element_text(family = "sans", size = 5, margin=margin(0,1,0,0)),
          panel.grid.major = element_line(linewidth = 0.5), 
          panel.grid.minor = element_line(linewidth = 0.3),
          plot.margin = margin(4,4,4,4),
          legend.spacing.x = unit(0.5, "cm"),
          legend.spacing.y = unit(0.5, "cm")
)

dev.off()

# PLP W18 Spinal - SobAM2 ####

A18S_SobAM2_CreNeg <- A6_18_S %>% filter(grepl('wk 18', ...1)) %>% filter(grepl('-', CRE.and.Treatment))
A18S_SobAM2_CreNeg_no0 <- A18S_SobAM2_CreNeg %>% select(-"Sample.number", -"...1")

# For no 0 df, drop PS(32:1), ePS(38:4) and ePS(40:4)
# https://stackoverflow.com/questions/72096913/remove-columns-that-have-one-zero-value
print(select(A18S_SobAM2_CreNeg_no0, where(~any(. == 0))))
A18S_SobAM2_CreNeg_no0 <- Filter(function(x) !any(x %in% 0), A18S_SobAM2_CreNeg_no0) 
log_A18S_SobAM2_CreNeg_no0 = log2(A18S_SobAM2_CreNeg_no0[,-ncol(A18S_SobAM2_CreNeg_no0)]) 
log_A18S_SobAM2_CreNeg_no0 <- cbind(log_A18S_SobAM2_CreNeg_no0, 'CRE.and.Treatment' = A18S_SobAM2_CreNeg$CRE.and.Treatment)

# Fix column names & remove extra rows
ttest_A18S_SobAM2_CreNeg_no0 <- transpose(log_A18S_SobAM2_CreNeg_no0, keep.names = 'Lipid species')
ttest_A18S_SobAM2_CreNeg_no0 <- ttest_A18S_SobAM2_CreNeg_no0[-nrow(ttest_A18S_SobAM2_CreNeg_no0),]

samples <- A18S_SobAM2_CreNeg$Sample.number
names <- c('Lipid species', samples)
colnames(ttest_A18S_SobAM2_CreNeg_no0) = c(names)

# Get p values from t test 
ttest_A18S_SobAM2_CreNeg_no0$"pvalue" <- unlist(lapply(1:nrow(ttest_A18S_SobAM2_CreNeg_no0), function(i) t.test((as.numeric(ttest_A18S_SobAM2_CreNeg_no0[i,2:7])), as.numeric(ttest_A18S_SobAM2_CreNeg_no0[i,8:11]))[c("p.value")]))

# calculate average values & sem for each species using CRE and treatment as factor
mean.sem_A18S_SobAM2_CreNeg_no0 <- data.table(log_A18S_SobAM2_CreNeg_no0)[, lapply(.SD, mean.sem), by=CRE.and.Treatment]
mean.sem_A18S_SobAM2_CreNeg_no0 <- transpose(mean.sem_A18S_SobAM2_CreNeg_no0[,2:(ncol(mean.sem_A18S_SobAM2_CreNeg_no0))], keep.names = "Lipid species")
# Make sure the order of columns is correct, some df will have experimental first
CRE_factor <- c("CRE-/control avg","CRE-/control sem","CRE-/SobAM2 avg","CRE-/SobAM2 sem")
names <- c('Lipid species', CRE_factor)
colnames(mean.sem_A18S_SobAM2_CreNeg_no0) = c(names)

# add means to main df
#VP_A18S_SobAM2_CreNeg_no0 <- cbind(ttest_A18S_SobAM2_CreNeg_no0, mean.sem_A18S_SobAM2_CreNeg_no0)
VP_A18S_SobAM2_CreNeg_no0 <- merge(ttest_A18S_SobAM2_CreNeg_no0, mean.sem_A18S_SobAM2_CreNeg_no0, by = 'Lipid species')
# Calculate fold change
VP_A18S_SobAM2_CreNeg_no0$"Log2FC" <- VP_A18S_SobAM2_CreNeg_no0$'CRE-/SobAM2 avg' - VP_A18S_SobAM2_CreNeg_no0$'CRE-/control avg'

#save df 
write_csv(VP_A18S_SobAM2_CreNeg_no0, "VP_csv/A18S_SobAM2_CreNeg_no0_Volcano_plot.csv")

## permFDP ####

VP_A18S_SobAM2_CreNeg_no0 <- read_csv("VP_csv/A18S_SobAM2_CreNeg_no0_Volcano_plot.csv")

pVals = VP_A18S_SobAM2_CreNeg_no0$pvalue
threshold = 0.05
myDesign = c(1,1,1,1,1,1,2,2,2,2)
intOnly = VP_A18S_SobAM2_CreNeg_no0 %>% select(-c("Lipid species","pvalue","CRE-/control avg","CRE-/control sem","CRE-/SobAM2 avg","CRE-/SobAM2 sem","Log2FC"))
intOnly <- mutate_all(intOnly, function(x) as.numeric(as.character(x))) 
corrThreshold_500 = permFDP::permFDP.adjust.threshold(pVals, threshold, myDesign, intOnly, 500)
corrThreshold_1000 = permFDP::permFDP.adjust.threshold(pVals, threshold, myDesign, intOnly, 1000)
corrThreshold_3000 = permFDP::permFDP.adjust.threshold(pVals, threshold, myDesign, intOnly, 3000)
corrThreshold_10000 = permFDP::permFDP.adjust.threshold(pVals, threshold, myDesign, intOnly, 10000)

print(corrThreshold_3000, digits = 20)
FDPthreshold[16,2] <- corrThreshold_3000
# corrThreshold_3000 = 0.0075435595634583199609
## EnhancedVolcano ####

pdf("FinalVP/VP_A18S_SobAM2_CreNeg_no0.pdf", width = 1.75, height = 1.24)

# Lipid species, p values, log2 FCs
toptable <- VP_A18S_SobAM2_CreNeg_no0[, c(1,ncol(VP_A18S_SobAM2_CreNeg_no0)-5,ncol(VP_A18S_SobAM2_CreNeg_no0))]
lab = toptable$"Lipid species"
x = "Log2FC"
y = "pvalue"
pCutoff = corrThreshold_3000
FCcutoff = 0.5

keyvals.colour <- ifelse(
  toptable$Log2FC > 0.5 & toptable$pvalue < pCutoff, '#FF00FF',
  ifelse(toptable$Log2FC <(-0.5) & toptable$pvalue < pCutoff, '#00FFFF',
         '#808080'))
# Gray - #A8B6CC - #DAECCF
# Magenta - #F05039 - #F32A50
# Cyan - #0090FF - #0882E0

keyvals.colour[is.na(keyvals.colour)] <- '#808080'
names(keyvals.colour)[keyvals.colour == '#FF00FF'] <- 'High FC & sig'
names(keyvals.colour)[keyvals.colour == '#808080'] <- 'NS'
names(keyvals.colour)[keyvals.colour == '#00FFFF'] <- 'Low FC & sig'

EnhancedVolcano(toptable, lab, x, y, selectLab = rownames(toptable)[which(names(keyvals.colour) %in% c('High', 'Low'))],
                xlab = bquote(~Log[2] ~ "Fold Change"),legendPosition = 'none',
                ylab = bquote(~-Log[10] ~ italic(p) ~ Value), colCustom = keyvals.colour,
                title = NULL, subtitle = NULL, pCutoff = pCutoff, pCutoffCol = y, FCcutoff = FCcutoff,
                drawConnectors = FALSE, xlim = c(min(toptable[[x]], na.rm = TRUE) - 0.2, max(toptable[[x]], na.rm = TRUE) + 0.2),
                ylim = c(0, max(-log10(toptable[[y]]), na.rm = TRUE) + 0.5), colAlpha = 1,
                cutoffLineWidth = 0.1, pointSize = 0.5,axisLabSize = 5, captionLabSize = 3, borderWidth = 0.1,
                vlineWidth = 0.1, hlineWidth = 0.1, caption = NULL, legendLabSize = 3, legendIconSize =1
) + theme(axis.text.x = element_text(face="plain", color="black", size=5, angle=0),
          axis.text.y = element_text(face="plain", color="black", size=5, angle=0),
          axis.ticks.length=unit(.05, "cm"), axis.ticks=element_line(linewidth=0.2),
          axis.title.x = element_text(family = "sans", size = 5, margin=margin(1,0,0,0)), 
          axis.title.y = element_text(family = "sans", size = 5, margin=margin(0,1,0,0)),
          panel.grid.major = element_line(linewidth = 0.5), 
          panel.grid.minor = element_line(linewidth = 0.3),
          plot.margin = margin(4,4,4,4),
          legend.spacing.x = unit(0.5, "cm"),
          legend.spacing.y = unit(0.5, "cm")
)

dev.off()

write_csv(FDPthreshold,"FDP_Threshold.csv")
```
